# Atom-economic enantioselective photoenzymatic radical hydroalkylation via single-electron oxidation of carbanions

**DOI:** 10.1101/2024.12.08.627421

**Authors:** Jin Zhu, Qiaoyu Zhang, Tao Gu, Binbin Chen, Mingzhe Ma, Xiaoyu Wang, Xiao Liu, Mingjie Ma, Binju Wang, Yajie Wang

## Abstract

Established strategies for enantioselective hydroalkylation for C(*sp*^3^)–C(*sp*^3^) bond formation usually require pre-functionalized substrates as radical precursors in both transition-metal and photoenzymatic catalysis. Based on a sequential proton transfer/electron transfer (PT/ET) strategy, here we show a cooperative photoenzymatic system consisting of a flavin-dependent ‘ene’-reductase (ER) and an organophotoredox catalyst fluorescein (FI) to achieve atom-economic enantiodivergent hydroalkylation of electron-deficient C(*sp*^3^)–H with olefins. Mechanistic studies revealed a pathway for radical intermediate formation via excited-state FI^*^-induced single-electron oxidation of carbanions under alkaline conditions. The overall catalytic efficiency is enhanced by the electron transfer (ET) between FMN_ox_ and FI^-•^, while the stereoselectivity is controlled by ERs through enantioselective hydrogen atom transfer (HAT). We anticipate that this mode of photoenzymatic catalysis will inspire new pathways for generating free radical intermediates and foster innovative strategies for achieving photoenzymatic new-to-nature reactions.

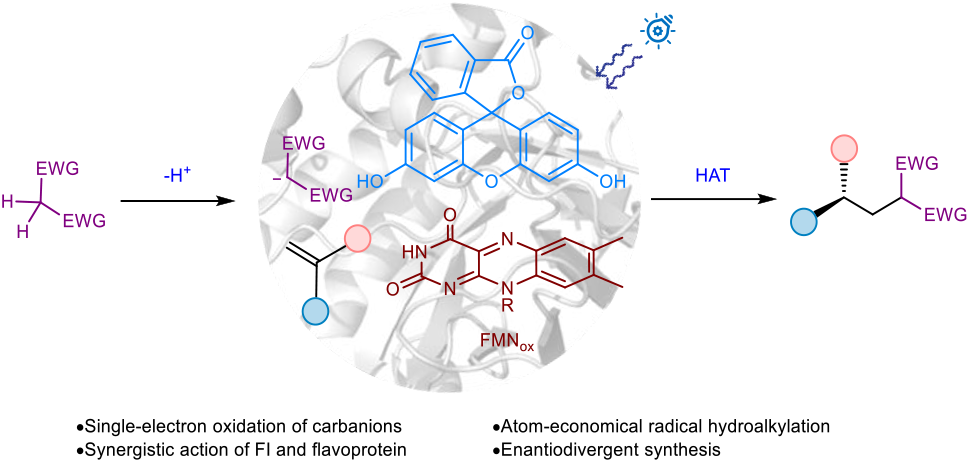

## Introduction

The formation of C(*sp*^3^)–C(*sp*^3^) bonds presents a significant synthetic challenge but is crucial in synthetic chemistry due to their fundamental role in constructing the carbon backbone of organic molecules^1-3^. Traditional chemocatalytic approaches include S_N_2-type substitution reactions of enolates with alkyl halides^4,5^ and transition metal-catalyzed cross-couplings^6,7^. The former strategy often requires strong bases and lacks monoalkylation control, while the latter is limited by β-hydrogen elimination in alkyl-metal species^1,2^. In contrast, radical hydroalkylation of cost-effective, readily available olefins has become an important strategy^8,9^. Common methods include single-electron reduction of halides^10-13^, proton-coupled electron transfer of diazo compounds^14^, and redox decarboxylation of carboxylic acids^15-17^ or esters^18-20^. However, these methods often require pre-functionalized substrates as radical precursors, impacting atom economy and necessitating pre-synthesis. While transition metal catalysis typically achieves enantioselectivity through metal hydride migratory insertion into olefins or radical trapping^12,21^, it struggles with remote stereocontrol at sites distant from the chiral environment^22,23^. State-of-the-art catalysis prioritizes maximum efficiency and sustainability through stringent adherence to step and atom economy, mild conditions, coupled with precise stereochemical control^24-26^, thus posing challenges for enantioselective radical hydroalkylation.

Recent advancements in photoenzymatic catalysis have shown remarkable enantioselectivity in the asymmetric hydrogen atom transfer (HAT) of prochiral radicals^27-39^, especially in remote stereocontrol^29,30^. Radicals are generated either through the single-electron reduction of an electron-donor-acceptor (EDA) complex (Fig. 1a and 1b)^27-34^ or via the single-electron oxidation of electron-rich precursors^35-39^. These methods have been successfully developed for selective hydroalkylation of halides^27-32^, NHPI esters^33^, diazo compounds^34^, and carboxylic acids^35^ with olefins, also largely relying on pre-functionalized substrates. In contrast, generating radicals through the homolytic cleavage of C–H bonds (Fig. 1c) in single-electron oxidation-initiated photoenzymatic catalysis shows excellent atom economy^36,37^. However, this approach has been restricted to the functionalization of electron-rich olefins and aromatics for lactonization^36^ and hydroarylation^37^. Therefore, developing atom-economic enantioselective hydroalkylation is of great scientific importance in both fields of transition metal catalysis and photoenzymatic catalysis.

**Fig. 1.**
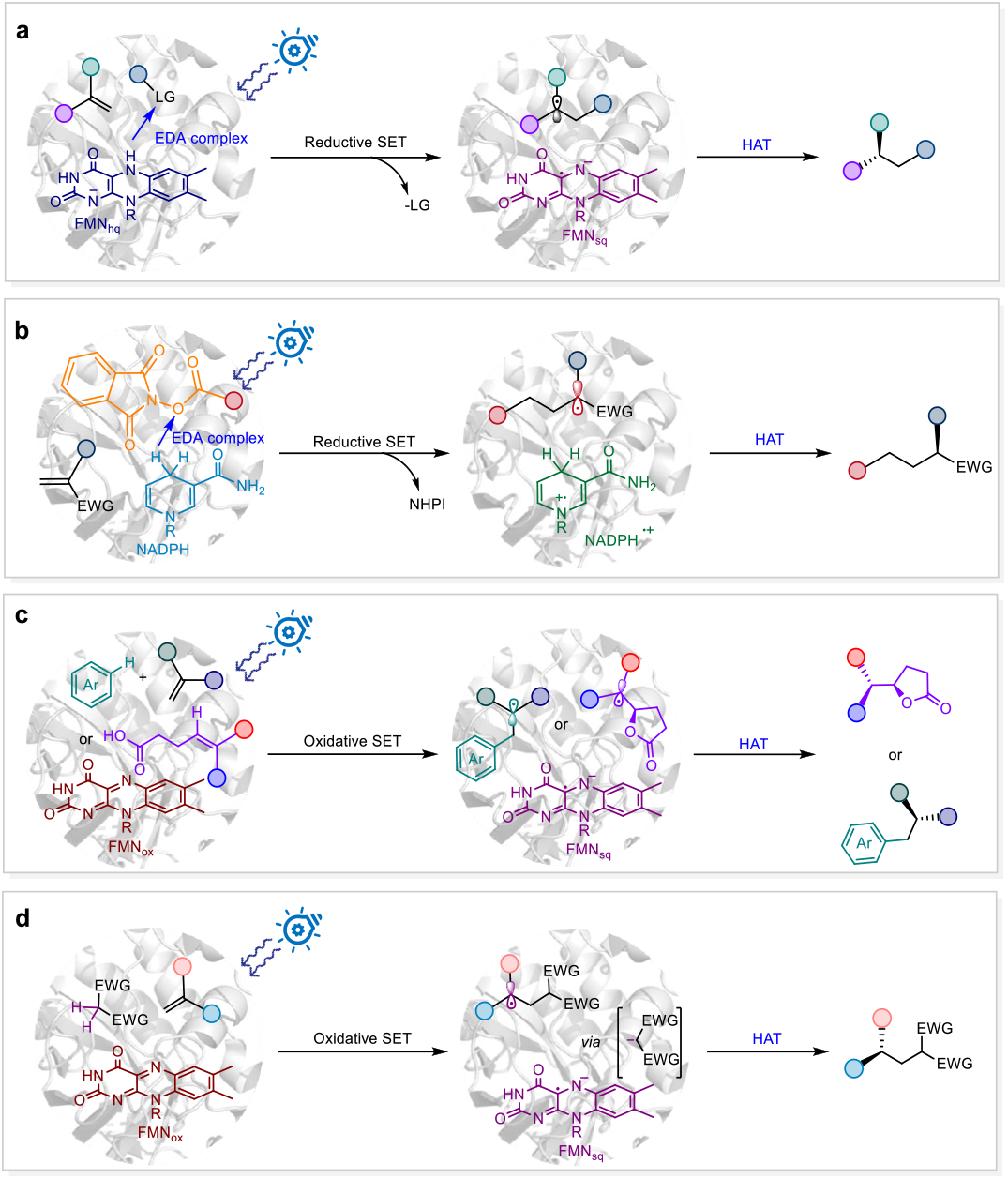
Different photoenzymatic modes for enantioselective hydrofunctionalization.

Given the critical role of cyano, carbonyl, and ester groups in late-stage functionalization^40^, the hydroalkylation of olefins with precursors containing these electron-withdrawing groups (EWG) is essential. Thus, electron-deficient C(*sp*^3^)–H bonds, activated by the adjacent EWGs, can serve as effective radical precursors for atom-economic hydroalkylation process. Although highly electron-deficient alkanes resist oxidation, they can be readily deprotonated to yield carbanions. Building on advances in synthetic chemistry, protic C(*sp*^3^)–H bonds can undergo radical activation through multiple pathways, including proton-coupled electron transfer (PCET), sequential PT/ET, and ET/PT^41^. This prompted our investigation of a PT/ET strategy, where electron-deficient substrates form carbanions via deprotonation, followed by single-electron oxidation to generate radical intermediates. Building on this hypothesis, we developed a cooperative photoenzymatic system comprising an ene-reductase (ER) and an organophotoredox fluorescein to achieve the enantioselective radical hydroalkylation of electron-deficient C(*sp*^3^)–H bonds with olefins. In this catalytic process, carbanions are generated under alkaline conditions and oxidized to radicals through an excited-state photocatalyst-induced single-electron transfer (SET). Stereoselectivity is achieved via an enantioselective HAT step of prochiral intermediates mediated by ER (Fig. 1d). This photoenzymatic system reveals a mechanism for radical formation that is rarely reported in photoredox catalysis, enabling an atom-economic and enantioselective radical hydroalkylation process.

## Results and Discussion

### Reaction development and engineering

While photoredox catalysis has successfully generated radicals from carbanions via single-electron oxidation^41-43^, this reactivity remains unexplored in biocatalysis systems. The key challenge lies in the high reactivity of carbanions, complicating control over single-electron oxidation versus competing pathways such as nucleophilic addition to electrophilic π-systems^44,45^. Highly electrophilic radicals can also quickly convert into carbanions through reductive radical polar crossover (RRPCO)^46,47^. To develop a photoenzymatic system for the enantioselective radical hydroalkylation of electron-deficient C(*sp*^3^)–H bonds with olefins, suitable enzymes must remain active in alkaline conditions necessary for carbanion formation. The excited ER-FMN_ox_^*^ must effectively oxidize carbanions into free radicals while minimizing undesirable pathways, such as carbanion nucleophilic addition, reductive quenching of FMN_ox_^*^ by the reaction buffer or nearby amino acids, and RRPCO (Supplementary Fig. 9).

We investigated the proposed reactions using malononitrile **1a** and α-methylstyrene **2a** as model substrates with a library of ERs at 1 mol% under 455 nm blue LED illumination. Most ERs showed no catalytic activity (Table 1, entry 1 and Supplementary Table 3), but MorB, OYE2, SYE1, SYE2, SYE3, and GluER achieved yields of 1.2% to 3%. Notably, SYE2 and GluER exhibited the best and opposite enantioselectivity for **3a**. GluER had (*S*)-selectivity with a 4:96 enantiomeric ratio (er) (Table 1, entry 2), while SYE2 showed (*R*)-selectivity with a 93.3:6.6 er (Table 1, entry 3). Thus, SYE2 and GluER were chosen for further investigation.

**Table 1.** Optimization of reaction conditions for photoenzymatic enantioselective C(*sp*^*3*^)-H functionalization.

| Entry | ER | Deviation from standard conditions<sup>a</sup> | Yield (%)<sup>b</sup> | er (%) (R:S)<sup>c</sup> |
| --- | --- | --- | --- | --- |
| 1 | ERs<sup>d</sup> | w/o FI | n.r. | n.a. |
| 2 | GluER | w/o FI | 1.3 | 4:96 |
| 3 | SYE2 | w/o FI | 2.6 | 93.3:6.6 |
| 4 | SYE2 | Tris buffer pH 7.4, w/o FI | n.r. | n.a. |
| 5 | SYE2 | Tris buffer pH 8, w/o FI | n.r. | n.a. |
| 6 | SYE2 | Tris buffer pH 10, w/o FI | 1.5 | 85:13 |
| 7 | SYE2 | w/o FI, 10 mol% FMN-Na | 2.6 | 93.3:6.6 |
| 8 | **SYE2** | **None** | **47.8** | **95.6:4.4** |
| 9 | **GluER** | **None** | **45.5** | **1.2:98.8** |
| 10 | SYE2 | Dark | n.r. | n.a. |
| 11 | SYE2 | Air | n.r. | n.a. |
| 12 | / | FMN-Na instead FI | n.r. | n.a. |
| 13 | / | None | n.r. | n.a. |
| 14 | / | 10 mol% FMN-Na + FI | 3 | racemate |

<sup>a</sup>Conditions: the total volume of standard reaction was 400 μL. <sup>b</sup>Yield was determined by gas chromatography. <sup>c</sup>er was determined by HPLC analysis on a chiral stationary phase, and the absolute configuration was established by theoretical calculations of electronic circular dichroism (ECD); see Supplementary Fig. 31 and Fig. 32. <sup>d</sup>LacER, KYE1, OYE1, OYE3, OPR1, SYE4, TOYE and YqjM were tested. n.r. No reaction; n.a. Not applicable.

To enhance the yield of **3a**, we tested various reaction conditions, including buffer systems, pH levels, and co-solvents (Supplementary Tables 4-5). Although these adjustments did not significantly improve the yield, we decided to proceed with Tris buffer at pH 9 and ethanol as a co-solvent. Notably, the reaction only occurs at pH 9 or above, indicating that strongly alkaline conditions are essential for initiating carbanion formation in the catalytic cycle (Table 1, entries 4-6).

Unlike other photoenzymatic systems^37,38^, adding extra FMN-Na did not enhance the yield of **3a** (Table 1, entry 7). This suggests that FMN_ox_^*^-induced single-electron oxidation of carbanions may be the rate-limiting step, indicating the need for photocatalysts with higher oxidative potential. We screened 18 transition metal and organic photocatalysts (Supplementary Tables 6-7) and identified fluorescein (FI-PC7), PC9, and PC12 as effective, increasing the yield of **3a** to 27%, 21%, and 19%, respectively, in both SYE2 and GluER-catalyzed reactions. The combination of 5 mol% FI with 1 mol% SYE2 or GluER gave the highest yields of **3a**, at 47.8% and 45.5%, respectively, without compromising enantioselectivity (Table 1, entries 8-9). Control experiments without light or in the presence of air did not produce the target product (Table 1, entries 10-11), supporting the visible-light-induced radical mechanism. Although neither FMN nor FI alone could catalyze the transformation (Table 1, entries 12-13), combining 10 mol% of both resulted in a 3% racemate of **3a** (Table 1, entry 14). These findings indicate that FI and FMN cooperate to catalyze the C(*sp*^*3*^)-H functionalization process, highlighting the essential role of photoenzymes in improving efficiency and controlling the stereochemistry of radical processes.

### Substrate scope investigations

We explored the substrate scope using SYE2 (ER1) and GluER to access both enantiomers of the products. While both enzymes showed excellent enantioselectivity for **3a**, this decreased significantly for most derivatives (Supplementary Fig. 33). To address this, we engineered GluER using site-saturated mutagenesis. Molecular docking of GluER’s crystal structure with **1a** and **2a** identified key residues in the substrate channel and FMN binding region, including T36, W66, H172, N175, Y177, T231, Q232, P239, R261, F269, and Y343 (see Supplementary Fig. 2). Site-saturated mutagenesis of these 11 sites led to GluER-Y343A (ER2), which showed improved enantioselectivity for both **3a** (up to R:S = 99.5:0.5) and other derivatives (Fig. 2).

**Fig. 2.**
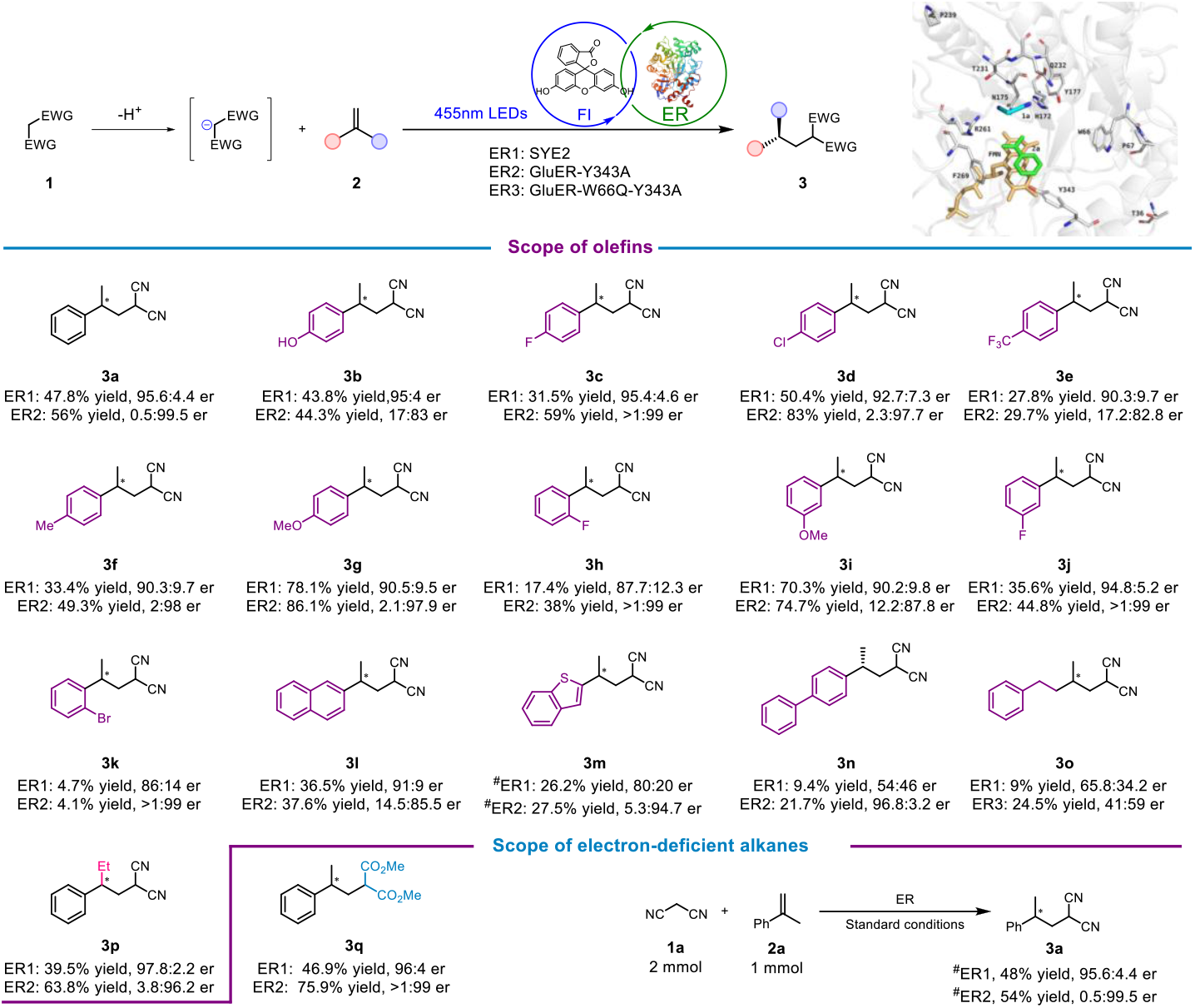
Substrate scope of enantioselective functionalization of electron-deficient C(*sp*^*3*^)-H bonds via hydroalkylation catalyzed by cooperative photoenzymatic system. Conditions: 1 (0.008 mmol), 2 (0.004 mmol), ER (1 mol%), FI (5 mol%), and EtOH (7.5 v/v%) in Tris buffer (50 mM, pH 9) were stirred at room temperature for 12 hours under N_2_ with 455 nm LEDs. The total reaction volume was 400 μL. Yields were determined by gas chromatography, and the er by HPLC on a chiral stationary phase. ^#^ Isolated yield was obtained by scaling up the reaction.

The photoenzymatic system enabled hydroalkylation of various substituted methyl styrenes with functional groups at the para-/meta-/ortho positions of the phenyl group (**3a-j**), producing stereodivergent products with moderate to good yields and up to >1:99 er. Although 1-bromo-2-(prop-1-en-2-yl)benzene (**3k**) had a relatively low yield, it exhibited good to excellent enantioselectivity. Additionally, 2-naphthyl (**3l**), benzothiophene (**3m**), and biphenyl (**3n**) demonstrated the compatibility of photoenzymatic catalysis with aromatic rings and heterocycle. Variations at the α position of styrene had minimal impact on outcomes, as seen with **3p**. However, replacing aromatic olefins with aliphatic olefins significantly reduced the yield of **3o**. Although the mutant GluER-W66Q-Y343A (ER3) improved the yield of **3o**, the enantioselectivity remained unsatisfactory. Beyond malononitrile **2a**, dimethyl malonate **2b** could also alkylate with **3q** with excellent enantioselectivity. Moreover, this photoenzymatic catalysis can be successfully scaled up to 1 mmol with minimal changes in yield and enantioselectivity.

### Mechanistic investigations

Based on the experimental results, we proposed a synergistic photoredox and biocatalytic mechanism for new-to-nature photoenzymatic C(*sp*^*3*^)-H functionalization (Fig. 3a and detailed in Supplementary Fig. 22). In this catalytic cycle, malononitrile **1a** is first deprotonated in alkaline solutions to form carbanion intermediate **Int. A**. The light-excited photocatalyst FI^*^ then oxidizes **Int. A** via SET, producing malononitrile free radical **Int. B** and fluorescein radical anion FI^-•^. **Int. B** subsequently attacks α-methylstyrene **2a**, forming a C-C bond and generating prochiral radical intermediate **Int. C**, alongside FMN_sq_ from FMN_ox_ capturing an electron from FI^-•^. Finally, **Int. C** undergoes ER-mediated enantioselective HAT, yielding enantioenriched **3a**, while FMNH^•^ is oxidized back to FMN_ox_, completing the catalytic cycle.

**Fig. 3.**
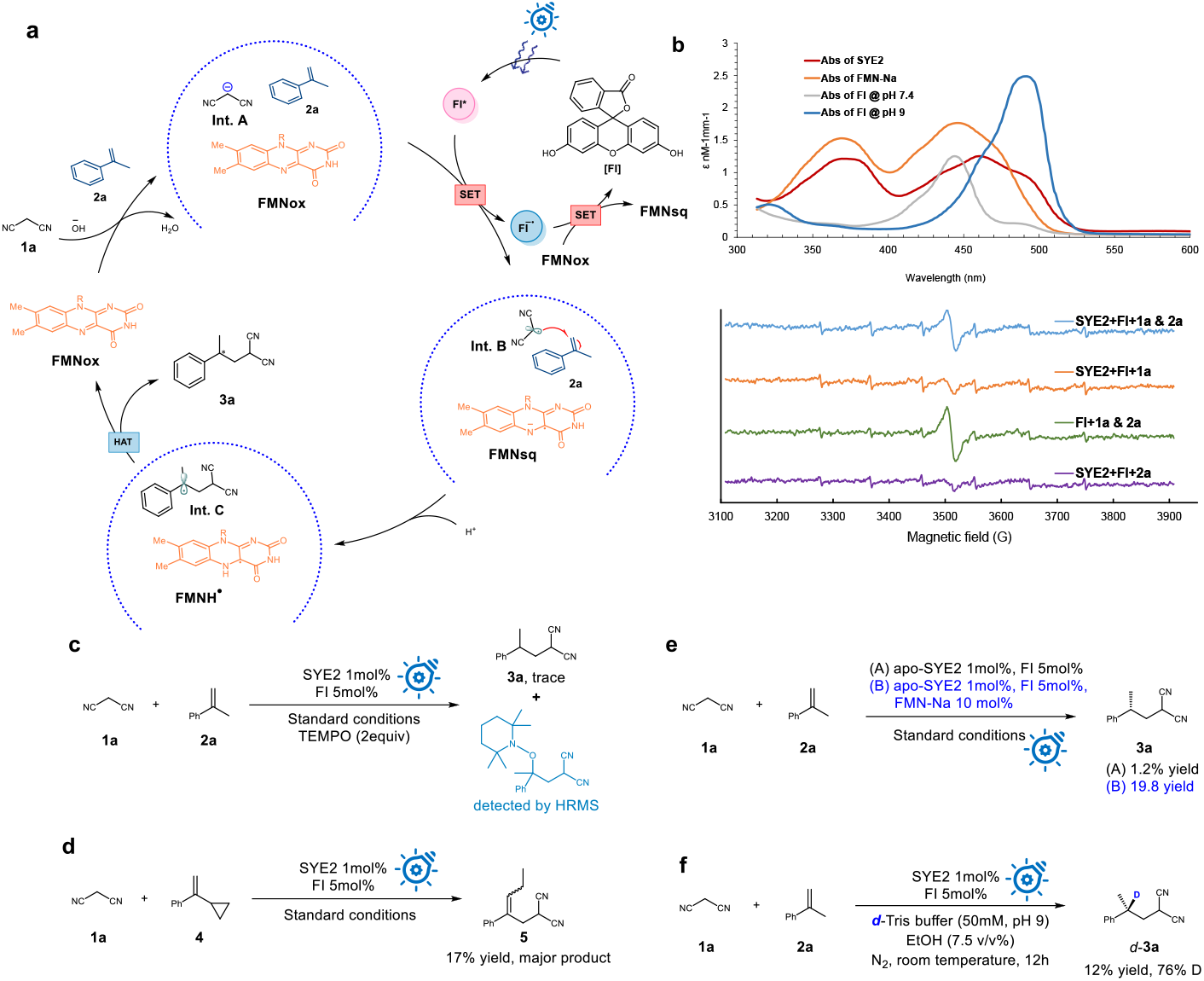
Mechanistic experiments and proposed catalytic cycle. **a**, Proposed catalytic cycle. **b**, UV-vis absorption (abs) spectra and room-temperature EPR experiments. **c**, TEMPO trapping experiment. **d**, Radical clock experiment. **e**, Photoenzymatic reactions catalysed by apo-flavoprotein. **f**, Deuterium-labelling experiment.

Several experiments were performed to support the proposed mechanism. The photophysical properties of SYE2 and FI were measured to identify visible-light-excited species. UV-vis absorption spectra show that SYE2 and FI absorb in the ranges of 400-520 nm and 420-530 nm, with maximum absorptions at 460 nm and 490 nm, respectively (Fig. 3b). This confirms that FI can be effectively excited by 455nm LED to initiate the catalytic cycle. Stern-Volmer luminescence quenching experiments also support the initial reductive quenching of excited FI^*^ by **1a** (**Int. A** → **Int. B**), as the fluorescence intensity of FI decreased upon adding **1a**, unlike with **2a** (Supplementary Fig. 10).

Several experiments were also performed to verify the generated radical intermediates. Firstly, the radical trapping experiment was performed by adding 2 equivalents of 2,2,6,6-tetramethylpiperidine-1-oxyl (TEMPO) to the reaction mixture under standard conditions. Only trace amounts of **3a** were detected, as the prochiral radical intermediate **Int. C** was trapped by TEMPO, confirmed by high-resolution mass spectrometry (HRMS) (Fig. 3c). Additionally, when α-cyclopropyl styrene **4** was subjected to the photoenzymatic system, the ring-opening product **5** was observed as the major product, supporting the involvement of **Int. C** (Fig. 3d). To further investigate the photo-oxidation step and clarify the resulting radical intermediates, electron paramagnetic resonance (EPR) experiments were conducted. With 5,5-dimethyl-1-pyrroline *N*-oxide (DMPO) as a radical trapping agent, slight changes in EPR signals were observed in the presence of **1a** compared to **2a** (Fig. 3b), indicating the formation of a potential **1a**-derived radical (**Int. B**), consistent with quenching experiments. The difference in EPR signals with and without **2a** suggests that the malononitrile radical **Int. B**, generated from the single-electron oxidation of the unstable carbanion **Int. A**, rapidly reacts with **2a** to form a more stable prochiral benzyl radical **Int. C** before being captured.^45^

Since apo-SYE2 (SYE2 without FMN) and FI alone did not catalyze the product formation, and adding extra FMN restored reactivity, we confirmed that the catalytic cycle requires the cooperative action of ER_FMN and FI (Fig. 3e and detailed in Supplementary Table 8 and Fig. 17). Finaly, deuterium-labelling experiments were conducted to investigate the final termination step (Fig. 3f). Using a deuterated buffer, the hydrogen at the stereogenic center was labelled at 76% D, suggesting that water may be involved in HAT steps.

### Computational studies

Extensive computational studies were performed to rationalize the mechanism proposed in Fig. 3a. Using hybrid cluster-continuum (HCC) model calculations, we found that **1a** undergoes facile deprotonation by OH− ions in aqueous solution, forming the anionic species **Int. A**. with a low energy barrier of 0.3 kcal/mol (Supplementary Figure 20). To ascertain that **Int. A**, rather than **1a**, serves as the reactant for the generation of the carbanion intermediates **Int. B**, density functional theory (DFT) calculations and time-dependent DFT (TD-DFT) calculations were conducted. The Gibbs reaction energy (ΔG) for the excited FI^*^-mediated oxidation of **1a** and **Int. A** were calculated to be +105.2 kcal/mol and −32.1 kcal/mol (Fig. 4a and Supplementary Fig. 15), respectively, supporting that photoinduced single-electron oxidation process occurs through carbanion intermediates **Int. A**.

**Fig. 4.**
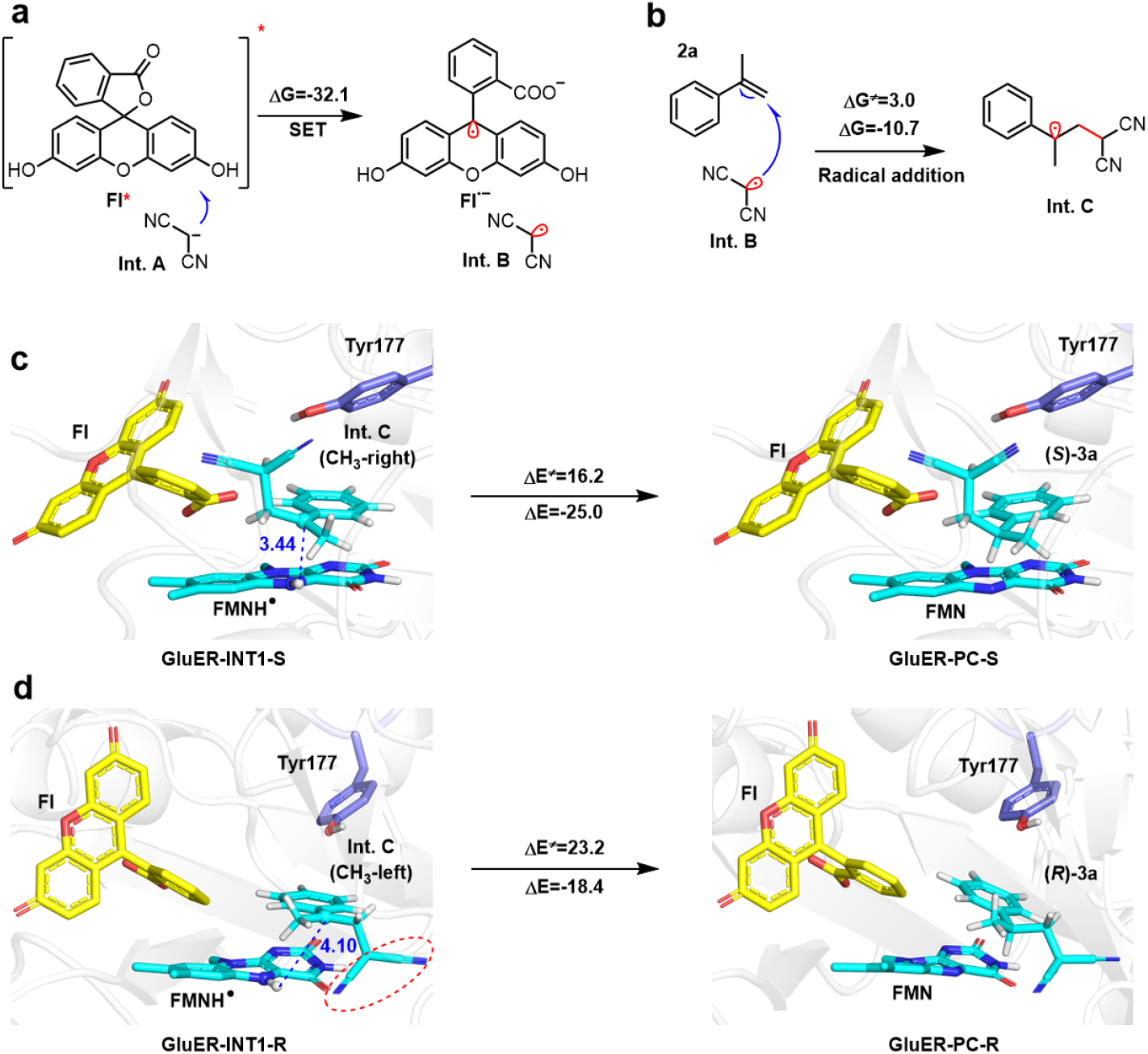
Mechanistic studies. **a**. SET from the excited FI^*^ to **Int. A. b**, The radical addition reaction between **Int. B** and **2a. c**, QM/MM-calculated energy of the HAT process from FMNH^•^ to the **Int. C** (CH_3_-right) result in the production of *S*-(**3a**). **d**, QM/MM-calculated energy of the HAT process from FMNH^•^ to the **Int. C** (CH_3_-left) result in the production of *R*-(**3a**).

Since EPR experiments with FI, **1a**, and **2a** revealed distinctive radical signals, suggesting that the formation of **Int. C** might be independent on ER participation. Quantum mechanics (QM) model calculations show that the radical addition between **Int. B** and **2a** has an energy barrier of 3.0 kcal/mol and an exothermicity of 10.7 kcal/mol (Fig. 4b), suggesting that formation of **Int. C** is quite facile. In addition, deuteration experiments and the pKa value of FMN_sq_ (∼8.5) suggested that the protonation of FMN_sq_ to form neutral FMNH^•^ is feasible at pH 9, with a Gibbs reaction energy (ΔG) of ∼0.7 kcal/mol. Starting from FMNH^•^, HAT from FMNH^•^ to **Int. C** is expected to yield the final product **3a**.

To elucidate the opposing enantioselectivity of GluER and SYE2, **Int. C** and FI were docked into the enzyme active sites. Molecular dynamics (MD) simulations indicate that FI was stabilized by Arg261 in GluER (Supplementary Fig. 23). **Int. C** adopts two distinct conformations (CH_3_-left and CH_3_-right) in GluER, corresponding to the *R* and *S* configurations, respectively. Moreover, the quantum mechanics /molecular mechanics (QM/MM)-calculated HAT barriers are 16.2 kcal/mol for the CH_3_-right and 23.2 kcal/mol for the CH_3_-left, aligning with the observed dominant product (*S*)-**3a** (Fig. 4c and Fig. 4d). This is likely due to steric repulsion between the bulky-CN substituent of **Int. C** and FMNH^•^ in the CH_3_-left conformation, which tends to enlarge the distance between the C3 atom of **Int. C** and the N5-H of FMNH^•^ (4.10 Å in CH_3_-left vs. 3.44 Å in CH_3_-right), thereby increasing the energy barrier for HAT in CH_3_-left conformation.

Unlike GluER, MD simulations reveal that the bulky-CN substituents in the CH_3_-left conformation of **Int. C** forms a stable hydrogen bond with Tyr180 in SYE2, which positions bulky-CN away from FMNH^•^ and facilitates the subsequent HAT process, that is favor of generating (*R*)-**3a** (Supplementary Fig. 26). Indeed, our calculations suggest that the HAT process in SYE2 involves a barrier of 17.2 kcal/mol in the CH_3_-left conformation and 22.0 kcal/mol in the CH_3_-right conformation, leading to the dominant formation of (*R*)-**3a** that agrees with experiments (Supplementary Fig. 26 and Fig. 27).

Furthermore, the newly formed FI^-•^ could reduce FMN_ox_ to FMN_sq_ and release the initial ground state FI. DFT calculations show that FI^-•^ has the lowest redox potential among all functional photocatalysts (PC1, PC2, PC5, FI, PC10, and PC12), making it highly prone to transfer an electron to FMN_ox_ to form FMN_sq_ (Supplementary Fig. 16). This suggests that ET between FMN_ox_ and FI^-•^ enhances the whole catalytic process, explaining why FI is the most effective among all tested photocatalysts.

## Conclusions

In summary, we have developed a synergistic photoenzymatic system for the enantioselective functionalization of electron-deficient C(*sp*^*3*^)-H bonds through radical intermediates generated by photoinduced single-electron oxidation of carbanions. This enantiodivergent strategy, demonstrated in 17 cases with enantiomeric ratios up to >1:99 and yields of up to 83%, offers potential for the one-step construction of dicyano and diester compounds with remote stereocenters. Detailed mechanistic experiments confirmed that the excited-state photocatalyst FI^*^ performs single-electron oxidation of carbanion rather than directly oxidizing the electron-deficient malononitrile **1a** to generate free radical intermediates. Unlike previous photoenzymatic systems, the ET between the additional photocatalyst FI and FMN_ox_ ensures catalytic cycle’s smooth operation. This novel mode of photoenzymatic catalysis is expected to inspire new strategies for forming free radical intermediates and enabling new-to-nature biotransformation.

## Methods

### General procedure for photoenzymatic reactions

Taking **1a** + **2a** → **3a** as an example, in a glovebox (O_2_ concentration <5 ppm),, solutions of ene-reductase (1 mol%), FI (10 μL, 2 mM stock in EtOH, 5 mol%), and substrates mix (20 μL, 400 mM **1a** and 200 mM **2a** in EtOH) were added to a 4 mL vial containing a magnetic stir bar and Tris buffer (50 mM, pH 9). The total volume of the reaction mixture was 400 μL and the final concentrations for EtOH was 7.5%. The vial was sealed with a screw cap, removed from the glove box, illuminated with blue LEDs, and stirred for 12 h with a cooling fan (see Supplementary Fig. 3 for reaction setup). For the reaction where the yield was determined by gas chromatography (GC), EtOAc (1.0 mL) and 10.0 μL of an internal standard stock (20 mg/mL of n-dodecane in EtOAc) were added to the reaction mixture and mixed thoroughly. For the reaction where the yield was determined by reverse phase HPLC, MeCN (1.0 mL) and 10.0 μL of an internal standard stock (16.8 mg/mL of 1,3,5-trimethoxybenzene in MeCN) were added to the reaction mixture and mixed thoroughly. The organic phase was separated and then analyzed by GC (or reverse phase HPLC), GC-MS and chiral HPLC. Product formation was confirmed by comparing retention times in GC/HPLC, as well as mass spectra and ^1^H-NMR spectra of the crude material with those of the reference. Yield determined by GC was relative to the n-dodecane internal standard, while yield determined by reverse phase HPLC was relative to the 1,3,5-trimethoxybenzene internal standard. Enantioselectivity was determined by chiral HPLC.

### System setup and MD simulations

The initial wild-type structure of GluER, was constructed based on the crystal structure (PDB code: 6O08). The SYE2 enzyme structure was modeled using AlphaFold2, and the position of FMN was confirmed by superimposing it with the GluER enzyme. Then, the **Int. C** and FI were docked into the active sites using AutoDock Vina tool. 100 ns MD simulations under the NPT ensemble. All MD simulations were performed with GPU version of Amber 18 package.

### QM/MM calculations for enzymatic reactions

A representative snapshot extracted from the clustering results in the MD trajectory was used for subsequent QM/MM calculations. All QM/MM calculations were performed using ChemShell combining Turbomole for the QM region and DL_POLY for the MM region. The electronic embedding scheme was used to account for the polarizing effect of the enzyme environment on the QM region.In all QM/MM geometry optimizations, the QM region was treated with the B3LYP functional using the double-zeta basis set def2-SVP for all atoms (labeled as B1). The energies were further corrected with the larger basis set def2-TZVP for all atoms (labeled as B2). Grimme’s D3BJ empirical dispersion was included in all calculations.

### DFT Calculations

All DFT calculations were performed with the Gaussian 16, Revision A.03 software. The initial structures were first optimized in conjunction with the SMD continuum solvation model at the B3LYP-D3/6-31G(d) level of theory. The solvation energies were further calculated at the M052X/6-31G* for all atoms. In particular, the solvent of water was used to simulate the reaction system.

### TDDFT Calculations

The initial structures were first optimized at B3LYP/6-31G(d) level. Then, the single-point TDDFT calculations at wB97XD/6-311+G(d,p) levels were performed to compute the three low-lying singlet excited states. All TDDFT calculations were performed with Gaussian 16, Revision A.03.

### Reporting summary

Further information on research design is available in the Nature Portfolio Reporting Summary linked to this article.

## Data availability

Data relating to the materials and methods, experimental procedures, mechanistic studies and computational calculations, HPLC spectra, and NMR spectra are available in the Supplementary Information or from the authors on reasonable request. The atomic coordinates of the optimized computational models are available in the Supplementary Data.

## Acknowledgements

This work was supported by the National Key R&D Program of China (2022YFA0912000), the Center of Synthetic Biology and Integrated Bioengineering (WU2022A006, WU2022A007, WU2023A009), the Research Center for Industries of the Future (WU2022C032). We thank Instrumentation and Service Center for Molecular Sciences (ISCMS) at Westlake University for the assistance in measurement and data interpretation. We thank Dr. Yinjuan Chen for the assistance with high resolution mass spectrometry (HRMS), Dr. Yuan Chen for the assistance with circular dichroism (CD), and Danyu Gu for the assistance with electron paramagnetic resonance (EPR).

## Author contributions

J.Z. and Y.W. conceived and designed the overall project. J.Z. performed all synthetic experiments and wet mechanism experiments. T.G., X.L., and M.-Z.M. created mutations. Q.Z., and X.W. carried out the computational studies and the calculation of absolute configuration (ECD) under the supervision of B.W. B.C. completed the calculation of redox potentials and Gibbs reaction energy (ΔG) for SET of Int. A. M.-J.M. performed the purification of some GluER variants. J.Z., B.W., and Y.W. wrote the paper with input of all authors.

## Competing interests

The authors declare no competing interests.

## Notes

### Competing Interest Statement

The authors have declared no competing interest.

### Summary of Updates

Abstract revised; Introduction revised; Figure 1 revised.

