## Supplementary FIgures and Figures for "Atom-economic enantioselective photoenzymatic radical hydroalkylation via single-electron oxidation of carbanions"

### 1. General Information

Unless otherwise noted, all chemicals and reagents for chemical reactions were obtained from commercial suppliers and used as received (Adamas, Aladdin, Bide Chemical, J&K Scientific and Macklin). All catalytic reactions were performed in a clear glass vial (4 mL) with magnetic stirring, illuminated with blue LED, and stirred for 12 h with a cooling fan or ROGER photo-reactor with a cooling pump (See **Supplementary Fig. 3** for reaction set-up). The alkenes are commercially available or synthesized by known methods. Flash column chromatography was performed using silica gel (200-300 mesh) from Aladdin. A blue LED (Kessil, PR160L-456nm, 40W) and photo-reactor (ROGER, 455nm LED, 15W) were used for all the photo-reactions. $^{1}H$, $^{13}C$ and $^{19}F$ NMR spectra were recorded on a Bruker BioSpin AVANCE NEO (600, 151 and 565 MHz, respectively) instrument, and are internally referenced to residual proton signals in $CDCl_3$ (7.26 ppm). $^{1}H$ NMR data are reported as follows: chemical shift (δ ppm), multiplicity (s = singlet, brs = broad singlet, d = doublet, t = triplet, q = quartet, m = multiplet, dd = doublet of doublet, dt = doublet of triplet, ddd = doublet of doublet of doublet), coupling constant (Hz), and integration. Data for $^{13}C$ NMR are reported in terms of chemical shift relative to $CDCl_3$ (77.16 ppm). High resolution mass spectra (HRMS) were obtained on Waters I-Class Plus/Syanpt-XS with electrospray ionization time-of-flight (ESI-TOF) and atmospheric pressure chemical ionization (APCI) detector. Enantiomeric ratio (er) was determined by chiral High Performance Liquid Chromatography (HPLC). UV detection was monitored at 210 nm at the same time. HPLC samples were dissolved in HPLC grade isopropanol (IPA) and hexane unless otherwise stated. HPLC analysis on chiral stationary phase was performed on an Agilent 1260 Infinity II LC System. Chiral IC, and IF columns were used to separate enantiomers (4.6 x 250 mm, 5 μm). Gas Chromatography Mass Spectrometry (GC-MS) analysis was performed on an Agilent 8890 system and HP-5 column was used.

*E. coli* DH5α cells, *E. coli* BL21 (DE3) stains were purchased from TransGen Biotech Co., Ltd. (Beijing, China). Plasmid pET22b, oligonucleotides for cloning and restriction enzymes were purchased from GenScript (Nanjing, China). Tryptone, yeast extract and FMN-Na were purchased from Aladdin Biochemical Technology Co., Ltd (Shanghai, China).

### 2. Cloning, Expression, Purification and Concentration Determination of Enzymes

For the expression of ene-reductases, an *E. coli* BL21(DE3) colony harboring pET28a or pET30a was inoculated in 5 mL LB medium containing 50 μg/mL kanamycin and grown overnight at 37 °C. This overnight culture was used to inoculate 500 mL of TB medium containing 50 μg/mL kanamycin, which was grown at 37 °C at 250 rpm until an $OD_{600}$ of ~ 0.6-0.8 was reached. Protein expression was subsequently induced with the addition of IPTG (for CsER, KYE1, OYE1, OYE2, OYE3, and YqjM: 0.1mM IPTG. For GluER, LacER, MorB, OPR1, OPR3, SYE1, SYE2, SYE3, SYE4, TOYE and YersER: 0.5mM IPTG). The induced culture was placed at 25 °C, 250 rpm, for 16 h for protein production. To obtain pure ERs, following expression, the cells were collected by centrifugation (4 °C, 6000 rpm for 15 min) and resuspended with lysis buffer (50 mM KPi buffer, pH = 7.4). FMN-Na (2 mg/mL) was added to the cell suspension. Then, the mixture was lysed by sonication and clarified by centrifugation (4 °C, 12000 rpm for 45 min). The protein was subsequently purified by affinity chromatography using a His-Trap column fitted to an ÄKTAxpress FPLC system (GE Health Life Sciences, Pittsburgh, PA). The purified protein was buffer exchanged against 50 mM Tris, pH 7.5 using an Amicon Ultra concentration tube with 30 kDa cut-off. After that, the proteins were stored in 20% glycerol as 100 μL aliquots at -80 °C.

The concentrations of ene-reductases were determined using the BCA protein assay kit.

DNA sequence of NAD(P)H: flavin oxidoreductase SYE2 form Shewanella oneidensis MR-1

ATGTTCAAGGCCTTTAAAGGCCACATGCTGGAAACCAAAAATCGTATTGTG

ATGGCACCGATGACCCGTAGCCGCAGTACCCAGCCGGGCGATATTCCGAAT

GAAATGATGGCAACCTATTATCGCCAGCGTGCCAGCGCCGGCCTGATTATTG

CCGAAGGCGCACCGGTGAGCGCCGTGGCACGTGGTTATAGCATGACCCCT

GGTATTTATACCCCGGCCCAGATTGAAGGTTGGAAAAAAGTTACCGAAGCA

GTTCATCAGGAAGGCGGCAAAATTTTTATTCAGCTGTGGCATGTGGGCCGT

CGTAGTCATAGCAGTGTTAGCGGTGCAGAACCGCTGGCCCCGAGCGCTATT

AAAATTCCGGATCAGGTTTTTGGTCCGCTGCCGGAAGGCGGTTTTGGTATG

ATTGAAACCCAGCAGCCGAAAGCAATGAGTGAACAGGATATTCAGGCAAC

CATTAGCGATTTTGTTCAGGCAGCCCAGAATGCAATGCTGGCCGGCTTTGAT

GGTGTTGAAGTGCATGCCGCCCACGGTTATCTGTTTGATACCTTTATGCGTC

TGGAAAGCAATCAGCGCCAGGATCGCTATGGTGGTAGTCAGGAAAATCGC

CTGCGTTTTCTGGTTGATACCCTGCAAGCACTGACCCAGACCATTGGTAGC

GGTCGCGTTGCCGTGCGCATTAGTCCGCATATTGGCGAAGGCTTTACCGGT

GATAATCCGGATATTATTCAGCTGACCCTGGCCCTGCTGCAAAAACTGCAAC

CGATGAATCTGGCCTATGTGCATTTTAGCGAAAATATTAGCCGCTATGTGGA

AGTGAGTGATGCCTTTCGCCAGCAGGTTCGTAGCGTTTATCAGCATCCGATT

ATGGTTGCCGGCAAACTGACCAAACAGAGCGCACAGCGTCTGCTGGATCA

GCATTATGCCGATTTTGTTGCCTTTGGTACACCGTTTGTGACCAATCCGGAT

CTGGTGGCACGTTTTGCCCATGATTGGCCGCTGACCGAATTTGATGCAGAT

GCACGTCTGACCCTGTATGGTGGCGGCGAAGCAGGCTATATTGATTATCCGG

TTTATCAGGCAAGTTTTTAA

Protein sequence of NAD(P)H: flavin oxidoreductase SYE2 form Shewanella oneidensis MR-1

MFKAFKGHMLETKNRIVMAPMTRSRSTQPGDIPNEMMATYYRQRASAGLIIA

EGAPVSAVARGYSMTPGIYTPAQIEGWKKVTEAVHQEGGKIFIQLWHVGRRSH

SSVSGAEPLAPSAIKIPDQVFGPLPEGGFGMIETQQPKAMSEQDIQATISDFVQ

AAQNAMLAGFDGVEVHAAHGYLFDTFMRLESNQRQDRYGGSQENRLRFLV

DTLQALTQTIGSGRVAVRISPHIGEGFTGDNPDIIQLTLALLQKLQPMNLAYVHF

SENISRYVEVSDAFRQQVRSVYQHPIMVAGKLTKQSAQRLLDQHYADFVAFG

TPFVTNPDLVARFAHDWPLTEFDADARLTLYGGGEAGYIDYPVYQASF

DNA sequence of 'Ene'-reductase from Gluconobacter Oxydans (GluER)

ATGGGCATGCACCACCATCACCACCACCCGACCCTTTTCGACCCCATCGAT

TTCGGACCTATCCACGCCAAGAATCGTATCGTCATGTCCCCCCTGACTCGCG

GTCGCGCTGACAAAGAGGCGGTTCCAACCCCCATTATGGCTGAATACTACG

CCCAACGCGCTTCGGCGGGTTTAATTATCACTGAAGCGACGGGGATTTCAC

GCGAAGGCTTAGGTTGGCCGTTTGCGCCGGGAATTTGGTCCGATGCACAGG

TTGAGGCGTGGAAACCTATCGTCGCGGGTGTCCATGCAAAGGGCGGCAAG

ATCGTATGTCAGCTTTGGCATATGGGCCGTATGGTACATTCTTCAGTTACAG

GGACGCAGCCCGTAAGCAGTTCCGCCACTACTGCTCCAGGTGAGGTTCAC

ACCTATGAGGGCAAGAAGCCCTTCGAACAAGCGCGTGCAATCGATGCTGC

AGACATCTCCCGCATCCTTAACGATTACGAAAATGCAGCACGTAATGCAATC

CGCGCGGGTTTCGATGGAGTGCAGATCCACGCAGCCAATGGCTACCTTATC

GATGAGTTTTTGCGTAACGGAACCAATCATCGCACCGATGAGTATGGGGGG

GTGCCGGAGAACCGTATTCGTTTCTTGAAAGAGGTAACAGAACGCGTCATC

GCGGCGATTGGCGCTGACCGTACGGGTGTGCGTCTGAGTCCAAACGGTGA

CACACAGGGTTGTATCGACAGTGCTCCCGAAACCGTTTTTGTTCCTGCCGC

AAAGCTTTTGCAAGATTTAGGGGTAGCGTGGCTTGAGCTGCGTGAACCTGG

TCCGAATGGTACGTTTGGAAAGACGGATCAACCAAAATTATCTCCACAAAT

CCGTAAGGTATTCCTTCGTCCATTGGTCTTAAATCAAGACTATACTTTTGAG

GCGGCACAGACGGCCCTGGCTGAGGGCAAGGCGGACGCTATTGCGTTTGG

CCGTAAGTTCATTTCAAATCCAGACTTGCCTGAGCGCTTTGCCCGTGGCATC

GCACTGCAACCAGACGATATGAAAACATGGTACTCCCAAGGCCCAGAGGG

TTACACAGACTATCCATCCGCAACTTCTGGGCCGAACTGA

Protein sequence of ‘Ene’-reductase from Gluconobacter Oxydans (GluER)

MGMHHHHHHPTLFDPIDFGPIHAKNRIVMSPLTRGRADKEAVPTPIMAEYYA

QRASAGLIITEATGISREGLGWPFAPGIWSDAQVEAWKPIVAGVHAKGGKIVC

QLWHMGRMVHSSVTGTQPVSSSATTAPGEVHTYEGKKPFEQARAIDAADISRI

LNDYENAARNAIRAGFDGVQIHAANGYLIDEFLRNGTNHRTDEYGGVPENRI

RFLKEVTERVIAAIGADRTGVRLSPNGDTQGCIDSAPETVFVPAAKLLQDLGV

AWLELREPGPNGTFGKTDQPKLSPQIRKVFLRPLVLNQDYTFEAAQTALAEGK

ADAIAFGRKFISNPDLPERFARGIALQPDDMKTWYSQGPEGYTDYPSATSGPN

DNA sequence of Caulobacter segnis Alkene Reductase (CsER)

ATGCCGAATTTGTTTGATCCGCTTCGTGTGGGAGACCTTAATTTGCCTAATC

GTGTCGTGATGGCACCCCTGACTCGCTTACGCGCTGGTCCTACACACATCC

CGAACGCTCTGATGGCAGAATACTATGGGCAGCGTGCAAGTGCAGGCTTAC

TTATTACGGAGGGAGTTCCAGTGGCGCCCCAAGGGGTTGGGTACGCTGGTG

TTCCTGGAATTTGGTCCAAGGAACAGACCGAAGGCTGGAAGCAAGTCACA

AAAGCTGTCCACGACAAGGGCGGCCGCATCTTCATGCAAATCTGGCACGTT

GGCCGCATCAGCGACCCGGAGTTGTTAAACGGAGAATTGCCGATTGCGCC

AAGTGCTATTGCCGCTAAAGGACATGTAAGCCTTTTACGCCCGCAACGCGA

TTACCCTACCCCCCGTGCACTTTCAACCGAGGAGGTGGCAGGAGTAGTCGA

AGCCTTCCGTCAGGGTGCTGAAAATGCTCAGGCAGCGGGCTTTGACGGGG

TCCAGTTGCATGGAGCTAACGGCTACCTTTTGGATCAGTTTTTACAGGACG

GGAGTAATCAACGCACGGATCAGTATGGGGGTTCGATTGAGAACCGTGCCC

GCCTGCTGTTGGAGGCAGCCGATGCGGCAATTAGCGTCTGGGGAGCAGAT

CGCGTAGGCGTGCACCTGGCCCCGCGTGCGGACTCCCATTCCATGGGTGAC

TCGAACCTGGCCGCGACCTTTGGTCACGTAGCGAAGGCATTAGGGGAGCG

CAAGATCGGTTTTGTCAGCGCACGCGAATATGAGGCCGCTGACTCTTTGGG

ACCGGATTTGAAGAAAGCATTCGGAGGAGTTTATATTGCGAATGAGAAATT

TGATCTTGCGTCTGCTAACGCCGCTATTGAGGCAGGCAAAGCGGATGCCAT

CGCGTTTGGCAAAGCCTACATCGCAAATCCCGATTTAGTGGAACGTCTTAA

AGCCGGGGCAGCTTTAAACACCCCGGATCCGGCGACTTTCTATGGCTTCGA

AAATGGTCCTCGCGGGTATACGGATTACCCTACCTTGGCTCAGGTCCGCGA

GCCCGCCCTCGAGCACCACCATCACCACCACTGA.

Protein sequence of Caulobacter segnis Alkene Reductase (CsER)

MPNLFDPLRVGDLNLPNRVVMAPLTRLRAGPTHIPNALMAEYYGQRASAGLL

ITEGVPVAPQGVGYAGVPGIWSKEQTEGWKQVTKAVHDKGGRIFMQIWHVG

RISDPELLNGELPIAPSAIAAKGHVSLLRPQRDYPTPRALSTEEVAGVVEAFRQ

GAENAQAAGFDGVQLHGANGYLLDQFLQDGSNQRTDQYGGSIENRARLLLE

AADAAISVWGADRVGVHLAPRADSHSMGDSNLAATFGHVAKALGERKIGFV

SAREYEAADSLGPDLKKAFGGVYIANEKFDLASANAAIEAGKADAIAFGKAY

IANPDLVERLKAGAALNTPDPATFYGFENGPRGYTDYPTLAQVREPALEHHHH

HH

DNA sequence of KYE1 from *Kluyveromyces lactis*

ATGAGCTTTATGAACTTTGAACCGAAACCGCTGGCGGATACCGATATTTTTA

AACCGATTAAAATTGGCAACACCGAACTGAAACATCGCGTGGTGATGCCG

GCGCTGACCCGCATGCGCGCGCTGCATCCGGGCAACGTGCCGAACCCGGA

TTGGGCGGTGGAATATTATCGCCAGCGCAGCCAGTATCCGGGCACCATGATT

ATTACCGAAGGCGCGTTTCCGAGCGCGCAGAGCGGCGGCTATGATAACGCG

CCGGGCGTGTGGAGCGAAGAACAGCTGGCGCAGTGGCGCAAAATTTTTAA

AGCGATTCATGATAACAAAAGCTTTGTGTGGGTGCAGCTGTGGGTGCTGGG

CCGCCAGGCGTTTGCGGATAACCTGGCGCGCGATGGCCTGCGCTATGATAG

CGCGAGCGATGAAGTGTATATGGGCGAAGATGAAAAAGAACGCGCGATTC

GCAGCAACAACCCGCAGCATGGCATTACCAAAGATGAAATTAAACAGTATA

TTCGCGATTATGTGGATGCGGCGAAAAAATGCATTGATGCGGGCGCGGATG

GCGTGGAAATTCATAGCGCGAACGGCTATCTGCTGAACCAGTTTCTGGATC

CGATTAGCAACAAACGCACCGATGAATATGGCGGCAGCATTGAAAACCGC

GCGCGCTTTGTGCTGGAAGTGGTGGATGCGGTGGTGGATGCGGTGGGCGC

GGAACGCACCAGCATTCGCTTTAGCCCGTATGGCGTGTTTGGCACCATGAG

CGGCGGCAGCGATCCGGTGCTGGTGGCGCAGTTTGCGTATGTGCTGGCGG

AACTGGAAAAACGCGCGAAAGCGGGCAAACGCCTGGCGTATGTGGATCTG

GTGGAACCGCGCGTGACCAGCCCGTTTCAGCCGGAATTTGAAGGCTGGTAT

AAAGGCGGCACCAACGAATTTGTGTATAGCGTGTGGAAAGGCAACGTGCT

GCGCGTGGGCAACTATGCGCTGGATCCGGATGCGGCGATTACCGATAGCAA

AAACCCGAACACCCTGATTGGCTATGGCCGCGCGTTTATTGCGAACCCGGA

TCTGGTGGAACGCCTGGAAAAAGGCCTGCCGCTGAACCAGTATGATCGCC

CGAGCTTTTATAAAATGAGCGCGGAAGGCTATATTGATTATCCGACCTATGA

AGAAGCGGTGGCGAAAGGCTATAAAAAA

Protein sequence of KYE1 from *Kluyveromyces lactis*

MSFMNFEPKPLADTDIFKPIKIGNTELKHRVVMPALTRMRALHPGNVPNPDW

AVEYYRQRSQYPGTMIITEGAFPSAQSGGYDNAPGVWSEEQLAQWRKIFKAI

HDNKSFVWVQLWVLGRQAFADNLARDGLRYDSASDEVYMGEDEKERAIRS

NNPQHGITKDEIKQYIRDYVDAAKKCIDAGADGVEIHSANGYLLNQFLDPISN

KRTDEYGGSIENRARFVLEVVDAVVDAVGAERTSIRFSPYGVFGTMSGGSDPV

LVAQFAYVLAELEKRAKAGKRLAYVDLVEPRVTSPFQPEFEGWYKGGTNEFV

YSVWKGNVLRVGNYALDPDAAITDSKNPNTLIGYGRAFIANPDLVERLEKGLP

LNQYDRPSFYKMSAEGYIDYPTYEEAVAKGYKK

DNA sequence of LacER

ATGTCGGGCTACCACTTCCTGAAGCCATTTACTTTTAAGCACCAAACTATAA

CGCTTAAAAACCGCATCGTCATTCCACCCATGACTACGAGACTTTCCTTCG

AGGATGGTACAGTTACCAGAGACGAGATTAGATACTATCAGCAACGGGCGG

GTGGCGTCGGTATGTTTATAACTGGTACTGCAAACGTCAACGCTCTTGGGA

AAGGCTTTGAAGGAGAATTATCGGTCGCGGACGATCGGTTCATTCCGGGCT

TGAGCAAATTGGCTGCAGCCATGAAGACTGGAGGGACCAAGGCTATTCTG

CAGATCTTTTCTGCCGGTCGCATGTCTAACAGCAAAATCTTGAGAGGGGAA

CAACCCGTGTCGGCATCAGCTGTGGCGGCGCCAAGAGCCGGGTACGAAAC

ACCTCGGGCGTTGACATCGGCTGAGATCGAAGCCACGATCCACGACTTTGG

GCAAGCTGTCCGTAGAGCAATCTTGGCGGGCTTCGATGGGATAGAATTGCA

TGGCGCCAATACATATTTGATCCAGCAATTTTATTCCCCTAACAGCAACCGG

CGTACCGATGAATGGGGAGGGGATAGAGACAAACGCATGCGGTTTCCCTTA

GCAGTGGTCCACGAGGCTGAAAAGGTGATAGCAACCATCGCGGATCGCCC

TTTCCTGCTTGGGTATCGGATCTCTCCTGAAGAACTGGAGCAACCGGGGAT

AACTCTTGATGACACTCTGGCCTTAATTGACGCTCTGAAACAAACGAAGAT

CGATTATTTACACGTTTCCCAGTCAGATGTCTGGAGAACTTCACTGCGTAAC

CCCGAGGATACAGCTATTATGAATGAGCAAATCCGTGATCATGTCGCAGGC

GCCTTCCCAGTTATCGTAGTAGGAGGAATCAAGACTCCAGCCGACGCTGAG

AAAGCTGCGGAATCTTTTGATTTAGTTGCTATAGGTCATGAAATGATACGTG

AGCCTCACTGGGTTCAAAAAGTACTGGACCACGACGAAAAGGCTATCCGT

TATCAAATTGCACCGGCGGACTTGGAAGAACTGGGCATCGCCCCTACGTTT

TTAGATTTTATCGAGAGCATCTCTGGTGGAGCCAAGGGGGTGCCCTTGACG

ACGGCGCAGTCGGTCACTAGCAGTAACGTCACACAAGACCTCGAGCACCA

CCATCACCACCACTGA

Protein sequence of LacER

MSGYHFLKPFTFKHQTITLKNRIVIPPMTTRLSFEDGTVTRDEIRYYQQRAGG

VGMFITGTANVNALGKGFEGELSVADDRFIPGLSKLAAAMKTGGTKAILQIFS

AGRMSNSKILRGEQPVSASAVAAPRAGYETPRALTSAEIEATIHDFGQAVRRAI

LAGFDGIELHGANTYLIQQFYSPNSNRRTDEWGGDRDKRMRFPLAVVHEAEK

VIATIADRPFLLGYRISPEELEQPGITLDDTLALIDALKQTKIDYLHVSQSDVWR

TSLRNPEDTAIMNEQIRDHVAGAFPVIVVGGIKTPADAEKAAESFDLVAIGHEM

IREPHWVQKVLDHDEKAIRYQIAPADLEELGIAPTFLDFIESISGGAKGVPLTTA

QSVTSSNVTQDLEHHHHHH

DNA sequence of Morphinone reductase (MorB) from Pseudomonas putida

ATGCCCGACACTTCTTTTTCGAATCCAGGACTTTTTACTCCTCTTCAGTTGG

GTAGTCTGTCTCTTCCAAATCGTGTCATAATGGCACCTTTAACCCGCTCACG

CACGCCAGATTCTGTACCTGGACGCCTTCAACAGATATACTATGGTCAACGC

GCCAGCGCCGGGTTAATCATCTCCGAAGCGACAAATATCAGTCCCACCGCT

CGGGGATACGTATACACGCCAGGCATTTGGACTGACGCTCAGGAGGCCGGT

TGGAAAGGTGTGGTCGAAGCTGTCCATGCTAAAGGGGGTCGTATAGCGTTG

CAGTTATGGCATGTCGGCCGGGTCTCTCATGAGCTGGTGCAGCCAGACGGC

CAACAACCCGTGGCACCATCCGCCTTAAAAGCCGAAGGGGCCGAGTGCTT

TGTCGAATTCGAGGATGGGACTGCTGGCCTGCACCCTACGTCAACTCCCAG

AGCCCTGGAGACAGATGAGATACCCGGTATTGTTGAAGATTACAGACAGGC

CGCGCAGCGTGCGAAGCGGGCCGGATTCGATATGGTAGAGGTCCACGCGG

CAAATGCTTGTCTTCCTAATCAGTTCTTGGCGACAGGAACCAATCGTCGCA

CAGACCAGTACGGTGGATCAATTGAGAACCGGGCTAGATTCCCATTAGAGG

TTGTCGATGCTGTAGCCGAGGTATTCGGGCCCGAAAGAGTGGGGATACGGC

TGACTCCTTTCCTGGAGTTATTTGGATTAACGGATGATGAACCCGAGGCAAT

GGCTTTTTACCTTGCGGGAGAATTAGACCGGCGTGGTTTAGCGTATTTACAC

TTTAATGAACCCGATTGGATAGGTGGGGACATCACGTACCCGGAAGGGTTT

CGTGAGCAAATGCGTCAACGGTTCAAGGGGGGGCTTATATATTGTGGAAAC

TACGACGCAGGTCGGGCCCAAGCCCGGCTTGACGACAATACAGCAGATGC

AGTGGCGTTTGGGCGTCCATTTATTGCCAACCCCGACTTGCCAGAACGTTT

CCGCTTAGGAGCAGCGCTGAACGAACCTGACCCCTCTACTTTTTACGGCGG

GGCAGAGGTGGGGTACACAGACTACCCGTTCCTGGACAACGGTCATGACC

GCCTGGGACTCGAGCACCACCATCACCACCACTGA

Protein sequence of Morphinone reductase (MorB) from Pseudomonas putida

MPDTSFSNPGLFTPLQLGSLSLPNRVIMAPLTRSRTPDSVPGRLQQIYYGQRAS

AGLIISEATNISPTARGYVYTPGIWTDAQEAGWKGVVEAVHAKGGRIALQLW

HVGRVSHELVQPDGQQPVAPSALKAEGAECFVEFEDGTAGLHPTSTPRALETD

EIPGIVEDYRQAAQRAKRAGFDMVEVHAANACLPNQFLATGTNRRTDQYGG

SIENRARFPLEVVDAVAEVFGPERVGIRLTPFLELFGLTDDEPEAMAFYLAGEL

DRRGLAYLHFNEPDWIGGDITYPEGFREQMRQRFKGGLIYCGNYDAGRAQA

RLDDNTADAVAFGRPFIANPDLPERFRLGAALNEPDPSTFYGGAEVGYTDYPF

LDNGHDRLGHHHHHH

DNA sequence of Old Yellow Enzyme 1 (OYE1)

ATGAGCTTTGTCAAGGACTTCAAGCCACAAGCACTTGGTGATACAAACCTT

TTTAAACCAATTAAGATTGGCAACAACGAACTGTTACATCGTGCAGTTATTC

CCCCTCTTACACGTATGCGTGCTTTGCACCCAGGTAATATTCCCAACCGTGA

TTGGGCGGTAGAATACTATACTCAACGCGCCCAGCGCCCGGGCACGATGAT

CATTACGGAAGGGGCATTTATCAGCCCCCAAGCTGGCGGGTATGATAACGC

ACCCGGTGTTTGGTCCGAAGAACAGATGGTGGAGTGGACGAAAATTTTCA

ATGCAATCCACGAAAAAAAATCGTTTGTCTGGGTACAGTTATGGGTCCTGG

GTTGGGCGGCATTTCCTGACAATTTGGCCCGCGACGGGTTACGCTACGATA

GCGCATCTGACAACGTCTTTATGGATGCCGAGCAAGAGGCTAAGGCGAAG

AAAGCCAACAACCCTCAGCACTCCTTGACAAAAGATGAAATTAAACAGTA

CATTAAGGAGTACGTGCAGGCTGCAAAAAACAGCATTGCTGCAGGTGCAG

ACGGTGTAGAGATTCACTCGGCTAATGGGTACCTGCTTAATCAATTTTTAGA

TCCTCACTCGAACACACGTACCGACGAATATGGGGGATCTATTGAGAATCG

CGCTCGTTTTACGTTGGAAGTGGTAGATGCATTGGTCGAGGCGATCGGGCA

CGAAAAAGTCGGATTACGTTTATCTCCTTATGGCGTGTTTAATTCAATGTCA

GGCGGAGCGGAGACTGGAATTGTCGCACAGTACGCTTACGTGGCGGGAGA

GCTGGAAAAACGTGCAAAGGCTGGAAAACGCCTGGCATTTGTACATTTAG

TGGAACCGCGTGTGACGAATCCTTTTCTTACTGAGGGGGAGGGCGAGTAC

GAAGGAGGGAGCAATGATTTCGTGTATAGTATTTGGAAGGGTCCTGTTATTC

GCGCTGGCAATTTCGCATTGCACCCAGAGGTGGTGCGTGAAGAAGTGAAA

GATAAACGCACGTTGATCGGCTACGGCCGTTTCTTTATTAGTAACCCAGACT

TGGTGGACCGTTTAGAGAAAGGTCTTCCCTTGAACAAATATGATCGTGACA

CCTTTTATCAGATGTCGGCGCACGGATACATTGATTACCCGACCTATGAAGA

GGCTTTAAAACTTGGTTGGGATAAGAAG

Protein sequence of Old Yellow Enzyme 1 (OYE1)

MSFVKDFKPQALGDTNLFKPIKIGNNELLHRAVIPPLTRMRALHPGNIPNRDW

AVEYYTQRAQRPGTMIITEGAFISPQAGGYDNAPGVWSEEQMVEWTKIFNAI

HEKKSFVWVQLWVLGWAAFPDNLARDGLRYDSASDNVFMDAEQEAKAKK

ANNPQHSLTKDEIKQYIKEYVQAAKNSIAAGADGVEIHSANGYLLNQFLDPHS

NTRTDEYGGSIENRARFTLEVVDALVEAIGHEKVGLRLSPYGVFNSMSGGAET

GIVAQYAYVAGELEKRAKAGKRLAFVHLVEPRVTNPFLTEGEGEYEGGSNDFV

YSIWKGPVIRAGNFALHPEVVREEVKDKRTLIGYGRFFISNPDLVDRLEKGLPL

NKYDRDTFYQMSAHGYIDYPTYEEALKLGWDKK

DNA sequence of OYE2

ATGCCCTTCGTGAAAGACTTCAAACCTCAAGCCCTGGGCGATACTAATTTAT

TTAAGCCAATTAAAATTGGAAACAATGAGTTGTTACACCGCGCTGTAATTCC

ACCCTTAACCCGCATGCGCGCCCAACATCCAGGGAACATCCCTAATCGCGA

TTGGGCAGTCGAGTACTATGCTCAGCGTGCTCAGCGTCCGGGTACCCTTAT

CATCACGGAAGGAACGTTTCCGTCGCCGCAATCGGGAGGGTATGACAACG

CTCCCGGTATCTGGTCGGAAGAACAGATTAAAGAATGGACCAAAATCTTTA

AAGCAATTCATGAGAATAAATCTTTCGCCTGGGTCCAACTTTGGGTCCTGG

GCTGGGCAGCCTTCCCTGACACATTGGCGCGTGACGGGCTTCGTTATGATA

GTGCTTCGGATAACGTGTATATGAATGCTGAACAAGAAGAAAAGGCAAAA

AAAGCAAACAATCCACAGCATTCGATTACTAAAGACGAGATTAAGCAGTAT

GTTAAGGAATACGTACAAGCAGCAAAGAATTCTATTGCCGCAGGGGCGGA

CGGGGTAGAAATCCACTCTGCTAATGGGTACTTGCTTAACCAGTTCCTGGA

CCCGCATTCAAACAACCGCACTGATGAGTACGGAGGGTCCATCGAAAATCG

TGCACGTTTTACTTTAGAGGTCGTAGATGCTGTAGTCGACGCGATTGGCCCT

GAGAAGGTAGGTTTGCGTTTAAGTCCTTATGGCGTGTTCAATTCAATGTCAG

GGGGCGCTGAAACAGGTATCGTCGCGCAGTACGCATACGTCTTGGGAGAG

CTGGAGCGTCGTGCTAAGGCTGGCAAGCGTTTAGCTTTTGTGCATTTAGTT

GAACCGCGCGTGACAAACCCCTTCTTGACGGAAGGCGAAGGAGAGTATAA

CGGAGGATCGAATAAATTTGCGTATTCCATTGGAAGGGCCCGATCATTCGTG

CCGGTAACTTTGCCTTACATCCCGAAGTTGTTCGCGAGGAAGTAAAAGACC

CACGTACCTTGATCGGGTATGGCCGTTTCTTTATTTCAAACCCCGACTTGGT

GGATCGCCTTGAAAAAGGTCTTCCCTTGAATAAGTATGACCGTGATACGTTC

TACAAAATGTCAGCCGAAGGTTACATCGACTACCCCACCTACGAAGAGGCT

TTGAAACTTGGTTGGGACAAGAACCACCACCATCACCACCACT

Protein sequence of OYE2

MPFVKDFKPQALGDTNLFKPIKIGNNELLHRAVIPPLTRMRAQHPGNIPNRDW

AVEYYAQRAQRPGTLIITEGTFPSPQSGGYDNAPGIWSEEQIKEWTKIFKAIHE

NKSFAWVQLWVLGWAAFPDTLARDGLRYDSASDNVYMNAEQEEKAKKANN

PQHSITKDEIKQYVKEYVQAAKNSIAAGADGVEIHSANGYLLNQFLDPHSNN

RTDEYGGSIENRARFTLEVVDAVVDAIGPEKVGLRLSPYGVFNSMSGGAETGI

VAQYAYVLGELERRAKAGKRLAFVHLVEPRVTNPFLTEGEGEYNGGSNKFAY

SIGRARSFVPVTLPYIPKLFARKKTHVPSGMAVSLFQTPTWWIALKKVFPISMT

VIRSTKCQPKVTSTTPPTKRLNLVGTRTTTITTT

DNA sequence of OYE3

ATGCCTTTCGTGAAGGGGTTCGAGCCGATCTCTCTGCGCGATACAAACTTG

TTCGAACCTATCAAGATCGGTAATACCCAATTAGCGCACCGTGCTGTAATGC

CCCCATTGACCCGTATGCGCGCGACGCACCCCGGTAATATTCCCAATAAAG

AGTGGGCGGCGGTCTACTATGGACAACGTGCGCAACGTCCTGGGACGATG

ATTATTACTGAAGGTACTTTTATTTCACCCCAAGCCGGCGGGTATGATAACG

CACCTGGAATCTGGAGTGATGAGCAAGTGGCTGAGTGGAAAAACATCTTC

CTTGCAATCCATGACTGCCAATCTTTTGCTTGGGTTCAGCTGTGGAGCTTAG

GATGGGCATCATTTCCAGATGTATTAGCCCGTGACGGTCTTCGTTATGATTGT

GCTTCAGACCGCGTGTATATGAACGCAACATTACAAGAAAAAGCCAAAGA

CGCAAACAACCTTGAGCACTCGCTGACTAAAGACGACATTAAGCAATACAT

TAAGGACTATATTCATGCAGCGAAGAATAGTATCGCTGCCGGAGCCGATGG

CGTGGAAATTCACAGCGCTAATGGCTACCTGCTGAACCAATTCTTAGACCC

CCATTCTAATAAACGCACTGATGAGTACGGGGGAACGATTGAGAACCGTGC

TCGTTTCACATTAGAGGTAGTCGATGCTTTGATTGAGACGATCGGCCCGGA

GCGCGTAGGCCTGCGTTTGTCCCCCTATGGGACCTTCAACAGTATGAGTGG

AGGGGCAGAGCCTGGAATCATTGCACAGTATAGCTATGTCCTGGGTGAATT

GGAAAAGCGTGCAAAAGCGGGCAAACGTCTTGCCTTCGTTCATCTGGTGG

AGCCGCGTGTTACCGACCCCTCCTTAGTTGAGGGAGAGGGAGAGTACAGT

GAGGGTACGAATGACTTCGCCTACAGCATCTGGAAGGGGCCCATCATTCGC

GCTGGCAATTACGCCTTGCACCCAGAAGTCGTCCGCGAGCAGGTAAAGGA

TCCACGTACACTGATCGGCTATGGGCGCTTCTTCATTTCAAATCCAGACTTG

GTCTACCGTCTGGAAGAGGGATTACCATTAAATAAATATGACCGCTCCACAT

TTTATACCATGTCGGCTGAGGGGTATACAGACTACCCCACCTATGAGGAAGC

AGTGGATCTTGGTTGGAACAAGAATCACCACCATCACCACCACTGA

Protein sequence of OYE3

MPFVKGFEPISLRDTNLFEPIKIGNTQLAHRAVMPPLTRMRATHPGNIPNKEWA

AVYYGQRAQRPGTMIITEGTFISPQAGGYDNAPGIWSDEQVAEWKNIFLAIHD

CQSFAWVQLWSLGWASFPDVLARDGLRYDCASDRVYMNATLQEKAKDANN

LEHSLTKDDIKQYIKDYIHAAKNSIAAGADGVEIHSANGYLLNQFLDPHSNKR

TDEYGGTIENRARFTLEVVDALIETIGPERVGLRLSPYGTFNSMSGGAEPGIIA

QYSYVLGELEKRAKAGKRLAFVHLVEPRVTDPSLVEGEGEYSEGTNDFAYSI

WKGPIIRAGNYALHPEVVREQVKDPRTLIGYGRFFISNPDLVYRLEEGLPLNKY

DRSTFYTMSAEGYTDYPTYEEAVDLGWNKNHHHHHH

DNA sequence of 12-Oxophytodienoate reductase 1 (OPR1) from Solanum lycopersicum

ATGGAGAATAAAGTTGTGGAGGAGAAACAAGTCGATAAAATCCCCTTGATG

TCACCGTGCAAGATGGGAAAGTTTGAACTTTGCCACCGTGTTGTCCTGGCT

CCGCTGACACGGCAACGCTCCTACGGGTATATACCGCAGCCCCATGCAATC

TTACATTACTCTCAGCGTTCAACCAACGGAGGGCTGTTGATAGGTGAAGCA

ACAGTCATTAGCGAAACGGGTATAGGCTATAAGGACGTACCCGGCATCTGG

ACTAAAGAACAAGTTGAGGCATGAAACCCATAGTAGACGCCGTACATGCA

AAGGGAGGGATCTTCTTCTGCCAGATATGGCATGTGGGTAGAGTGAGTAAT

AAGGACTTCCAACCTAACGGTGAGGATCCCATTAGCTGTACCGATCGGGGA

TTAACGCCACAGATACGGTCAAATGGTATTGACATAGCTCATTTTACAAGAC

CTAGACGTCTTACCACGGACGAGATCCCACAAATTGTCAACGAATTCCGCG

TGGCGGCTAGAAATGCCATCGAAGCAGGATTCGACGGCGTAGAGATACAC

GGAGCACACGGTTATCTGATAGACCAGTTCATGAAAGACCAAGTTAATGAC

CGGTCCGATAAGTATGGAGGATCTCTGGAAAACCGGTGTCGGTTCGCCTTG

GAGATTGTTGAAGCCGTCGCTAATGAAATCGGAAGCGACCGTGTCGGAATA

CGCATTAGTCCATTCGCGCACTACAATGAGGCAGGGGATACCAATCCCACT

GCGCTGGGTTTATATATGGTGGAGAGCCTGAATAAATACGATTTAGCATATTG

TCATGTAGTGGAACCTCGCATGAAAACTGCTTGGGAAAAAATTGAATGCAC

TGAGAGTCTTGTTCCGATGCGTAAAGCGTACAAGGGAACGTTCATAGTAGC

TGGGGGTTATGATCGGGAGGACGGGAACGGGCCCTGATAGAAGACCGGGC

CGACCTTGTCGCATACGGACGTTTGTTCATATCCAACCCAGATTTACCGAAA

CGTTTTGAGTTAAACGCTCCCCTGAATAAATACAATCGTGACACGTTCTATA

CTTCTGATCCAATCGTGGGTTATACGGACTATCCGTTTTTAGAGACGATGAC

GCTCGAGCACCACCACCACCACCACTGA

Protein Sequence of 12-Oxophytodienoate reductase 1 (OPR1) from Solanum lycopersicum

MENKVVEEKQVDKIPLMSPCKMGKFELCHRVVLAPLTRQRSYGYIPQPHAIL

HYSQRSTNGGLLIGEATVISETGIGYKDVPGIWTKEQVEAWKPIVDAVHAKGG

IFFCQIWHVGRVSNKDFQPNGEDPISCTDRGLTPQIRSNGIDIAHFTRPRRLTTD

EIPQIVNEFRVAARNAIEAGFDGVEIHGAHGYLIDQFMKDQVNDRSDKYGGSL

ENRCRFALEIVEAVANEIGSDRVGIRISPFAHYNEAGDTNPTALGLYMVESLNK

YDLAYCHVVEPRMKTAWEKIECTESLVPMRKAYKGTFIVAGGYDREDGNRAL

IEDRADLVAYGRLFISNPDLPKRFELNAPLNKYNRDTFYTSDPIVGYTDYPFLE

TMTLEHHHHHH

DNA sequence of OPR3

ATGACCGCGGCGCAGGGCAACAGCAACGAAACCCTGTTTAGCAGCTATAA

AATGGGCCGCTTTGATCTGAGCCATCGCGTGGTGCTGGCGCCGATGACCCG

CTGCCGCGCGCTGAACGGCGTGCCGAACGCGGCGCTGGCGGAATATTATGC

GCAGCGCACCACCCCGGGCGGCTTTCTGATTAGCGAAGGCACCATGGTGA

GCCCGGGCAGCGCGGGCTTTCCGCATGTGCCGGGCATTTATAGCGATGAAC

AGGTGGAAGCGTGGAAACAGGTGGTGGAAGCGGTGCATGCGAAAGGCGG

CTTTATTTTTTGCCAGCTGTGGCATGTGGGCCGCGCGAGCCATGCGGTGTAT

CAGCCGAACGGCGGCAGCCCGATTAGCAGCACCAACAAACCGATTAGCGA

AAACCGCTGGCGCGTGCTGCTGCCGGATGGCAGCCATGTGAAATATCCGAA

ACCGCGCGCGCTGGAAGCGAGCGAAATTCCGCGCGTGGTGGAAGATTATT

GCCTGAGCGCGCTGAACGCGATTCGCGCGGGCTTTGATGGCATTGAAATTC

ATGGCGCGCATGGCTATCTGATTGATCAGTTTCTGAAAGATGGCATTAACGA

TCGCACCGATCAGTATGGCGGCAGCATTGCGAACCGCTGCCGCTTTCTGAA

ACAGGTGGTGGAAGGCGTGGTGAGCGCGATTGGCGCGAGCAAAGTGGGC

GTGCGCGTGAGCCCGGCGATTGATCATCTGGATGCGACCGATAGCGATCCG

CTGAGCCTGGGCCTGGCGGTGGTGGGCATGCTGAACAAACTGCAGGGCGT

GAACGGCAGCAAACTGGCGTATCTGCATGTGACCCAGCCGCGCTATCATGC

GTATGGCCAGACCGAAAGCGGCCGCCAGGGCAGCGATGAAGAAGAAGCG

AAACTGATGAAAAGCCTGCGCATGGCGTATAACGGCACCTTTATGAGCAGC

GGCGGCTTTAACAAAGAACTGGGCATGCAGGCGGTGCAGCAGGGCGATGC

GGATCTGGTGAGCTATGGCCGCCTGTTTATTGCGAACCCGGATCTGGTGAG

CCGCTTTAAAATTGATGGCGAACTGAACAAATATAACCGCAAAACCTTTTAT

ACCCAGGATCCGGTGGTGGGCTATACCGATTATCCGTTTCTGGCGCCGTTTA

GCCGCCTG

Protein Sequence of OPR3

MTAAQGNSNETLFSSYKMGRFDLSHRVVLAPMTRCRALNGVPNAALAEYYA

QRTTPGGFLISEGTMVSPGSAGFPHVPGIYSDEQVEAWKQVVEAVHAKGGFIF

CQLWHVGRASHAVYQPNGGSPISSTNKPISENRWRVLLPDGSHVKYPKPRALE

ASEIPRVVEDYCLSALNAIRAGFDGIEIHGAHGYLIDQFLKDGINDRTDQYGGS

IANRCRFLKQVVEGVVSAIGASKVGVRVSPAIDHLDATDSDPLSLGLAVVGML

NKLQGVNGSKLAYLHVTQPRYHAYGQTESGRQGSDEEEAKLMKSLRMAYN

GTFMSSGGFNKELGMQAVQQGDADLVSYGRLFIANPDLVSRFKIDGELNKYN

RKTFYTQDPVVGYTDYPFLAPFSRL

DNA sequence of SYE1

ATGACCCAGAGTCTGTTTCAGCCGATTACCCTGGGTGCCCTGACCCTGAAA

AATCGCATTGTTATGCCGCCGATGACCCGTAGTCGCGCCAGTCAGCCGGGC

GATGTTGCCAATCACATGATGGCCACCTATTATGCACAGCGTGCAAGCGCC

GGCCTGATTGTTAGTGAAGGTACACAGATTAGTCCGACCGCAAAAGGCTAT

GCATGGACCCCTGGTATTTTTACCCCGGAACAGATTGCCGGTTGGCGTATTG

TTACCGAAGCAGTGCATGCCAAAAATGGCGTTATTTTTGCCCAGCTGTGGC

ATGTTGGTCGTGTGACCCATCCGGATAATATTGGCGGCCAGCAGCCGATTAG

TAGCAGCGCCCTGAAAGCCGAAAATGTTAAAGTGTTTGTGGATAATGGCAC

CGATGAACCGGGCTTTGTTGATGTTGTGGCACCGCGCGCCATGACCAAAGC

AGATATTGCACAGGTGATTGCAGATTATCGCCAGGCAGCACTGAATGCAAT

GGCTGCCGGTTTTGATGGCATTGAACTGCATGCAGCCAATGGTTATCTGATT

AATCAGTTTATTGACAGTGAGGCCAATAATCGTAGTGATGAATATGGTGGCA

GCCTGGAAAATCGCCTGCGTTTTCTGGATGAAGTTGTGGCAGCCCTGGTGG

ATGCCATTGGCGCAGAACGCGTGGGCGTGCGCCTGGCACCTCTGACAACC

CTGAATGGTACAGTGGATGCAGATCCGATTCTGACCTATACCGCCGCAGCA

GCACTGCTGAATAAACATCGCATTGTTTATCTGCATATCGCAGAAGTTGATT

GGGATGATGCCCCGGATACCCCGCGTAGTTTTAAACAGGCACTGCGTGAAG

CCTATCAGGGCGTGCTGATTTATGCCGGTCGCTATAATGCACATACCGCAGA

ACAGGCCATTAATGATGGTCTGGCCGATATGATTGGTTTTGGTCGCCCGTTT

ATTGCAAATCCGGATCTGCCGGAACGTCTGAAATATGGTTATCCGCTGGCCG

AACATGATCCGACCACCCTGTTTGGTGGTGGTGAAAAAGGTCTGACCGATT

ATCCGACCTATCAGGCCTAA

Protein sequence of SYE1

MTQSLFQPITLGALTLKNRIVMPPMTRSRASQPGDVANHMMATYYAQRASAG

LIVSEGTQISPTAKGYAWTPGIFTPEQIAGWRIVTEAVHAKNGVIFAQLWHVGR

VTHPDNIGGQQPISSSALKAENVKVFVDNGTDEPGFVDVVAPRAMTKADIAQ

VIADYRQAALNAMAAGFDGIELHAANGYLINQFIDSEANNRSDEYGGSLENR

LRFLDEVVAALVDAIGAERVGVRLAPLTTLNGTVDADPILTYTAAAALLNKHR

IVYLHIAEVDWDDAPDTPRSFKQALREAYQGVLIYAGRYNAHTAEQAINDGL

ADMIGFGRPFIANPDLPERLKYGYPLAEHDPTTLFGGGEKGLTDYPTYQA

DNA sequence of SYE3

ATGAGTCTGTTTACCGCCTATGAAAGCGCAGCACTGACCCTGCAAAATCGC

ATTGTGATGGCACCGATGACCCGTGCCCGTACCACCCAGCCGGGTAATATTC

CGAATGATCTGATGGCCCAGTATTATGCACAGCGCGCCAGCGCAGGTCTGA

TTATTACCGAAGCCACCCAGATTAGTGATGATAGTCAGGGCTATAGTTTTAC

CCCTGGTGTGTATACCGAAGCCCAGGTTGATGGCTGGAAAAAAGTGACCG

CCGCAGTGCATGAAGCCGGCGGCAAAATTTTTAATCAGATTTGGCATGTGG

GTCGCGTGAGTCATAGCATTTTTCAGCAGGGCAATGCCCCGATTGCCCCGA

GCGCCATTGCCCCTGTTGGCACCAAAGTGTGGATTGTGGATGAAGCCCATC

CGGAAGGCCAGATGGTTGATTGTCCGGAACCGCGTGAAATGACCCAGGCA

GATATTGATCGCGTGGTGAGCGATTTTGCCAATGCAGGTGCCAATGCCGTTG

CAGCCGGTTTTGATGGCATTGAAATTCATGGCGGCAATGGTTATCTGATTGA

TCAGTTTCTGCGCACCAATAGCAATCATCGCACCGATGCATACGGTGGTAGT

CCGGAAAAACGTATTCGCTTTCTGCTGGAAGTTGTGGAAGCCGTTAGCGCA

AAAATTGGCGCAGATAAAGTTGGCGTGCGCCTGGCACCGTATGTTACCTTT

AAAGATATGGCATGTCCGGAAATTGTGGAAACCATTCTGCTGGCAGCAAAA

CAGCTGAGTGCCTTTGGTATTGCCTATCTGCATCTGAGCGAAGCAGATTGG

GATGATGCACCGCAGGTTCCGGAAAGTTTTCGCGTTGAACTGCGTAAAGTT

TTTAAAGGTAGCATTATTGTGGCCGGTCGTTATGATGTGGAACGCGCAACC

GAAGTTATTGAAAAAGGCTATGCCGATCTGGTGGCATTTGGCCGTGCATTTA

TTGCCAATCCGGATCTGCCGTATCGTCTGGCCAATCAGCTGCCGCTGAGTCC

GTTTGATAAAGGTCCGCTGTTTGGTGGCAGTGCCGCCGGCTATACCGATTAT

CCGAGCTATCAGGCAGCCCTGCGTGCAGTGATTCGTAGCAGCGATGATGAA

GTTGCATAA

Protein sequence of SYE3

MSLFTAYESAALTLQNRIVMAPMTRARTTQPGNIPNDLMAQYYAQRASAGLII

TEATQISDDSQGYSFTPGVYTEAQVDGWKKVTAAVHEAGGKIFNQIWHVGRV

SHSIFQQGNAPIAPSAIAPVGTKVWIVDEAHPEGQMVDCPEPREMTQADIDRV

VSDFANAGANAVAAGFDGIEIHGGNGYLIDQFLRTNSNHRTDAYGGSPEKRIR

FLLEVVEAVSAKIGADKVGVRLAPYVTFKDMACPEIVETILLAAKQLSAFGIA

YLHLSEADWDDAPQVPESFRVELRKVFKGSIIVAGRYDVERATEVIEKGYADL

VAFGRAFIANPDLPYRLANQLPLSPFDKGPLFGGSAAGYTDYPSYQAALRAVI

RSSDDEVA

DNA sequence of SYE4

ATGACCATTGAAAACGCAGTTAATAGCGTGGGTAATCTGTTTGATACCTATA

AACTGAATGACACCATTACCCTGAAAAATCGCATTCTGATGGCCCCGCTGA

CCCGCTGCATGGCAGATGCAGATCTGGTGCCGACCGATGATATGGTGGCCT

ATTATGCACGTCGTGCAGAAGCAGGTCTGATTATTAGTGAAGCCACCATTAT

TCGCCCGGATGCACAGGGTTATCCGAATACCCCTGGTATTTTTACCCAGGGT

CAGATTGCAGGTTGGCGCAAAGTGACCGATGCAGTTCATGCAAATGGTGG

CAAAATTTTTGTTCAGCTGTGGCATACCGGCCGCGTGGCACATCCGCATTTT

TTTGGCGGCGGCGATGTGCTGGCACCGAGTGCACAGAAAATTGAAGGCAG

CGTGCCGCGTATGCGCGAACTGACCTATGTGACCCCGAAAGCAGCAACCG

TGGAAGATATTCAGGGTCTGGTTCGTGATTATGCAAAAGCAGCCGAAAATG

CAATTGAAGCAGGCTTTGATGGTGTTGAAATTCATGGCGCAAATGGCTATCT

GATTGATGAATTTCTGCATCATGATAGTAACCGCCGCACCGATGAATATGGT

GGCACCCCGGCAAATATGAGTCGTTTTGCACTGGAAGTTGTGGATGCCATT

GCAGCCCGTATTGGCAAAGATCGTACCGGCCTGCGTATTAGTCCGGGTGCC

TATTTTAATATGGCCAGTGATAGCCGTGATCGTGCCGTGTTTGATTATCTGCT

GCCGGAACTGGAAAAACGCGATCTGGCCTTTGTGCATATTGGCATTTTTGAT

GATAGCATGGAATTTGATTACCTGGGTGGCAGCGCCAGTAGTTATGTGCGTG

CCCATTATGGCAAAACCCTGGTGGGTGTTGGTAGTTATAGTGCCCAGACCG

CCAGTAAAGCCATTGCCGAAGATAAATTTGATCTGATTGCAATTGGTCGCCC

GTTTATTGCAAATCCGGATTATGTGGCAAAAGTTCGCAAAGGTGAAGAACT

GGTTGCATATAGCGATGAAATGCTGGCCACCCTGATTTAA

Protein sequence of SYE4

MTIENAVNSVGNLFDTYKLNDTITLKNRILMAPLTRCMADADLVPTDDMVAY

YARRAEAGLIISEATIIRPDAQGYPNTPGIFTQGQIAGWRKVTDAVHANGGKIF

VQLWHTGRVAHPHFFGGGDVLAPSAQKIEGSVPRMRELTYVTPKAATVEDIQ

GLVRDYAKAAENAIEAGFDGVEIHGANGYLIDEFLHHDSNRRTDEYGGTPAN

MSRFALEVVDAIAARIGKDRTGLRISPGAYFNMASDSRDRAVFDYLLPELEKR

DLAFVHIGIFDDSMEFDYLGGSASSYVRAHYGKTLVGVGSYSAQTASKAIAED

KFDLIAIGRPFIANPDYVAKVRKGEELVAYSDEMLATLI

DNA sequence of TOYE

ATGAGCATTCTGCATATGCCGCTGAAAATTAAAGATATTACCATTAAAAACC

GCATTATGATGAGCCCGATGTGCATGTATAGCGCGAGCACCGATGGCATGCC

GAACGATTGGCATATTGTGCATTATGCGACCCGCGCGATTGGCGGCGTGGG

CCTGATTATGCAGGAAGCGACCGCGGTGGAAAGCCGCGGCCGCATTACCG

ATCATGATCTGGGCATTTGGAACGATGAACAGGTGAAAGAACTGAAAAAA

ATTGTGGATATTTGCAAAGCGAACGGCGCGGTGATGGGCATTCAGCTGGCG

CATGCGGGCCGCAAATGCAACATTAGCTATGAAGATGTGGTGGGCCCGAGC

CCGATTAAAGCGGGCGATCGCTATAAACTGCCGCGCGAACTGAGCGTGGA

AGAAATTAAAAGCATTGTGAAAGCGTTTGGCGAAGCGGCGAAACGCGCGA

ACCTGGCGGGCTATGATGTGGTGGAAATTCATGCGGCGCATGGCTATCTGAT

TCATGAATTTCTGAGCCCGCTGAGCAACAAACGCAAAGATGAATATGGCAA

CAGCATTGAAAACCGCGCGCGCTTTCTGATTGAAGTGATTGATGAAGTGCG

CAAAAACTGGCCGGAAAACAAACCGATTTTTGTGCGCGTGAGCGCGGATG

ATTATATGGAAGGCGGCATTAACATTGATATGATGGTGGAATATATTAACATG

ATTAAAGATAAAGTGGATCTGATTGATGTGAGCAGCGGCGGCCTGCTGAAC

GTGGATATTAACCTGTATCCGGGCTATCAGGTGAAATATGCGGAAACCATTA

AAAAACGCTGCAACATTAAAACCAGCGCGGTGGGCCTGATTACCACCCAG

GAACTGGCGGAAGAAATTCTGAGCAACGAACGCGCGGATCTGGTGGCGCT

GGGCCGCGAACTGCTGCGCAACCCGTATTGGGTGCTGCATACCTATACCAG

CAAAGAAGATTGGCCGAAACAGTATGAACGCGCGTTTAAAAAA

Protein sequence of TOYE

MSILHMPLKIKDITIKNRIMMSPMCMYSASTDGMPNDWHIVHYATRAIGGVG

LIMQEATAVESRGRITDHDLGIWNDEQVKELKKIVDICKANGAVMGIQLAHAG

RKCNISYEDVVGPSPIKAGDRYKLPRELSVEEIKSIVKAFGEAAKRANLAGYD

VVEIHAAHGYLIHEFLSPLSNKRKDEYGNSIENRARFLIEVIDEVRKNWPENKP

IFVRVSADDYMEGGINIDMMVEYINMIKDKVDLIDVSSGGLLNVDINLYPGYQ

VKYAETIKKRCNIKTSAVGLITTQELAEEILSNERADLVALGRELLRNPYWVLH

TYTSKEDWPKQYERAFKK

DNA sequence of Ene-reductase from Yersenia bercovieri (YersER)

ATGAAGACGGCTAAGTTATTCAGTCCTCTTAAGGTGGGCGCGTTGACCCTG

CCTAATCGGGTTTTCATGGCTCCGCTTACGAGACTTCGGTCTATTGAACCTG

GGGACATTCCAACCCCCTTAATGGCTGAGTACTACCGCCAACGTGCCTCGG

CGGGGTTAATAATAACCGAAGCGACCCAAATAAGCTTCCAGGCGAAAGGTT

ACGCCGGTGCGCCGGGCTTACACACGCAGGAACAATTAAACGCTTGGAAG

AAGATTACGCAAGCTGTCCACGAGGAAGGTGGACACATTGCCGTTCAGTT

ATGGCACGTGGGCCGCATCTCGCATAGCTCGCTGCAGCCAGGACAACAAG

CACCAGTGGCCCCTTCCGCGATTGCGGCTGATACGAGAACGACGGTACGC

GATGAGAATGGGGCATGGGTACGTGTCCCCTGCTCGACGCCACGCGCGTTG

GAGACTGAGGAGATACCTGGTATTATAAATGATTTCCGTCAGGCAACCGCTA

ACGCTAGAGAGGCAGGCTTTGATTACATAGAATTACACGCCGCGCATGGTT

ACCTGTTGCATCAGTTTATGAGTCCTGCTAGTAATCAGCGGACAGACCAGT

ACGGAGGGTCCATAGAAAATCGGACCCGGTTGACGTTGGAGGTCGTCGAC

GCCACCGCAGCCCAATGGTCCGCCGAGCGGATAGGCATCCGTATAAGTCCA

CTTGGTCCTTTTAATGGGCTTGACAACGGGGAAGACCAGGAGGAGGCCGC

GCTGTATTTAATCGATGAACTGAACAAACGGCATATCGCTTATCTGCATATCT

CAGAACCGGACTGGGCAGGAGGGAAGCCTTACAGTGAAGCGTTCAGAGA

CGCAGTCCGTGCTCGTTTCAAAGGGGTAATCATTGGCGCAGGAGCATATAC

CGCCGAGAAAGCAGAGGAACTTATAGAGAAGGGCTTCATTGACGCGGTGG

CTTTTGGACGTTCATATATCTCCAACCCAGACCTTGTGGCGAGATTACAGCA

GCATGCCCCCTTGAATGAACCAGATGGAGAAACGTTTTACGGAGGAGGGG

CAAAAGGATATACTGATTATCCTACACTGCTCGAGCACCACCATCACCACCA

CTGA

Protein Sequence of Ene-reductase from Yersenia bercovieri (YersER)

MKTAKLFSPLKVGALTLPNRVFMAPLTRLRSIEPGDIPTPLMAEYYRQRASAG

LIITEATQISFQAKGYAGAPGLHTQEQLNAWKKITQAVHEEGGHIAVQLWHVG

RISHSSLQPGQQAPVAPSAIAADTRTTVRDENGAWVRVPCSTPRALETEEIPGII

NDFRQATANAREAGFDYIELHAAHGYLLHQFMSPASNQRTDQYGGSIENRTR

LTLEVVDATAAQWSAERIGIRISPLGPFNGLDNGEDQEEAALYLIDELNKRHIA

YLHISEPDWAGGKPYSEAFRDAVRARFKGVIIGAGAYTAEKAEELIEKGFIDAV

AFGRSYISNPDLVARLQQHAPLNEPDGETFYGGGAKGYTDYPTLLEHHHHHH

DNA sequence of NADPH dehydrogenase from Bacillus subtilis (YqjM)

ATGCACCACCATCACCACCACGCCCGTAAGCTGTTCACGCCCATCACCATT

AAGGATATGACTTTGAAAAACCGTATCGTTATGAGTCCCATGTGCATGTACA

GCAGCCATGAAAAAGACGGAAAATTAACTCCGTTTCATATGGCGCATTATAT

CAGTCGTGCAATCGGCCAAGTTGGTCTTATTATCGTGGAGGCAAGTGCCGT

AAATCCCCAGGGACGTATTACGGATCAAGATTTGGGTATCTGGAGCGATGA

ACACATCGAAGGCTTCGCGAAGCTGACAGAACAGGTTAAGGAACAAGGGT

CTAAGATCGGCATTCAACTGGCCCACGCCGGACGTAAGGCTGAATTGGAG

GGTGACATCTTTGCTCCATCTGCTATCGCGTTTGACGAGCAATCTGCGACTC

CGGTCGAGATGAGCGCTGAGAAGGTGAAGGAAACAGTGCAAGAGTTCAA

GCAGGCAGCAGCACGTGCGAAGGAAGCAGGGTTCGATGTGATTGAGATCC

ATGCAGCACATGGTTATCTGATTCACGAGTTTCTGTCCCCTCTGTCAAACCA

TCGCACCGATGAGTATGGAGGAAGCCCTGAGAATCGCTATCGCTTCCTGCG

TGAAATTATCGATGAAGTTAAACAGGTTTGGGACGGTCCGCTTTTTGTGCG

CGTGTCTGCCTCAGATTACACGGATAAGGGCTTGGATATTGCTGACCACATC

GGGTTCGCAAAGTGGATGAAGGAGCAAGGAGTGGACTTAATTGATTGCAG

CAGCGGGGCTTTAGTACACGCGGACATTAACGTATTCCCGGGCTACCAAGT

TTCCTTTGCAGAAAAGATCCGCGAACAAGCGGATATGGCAACAGGTGCTGT

TGGGATGATCACGGACGGTTCGATGGCCGAGGAAATCCTTCAAAACGGCC

GTGCCGACTTGATCTTTATCGGTCGTGAATTACTTCGCGACCCTTTTTTTGC

TCGCACCGCAGCGAAACAATTAAATACGGAAATTCCTGCACCAGTGCAATA

C GAGCGTGGTTGGTGA

Protein sequence of NADPH dehydrogenase from Bacillus subtilis (YqjM)

MHHHHHHARKLFTPITIKDMTLKNRIVMSPMCMYSSHEKDGKLTPFHMAHYI

SRAIGQVGLIIVEASAVNPQGRITDQDLGIWSDEHIEGFAKLTEQVKEQGSKIGI

QLAHAGRKAELEGDIFAPSAIAFDEQSATPVEMSAEKVKETVQEFKQAAARA

KEAGFDVIEIHAAHGYLIHEFLSPLSNHRTDEYGGSPENRYRFLREIIDEVKQV

WDGPLFVRVSASDYTDKGLDIADHIGFAKWMKEQGVDLIDCSSGALVHADIN

VFPGYQVSFAEKIREQADMATGAVGMITDGSMAEEILQNGRADLIFIGRELLR

DPFFARTAAKQLNTEIPAPVQYERGW

DNA sequence of XenB

ATGACCACGCTTTTCGATCCGATCACCCTGGGCGACCTGCAACTGCCCAAC

CGCATCATCATGGCCCCACTCACCCGCTGCCGTGCCGACGAAGGCCGCGTG

CCCAATGCGCTGATGGCCGAATACTACGTGCAGCGCGCCAGCGCCGGGCTG

ATCCTCAGCGAGGCGACGTCGGTCAGTGCCATGGGGGTCGGCTACCCGGAT

ACCCCCGGTATCTGGAACGACGAGCAGGTACGTGGCTGGAACAACGTCAC

CAAGGCTGTGCATGCCGCAGGCGGACGCATCTTCCTGCAACTGTGGCACG

TTGGCCGTATCTCCCACCCCAGCTACCTGAACGGCGAGCTGCCGGTGGCCC

CCAGCGCGATCCAGCCCAAGGGGCATGTAAGCCTGGTGCGCCCGTTGAGT

GATTACCCCACCCCACGCGCGCTGGAAACCGAAGAAATCATCGACATCGTC

GAGGCTTACCGCAGCGGTGCCGAGAATGCCAAGGCGGCTGGCTTTGATGG

TGTAGAAATCCACGGCGCCAACGGTTACCTGCTCGACCAGTTCCTGCAAAG

CAGCACCAACCAGCGCACTGACCGCTATGGCGGCTCGCTGGAAAACCGTG

CACGCCTGCTACTGGAAGTGACCGATGCAGCCATTGAAGTGTGGGGCGCG

AACCGCGTGGGCGTGCACCTGGCACCGCGTGCCGATGCGCACGACATGGG

CGATGCCGATCGCGCCGAAACCTTTACCTACGTGGCCCGTGAACTGGGCAA

GCGTGGCATTGCCTTTATCTGCTCGCGGGAAAGGGAAGCCGATGACAGCAT

TGGTCCACTGATCAAGGAAGCGTTTGGCGGCCCGTACATCGTCAATGAGCG

GTTTGACAAAGCCAGTGCCAATGCGGCGCTGGCCAGCGGCAAGGCCGATG

CTGTGGCGTTTGGCGTGCCGTTCATCGCCAACCCTGATTTGCCGGCGCGGC

TGGCGGCGGATGCGCCACTGAACGAGGCGCGACCCGAGACCTTCTATGGC

AGGGGCCCGGTGGGGTACATCGATTACCCGCGGTTGGGATCCCATCATCAT

CATCATCATTGA

Protein sequence of XenB

MTTLFDPITLGDLQLPNRIIMAPLTRCRADEGRVPNALMAEYYVQRASAGLIL

SEATSVSAMGVGYPDTPGIWNDEQVRGWNNVTKAVHAAGGRIFLQLWHVG

RISHPSYLNGELPVAPSAIQPKGHVSLVRPLSDYPTPRALETEEIIDIVEAYRSGA

ENAKAAGFDGVEIHGANGYLLDQFLQSSTNQRTDRYGGSLENRARLLLEVTD

AAIEVWGANRVGVHLAPRADAHDMGDADRAETFTYVARELGKRGIAFICSR

EREADDSIGPLIKEAFGGPYIVNERFDKASANAALASGKADAVAFGVPFIANPD

LPARLAADAPLNEARPETFYGRGPVGYIDYPRLGSHHHHHH

### 3. Directed Evolution of SYE2 and GluER

#### 3.1 Cloning and Site-saturation Mutagenesis

pET-28a(+) was used as the cloning and expression vector for SYE2 while pET-30a(+) was used as the cloning and expression vector for GluER described in this study. Genes of ene-reductases (ERs) were codon optimized using *E. coli* as the host organism and purchased as gBlocks from Genscript as plasmids in a desired vector. The genes of ERs used in this study were either cloned into pET-30a(+) between Nde I and Hind III (N-terminal 6X His tag) or cloned into pET-28a(+) between Nde I and Xho I (C-terminal 6X His tag). Site-saturation mutagenesis was performed using the "22c-trick" method as described by Kille et al[1]. The PCR products were ligated using a Gibson mix prepared from 5X isothermal (ISO) reaction buffer (25% PEG-8000, 500 mM Tris-HCl pH 7.5, 50 mM $MgCl_2$, 50 mM DTT, 1 mM each of the dNTPs, and 5 mM NAD), T5 exonuclease, Phusion DNA polymerase, and Taq DNA ligase[2]. The ligation mixture was used directly to transform chemically competent *E. coli* BL21(DE3) cells (TransGen).

#### 3.2 Expression of ER Variants in 96-Well Plates

Single colonies from LBkan agar plates were picked using sterile toothpicks and cultured in deep-well 96-well plates containing LBkan (1000 μL/well), and ERs were expressed using the addition of 4% (v/v) auto inducing mix (sterile filtered mixture of 1.25% glucose, 5% lactose and 15% glycerol) at 30 ºC, 800 rpm for 24 h.

#### 3.3 Reaction Screening in 96-Well Plate Format

E. coli (E. cloni BL21(DE3)) cells in deep-well 96-well plates were pelleted (3,000 g, 10 min, 4 °C) using an Eppendorf tabletop centrifuge 5910R and resuspended in Tris buffer (50mM, pH 9, 360 μL/well) by gentle shaking using a Fisher Scientific microplate shaker (1000 rpm, 5 min), and the 96-well plate was then transferred into a Coy anaerobic chamber. In the anaerobic chamber, the bacterial solution in the 96-deep-well plate is transferred to a metal 96-well plate equipped with magnetic stirrers and glass tubes using an Eppendorf Xplorer 12-channel pipette (50-1200 μL), 20 μL of the substrates stock solution (**1a** 800 mM, **2a** 400mM in EtOH) was added into each well using an Eppendorf Xplorer 12-channel pipette (1-20 μL). Finally, 10 μL FMN-Na (40 mM in $H_2O$, 10 mol%) and 10 μL **FI** (20 mM in EtOH, 5 mol%) were added to each well using an Eppendorf Xplorer 12-channel pipette (0.5-10 μL). After tightening the screws of the metal cover, the metal 96-well plate was taken out of the anaerobic chamber and fixed in a ROGER 96-position parallel photoreactor (see Supplementary Fig. 1). After 12 hours of blue LED exposure, the 96-well plate was taken out of 96-position parallel photoreactor, the screws were loosened and the cover was opened. The reaction mixture in each well was transferred to a 2 mL centrifuge tube, 1 mL EtOAc and 10 μL dodecane (20 mg/mL in EtOAc) as internal standard were added, vortexed for 1 min, and centrifuged at 12,000 rpm for 4 min. The supernatant was taken and transferred to a centrifuge tube containing about 20 mg of anhydrous sodium sulfate. The supernatant was vortexed for 1 min, followed by centrifugation at 12,000 rpm for 2 min. Finally, a syringe was used to inject the supernatant into a 2.5 mL liquid phase vial through a 0.22 μm organic filter membrane. The yield was determined by gas chromatography (GC) and enantiomeric ratio (er) was determined by chiral HPLC.

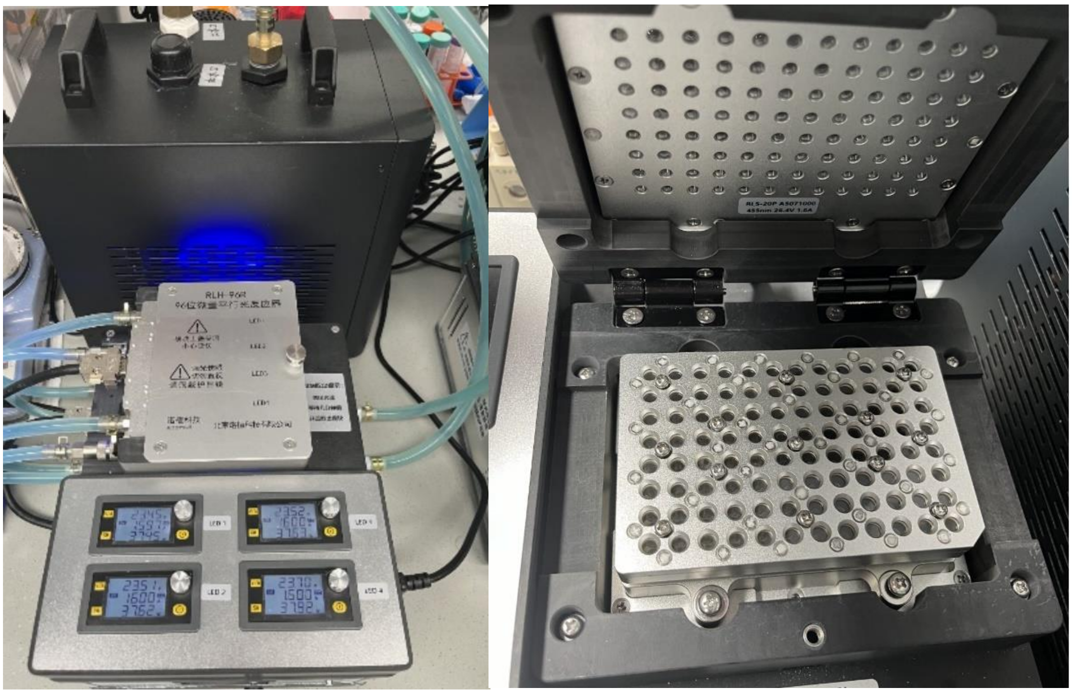

**Supplementary Fig. 1 | 96 well plate photoreactor.**

#### 3.4 Summary of Directed Evolution of ERs

We conducted directed evolution of SYE2 and GluER respectively using **1a** + **2a** → **3a** as the model reaction, hoping to obtain higher yields and better enantioselectivities. However, after we performed site-saturated mutagenesis on a total of 54 sites in the active and substrate binding sites of SYE2, no beneficial mutations were obtained. For GluER, we investigated 11 sites within 3 Å region of FMN binding site and substrate channels, including T36, W66, H172, N175, Y177, T231, Q232, P239, R261, F269 and Y343. The mutation sites and primers were summarized in Supplementary Table 1 and 2.

**Supplementary Table 1 | Primers used in directed evolution of SYE2.**

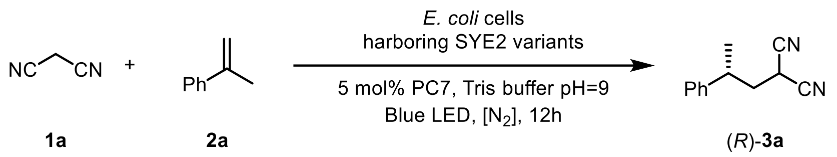

| Sites | Primer Name | Primer Sequence |
|---|---|---|
| A19 | 19-F | AATCGTATTGTGATGNNKCCGAT |
| | 19-R | CATCACAATACGATTTTTGG |
| P20 | 20-F | CGTATTGTGATGGCANNKATGAC |
| | 20-R | TGCCATCACAATACGATTTT |
| M21 | 21-F | ATTGTGATGGCACCGNNKACCCGTA |
| | 21-R | CGGTGCCATCACAATACGAT |
| T22 | 22-F | GTGATGGCACCGATGNNKCGTAGC |

| | | |
|---|---|---|
| | 22-R | CATCGGTGCCATCACAATAC |
| R23 | 23-F | ATGGCACCGATGACCNNKAGCCGCAGT |
| | 23-R | GGTCATCGGTGCCATCACAA |
| S24 | 24-F | GCACCGATGACCCGTNNKCGCAGT |
| | 24-R | ACGGGTCATCGGTGCCATCA |
| E53 | 53-F | GGCCTGATTATTGCCNNKGGCGCACC |
| | 53-R | GGCAATAATCAGGCCGGCGC |
| G54 | 54-F | CTGATTATTGCCGAANNKGCACCG |
| | 54-R | TTCGGCAATAATCAGGCCGG |
| G63 | 63-F | AGCGCCGTGGCACGTNNKTATAGCA |
| | 63-R | ACGTGCCACGGCGCTCACCG |
| Y64 | 64-F | GCCGTGGCACGTGGTNNKAGCATGA |
| | 64-R | ACCACGTGCCACGGCGCTCA |
| T67 | 67-F | CGTGGTTATAGCATGNNKCCTGG |
| | 67-R | CATGCTATAACCACGTGCCA |
| Q96 | 96-F | GGCAAAATTTTATTNNKCTGTGG |
| | 96-R | AATAAAAATTTTGCCGCCTT |
| W98 | 98-F | ATTTTTATTCAGCTGNNKCATGT |
| | 98-R | CAGCTGAATAAAAATTTTGC |
| V100 | 100-F | ATTCAGCTGTGGCATNNKGGCCGTC |
| | 100-R | ATGCCACAGCTGAATAAAAA |
| R103 | 103-F | TGGCATGTGGGCCGTNNKAGTCATAGCAG |

| | | |
|---|---|---|
| | 103-R | ACGGCCCACATGCCACAGCTGAAT |
| Q124 | 124-F | GCTATTAAAATTCCGGATNNKGTTTTTGGTCCGC |
| | 124-R | ATCCGGAATTTTAATAGCGCTCGGGG |
| F126 | 126-F | ATTCCGGATCAGGTTNNKGGTCCG |
| | 126-R | AACCTGATCCGGAATTTTAA |
| E173 | 173-F | GGCTTTGATGGTGTTNNKGTGCAT |
| | 173-R | AACACCATCAAAGCCGGCCA |
| H175 | 175-F | GATGGTGTTGAAGTGNNKGCCGCCC |
| | 175-R | CACTTCAACACCATCAAAGC |
| A176 | 176-F | GGTGTTGAAGTGCATNNKGCCCACG |
| | 176-R | ATGCACTTCAACACCATCAA |
| A177 | 177-F | GTTGAAGTGCATGCCNNKCACGGT |
| | 177-R | GGCATGCACTTCAACACCAT |
| H178 | 178-F | GAAGTGCATGCCGCCNNKGGTTA |
| | 178-R | GGCGGCATGCACTTCAACAC |
| G179 | 179-F | GTGCATGCCGCCCACNNKTATCT |
| | 179-R | GTGGGCGGCATGCACTTCAA |
| Y180 | 180-F | CATGCCGCCCACGGTNNKCTGTT |
| | 180-R | ACCGTGGGCGGCATGCACTT |
| R227 | 227-F | GGTCGCGTTGCCGTGNNKATTAGT |
| | 227-R | CACGGCAACGCGACCGCTAC |
| S229 | 229-F | GTTGCCGTGCGCATTNNKCCGCA |

| | | |
|---|---|---|
| | 229-R | AATGCGCACGGCAACGCGAC |
| I232 | 232-F | CGCATTAGTCCGCATNNKGGCGAAG |
| | 232-R | ATGCGGACTAATGCGCACGG |
| E234 | 234-F | AGTCCGCATATTGGCNNKGGCTTT |
| | 234-R | GCCAATATGCGGACTAATGC |
| G235 | 235-F | CCGCATATTGGCGAANNKTTTACC |
| | 235-R | TTCGCCAATATGCGGACTAA |
| H263 | 263-F | AATCTGGCCTATGTGNNKTTTAG |
| | 263-R | CACATAGGCCAGATTCATCG |
| S265 | 265-F | GCCTATGTGCATTTTNNKGAAAATATTAGCCGC |
| | 265-R | AAATGCACATAGGCCAGATTCATCGGTTGCAGT |
| E266 | 266-F | TATGTGCATTTTAGCNNKAATAT |
| | 266-R | GCTAAAATGCACATAGGCCA |
| I268 | 268-F | CATTTTAGCGAAAATNNKAGCCGCTATGTGGAA |
| | 268-R | TTTTCGCTAAAATGCACATAGGCCAGATTCATC |
| S269 | 269-F | TTTAGCGAAAATATTNNKCGCTATGTGGAAGTG |
| | 269-R | ATATTTTCGCTAAAATGCACATAGGCCAGATTC |
| R270 | 270-F | GCGAAAATATTAGCNNKTATGTGGAAGTGAGTG |
| | 270-R | GCTAATATTTTCGCTAAAATGCACATAGGCC |
| V272 | 272-F | AATATTAGCCGCTATNNKGAAGT |
| | 272-R | ATAGCGGCTAATATTTTCGC |
| E273 | 273-F | ATTAGCCGCTATGTGNNKGTGAGT |

| | | |
|---|---|---|
| | 273-R | CACATAGCGGCTAATATTTT |
| V274 | 274-F | AGCCGCTATGTGGAANNKAGTGA |
| | 274-R | TTCCACATAGCGGCTAATAT |
| S275 | 275-F | CGCTATGTGGAAGTGNNKGATG |
| | 275-R | CACTTCCACATAGCGGCTAA |
| D276 | 276-F | TATGTGGAAGTGAGTNNKGCCTTT |
| | 276-R | ACTCACTTCCACATAGCGGC |
| A277 | 277-F | GTGGAAGTGAGTGATNNKTTTCGC |
| | 277-R | ATCACTCACTTCCACATAGC |
| A293 | 293-F | CATCCGATTATGGTTNNKGGCAAA |
| | 293-R | AACCATAATCGGATGCTGAT |
| G294 | 294-F | CCGATTATGGTTGCCNNKAAACT |
| | 294-R | GGCAACCATAATCGGATGCT |
| K295 | 295-F | ATTATGGTTGCCGGCNNKCTGACCAAACAGAGC |
| | 295-R | CCGGCAACCATAATCGGATGCTGATAAACGCTA |
| T297 | 297-F | GTTGCCGGCAAACTGNNKAAACAGAGCGCACAG |
| | 297-R | AGTTTGCCGGCAACCATAATCGGATGCTGATAA |
| A314 | 314-F | TATGCCGATTTTGTTNNKTTTGGTACACCGTTT |
| | 314-R | ACAAAATCGGCATAATGCTGATCCAGCAGACGC |
| G316 | 316-F | GATTTTGTTGCCTTTNNKACACC |
| | 316-R | AAAGGCAACAAAATCGGCAT |
| T317 | 317-F | TTTGTTGCCTTTGGTNNKCCGTTT |

| | | |
|---|---|---|
| | 317-R | ACCAAAGGCAACAAAATCGG |
| P318 | 318-F | GTTGCCTTTGGTACANNKTTTGTGACCAATCCG |
| | 318-R | GTACCAAAGGCAACAAAATCGGCATAATGCTGA |
| V320 | 320-F | TTTGGTACACCGTTTNNKACCAA |
| | 320-R | AAACGGTGTACCAAAGGCAA |
| T321 | 321-F | GGTACACCGTTTGTGNNKAATCC |
| | 321-R | CACAAACGGTGTACCAAAGG |
| F338 | 338-F | TGGCCGCTGACCGAANNKGATGCAGATGCACGT |
| | 338-R | TCGGTCAGCGGCCAATCATGGGCAAAACGTGCC |
| R343 | 343-F | AATTTGATGCAGATGCANNKCTGACCCTGTATG |
| | 343-R | TGCATCTGCATCAAATTCGGTCAGCG |
| L346 | 346-F | GATGCACGTCTGACCNNKTATGGTGGCGGCGAA |
| | 346-R | GTCAGACGTGCATCTGCATCAAATTCGGTCAGC |
| Y347 | 347-F | GCACGTCTGACCCTGNNKGGTGGCGGCGAAGCA |
| | 347-R | AGGGTCAGACGTGCATCTGCATCAAATTCGGTC |

**Supplementary Table 2 | Primers used in directed evolution of GluER.**

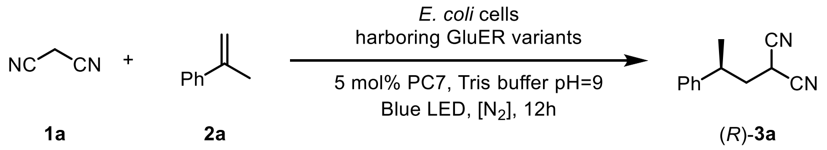

| Sites | Primer Name | Primer Sequence |
|---|---|---|
| W66A | 66A-F | CGCGAAGGCTTAGGTGCGCCGTTTGCGCCG |
| | 66A-R | ACCTAAGCCTTCGCGTGAAATCCCC |
| W66Q | 66Q-F | CGCGAAGGCTTAGGTCAACCGTTTGCGCCG |
| | 66Q-R | ACCTAAGCCTTCGCGTGAAATCCCC |
| Y343A | 343A-F | CGATATGAAAACATGGGCCTCCCAAGGCCCAG AG |
| | 343A-R | CCATGTTTTCATATCGTCTGGTTGCAGTGC |

#### 3.5 Protein Engineering and Molecular Docking Results

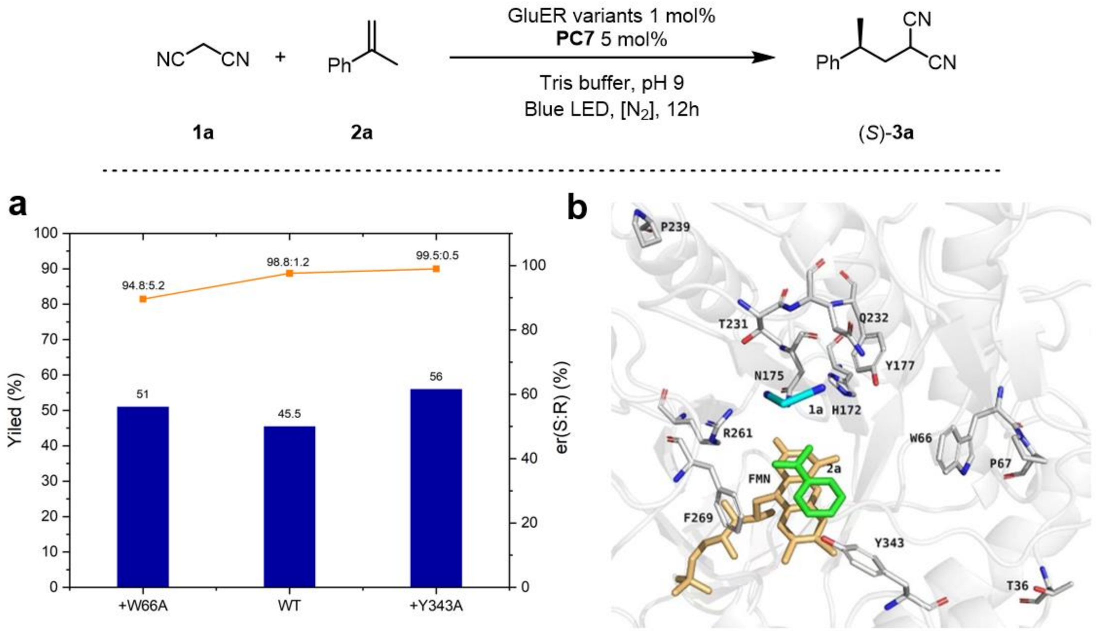

**Supplementary Fig. 2 | a, Protein engineering. b, molecular docking.** All reactions were carried out with **1a** (0.008 mmol), **2a** (0.004 mmol), GluER-variants (1 mol%), FI (5 mol%) and EtOH (7.5 v/v%) in Tris buffer (50 mM, pH 9) were stirred at room temperature for 12 h under an $N_2$ atmosphere with 455nm LEDs. The total volume of reaction was 400 μL. Yield was determined by gas chromatography and the enantiomeric ratio was determined by HPLC analysis on a chiral stationary phase. Molecular docking of substrates **1a** (cyan and blue) and **2a** (green) into GluER-WT (PDB ID: 6O08) with key active-site residues shown in grey and FMN in yellow. FMN-Na = sodium riboflavin 5-monophosphate.

### 4. Preparation of Starting Materials and Corresponding Racemic Hydroalkylated Products

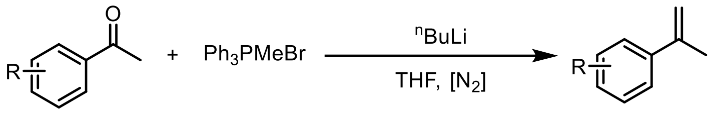

**2**

**Supplementary Scheme 1 | General methods for the synthesis of olefins.**

Substrates **2e** - **2k**, **2m** - **2o** were prepared according to literatures.[3] Other substrates were commercially available and used without any purification.

Methyltriphenylphosphonium bromide (1.5 mmol) in dry THF (10 mL) under $N_2$ atmosphere was cooled to 0 °C. Then, $^{n}$BuLi (1.6 M solution in hexane, 1.5 mmol) was added slowly to the solution. After, the resulting orange mixture was maintained at 0 °C for 0.5-1 h, a solution of the corresponding ketone (1.0 mmol) in dry THF was added dropwise at 0 °C. The reaction was allowed to warm up to rt, stirred overnight (monitored by TLC), and finally quenched with sat. $NH_4Cl$. The resulting mixture was extracted with DCM. The combined organic phase was washed with brine, dried over $Na_2SO_4$, and concentrated under reduced pressure. The resulting crude product was purified by flash column chromatography (hexane/ ethyl acetate 20:1) to give the corresponding alkenes **2**.

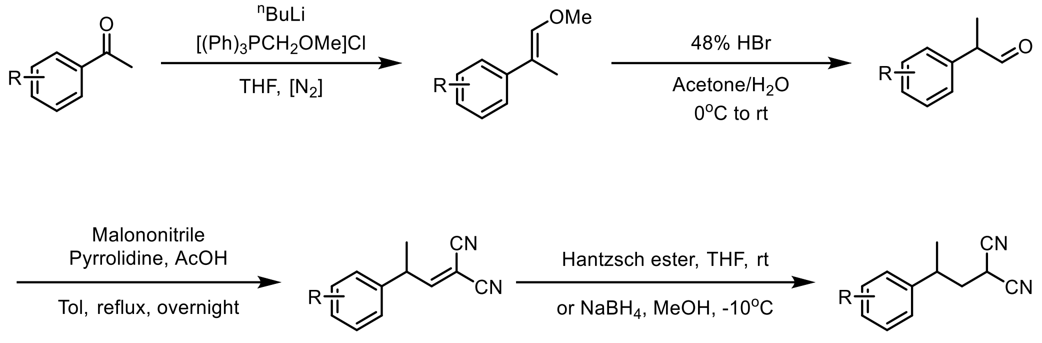

3

**Supplementary Scheme 2 | Synthesis of racemic products.**

In a two-necked round bottom flask under $N_2$ atmosphere, a suspension of the phosophonium chloride (15 mmol) in dry THF was cooled to 0 °C. With continuous stirring, $^n$BuLi (15 mmol) was added slowly to the solution. This reaction mixture was allowed to stir for 0.5-1 h, and a solution of the ketone substrate (10 mmol) in dry THF was added dropwise through a dropping funnel. The mixture was allowed to stir for another 30 minutes at 0 °C and was warmed up to ambient temperature, after which it was allowed to stir for another 16 h, and finally quenched with sat. $NH_4Cl$. The resulting mixture was extracted with DCM. The combined organic phase was washed with brine and dried over $Na_2SO_4$. All solvents were removed and the residue was purified using column chromatography (hexane/ ethyl acetate 50:1 or 20:1) to give the enol ether intermediate as colorless oil.[4]

The enol ether intermediate was dissolved in 20 mL acetone and 5 mL water. The solution was cooled to 0 °C using ice bath with continuous stirring. 3 - 4 mL of HBr (48 %) was added dropwise and the reaction was left to stir at 0 °C for another 30 minutes before it was allowed to warm up to ambient temperature. After which the reaction mixture was left to stir for another 16 h. The reaction was quenched carefully with the addition of $NaHCO_3$ and extracted with DCM (20 mL x 4). The combined organic phase was washed with brine and dried over $Na_2SO_4$. All solvents were removed and the residue was purified using column chromatography (hexane/ ethyl acetate 15:1) to give the aldehyde as colorless oil.[5]

The racemic products **3** were prepared according to the literature procedures from the corresponding aldehydes through a sequence of a Knoevenagel condensation[6,7] followed by a Hantzsch ester[8] or $NaBH_4$ reduction[9]. The solution of aromatic aldehyde (10 mmol, 1.0 equiv), malononitrile (10 mmol, 1.0 equiv), pyrrolidine (2 mmol, 0.2 equiv), and acetic acid (2 mmol, 0.2 equiv) in toluene (0.4 M) was refluxed in an oil bath for 18 h. Upon completion, evaporation of the solvent gave the crude condensation product, which was purified by silica gel column chromatography. Then, to the respective alkene (2 mmol, 1.0 equiv) in methanol (20 mL) at -10 °C, $NaBH_4$ (2.2 mmol, 1.1 equiv) was added slowly. After stirring for 15 min, the reaction mixture was quenched with $H_2O$ (2 mL), concentrated under reduced pressure, diluted with water, and extracted with DCM (15 mL × 3). The combined organic layers were dried over $Na_2SO_4$, filtered, and concentrated in vacuo. The product was obtained after purification by column chromatography on silica gel (hexane/ ethyl acetate 8:1) to provide **3**.

### 5. Typical Procedure for Photoenzymatic Reactions

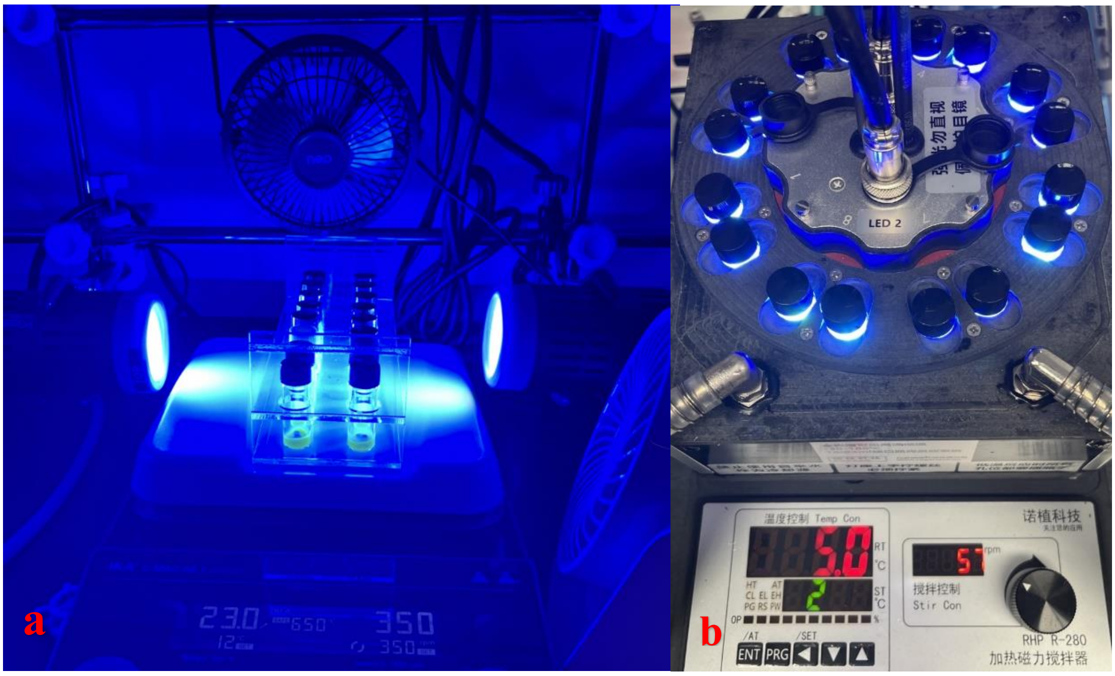

**Supplementary Fig. 3 | Photoenzymatic reaction set -up. a**. Reaction vials were placed on the stirplate, in a distance of ~ 10 cm from a 40 W 456 nm Blue LED lamp, with a cooling fan; **b**. Reaction vials were placed in ROGER photo-reactor, with a cooling pump.

All enzymatic reactions were assembled in a glove box with $O_2$ concentration below 5 ppm. Taking **1a** + **2a** → **3a** as an example, to a 4 mL vial containing a magnetic stir bar, solutions of ene-reductase (1 mol%), FI (10 µL, 2 mM stock in EtOH, 5 mol% for **1a** + **2a** → **3a**), and substrates mix (20 µL, 400 mM **1a** and 200 mM **2a** in EtOH) were added to Tris buffer (50 mM, pH 9). The total volume of the reaction mixture was 400 µL and the final concentrations for EtOH was 7.5%. The vial was sealed with screw cap, removed from the glove box, illuminated with Blue LEDs, and stirred for 12 h with a cooling fan (See Supplementary Fig. 3 for reaction set-up).

For the reaction that yield was determined by gas chromatography (GC), EtOAc (1.0 mL) and 10.0 μL of an internal standard stock (20 mg/mL of n-dodecane in EtOAc) were added to the reaction mixture and mixed thoroughly. For the reaction that yield was determined by reverse phase HPLC, MeCN (1.0 mL) and 10.0 μL of an internal standard stock (16.8 mg/mL 1,3,5-trimethoxybenzene in MeCN) were added to the reaction mixture and mixed thoroughly. The organic phase was separated and then analysed by GC (or reverse phase HPLC), GC-MS and chiral HPLC. The product formation was confirmed by the comparisons of retention times in GC/HPLC, mass spectra and $^{1}$H NMR spectra of the crude material and the reference. Yield determined by GC was relative to the n-dodecane internal standard. Yield determined by reverse phase HPLC was relative to the 1,3,5-trimethoxybenzene internal standard. Enantioselectivity was determined by chiral HPLC.

All the 0.004 mmol reactions were repeated at least twice independently using different batches of enzyme. Selected examples were reproduced by another person from the same group. GC and reverse phase HPLC yields were typically within 10% error among each run, and enantioselectivity was almost identical in duplicate runs (within 2% error bar).

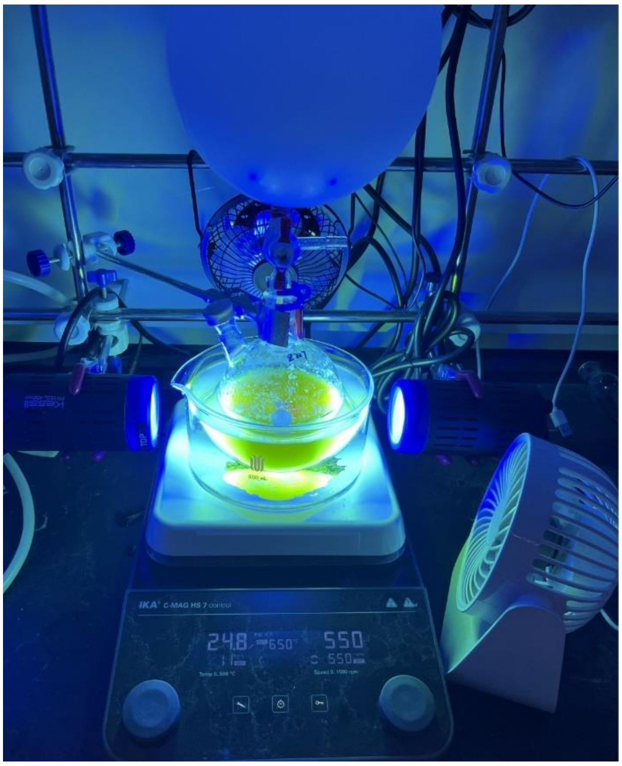

**Supplementary Fig. 4 | The setup for photoenzymatic large scale reaction.**

**Supplementary Table 3 | Effect of ERs for the reaction 1a + 2a → 3a.**

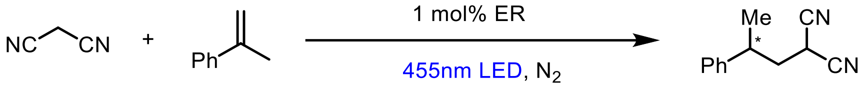

1a 2a 3a

| Entry | ERs | Yield (%) | er (%) (R:S) |
|---|---|---|---|
| 1 | CsER | 1.2 | 70.2:29.8 |
| 2 | KYE1 | n.r. | n.a. |
| 3 | LacER | n.r. | n.a. |
| 4 | MorB | 2 | 58.8:41.2 |
| 5 | OYE2 | 2.5 | 40.9:59.1 |
| 6 | OYE3 | n.r. | n.a. |
| 7 | OPR1 | n.r. | n.a. |
| 8 | OPR3 | n.r. | n.a. |
| 9 | SYE1 | 1.4 | 46:54 |
| 10 | SYE2 | n.r. | n.a. |
| 11 | SYE3 | 2.7 | 56.3:43.7 |
| 12 | SYE4 | n.r. | n.a. |
| 13 | TOYE | n.r. | n.a. |
| 14 | YersER | 1.5 | 87.5:12.5 |
| 15 | YqjM | n.r. | n.a. |
| 16 | XenB | 1.7 | 85.7:14.3 |

Typical Conditions: **1a** (0.008 mmol), **2a** (0.004 mmol), ERs (1 mol%), in a solvent mixture of Tris buffer (50 mM, pH 9) / EtOH (5 % v/v) were stirred for 12 hours under inert atmosphere with 455nm LEDs. The total volume of the reaction was 400 μL. Yield was determined by GC. Enantiomeric ratio was determined by HPLC analysis on a chiral stationary phase. n.r. No reaction; n.a. Not applicable.

**Supplementary Table 4 | Effect of the buffer type and pH for the reaction 1a + 2a → 3a.**

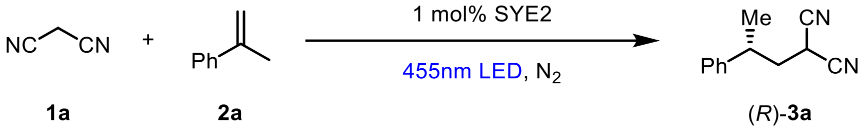

| Entry | Buffer | pH | Yield (%) | er (%) (R:S) |
|---|---|---|---|---|
| 1 | 50 mM Tris | 7.4 | n.r. | n.a. |
| 2 | 50 mM Tris | 8 | n.r. | n.a. |
| 3 | 50 mM Tris | 9 | 2.6 | 93:7 |
| 4 | 50 mM Tris | 10 | 1.5 | 85:13 |
| 5 | 50 mM KPi | 9 | 2.5 | 90.6:9.4 |
| 6 | 50 mM M9-N | 9 | 1.3 | n.d. |
| 7 | 50 mM PBS | 9 | 1.8 | n.d. |

Typical Conditions: **1a** (0.008 mmol), **2a** (0.004 mmol), ERs (1 mol%), in a solvent mixture of Tris buffer (50 mM, pH 9) / EtOH (5 % v/v) were stirred for 12 hours under inert atmosphere with 455nm LEDs. The total volume of the reaction was 400 μL. Yield was determined by GC. Enantiomeric ratio was determined by HPLC analysis on a chiral stationary phase. n.r. No reaction; n.a. Not applicable.

**Supplementary Table 5 | Effect of the co-solvents for the reaction 1a + 2a → 3a.**

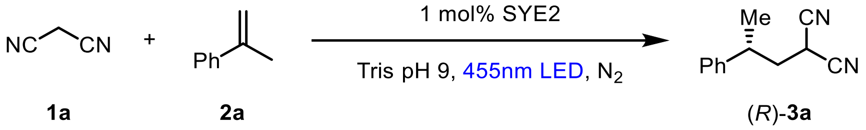

| Entry | co-Solvent | Yield (%) | er (%) (R:S) |
|---|---|---|---|
| 1 | EtOH | 2.6 | 93:7 |
| 2 | DMSO | 1.2 | 92:8 |
| 3 | DMF | 0.7 | 79.4:20.6 |
| 4 | THF | 2 | 91.2:8.8 |
| 5 | MeCN | 1.2 | 88:12 |
| 6 | iPrOH | 0.6 | 92.6:7.4 |
| 7 | MeOH | 0.7 | 90.5:9.5 |
| 8 | Acetone | n.r. | n.a. |

Typical Conditions: **1a** (0.008 mmol), **2a** (0.004 mmol), SYE2 (1 mol%), in a solvent mixture of 50mM Tris buffer (pH 9) / co-solvent (5 % v/v) were stirred for 12 hours under inert atmosphere with 455nm LEDs. The total volume of the reaction was 400 μL. Yield was determined by GC. Enantiomeric ratio was determined by HPLC analysis on a chiral stationary phase. n.r. No reaction; n.a. Not applicable.

**Supplementary Table 6 | Effect of the photo catalysts (PC) for the reaction 1a + 2a → 3a catalyzed by SYE2.**

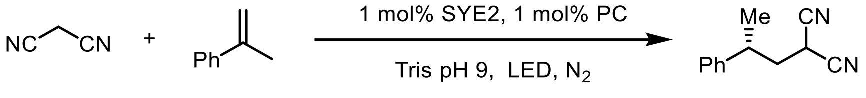

| Entry | PC | LED | Yield (%) | er (%) (R:S) |
|---|---|---|---|---|
| 1 | / | 455nm | 2.6 | 93:7 |
| 2 | FMN-Na | 455nm | 2.4 | 93:7 |
| 3 | 1 | 455nm | 10.4 | 79.8:20.2 |
| 4 | 2 | 455nm | 1.5 | n.d. |
| 5 | 3 | 455nm | n.r. | n.a. |
| 6 | 4 | 455nm | n.r. | n.a. |
| 7 | 5 | 500nm | 5.7 | 84.4:15.7 |
| 8 | 6 | 500nm | n.r. | n.a. |
| 9 | 7 (FI) | 500nm | 27.7 | 96.1:3.9 |
| 10 | 8 | 500nm | 1 | n.d. |
| 11 | 9 | 500nm | 20.4 | 94.8:5.2 |
| 12 | 10 | 540nm | n.r. | n.a. |
| 13 | 11 | 540nm | 5.1 | 95.6:4.4 |
| 14 | 12 | 540nm | 17.8 | 94.8:5.2 |
| 15 | 13 | 455nm | 0.6 | n.d. |
| 16 | 14 | 455nm | 2 | n.d. |

| | | | | |
|---|---|---|---|---|
| 17 | 15 | 455nm | n.r. | n.a. |
| 18 | 16 | 455nm | 0.9 | n.d. |
| 19 | 17 | 455nm | 0.4 | n.d. |
| 20 | 18 | 455nm | 0.9 | n.d. |
| 21 | 7 (FI) | 455nm | 27 | 95.6:4.4 |
| 22[a] | 7 (FI) | 455nm | 47.8 | 95.6:4.4 |
| 23[b] | 7 (FI) | 455nm | 45.5 | 95.6:4.4 |

Typical Conditions: **1a** (0.008 mmol), **2a** (0.004 mmol), SYE2 (1 mol%), **PC** (1 mol%) in a solvent mixture of buffer / EtOH (7.5 % v/v) were stirred for 12 hours under inert atmosphere with the irradiation of LEDs. The total volume of the reaction was 400 μL. Yield was determined by GC. Enantiomeric ratio was determined by HPLC analysis on a chiral stationary phase. [a] 5mol% FI was used. [b] 10 mol% FI was used. n.r. No reaction; n.a. Not applicable; n.d. Not determined.

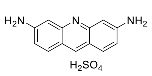

**PC1**
CAS 1811-28-5
444 nm

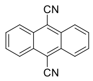

**PC2**
CAS 1217-45-4
450 nm

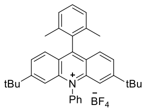

**PC3**
CAS 1810004-87-5
455 nm

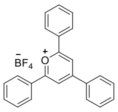

**PC4**
CAS 448-61-3
462 nm

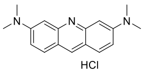

**PC5**
CAS 65-61-2
492 nm

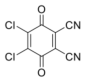

**PC6**
CAS 84-58-2
455 nm

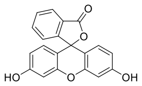

**PC7** (Fl)
CAS 2321-07-5
496 nm

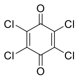

**PC8**
CAS 118-75-2
510 nm

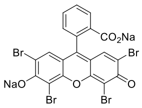

**PC9**
CAS 17372-87-1
514 nm

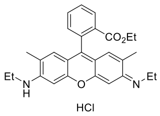

**PC10**
CAS 989-38-8
524 nm

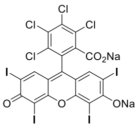

**PC11**
CAS 632-69-9
546 nm

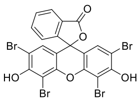

**PC12**
CAS 15086-94-9
524 nm

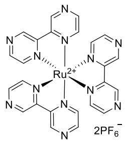

**PC13**
CAS 80907-56-8
455nm

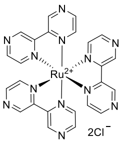

**PC14**
CAS 14323-06-9
450nm

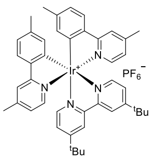

**PC15**
CAS 1607469-49-7
465nm

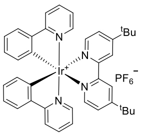

**PC16**
CAS 676525-77-2
455nm

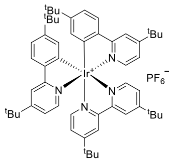

**PC17**
CAS 808142-80-5
465nm

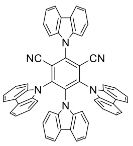

**PC18**
CAS 1416881-52-1
365nm

**Supplementary Fig. 5 | Structure, CAS number, and $\lambda_{max}$ of different photo catalysts.**

**Supplementary Table 7 | Effect of the photo catalysts (PC) for the reaction 1a + 2a → 3a catalyzed by GluER.**

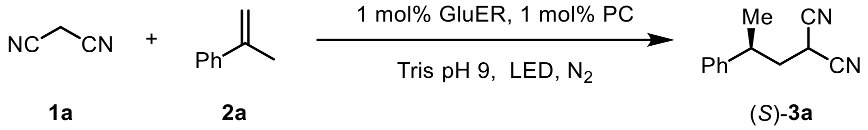

| Entry | PC | LED | Yield (%) | er (%) (R:S) |
|---|---|---|---|---|
| 1 | / | 455nm | 1.5 | 3:95 |
| 2 | FMN-Na | 455nm | 1.3 | 3:95 |
| 3 | 1 | 455nm | 18.5 | 2.5:97.5 |
| 4 | 2 | 455nm | 0.9 | n.d. |
| 5 | 3 | 455nm | n.r. | n.a. |
| 6 | 4 | 455nm | 1 | n.d. |
| 7 | 5 | 500nm | 12.5 | 3.7:96.3 |
| 8 | 6 | 500nm | n.r. | n.a. |
| 9 | 7 (FI) | 500nm | 28.8 | 1.7:98.3 |
| 10 | 8 | 500nm | 0.2 | n.d. |
| 11 | 9 | 500nm | 22.5 | 2.2:97.8 |
| 12 | 10 | 540nm | 1.3 | n.d. |
| 13 | 11 | 540nm | 4.9 | 4.8:95.2 |
| 14 | 12 | 540nm | 21.2 | 2.3:97.7 |
| 15 | 13 | 455nm | 0.3 | n.d. |
| 16 | 14 | 455nm | 0.4 | n.d. |

| | | | | |
|---|---|---|---|---|
| 17 | 15 | 455nm | 0.9 | n.d. |
| 18 | 16 | 455nm | n.r. | n.a. |
| 19 | 17 | 455nm | n.r. | n.a. |
| 20 | 18 | 455nm | n.r. | n.a. |
| 21 | 7 (FI) | 455nm | 27.8 | 1.7:98.3 |
| 22[a] | 7 (FI) | 455nm | 45.5 | 1.7:98.3 |
| 23[b] | 7 (FI) | 455nm | 42.9 | 1.7:98.3 |

Typical Conditions: **1a** (0.008 mmol), **2a** (0.004 mmol), GluER (1 mol%), **PC** (1 mol%) in a solvent mixture of buffer / EtOH (7.5 % v/v) were stirred for 12 hours under inert atmosphere with the irradiation of LEDs. The total volume of the reaction was 400 μL. Yield was determined by GC. Enantiomeric ratio was determined by HPLC analysis on a chiral stationary phase. [a] 5mol% FI was used. [b] 10 mol% FI was used. n.r. No reaction; n.a. Not applicable; n.d. Not determined.

### 6. Mechanistic Studies

#### 6.1 Proposed Catalytic Cycle

To achieve this enantioselective photoenzymatic olefin hydroalkylation reaction via single-electron oxidation of carbanions, we initially proposed the following catalytic cycle: Electron-deficient alkane **1** is firstly deprotonated in alkaline solutions to form carbanion intermediate **Int. 1**. Subsequently, flavin mononucleotide (FMNox) is excited by visible light and undergo the single electron oxidation of **Int. 1**, generating a free radical intermediate **Int. 2**, together with semiquinone state flavin cofactor FMNsq. Then **Int. 2** attacks olefins, constructing C-C bonds and generating a prochiral radical intermediate **Int. 3**. Finally, **Int. 3** is quenched by ER mediated enantioselective HAT to afford enantioenriched **3**.

In addition to the catalytic cycle we envisioned, this photoenzymatic catalysis might have the following possible pathways: First, when the EWG is carbonyl group (C=O), carbanion intermediates could rapidly attack to C=O namely nucleophilic addition instead of being oxidized by excited FMNox* (Supplementary Fig. 6, path a). Second, the visible light excited, quinone state flavin is prone to be reduced by reaction buffer or adjacent amino acids (Supplementary Fig. 6, path b). Third, once reducing power exists in the system, the free radical intermediates **Int. 2** will be rapidly reduced to the carbanion in a manner of radical polarity crossover (RPC) (Supplementary Fig. 6, path c). Finally, the equilibrium concentration of free unbound flavin in a flavoprotein solution may cause a racemic background reaction.

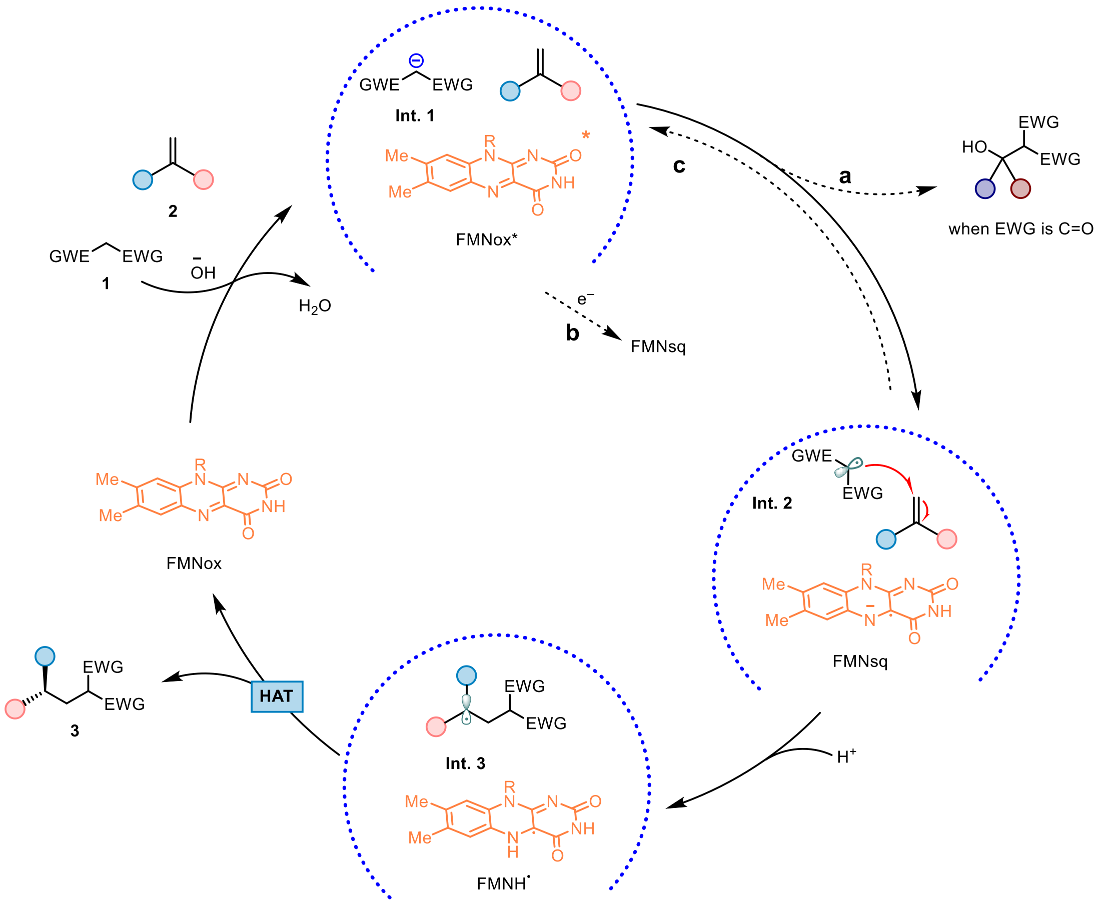

**Supplementary Fig. 6 | Designed catalytic cycle and potential undesirable pathways (dash arrows).** a. nucleophilic addition of carbanion, when EWG is carbonyl; b. reductive quenching of FMNox* by either reaction buffer or adjacent amino acids; c. radical polarity crossover.

### 6.2 UV-Vis Absorption and Luminescence Spectra

To identify the visible-light-excited species, the photophysical properties of the ene-reductases and external photocatalyst fluorescein (FI, PC7) were measured. With a Synergy H1 microplate reader from BioTek, several absorption spectra were recorded. First, the yellow protein SYE2 (0.02 mM in Tris buffer/glycerol mixture, related to the reaction conditions) were recorded (Supplementary Fig. 7, blue solid curve). A strong absorption band among the visible light region with a maximum absorption wavelength at 460 nm indicates the SYE2 bound FMN quinone state. Then, spectral data of the free FMN-Na were recorded (Supplementary Fig. 7, black dash curve). In addition, spectral data of FI were also recorded (Supplementary Fig. 7, red solid curve). A strong absorption band among the visible light region with a maximum absorption wavelength at 490 nm indicates the SYE2 bound excited state FI*. The absorbance spectra of SYE2 with the addition of substrates **1a** and/or **2a** were recorded too. As shown in Supplementary Fig. 9, the addition of substrates has little influence on the spectra. In addition, spectral data of the mixture of SYE2 and FI were recorded, giving a similar spectrum. Overall, SYE2 and FI absorbs visible light at 400-520 nm region with a maximum absorption at 490 nm, slightly red-shifted and enhanced compared with free FMN-Na.

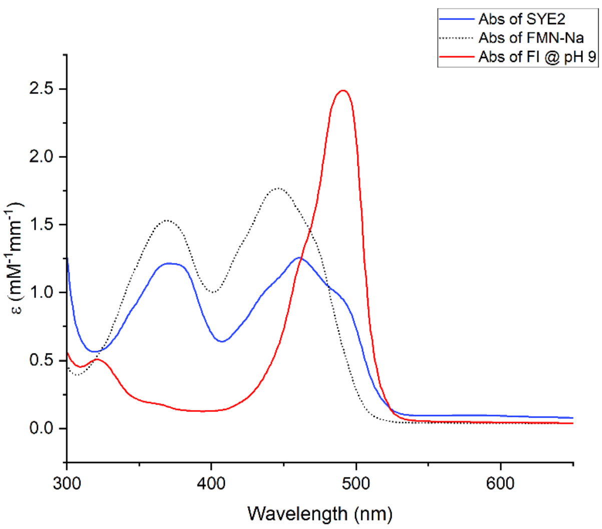

**Supplementary Fig. 7 | UV-Vis absorption spectra.** 1) The absorbance spectra of SYE2 (blue solid curve); 2) The absorbance spectra of FMN-Na (black dash curve); 3) The absorbance spectra of FI (red solid curve).

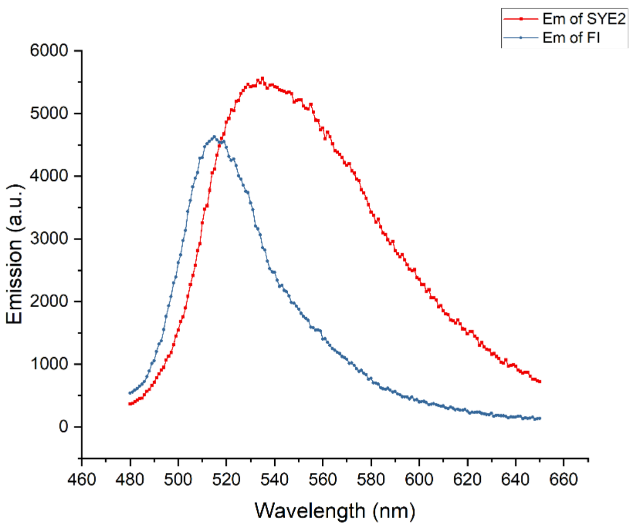

**Supplementary Fig. 8 | UV-Vis luminescence spectra.** 1) The luminescence spectra of SYE2 (red solid curve); 2) The luminescence spectra of FI (cyan solid curve).

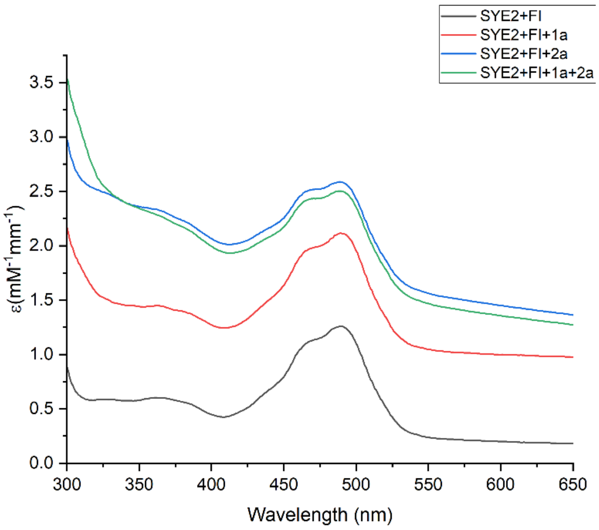

**Supplementary Fig. 9 | UV-vis absorbance spectra of SYE2, FI with substrates.** 1) SYE2+FI (SYE2 + FI curve, black solid curve); 2) SYE2, FI with the addition of 1a (SYE2 + FI + **1a** curve, red solid curve); 3) SYE2, FI with the addition of **2a** (SYE2 + FI + **2a** curve, blue solid curve); 4) standard conditions (SYE2 + FI + **1a** + **2a** curve, green solid curve).

**Conclusion**: The results support that the ene-reductase bound FI is excited with blue LEDs and turns to a highly oxidizing species FI*, which then initiates the reaction.

#### 6.3 Stern-Volmer Luminescence Quenching Studies

To identify the key species that interacts with the visible-light-excited ene-reductase bound FI, Stern-Volmer luminescence quenching experiments were designed and conducted. The photoenzyme, potential quenchers and Tris buffer (50 Mm, pH = 9) were weighed into vials inside a glove box under nitrogen atmosphere. The emission spectra were recorded using a Synergy H1 microplate reader from BioTek. SYE2 was excited at 460 nm and the emission intensity was collected at 530 nm. In a typical experiment, to a 20 μM solution of SYE2 in Tris buffer (50 mM, pH = 9) and 15 ppm FI in Tris buffer (50 mM, pH = 9) was added the appropriate amount of **1a**, in a quartz ELISA Plate sealed with 3M adhesive tape. Then, the emission of the sample was collected. As shown in Supplementary Fig. 10, the measured luminescence intensity of ER3 decreases with the addition of **1a** rather than **2a**.

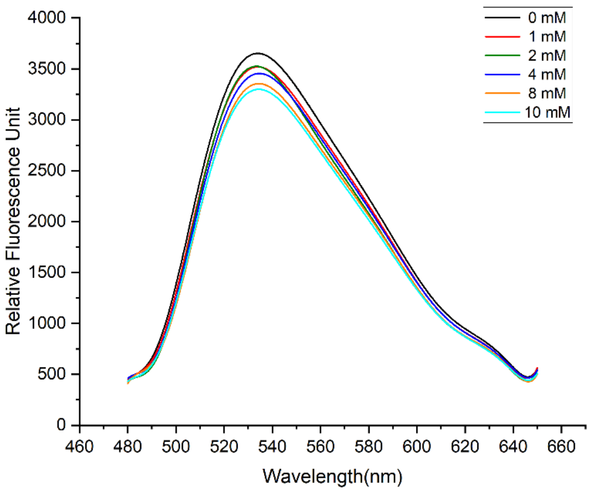

**Supplementary Fig. 10 | Stern-Volmer luminescence quenching studies.**

**Conclusion**: These results support a reductive quenching pathway that excited FI* was single electron reduced by substrate **1a**.

#### 6.4 Determination of Oxidation Potential

To verify that electron-deficient malononitrile cannot be directly oxidized under this photoenzyme synergistic catalytic system, we measured the oxidation potential of **1a** under the same conditions as previously reported[10] for the oxidation potential of α-methylstyrene. Electrochemical potentials were obtained with a standard set of conditions to maintain internal consistency. Cyclic voltammograms were performed on a CH Instruments Electrochemical Workstation model CHI760E at room temperature under anaerobic conditions and results were shown in Supplementary Fig. 11 and Fig. 12. Samples were prepared with 0.05 mmol of substrate in 5 mL of 0.1 M tetrabutylammonium hexafluorophosphate in dry, degassed acetonitrile. Measurements employed a glassy carbon working electrode, a Pt counter electrode, and an Ag/AgCl reference electrode, and a scan rate of 100 mV/s.

$E_{p/2}$=2.15V vs Ag/AgCl (2.63 vs SCE)

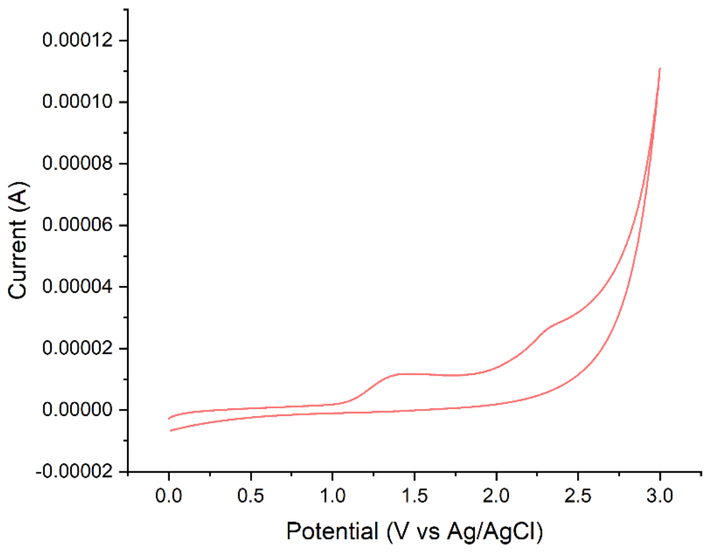

**Supplementary Fig. 11 | Cyclic voltammetry curve of 1a.**

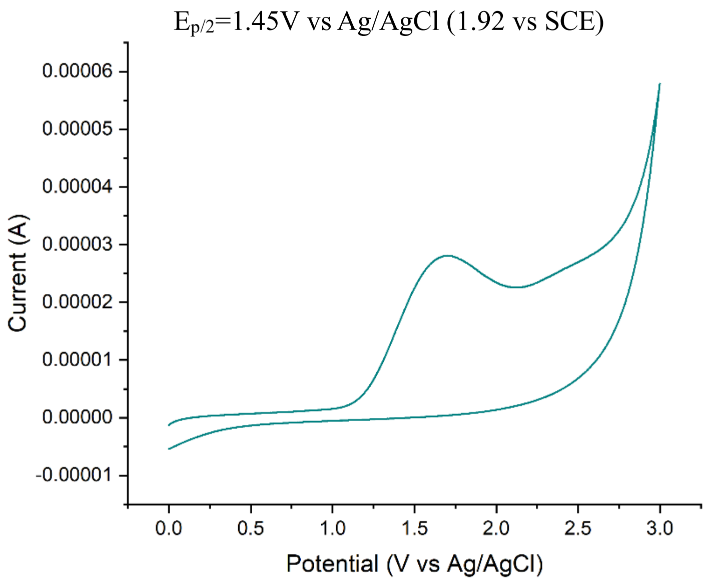

**Supplementary Fig. 12 | Cyclic voltammetry curve of 2a.**

**Conclusion**: The oxidation potential of **2a** we actually measured is consistent with previously reported (1.91V vs SCE)[10]. The oxidation potential of **1a** is 2.63 V (vs SCE), which proves that both excited FI*and FMNox* cannot directly oxidize **1a**.

### 6.5 Calculation Results of Oxidative Potential and Gibbs Free Energy

To illustrate the role of exogenously added photocatalysts, we selected some photocatalysts and calculated their oxidation potentials and Gibbs free energy, and the results were shown in Supplementary Fig. 13-16.

All DFT calculations were performed with the Gaussian 16, Revision A.03 software 27. The initial structures were first optimized in conjunction with the SMD continuum solvation model at the B3LYP-D3/6-31G(d) level of theory. The solvation energies were further calculated at the M052X/6-31G* for all atoms. In particular, the solvent of water was used to simulate the reaction system.

The initial structures were first optimized at B3LYP/6-31G(d) level. Then, the single-point TDDFT calculations at wB97XD/6-311+G(d,p) levels were performed to compute the three low-lying singlet excited states. All TDDFT calculations were performed with Gaussian 16, Revision A.03.

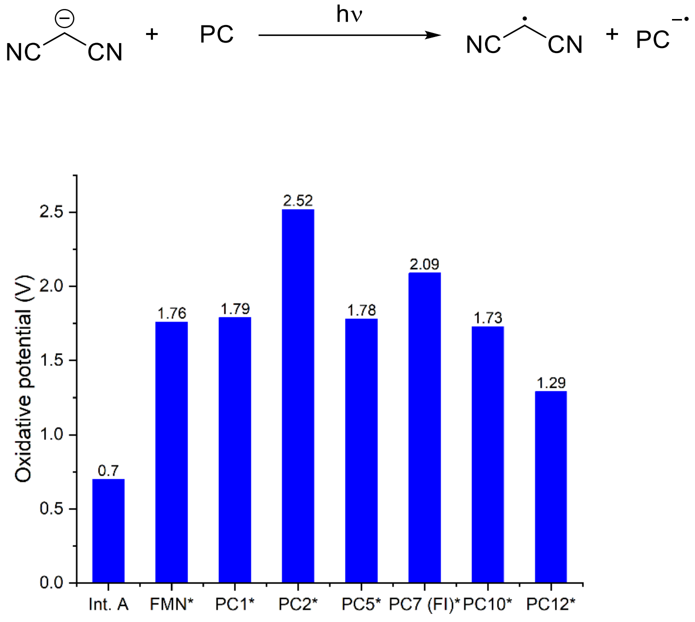

**Supplementary Fig. 13 | The calculated oxidative potentials of excited state photocatalysts.**

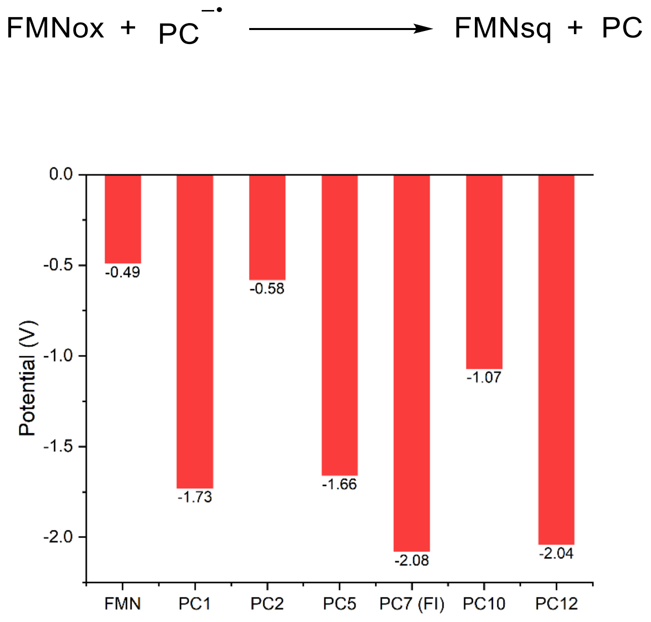

**Supplementary Fig. 14 | The calculated potentials of electron transfer between FMNox with photocatalyst radical anions**

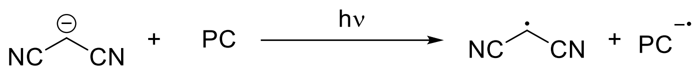

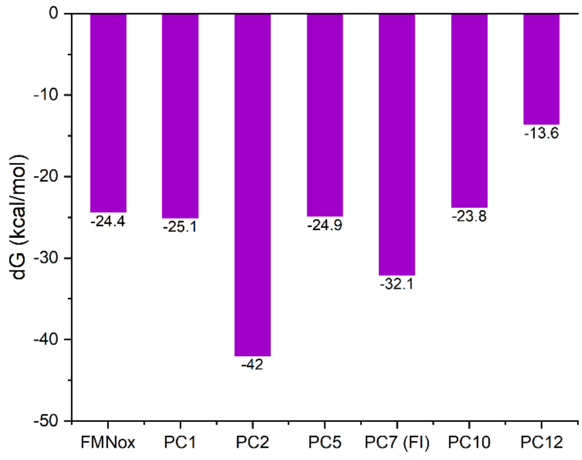

**Supplementary Fig. 15 | The calculated Gibbs free energy changes (dG) of excited state photocatalysts with Int. A.**

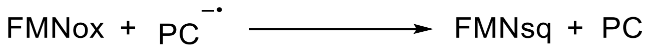

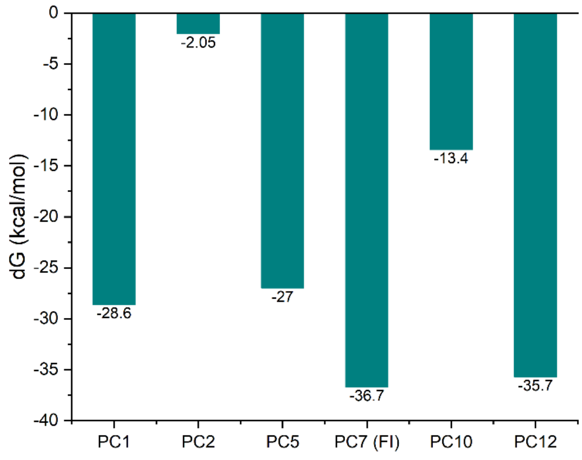

**Supplementary Fig. 16 | The calculated Gibbs free energy changes (dG) of electron transfer between FMNox with photocatalyst radical anions.**

**Conclusion:** The single electron oxidation of carbanions by excited FI and the electron transfer between FI radical anions and FMNox are both thermodynamically favorable processes.

### 6.6 The Roles of Protein-bound FMN by Preparation of Apo Flavoprotein

To explore the roles of FMN and externally added photocatalyst FI, we conducted the following control experiments (as shown in Supplementary Table 8) to verify whether the reaction undergoes photocatalytic, FMN-mediated single electron transfer.

To obtain the apo flavoprotein SYE2, we used a previously reported method[11] to remove the cofactor FMN that is endogenously expressed and bound to the protein. By using the typical procedure described in Section 2, after the expression and lysis of SYE2, no additional FMN-Na was added. After loading the protein solution at a flow rate of 0.2 ml/min, the column outlet was connected with the pump inlet, creating a loop to ensure maximal binding of protein to the column. After overnight circulation (0.2 ml/min) at 4 °C in the dark, the loop was disconnected. Protein-bound FMN was then removed by washing the column with 1500 mL Tris buffer (20 mM, pH 7.5, containing 0.5 M NaCl, 2 M KBr and 2 M urea), resulting in a column-bound apo form of SYE2. The column was washed with 10 column volumes of a 20 mM Tris buffer pH 7.5 containing 0.5 M NaCl and 10 mM imidazole. Finally, the apo-SYE2 was eluted by elution buffer (20 mM Tris, containing 0.5 M NaCl, 0.5 M imidazole), and the purified apo-SYE2 was shown in Supplementary Fig. 17.

**Supplementary Table 8 | Control experiments of the reaction for 1a + 2a → 3a catalyzed by apo-SYE2.**

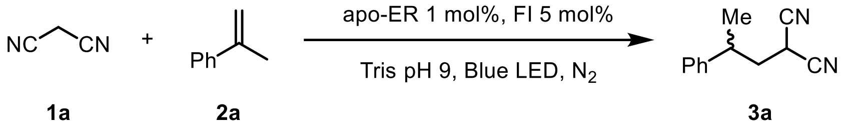

| Entry | ER | FMN-Na | Yield (%) |
|---|---|---|---|
| 1 | Apo-SYE2 | / | 1.2 |
| 2 | Apo-SYE2 | 10 mol% | 19.8 |

Typical Conditions: **1a** (0.008 mmol), **2a** (0.004 mmol), apo-SYE2 (1 mol%), Fl (5 mol%) in a solvent mixture of buffer / EtOH (7.5 % v/v) were stirred for 12 hours under inert atmosphere with the irradiation of blue LEDs; total volume of the reaction was 0.4 mL. Yield was determined by GC.

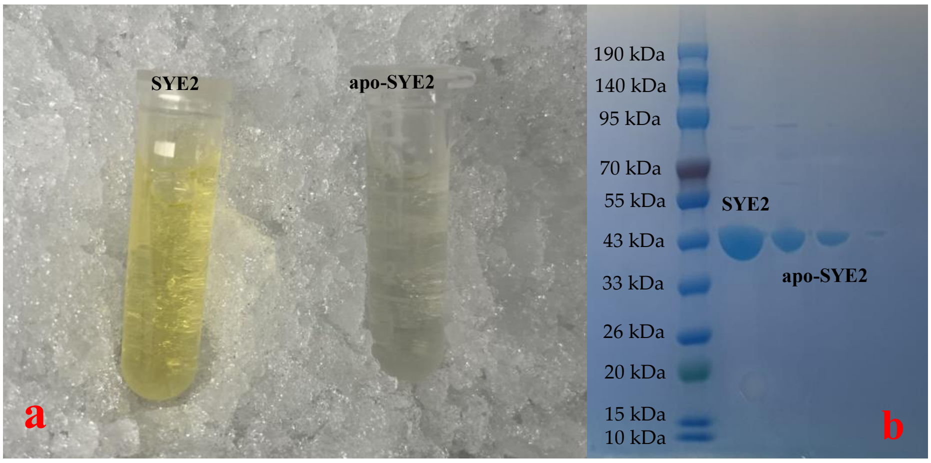

**Supplementary Fig. 17 | Pictures of SYE2 and aop-SYE2. a**. The purified enzymes of SYE and apo-SYE2; **b**. SDS-PAGE of SYE2 and apo-SYE2.

**Conclusion**: The catalytic cycle requires the participation of FMN to proceed successfully, and FMN may participate in the electron transfer between the radical anion and the ground state photocatalyst.

#### 6.7 Electron Paramagnetic Resonance (EPR) Spin Trapping Experiments

To explore the relatively short-lived substrate-derived radical intermediates, room temperature EPR spin trapping experiments were designed and performed with the addition of DMPO (5,5-dimethyl-1-pyrroline-*N*-oxide) as spin trap.

EPR spin-trapping experiments were carried out at room temperature using a Bruker X-band E500 spectrometer equipped with a super-high sensitivity ER4122SHQE resonator. The experimental parameters were set to 0.2 mW of incident microwave power, 100 kHz field modulation and 1 G modulation amplitude. For each sample, 20 scans were taken to increase the signal-to-noise ratio.

(1) When the reaction was performed without substrates, no free radical EPR signal was observed. (Supplementary Fig. 18, entry 1).

(2) When the reaction was performed without SYE2, compared with the standard conditions, free radical signal barely changes. (Supplementary Fig. 18, entry 3).

(3) When the reaction was carried out under standard conditions, there were obvious free radical signals of carbon-centred radical. (Supplementary Fig. 18, entry 5).

(4) In order to better understand the substrate that produce free radical signal, we designed the corresponding control experiments. As shown in the Supplementary Fig. 16, with the addition of DMPO as free radical spin-trapping agent, EPR signals were significantly enhanced at room temperature when **1a** instead of **2a** was added (entry 2 and 4).

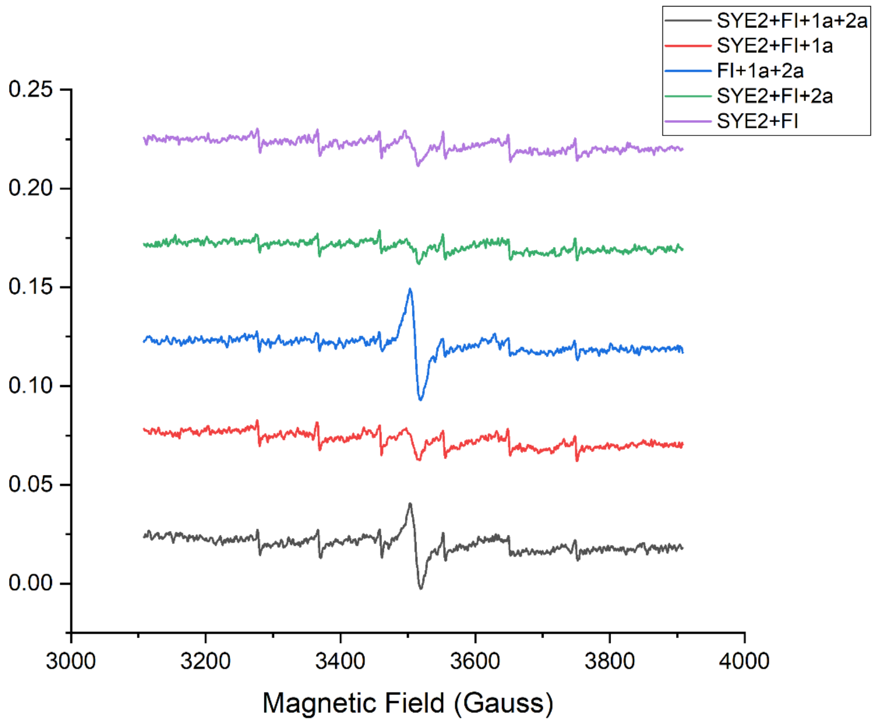

**Supplementary Fig. 18 | Room temperature EPR spin trapping experiments.**

**Conclusion**: These results indicate the formation of a possible **1a**-derived carbon-centered radical being consistent with quenching experiments.

### 6.8 TEMPO Trapping Experiment

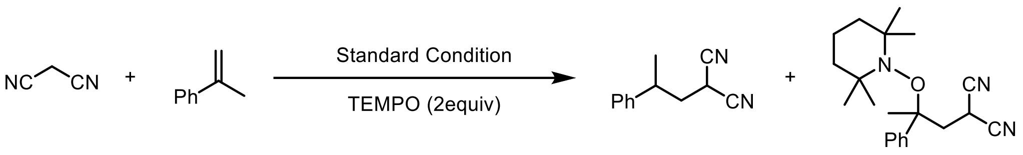

**1a** **2a** **3a,** trace detected by HRMS

To gain insight into the reaction mechanism, 2 equiv. of 2,2,6,6-tetramethylpiperidine-1-oxyl (TEMPO) was added to the reaction mixture of **1a** and **2a** under otherwise standard conditions and only trace amounts of **3a** was detected with the prochiral radical intermediate being trapped by TEMPO, which was detected by high-resolution mass spectrometry (HRMS). HRMS (ESI) calculated for $C_{21}H_{30}N_3O$ [M+H$^+$]: 340.2383, found: 340.2389.

UPLC-HRMS mode: I-class Synapt XS mobile phase A: $H_2O$ (0.1% FA); B: CAN; column temperature 40 degrees; C18 column: 1.8 um, 2.1mm*50mm.

TIC and EIC@340.2389Da:

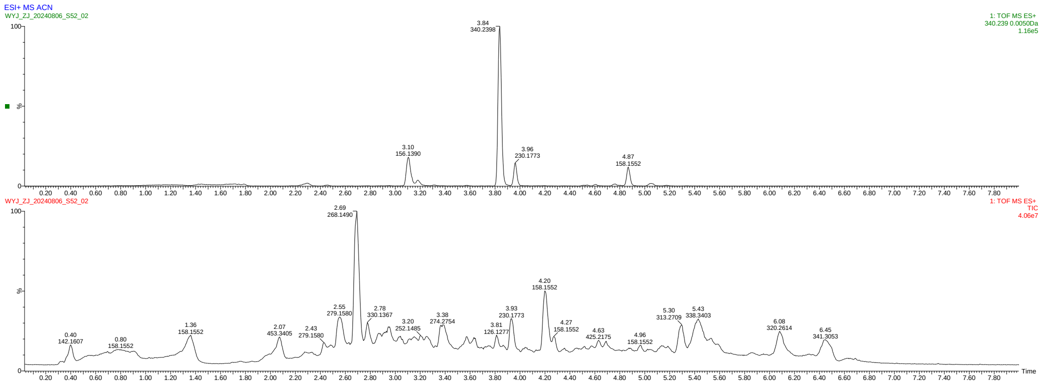

HRMS@3.838 min:

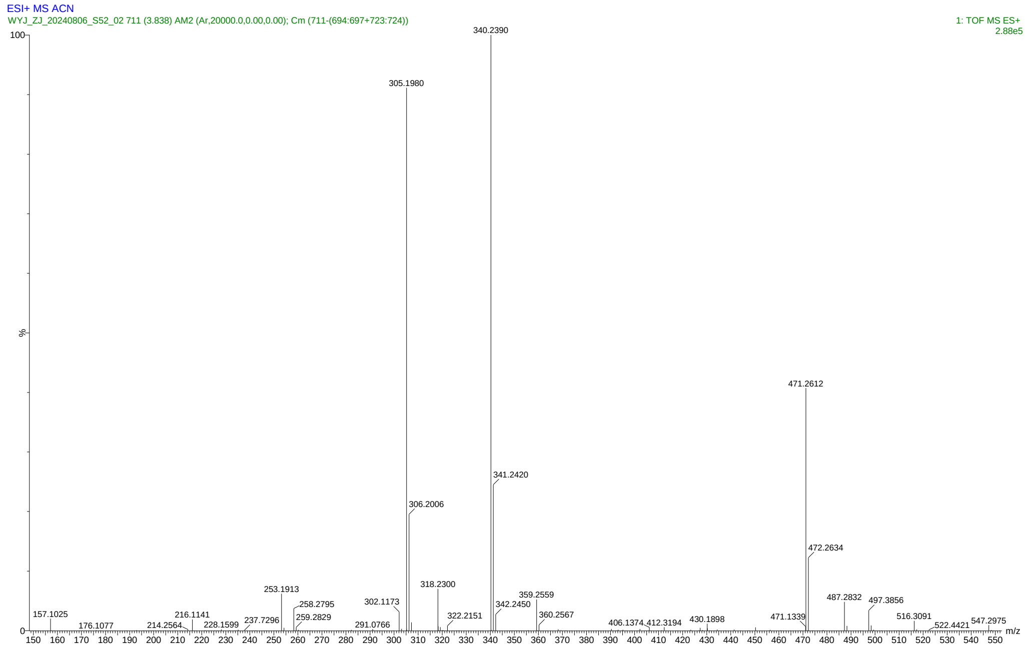

Results of predicted MS and actual measured MS:

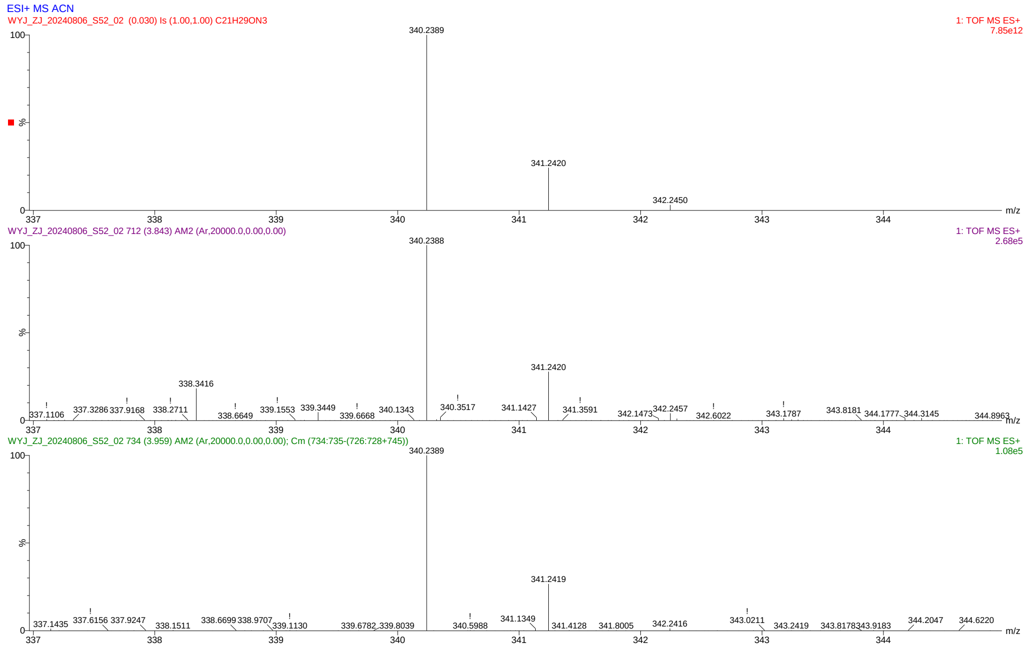

### 6.9 Radical Clock Experiment

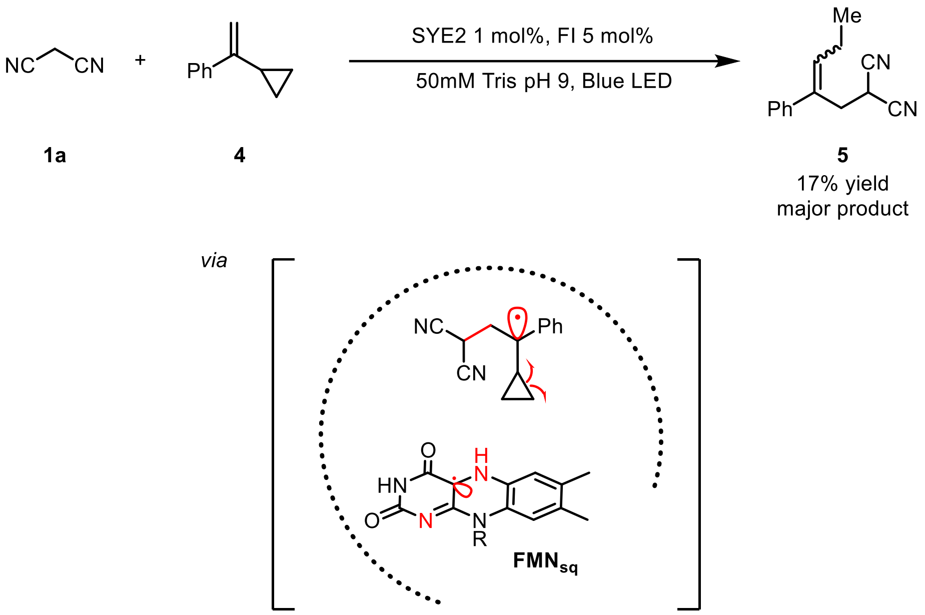

To probe the prochiral radical intermediate, a radical clock experiment was designed and performed.

By using the typical procedure described in Section 5, the reaction of malononitrile **1a** (0.008 mmol) and (1-cyclopropylvinyl)benzene **4** (0.004 mmol) catalyzed by SYE2 (1 mol%) afforded 2-(2-phenylpent-2-en-1-yl)malononitrile **5** in 17% GC yield (average of duplicate runs) as the major product. *E/Z* ratio was determined by GC-MS as 2:1.

**Conclusion**: The result from the radical clock experiment supports the generation of prochiral radical intermediate.

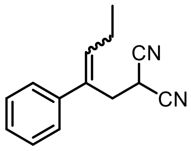

**5**

2-(2-phenylpent-2-en-1-yl) malononitrile **5**.

**5** is a colorless oil.

$^1$H NMR (600 MHz, $CDCl_3$) δ 7.36 – 7.27 (m, 3H), 7.13 – 7.04 (m, 2H), 5.75 (t, *J* = 7.4 Hz, 1H), 3.42 (t, *J* = 7.8 Hz, 1H), 2.95 (d, *J* = 7.9 Hz, 2H), 1.95 (p, *J* = 7.5 Hz, 2H), 0.90 (t, *J* = 7.5 Hz, 3H).

$^{13}$C NMR (151 MHz, $CDCl_3$) δ 136.87, 135.92, 131.05, 127.97, 127.42, 127.11, 125.59, 111.24, 39.75, 21.40, 21.02, 13.03.

GC calibration curve for **5** is displayed below:

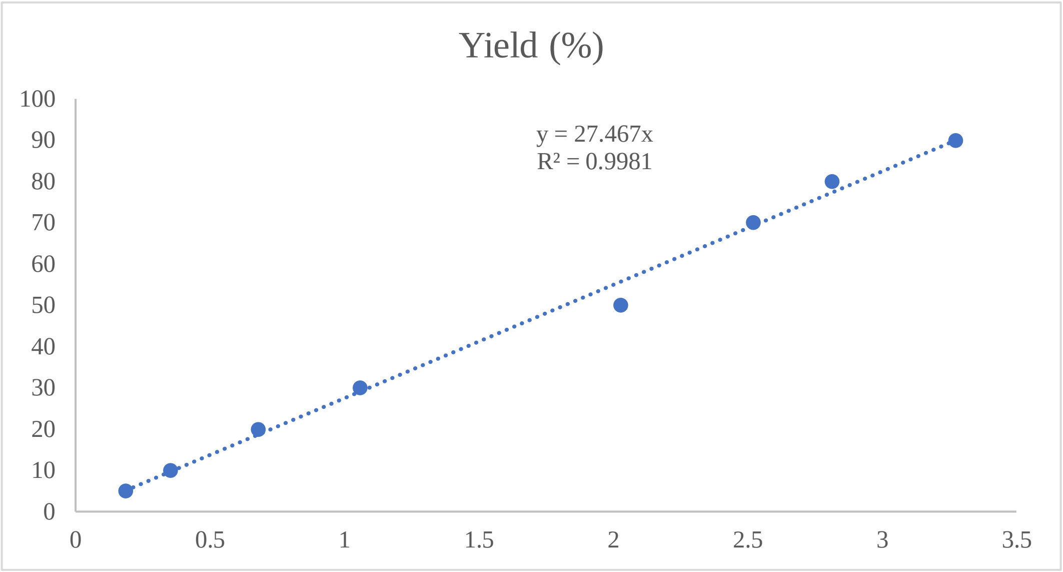

GC-MS spectra of the crude mixture of the run_1, which confirms the formation of **5** as the major product and determines a *Z/E* ratio of 2:1.

GC-MS trace of the reaction mixture catalyzed by SYE2:

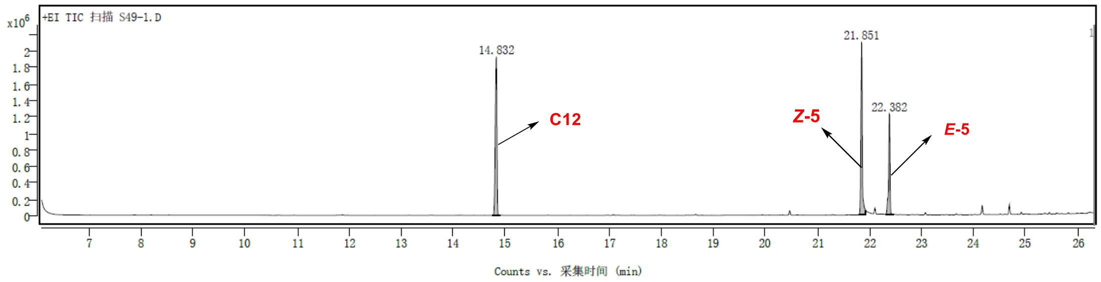

GC-MS trace of **5**:

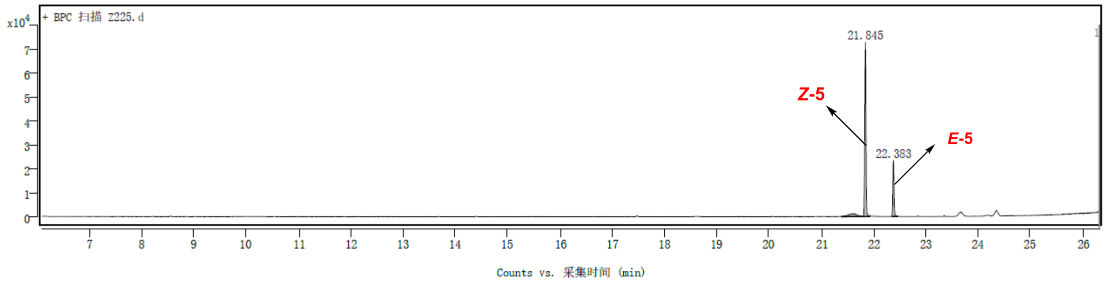

MS spectrum of the compound at 21.851 min (*Z*-**5**):

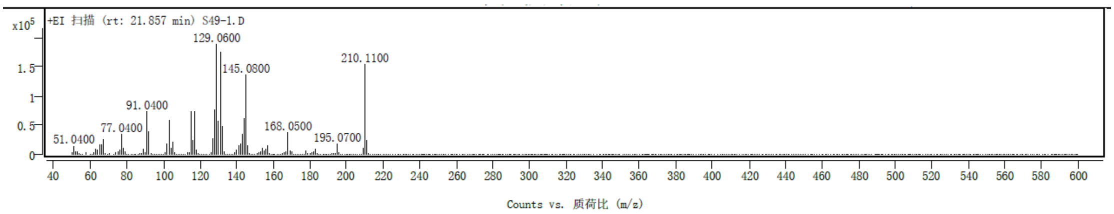

MS spectrum of the compound at 22.382 min (*E*-**5**):

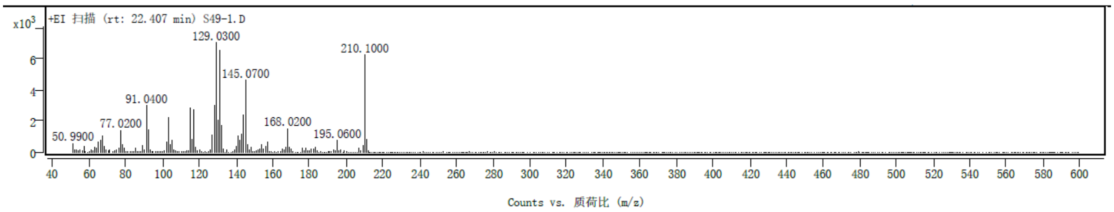

### 6.10 Isotopic Labelling Experiments

To get a better understanding of the terminating HAT step, the reaction of **1a** + **2a** → **3a** was conducted using a deuterated Tris buffer, duplicate was set-up in parallel and the deuterated product *d*-**3a** was obtained in a lower yield of 12% which indicates HAT process might be rate-limiting, and the newly-formed stereocenter in compound *d*-**3a** was partially labelled with deuterium (76% D).

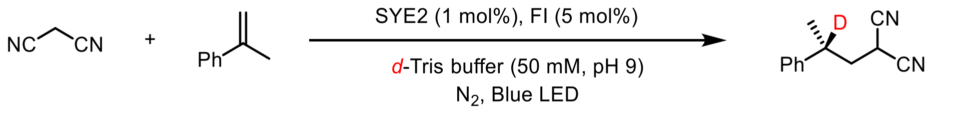

$^1$H NMR traces of the deuterium labelling experiments were displayed in Supplementary Fig. 19.

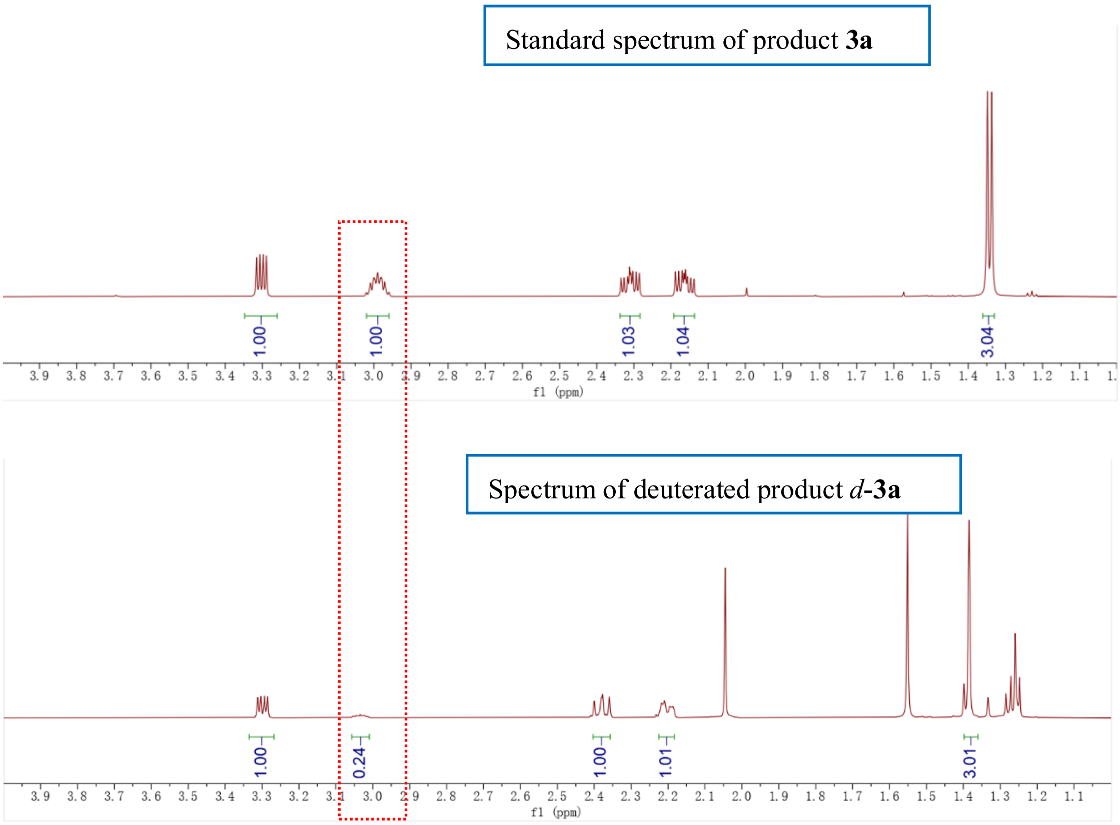

**Supplementary Fig. 19 | $^{1}$H NMR trace of the deuterium labelling experiments.**

Deuterated Tris buffer was prepared by dissolving Tris (Hydroxymethyl) Aminomethane Hydrochloride (Tris-HCl) in $D_2O$ (50 mM) and adjusting pH to 9 with deuterium sodium hydroxide (1 M in $D_2O$).

### 7. Electronic Structure Calculations

#### 7.1 System Setup and MD Simulations

The initial wild-type structure of GluER, was constructed based on the crystal structure (PDB code: 6O08).[12] The SYE2 enzyme structure was modeled using AlphaFold2,[13] and the FMN position was confirmed by superimposing it with the GluER enzyme. Then, the **Int. C** and FI were docked into the active site using AutoDock Vina tool,[14] we assigned the protonation states of titratable residues (His, Glu, Asp) based on pKa values from the PROPKA software[15,16] in combination with a careful visual inspection of local hydrogen-bonded networks. Notably, the phosphate groups of FMN were assigned to be fully deprotonated. For GluER enzyme, His89, His101, His107 and His172 were protonated at the ε position, His 14 and His 189 was protonated at the δ position, His128 were both protonated at ε position and δ position, all the Asp and Glu residues were deprotonated. For SYE2 enzyme, His8, His87, His99, His105, His175, His178, His231 and His288 were protonated at the ε position, His 263, His 308 and His 331 were protonated at the δ position, all the Asp and Glu residues were deprotonated. The Amber ff14SB force field[17] was employed for the protein residues, while the general AMBER GAFF force field[18] was used for **Int. C**, FI and FMNHsq. The partial atomic charges of **Int. C**, FI and FMNHsq were obtained from the RESP calculations at the B3LYP/6-31G(d,p) level of theory. Sodium ions were added to the protein surface to neutralize the total charge of the systems. Finally, the resulting system was solvated in a rectangular box of TIP3P[19] waters extending up to a minimum distance of 16 Å from the protein surface.

After proper setup, the whole system was fully minimized using a combined steepest descent and conjugate gradient method. Then, the system was gently annealed from 0 to 300 K under canonical ensemble for 50 ps with a weak restraint of 25 kcal/mol/Å on protein. To achieve a uniform density after heating dynamics, 1 ns of density equilibration was performed under the NPT ensemble at the target temperature of 300 K and target pressure of 1.0 atm. Afterward, we performed 100 ns MD simulations under the NPT ensemble. All MD simulations were performed with GPU version of Amber 18 package.[20]

### 7.2 QM/MM Calculations for Enzymatic Reactions

A representative snapshot extracted from the clustering results in the MD trajectory was used for subsequent QM/MM calculations.[21,22] All QM/MM calculations were performed using ChemShell[23,24] combining Turbomole[25] for the QM region and DL_POLY[26,27] for the MM region. The electronic embedding scheme[28] was used to account for the polarizing effect of the enzyme environment on the QM region. Hydrogen link atoms with the charge-shift model were applied to treat the QM/MM boundary. For whole reactions, the QM region consists of the FMNHsq and **Int. C**. In all QM/MM geometry optimizations, the QM region was treated with the B3LYP functional using the double-zeta basis set def2-SVP[29] for all atoms (labeled as B1). The energies were further corrected with the larger basis set def2-TZVP for all atoms (labeled as B2). Grimme's D3BJ empirical dispersion was included in all calculations.[30,31]

### 7.3 DFT Calculations

All DFT calculations were performed with the Gaussian 16,[32] Revision A.03 software. The initial structures were first optimized in conjunction with the SMD continuum solvation model at the B3LYP-D3/6-31G(d) level of theory.[33,34] The solvation energies were further calculated at the M052X/6-31G* for all atoms. In particular, the solvent of water was used to simulate the reaction system.

### 7.4 TDDFT Calculations

The initial structures were first optimized at B3LYP/6-31G(d) level. Then, the single-point TDDFT calculations at wB97XD/6-311+G(d,p) levels were performed to compute the three low-lying singlet excited states. All TDDFT calculations were performed with Gaussian 16, Revision A.03.[32]

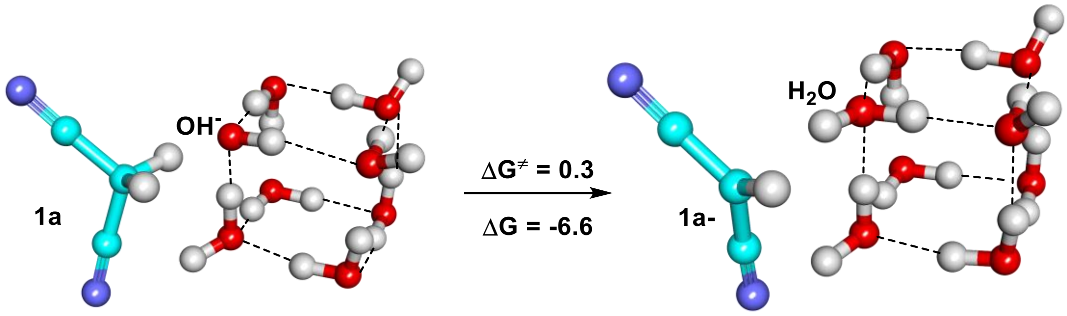

**Supplementary Fig. 20 | QM(B3LYP-D3/6-311+G(d)) calculated Gibbs free energy for the deprotonation of 1a in water solution with hybrid cluster-continuum (HCC) model.** Our calculated results show that **1a** was easily deprotonated by $OH^-$ in the solvent to generate the anionic **1a** (**1a**$^-$) involving a low energy barrier of 0.3 kcal/mol.

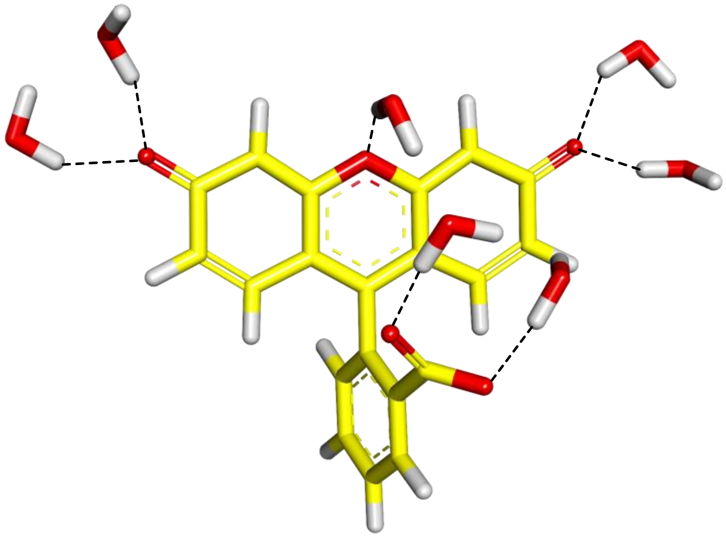

**λ = 465.2**

**Supplementary Fig. 21 | Maximum absorption wavelength of fluorescein calculated by using TDDFT (B3LYP-D3/6-311+G(d)) method.** In pH = 9, the maximum absorption wavelength of divalent anion fluorescein (FI) is 465.2 nm.

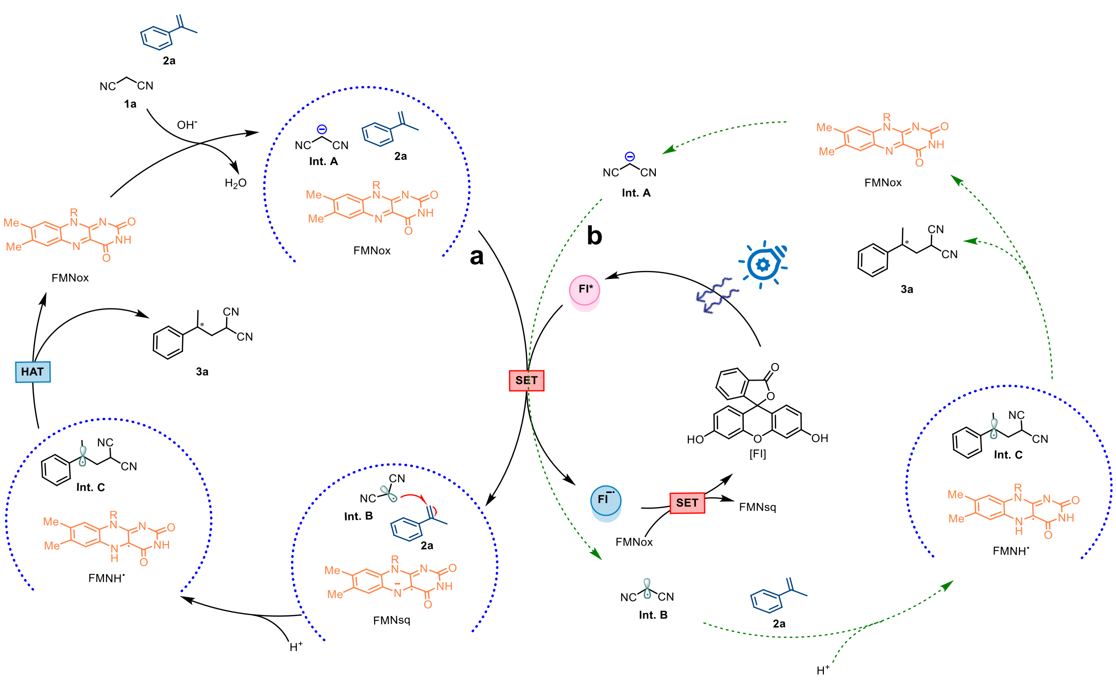

**Supplementary Fig. 22 | Proposed catalytic cycle. a**, SET oxidation of carbanions occurs within enzyme's active site (black solid curve). **b**, SET oxidation of carbanions locates outside the enzyme active site and the resulting prochiral radicals **Int. C** diffuse into the active site (green dash curve).

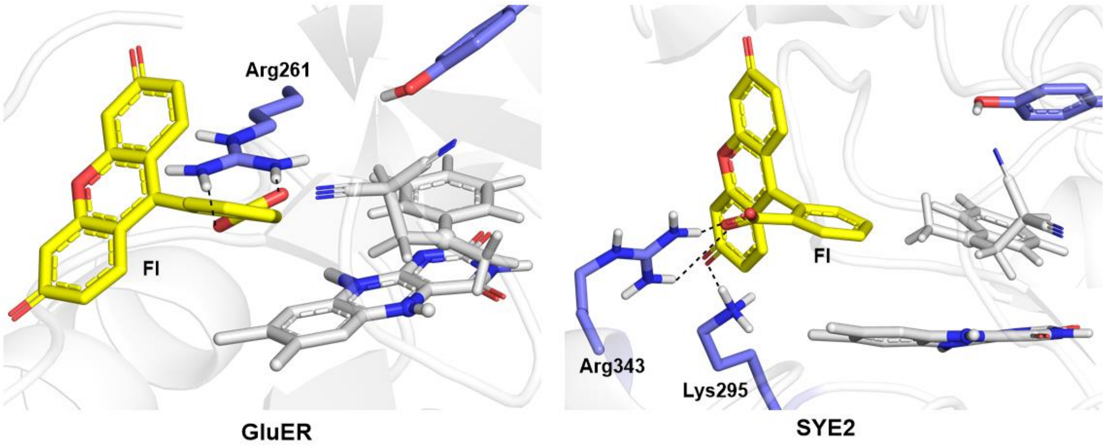

**Supplementary Fig. 23 | Representative conformations of FI in GluER and SYE2 enzymes in MD trajectories.** The results of MD simulations showed that FI was stabilized by the Arg261 residue within the active pocket of GluER and by the Arg343 and Lys295 residues in SYE2.

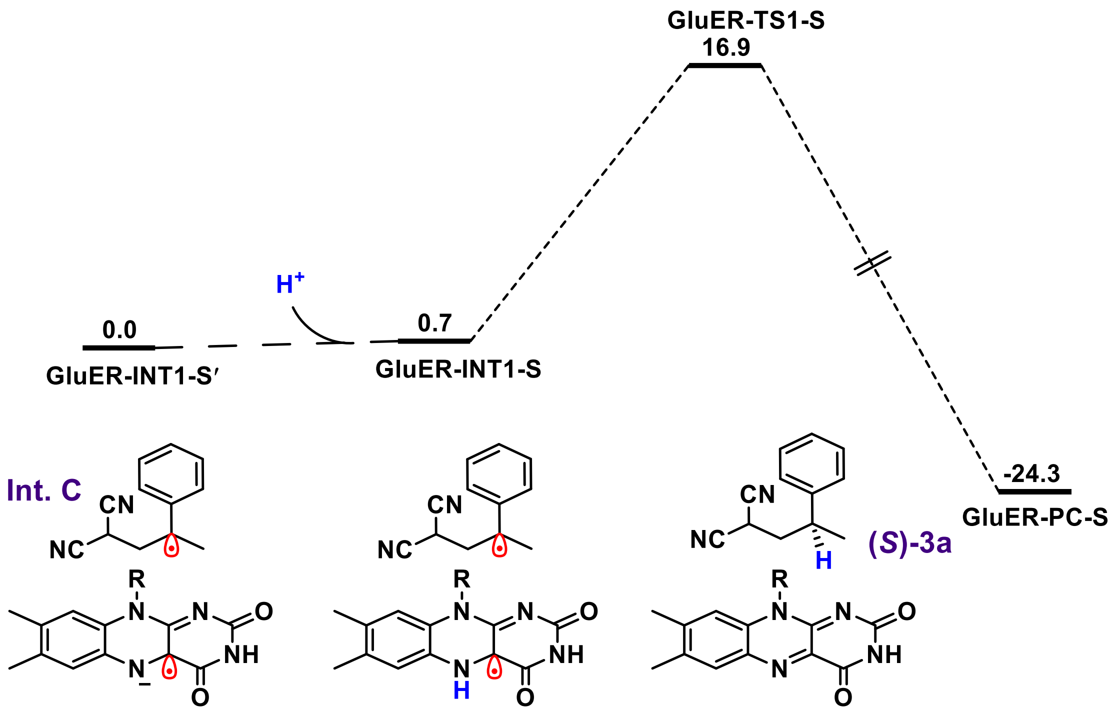

**Supplementary Fig. 24 | Calculated pathway from $CH_3$-right conformation of Int. C to (*S*)-3a catalyzed by GluER.** The QM(B3LYP-D3/B2)/MM calculated energy profile (in kcal/mol) for the generation of the product (*S*)-**3a** from the $CH_3$-right conformation of **Int. C**.

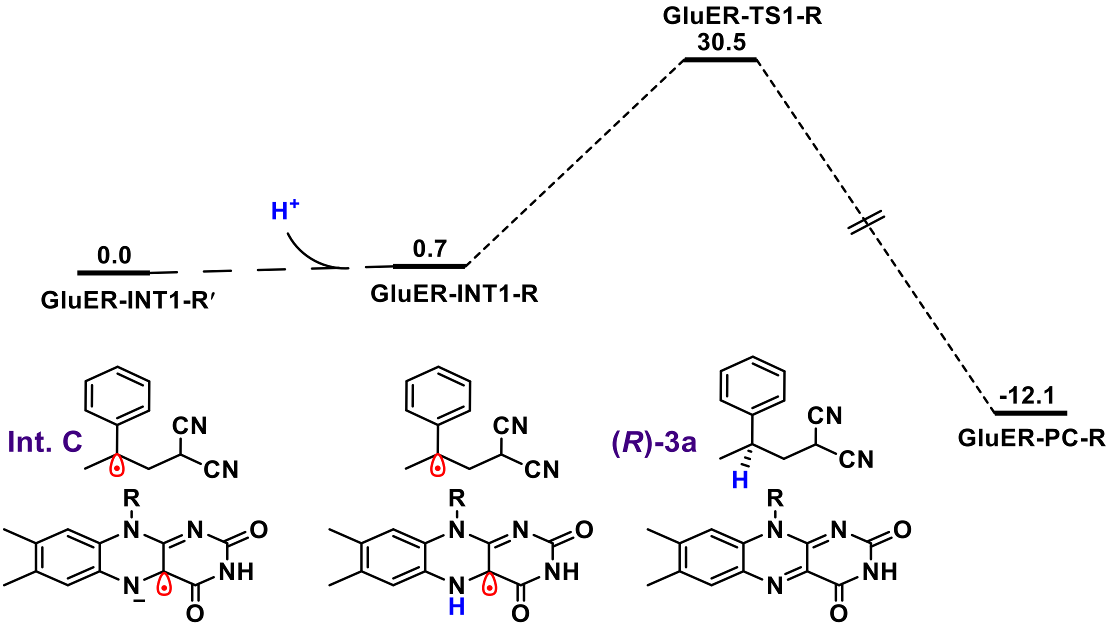

**Supplementary Fig. 25 | Calculated pathway from $CH_3$-left conformation of Int. C to (*R*)-3a catalyzed by GluER.** The QM(B3LYP-D3/B2)/MM calculated energy profile (in kcal/mol) for the generation of the product (*R*)-**3a** from the $CH_3$-left conformation of **Int. C**.

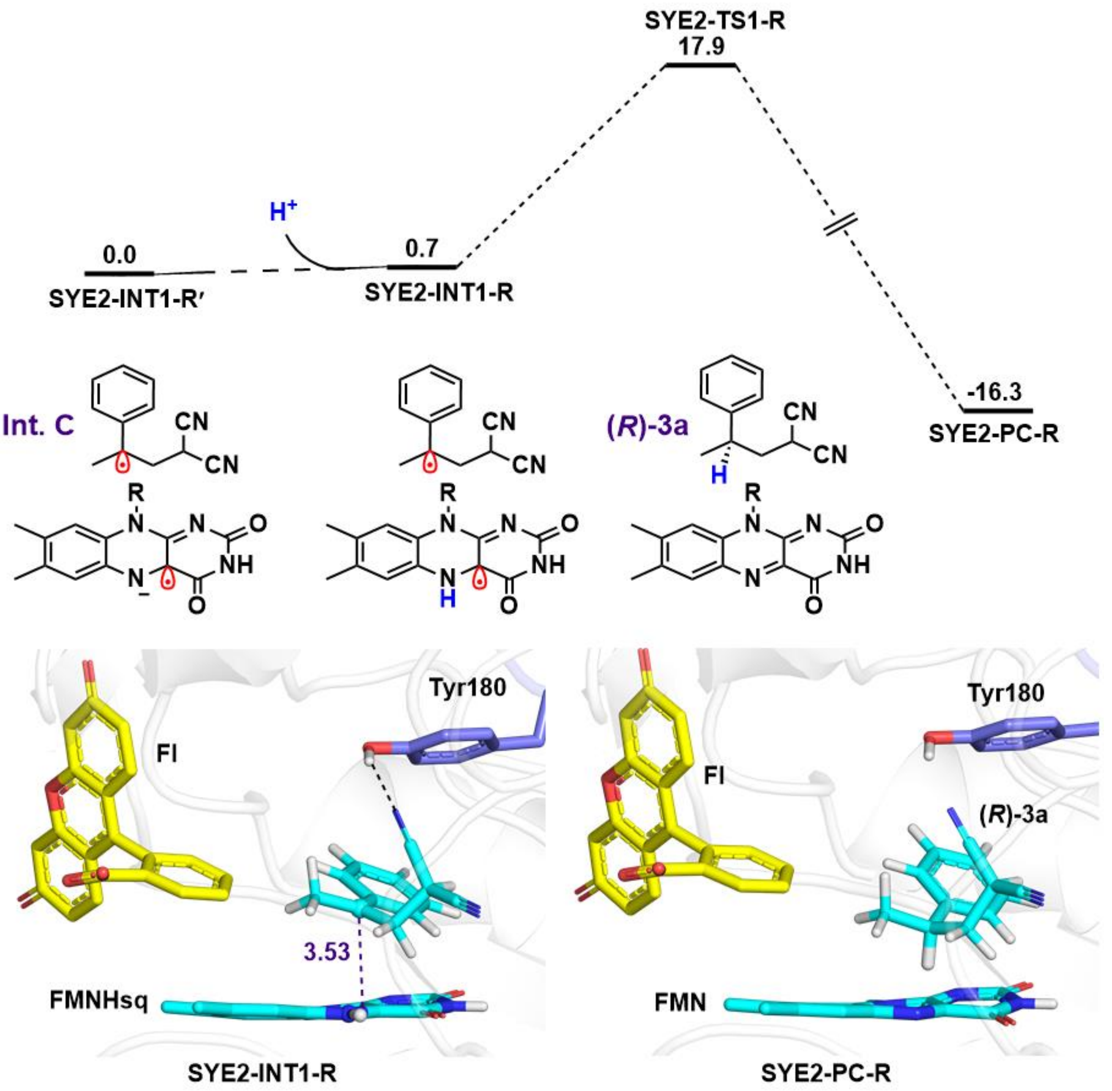

**Supplementary Fig. 26 | Calculated pathway from $CH_3$-left conformation of Int. C to (*R*)-3a catalyzed by SYE2.** The QM(B3LYP-D3/B2)/MM calculated energy profile (in kcal/mol) for the generation of the product (*R*)-**3a** from the $CH_3$-left conformation of **Int. C**.

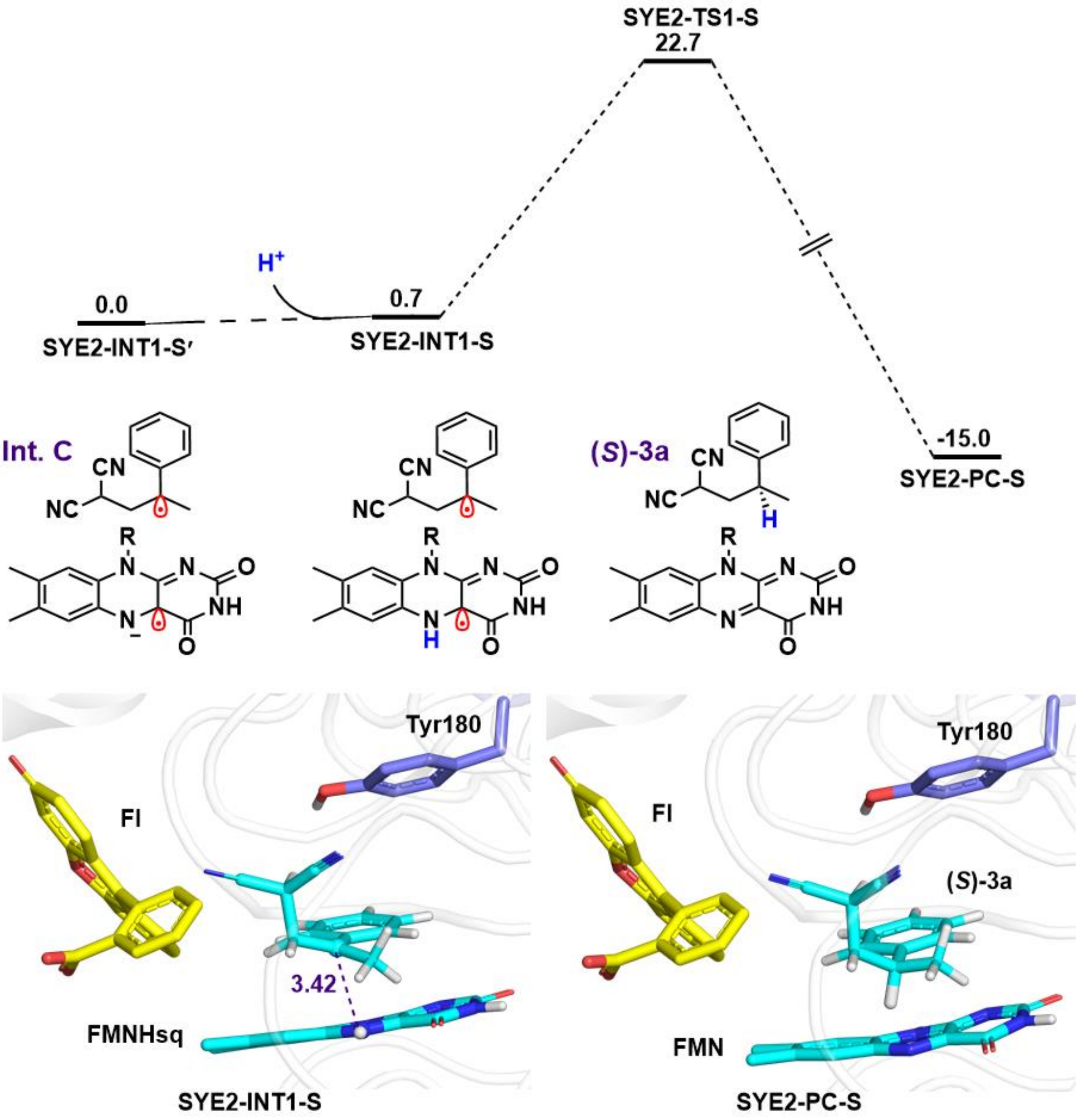

**Supplementary Fig. 27 | Calculated pathway from $CH_3$-right conformation of Int. C to (*S*)-3a catalyzed by SYE2.** The QM(B3LYP-D3/B2)/MM calculated energy profile (in kcal/mol) for the generation of the product (*S*)-**3a** from the $CH_3$-right conformation of **Int. C**.

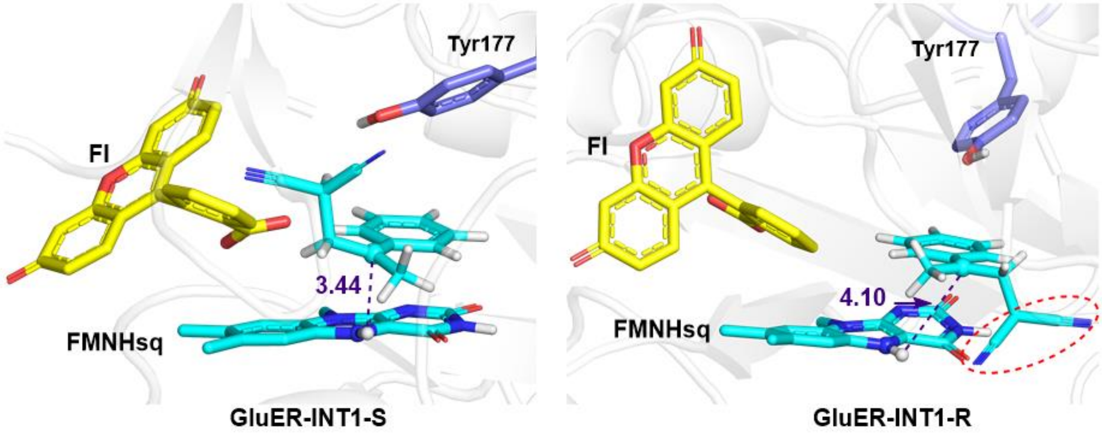

**Supplementary Fig. 28 | Structures of Int. C with *S* and *R* configurations in GluER.**

Since only the HAT process in the reaction depends on the enzyme, this step will determine the configuration of product **3a**. Regarding the HAT process, our findings suggest that the barrier correlates with the proximity of the prochiral carbon-centered radical (**Int. C**) to neutral FMNHsq. In the $CH_3$-left conformation of **Int. C** (GluER-INT1-R), the larger -CN substituent is positioned closer to FMNHsq, which increases the distance between the target C of Int. C and the N5-H of FMNHsq (3.44 Å in $CH_3$-right vs in 4.10 Å $CH_3$-left), thus increasing the energy barrier of the HAT process.

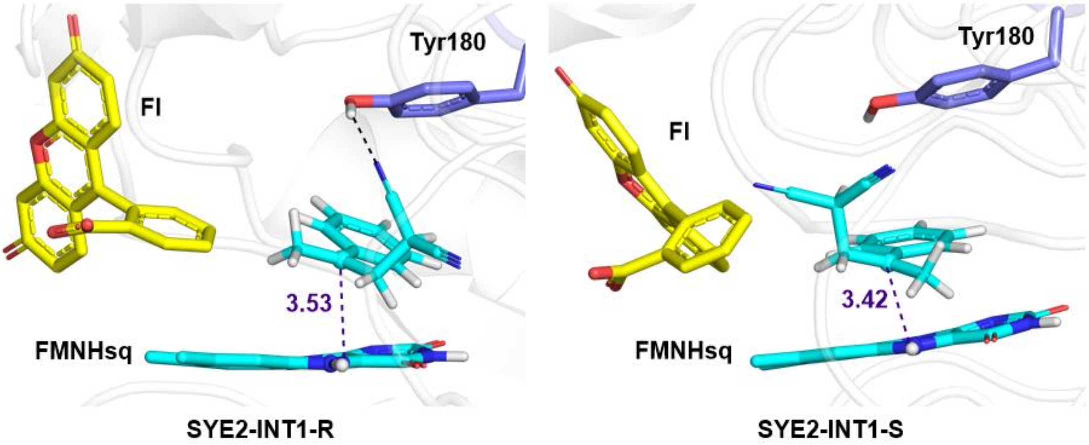

**Supplementary Fig. 29 | Structures of Int. C with *R* and *S* configurations in SYE2.** Unlike GluER, in the active site of SYE2, the larger -CN substituent in the $CH_3$-left conformation of Int. C (SYE2-INT1-R) forms a stable hydrogen bond with Tyr180, positioning -CN away from FMNHsq and thereby facilitating the subsequent HAT process and resulting in the formation of (*R*)-**3a**.

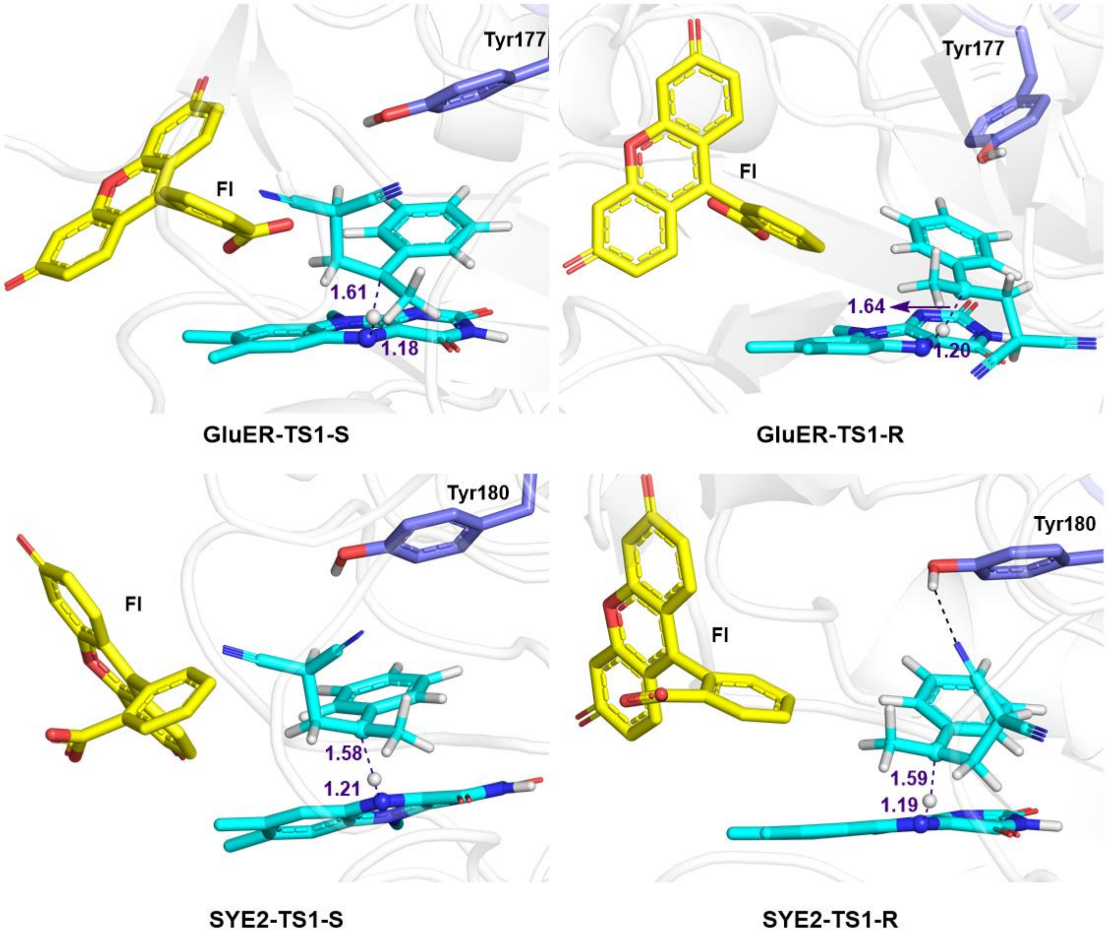

**Supplementary Fig. 30 | Structures of transition states (TS1) with *R* and *S* configurations in GluER and SYE2.** QM(B3LYP-D3/B1)/MM optimized structures of GluER-TS1-S, GluER-TS1-R, SYE2-TS1-S and SYE2-TS1-R. The distances are given in Å.

### 8. Analytical Data for the Substrates

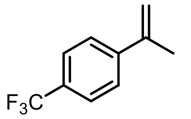

**2e**

1-(Prop-1-en-2-yl)-4-(trifluoromethyl) benzene **2e**

$^1$H NMR (600 MHz, $CDCl_3$) δ 7.56 (q, J = 8.5 Hz, 4H), 5.43 (s, 1H), 5.20 – 5.17 (m, 1H), 2.16 (s, 3H).

$^{13}$C NMR (151 MHz, $CDCl_3$) δ 144.80, 142.25, 129.72, 129.50, 129.29, 129.07, 127.01, 125.82, 125.26, 125.23, 125.21, 125.18, 123.41, 121.61, 114.58, 21.66.

$^{19}$F NMR (565 MHz, $CDCl_3$) δ -62.52.

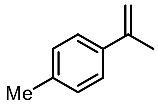

**2f**

2-p-Tolylpropene **2f**

$^1$H NMR (600 MHz, $CDCl_3$) δ 7.39 – 7.31 (m, 2H), 7.16 – 7.09 (m, 2H), 5.34 – 5.32 (m, 1H), 5.02 (d, J = 1.5 Hz, 1H), 2.33 (s, 3H), 2.13 (s, 3H).

$^{13}$C NMR (151 MHz, $CDCl_3$) δ 143.08, 138.36, 137.13, 128.91, 125.37, 111.55, 21.84, 21.07.

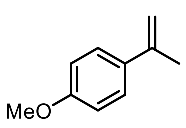

**2g**

1-Methoxy-4-(1-methylethenyl) benzene **2g**

$^{1}$H NMR (600 MHz, $CDCl_3$) δ 7.43 – 7.37 (m, 2H), 6.89 – 6.79 (m, 2H), 5.28 (d, J = 1.1 Hz, 1H), 4.98 (d, J = 1.5 Hz, 1H), 3.78 (s, 3H), 2.12 (s, 3H).

$^{13}$C NMR (151 MHz, $CDCl_3$) δ 158.05, 141.51, 132.71, 125.55, 112.51, 109.60, 54.19, 20.85.

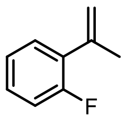

**2h**

1-Fluoro-2-(prop-1-en-2-yl) benzene **2h**

$^{1}$H NMR (600 MHz, $CDCl_3$) δ 7.30 (td, J = 7.7, 1.8 Hz, 1H), 7.24 – 7.20 (m, 1H), 7.09 (td, J = 7.5, 1.2 Hz, 1H), 7.03 (ddd, J = 11.2, 8.2, 1.2 Hz, 1H), 5.24 – 5.22 (m, 2H), 2.15 (d, J = 1.3 Hz, 3H).

$^{13}$C NMR (151 MHz, $CDCl_3$) δ 160.82, 159.18, 140.22, 130.33, 130.24, 129.41, 129.38, 128.68, 128.63, 123.90, 123.88, 116.59, 116.56, 115.90, 115.74, 23.09, 23.06.

$^{19}$F NMR (565 MHz, $CDCl_3$) δ -114.49.

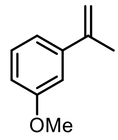

**2i**

1-Methoxy-3-(prop-1-en-2-yl) benzene **2i**

$^{1}$H NMR (600 MHz, $CDCl_3$) δ 7.24 (t, J = 8.0 Hz, 1H), 7.06 (ddd, J = 7.7, 1.7, 1.0 Hz, 1H), 7.00 (dd, J = 2.6, 1.7 Hz, 1H), 6.81 (ddd, J = 8.2, 2.6, 0.9 Hz, 1H), 5.36 (d, J = 0.8 Hz, 1H), 5.09 – 5.07 (m, 1H), 3.81 (s, 3H), 2.14 (s, 3H).

$^{13}$C NMR (151 MHz, $CDCl_3$) δ 159.56, 143.23, 142.86, 129.17, 118.13, 112.68, 112.63, 111.56, 55.21, 21.87.

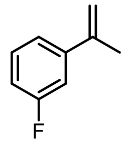

**2j**

1-Fluoro-3-(1-methylethenyl) benzene **2j**

$^{1}$H NMR (600 MHz, $CDCl_3$) δ 7.28 – 7.20 (m, 2H), 7.14 (dt, J = 10.7, 2.1 Hz, 1H), 6.97 – 6.90 (m, 1H), 5.38 (d, J = 1.3 Hz, 1H), 5.11 (d, J = 1.6 Hz, 1H), 2.12 (s, 3H).

$^{13}$C NMR (151 MHz, $CDCl_3$) δ 163.75, 162.13, 143.63, 143.58, 142.22, 142.20, 129.65, 129.60, 121.17, 121.15, 114.22, 114.08, 113.50, 112.56, 112.41, 21.67.

19F NMR (565 MHz, $CDCl_3$) δ -113.68.

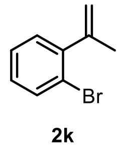

**2k**

1-bromo-2-(prop-1-en-2-yl) benzene **2k**

$^1$H NMR (600 MHz, $CDCl_3$) δ 7.54 (dd, J = 8.1, 1.3 Hz, 1H), 7.27 – 7.22 (m, 1H), 7.18 (dd, J = 7.6, 1.8 Hz, 1H), 7.09 (td, J = 7.7, 1.8 Hz, 1H), 5.22 (d, J = 1.6 Hz, 1H), 4.95 – 4.92 (m, 1H), 2.09 (s, 3H).

$^{13}$C NMR (151 MHz, $CDCl_3$) δ 145.78, 144.81, 132.73, 129.70, 128.34, 127.21, 121.53, 116.00, 23.53.

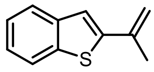

**2m**

2-(prop-1-en-2-yl) benzo[b]thiophene **2m**

$^1$H NMR (600 MHz, $CDCl_3$) δ 7.74 – 7.70 (m, 1H), 7.65 (dd, *J* = 7.1, 2.0 Hz, 1H), 7.30 – 7.23 (m, 2H), 7.13 (s, 1H), 5.47 (s, 1H), 5.09 (s, 1H), 2.18 (s, 3H).

$^{13}$C NMR (151 MHz, $CDCl_3$) δ 145.67, 140.44, 139.15, 137.66, 124.72, 124.36, 123.61, 122.14, 120.71, 114.13, 21.41.

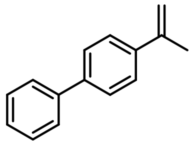

**2n**

4-(Prop-1-en-2-yl)-1,1'-biphenyl **2n**

$^1$H NMR (600 MHz, $CDCl_3$) δ 7.87 – 7.70 (m, 6H), 7.63 (t, *J* = 7.6 Hz, 2H), 7.54 (t, *J* = 7.4 Hz, 1H), 5.67 (s, 1H), 5.34 (s, 1H), 2.40 (s, 3H).

$^{13}$C NMR (151 MHz, $CDCl_3$) δ 142.93, 140.94, 140.38, 140.28, 129.01, 127.49, 127.18, 127.13, 126.14, 112.68, 22.00.

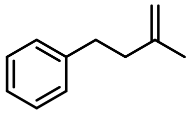

**2o**

(3-methylbut-3-en-1-yl) benzene **2o**

$^{1}$H NMR (600 MHz, $CDCl_3$) δ 7.26 (t, *J* = 7.7 Hz, 2H), 7.20 – 7.14 (m, 3H), 4.72 (d, *J* = 16.4 Hz, 2H), 2.76 – 2.72 (m, 2H), 2.33 – 2.29 (m, 2H), 1.76 (s, 3H).

$^{13}$C NMR (151 MHz, $CDCl_3$) δ 145.43, 142.31, 128.43, 128.40, 125.88, 110.34, 39.73, 34.37, 22.72.

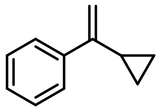

**4**

(1-cyclopropylvinyl) benzene **4**

$^{1}$H NMR (600 MHz, $CDCl_3$) δ 7.59 (dd, *J* = 7.6, 1.7 Hz, 2H), 7.33 (dd, *J* = 8.5, 6.8 Hz, 2H), 7.27 (t, *J* = 7.3 Hz, 1H), 5.27 (s, 1H), 4.93 (s, 1H), 1.64 (ddd, *J* = 13.8, 8.4, 5.3 Hz, 1H), 0.84 – 0.80 (m, 2H), 0.59 (td, *J* = 6.0, 4.1 Hz, 2H).

$^{13}$C NMR (151 MHz, $CDCl_3$) δ 149.41, 141.69, 128.20, 127.49, 126.17, 109.06, 15.69, 6.73.

### 9. ECD Spectra and Structural Refinement

To obtain the absolute configuration of the enantioenriched products, we measured the circular dichroism of the enantiomer **3a** catalyzed by SYE2 and GluER respectively with a Chirascan V100. Conformational search of investigated compounds has been carried out by using Molclus software (version 1.9.9.4) together with Gaussian 09 software package. All stable conformations obtained are then optimized using DFT at the PBE1PBE /6-31G(d) level, and the harmonic vibrational frequencies of each conformation were calculated at the same level. The self-consistent reaction field (SCRF) and SMD solvation model was adopted to evaluate the effect of solvent (Methanol). The energy of each conformer and Boltzmann distribution was calculated. Based on the Boltzmann distribution of each conformer, the ECD of each represent conformer was calculated by TDDFT at the pbe1pbe/def2tzvp level and the weighted ECD spectrums were plotted by Multiwfn software.[12]

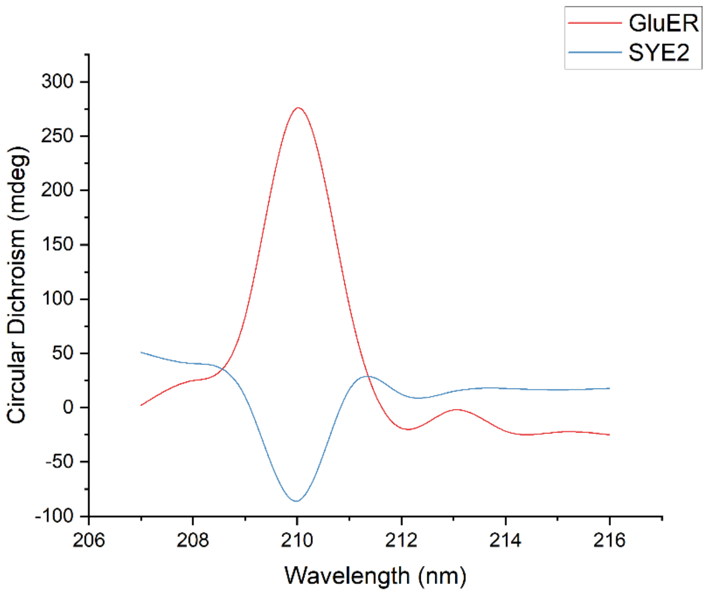

**Supplementary Fig. 31 | The experimental ECD of 3a catalyzed by SYE2 and GluER.**

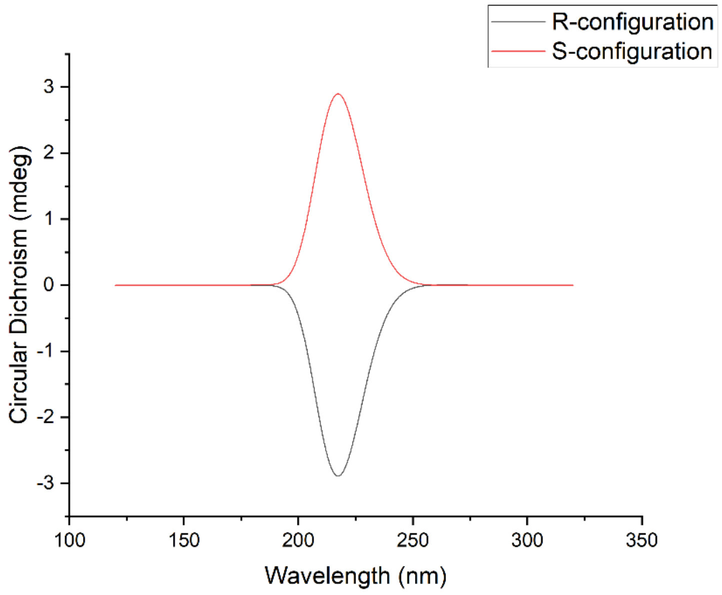

**Supplementary Fig. 32 | The predicted ECD of (*R*) and (*S*)-3a.**

**Conclusion**: The enantioenriched product obtained by SYE2 is (*R*)-enantiomer while GluER offers (*S*)-enantiomer.

### 10. Substrate Scope of Enantiodivergent Hydroalkylation Catalyzed by SYE2-WT and GluER-WT

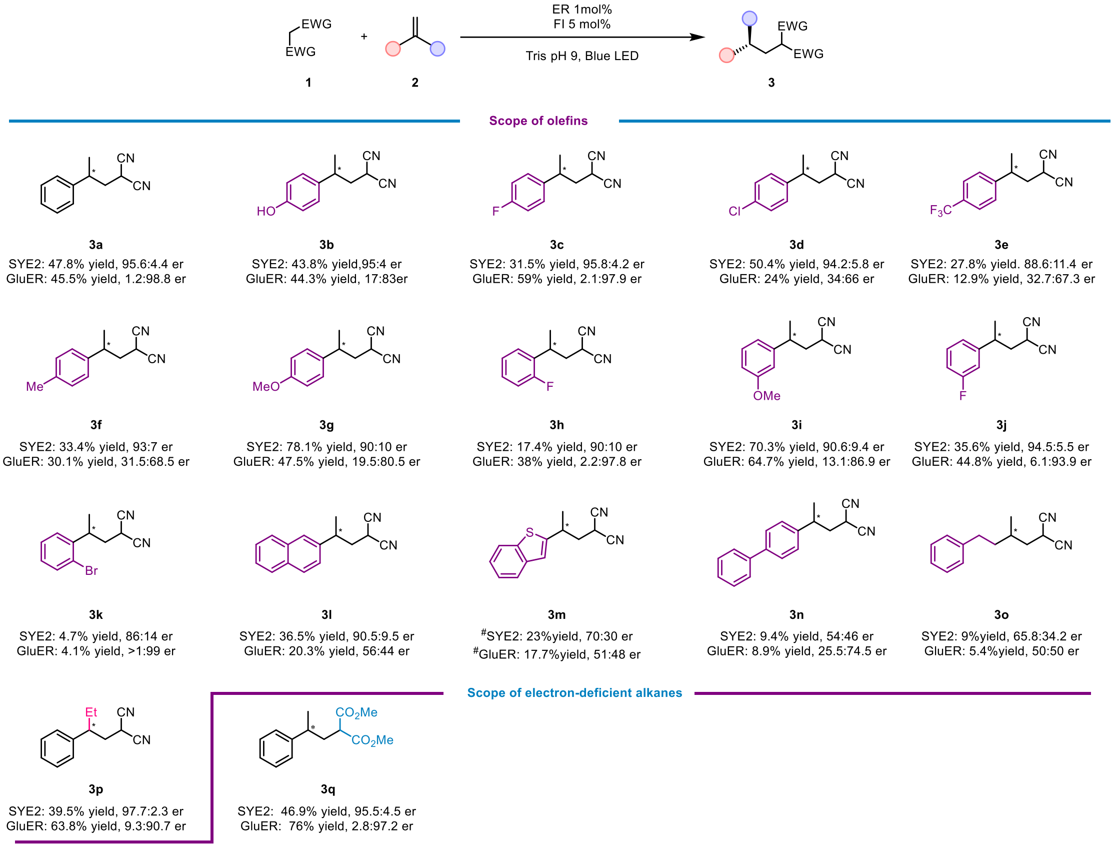

**Supplementary Fig. 33 | Substrate Scope of Radical Hydroalkylation Catalyzed by SYE2-WT and GluER-WT.** Conditions: **1** (0.008 mmol), **2** (0.004 mmol), ER (1 mol%), **FI** (5 mol%) and EtOH (7.5 v/v%) in Tris buffer (50 mM, pH 9) were stirred at room temperature for 12 h under an $N_2$ atmosphere with 455nm LEDs. The total volume of reaction was 400 µL. Yield was determined by gas chromatography and the enantiomeric ratio was determined by HPLC analysis on a chiral stationary phase. # Isolation yield was obtained by scaling up the reaction.

### 11. Experimental and Characterization Data of Products

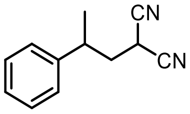

**3a**

2-(2-Phenyl-propyl) malononitrile **3a**

By using the typical procedure described in Section 5, the reaction of malononitrile **1a** (0.008 mmol) and prop-1-en-2-ylbenzene **2a** (0.004 mmol) catalyzed by SYE2 (1 mol%) with FI (5 mol%) afforded **3a** in 47.8% GC yield (average of duplicate runs), with a 95.6:4.4 er. The corresponding reaction catalyzed by GluER-Y343A (1 mol%) afforded **3a** in 56% GC yield (average of duplicate runs), with a 0.5:99.5 er (the other enantiomer as the major).

GC calibration curve for **3a** is displayed below:

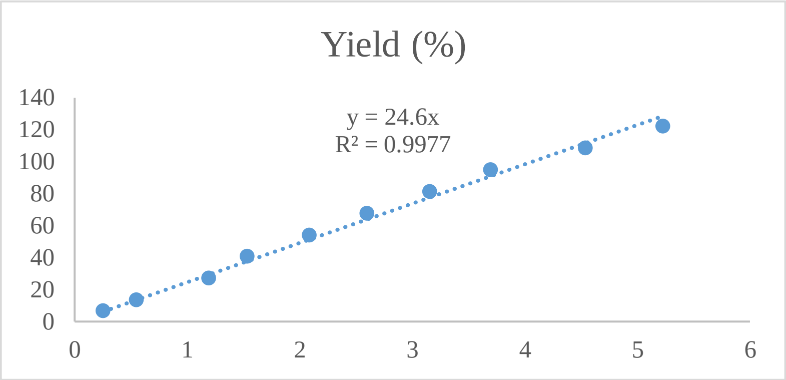

Enantiomeric ratio was determined by chiral HPLC using a Chiralpak IF column (4.6 x 250 mm, 5 μm), (HPLC: IF, 210 nm, n-hexane/isopropanol = 95:5, flow rate 1 mL/min, 25 °C, Retention Time: 7.416 min, 7.892 min).

HPLC trace of rac-**3a**:

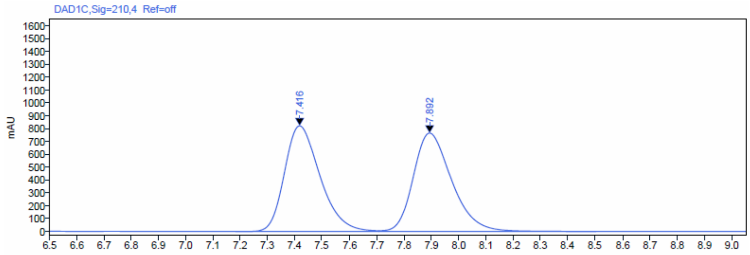

Signal: DAD1C,Sig=210,4 Ref=off

| RT [min] | Type | Width [min] | Area | Height | Area% | Name |
|---|---|---|---|---|---|---|
| 7.416 | BV | 0.53 | 7420.71 | 821.71 | 49.83 | |
| 7.892 | VBA | 0.79 | 7472.34 | 764.92 | 50.17 | |

HPLC trace of enantioenriched-**3a** obtained from the reaction catalyzed by SYE2:

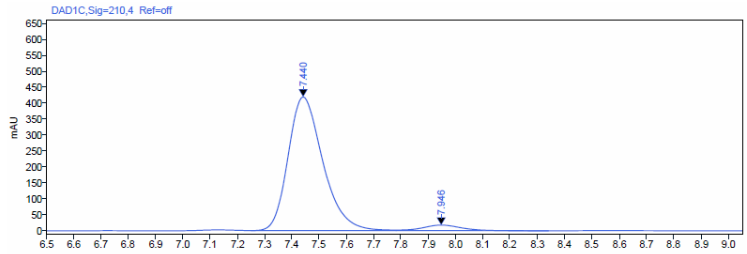

Signal: DAD1C,Sig=210,4 Ref=off

| RT [min] | Type | Width [min] | Area | Height | Area% | Name |
|---|---|---|---|---|---|---|
| 7.440 | BV | 0.53 | 3707.48 | 419.38 | 95.62 | |
| 7.946 | VB | 0.56 | 169.95 | 17.45 | 4.38 | |

HPLC trace of enantioenriched-**3a** obtained from the reaction catalyzed by GluER-Y343A:

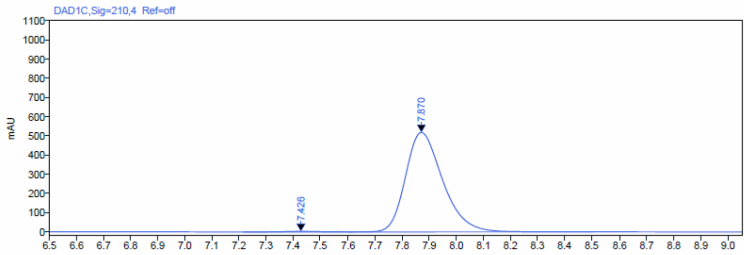

Signal: DAD1C,Sig=210,4 Ref=off

| RT [min] | Type | Width [min] | Area | Height | Area% | Name |
|---|---|---|---|---|---|---|
| 7.426 | BV | 0.40 | 24.51 | 2.89 | 0.50 | |
| 7.870 | VBA | 0.89 | 4925.28 | 518.03 | 99.50 | |

Rac-**3a** is a colorless oil.

$^1$H NMR (600 MHz, $CDCl_3$) δ 7.36 (t, J = 7.6 Hz, 2H), 7.31 – 7.26 (m, 1H), 7.22 – 7.19 (m, 2H), 3.30 (dd, J = 10.9, 5.1 Hz, 1H), 3.05 – 2.99 (m, 1H), 2.36 (ddd, J = 13.6, 10.9, 4.8 Hz, 1H), 2.19 (ddd, J = 13.6, 11.0, 5.1 Hz, 1H), 1.38 (d, J = 7.0 Hz, 3H).

$^{13}$C NMR (151 MHz, $CDCl_3$) δ 142.11, 129.36, 127.75, 126.85, 112.77, 112.31, 38.81, 37.53, 21.83, 21.03.

HRMS (APCI, m/z) calcd for $C_{12}H_{13}N_2$ $[M+H]^+$: 185.1079, found: 185.1096.

HRMS spectrum of **3a**:

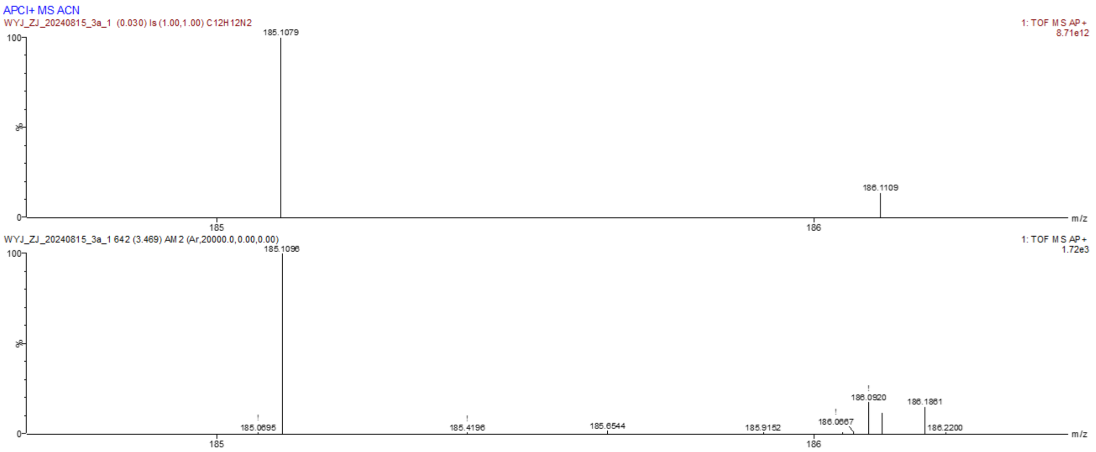

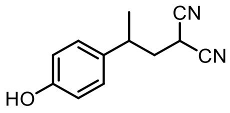

**3b**

2-(2-(4-Hydroxy-phenyl) propyl) malononitrile **3b**

By using the typical procedure described in Section 5, the reaction of malononitrile **1a** (0.008 mmol) and 4-(prop-1-en-2-yl)phenol **2b** (0.004 mmol) catalyzed by SYE2 (1 mol%) with FI (5 mol%) afforded **3b** in 43.8% reverse phase HPLC yield (average of duplicate runs), with a 95:4 er. The corresponding reaction catalyzed by GluER-Y343A (1 mol%) afforded **3b** in 44.3% reverse phase HPLC yield (average of duplicate runs), with a 17:83 er (the other enantiomer as the major).

Reverse phase HPLC calibration curve for **3b** is displayed below:

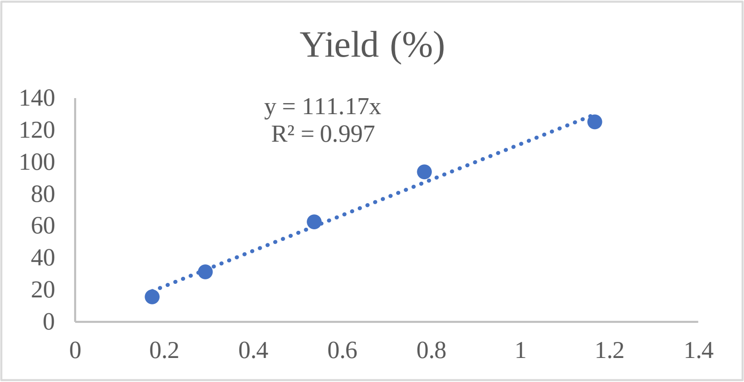

Enantiomeric ratio was determined by chiral HPLC using a Chiralpak IC column (4.6 x 250 mm, 5 μm), (HPLC: IF, 210 nm, n-hexane/isopropanol = 90:10, flow rate 1 mL/min, 25 °C, Retention Time: 28.445 min, 38.394 min).

HPLC trace of rac-**3b**:

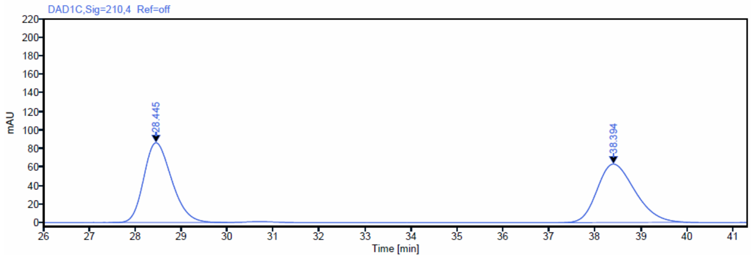

| Signal: | DAD1C,Sig=210,4 Ref=off | | | | | |
|---|---|---|---|---|---|---|
| RT [min] | Type | Width [min] | Area | Height | Area% | Name |
| 28.445 | BBA | 2.49 | 3630.60 | 86.08 | 50.19 | |
| 38.394 | BBA | 2.81 | 3602.55 | 63.08 | 49.81 | |

HPLC trace of enantioenriched-**3b** obtained from the reaction catalyzed by SYE2:

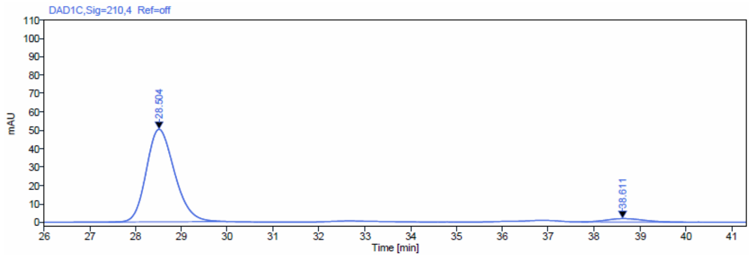

| Signal: | DAD1C,Sig=210,4 Ref=off | | | | | |
|---|---|---|---|---|---|---|
| RT [min] | Type | Width [min] | Area | Height | Area% | Name |
| 28.504 | BBA | 2.60 | 2153.34 | 50.28 | 95.33 | |
| 38.611 | BBA | 2.28 | 105.44 | 1.88 | 4.67 | |

HPLC trace of enantioenriched-**3b** obtained from the reaction catalyzed by GluER-Y343A:

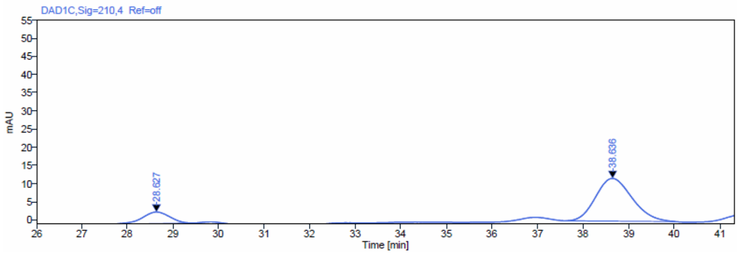

| Signal: | DAD1C,Sig=210,4 Ref=off | | | | | |
|---|---|---|---|---|---|---|
| RT [min] | Type | Width [min] | Area | Height | Area% | Name |
| 28.627 | BB | 1.93 | 134.46 | 3.19 | 17.08 | |
| 38.636 | BBA | 2.39 | 652.78 | 11.74 | 82.92 | |

Rac-**3b** is a yellow oil.

$^1$H NMR (600 MHz, $CDCl_3$) δ 7.01 – 6.95 (m, 2H), 6.76 – 6.70 (m, 2H), 6.02 (s, 1H), 3.30 (dd, J = 10.7, 5.1 Hz, 1H), 2.86 (tt, J = 11.6, 6.8 Hz, 1H), 2.25 (ddd, J = 13.7, 10.7, 4.7 Hz, 1H), 2.04 (ddd, J = 13.7, 11.1, 5.1 Hz, 1H), 1.25 (d, J = 7.1 Hz, 3H).

$^{13}$C NMR (151 MHz, $CDCl_3$) δ 154.18, 132.93, 127.10, 115.16, 111.91, 111.44, 37.75, 35.79, 21.00, 20.05.

HRMS (ESI, m/z) calcd for $C_{12}H_{13}N_2O$ $[M+H]^+$: 201.1028, found: 201.1044.

HRMS spectrum of **3b**:

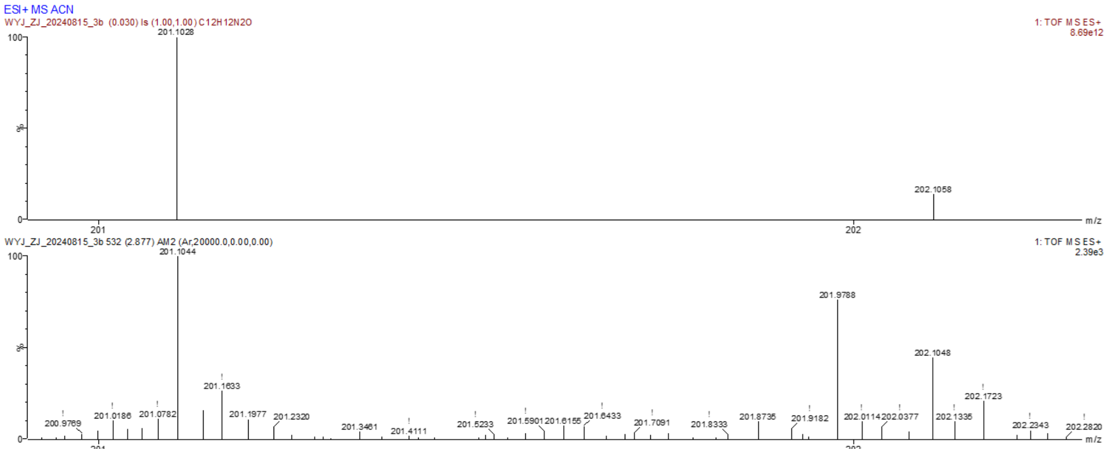

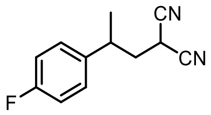

**3c**

2-(2-(4-Fluorophenyl) propyl) malononitrile **3c**

By using the typical procedure described in Section 5, the reaction of malononitrile **1a** (0.008 mmol) and 1-fluoro-4-(prop-1-en-2-yl)benzene **2c** (0.004 mmol) catalyzed by SYE2 (1 mol%) with FI (5 mol%) afforded **3c** in 31.5% reverse phase HPLC yield (average of duplicate runs), with a 95.4:4.6 er. The corresponding reaction catalyzed by GluER-Y343A (1 mol%) afforded **3c** in 59% reverse phase HPLC yield (average of duplicate runs), with a > 1:99 er (the other enantiomer as the major).

Reverse phase HPLC calibration curve for **3c** is displayed below:

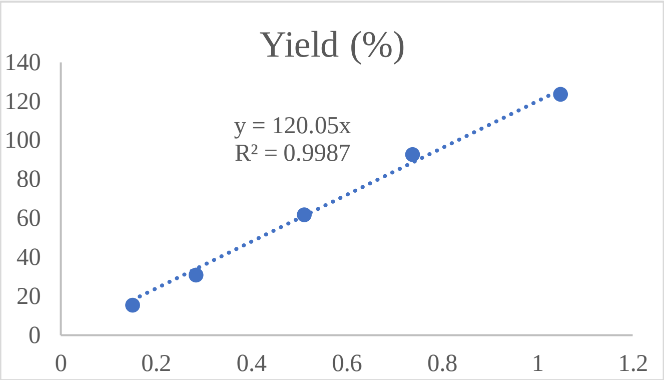

Enantiomeric ratio was determined by chiral HPLC using a Chiralpak IC column (4.6 x 250 mm, 5 μm), (HPLC: IF, 210 nm, n-hexane/isopropanol = 90:10, flow rate 1 mL/min, 25 °C, Retention Time: 10.063 min, 11.485 min).

HPLC trace of rac-**3c**:

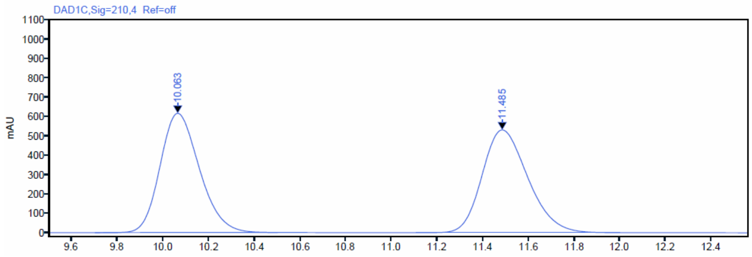

Signal: DAD1C,Sig=210,4 Ref=off

| RT [min] | Type | Width [min] | Area | Height | Area% | Name |
|---|---|---|---|---|---|---|
| 10.063 | BBA | 0.80 | 7363.45 | 615.17 | 49.97 | |
| 11.485 | BBA | 0.89 | 7373.34 | 530.25 | 50.03 | |

HPLC trace of enantioenriched-**3c** obtained from the reaction catalyzed by SYE2:

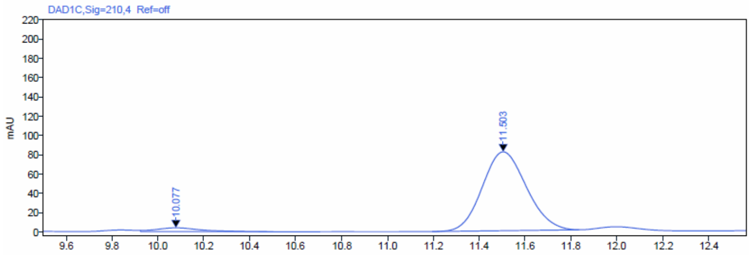

Signal: DAD1C,Sig=210,4 Ref=off

| RT [min] | Type | Width [min] | Area | Height | Area% | Name |
|---|---|---|---|---|---|---|
| 10.077 | VBA | 0.58 | 53.09 | 3.93 | 4.64 | |
| 11.503 | BB | 0.64 | 1090.52 | 81.69 | 95.36 | |

HPLC trace of enantioenriched-**3c** obtained from the reaction catalyzed by GluER-Y343A:

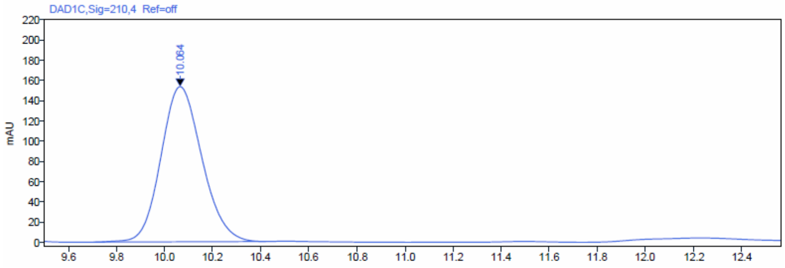

Signal: DAD1C,Sig=210,4 Ref=off

| RT [min] | Type | Width [min] | Area | Height | Area% | Name |
|---|---|---|---|---|---|---|
| 10.064 | BB | 0.71 | 1813.31 | 153.19 | 100.00 | |

Rac-**3c** is a colorless oil.

$^{1}$H NMR (600 MHz, $CDCl_3$) δ 7.21 – 7.17 (m, 2H), 7.08 – 7.04 (m, 2H), 3.33 (dd, J = 10.6, 5.2 Hz, 1H), 3.07 – 3.01 (m, 1H), 2.36 (ddd, J = 13.7, 10.6, 4.9 Hz, 1H), 2.17 (ddd, J = 13.7, 10.9, 5.2 Hz, 1H), 1.37 (d, J = 7.0 Hz, 3H).

$^{13}$C NMR (151 MHz, $CDCl_3$) δ 162.96, 161.33, 137.89, 128.42, 116.34, 116.20, 112.64, 112.21, 38.81, 36.91, 21.97, 21.04.

$^{19}$F NMR (565 MHz, CDCl3) δ -114.44.

HRMS (APCI, m/z) calcd for $C_{12}H_{12}FN_2$ $[M+H]^+$: 203.0984, found: 203.1031.

HRMS spectrum of **3c**:

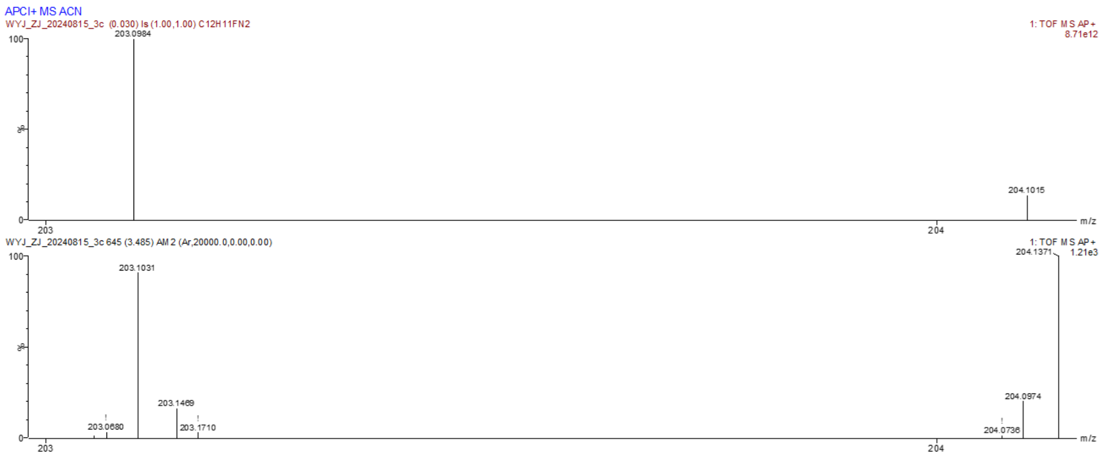

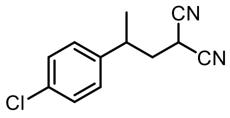

**3d**

2-(2-(4-Chlorophenyl) propyl) malononitrile **3d**

By using the typical procedure described in Section 5, the reaction of malononitrile **1a** (0.008 mmol) and 1-chloro-4-(prop-1-en-2-yl)benzene **2d** (0.004 mmol) catalyzed by SYE2 (1 mol%) with FI (5 mol%) afforded **3d** in 50.4% reverse phase HPLC yield (average of duplicate runs), with a 92.7:7.3 er. The corresponding reaction catalyzed by GluER-Y343A (1 mol%) afforded **3d** in 83% reverse phase HPLC yield (average of duplicate runs), with a 2.3:97.7 er (the other enantiomer as the major).

Reverse phase HPLC calibration curve for **3d** is displayed below:

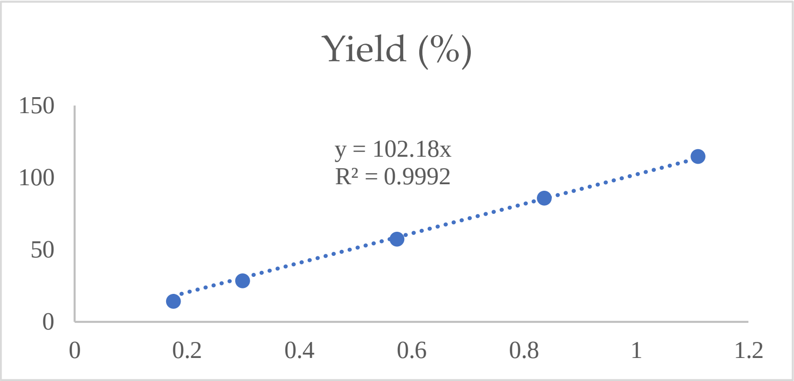

Enantiomeric ratio was determined by chiral HPLC using a Chiralpak IC column (4.6 x 250 mm, 5 μm), (HPLC: IF, 210 nm, n-hexane/isopropanol = 90:10, flow rate 1 mL/min, 25 °C, Retention Time: 10.831 min, 12.314 min).

HPLC trace of rac-**3d**:

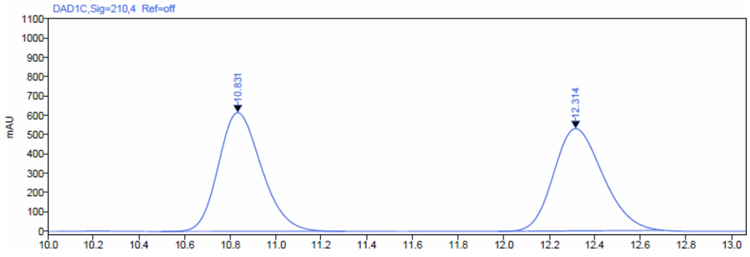

| Signal: | DAD1C,Sig=210,4 Ref=off | | | | | |
|---|---|---|---|---|---|---|
| RT [min] | Type | Width [min] | Area | Height | Area% | Name |
| 10.831 | BBA | 0.81 | 8078.72 | 614.50 | 50.40 | |
| 12.314 | BBA | 0.73 | 7951.83 | 529.37 | 49.60 | |

HPLC trace of enantioenriched-**3d** obtained from the reaction catalyzed by SYE2:

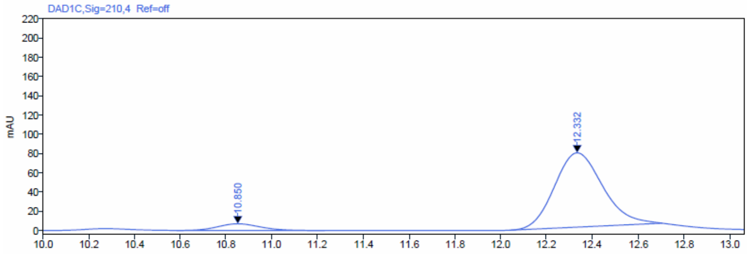

| Signal: | DAD1C,Sig=210,4 Ref=off | | | | | |
|---|---|---|---|---|---|---|
| RT [min] | Type | Width [min] | Area | Height | Area% | Name |
| 10.850 | BBA | 0.65 | 88.48 | 6.92 | 7.26 | |
| 12.332 | BBA | 0.67 | 1129.65 | 77.11 | 92.74 | |

HPLC trace of enantioenriched-**3d** obtained from the reaction catalyzed by GluER-Y343A:

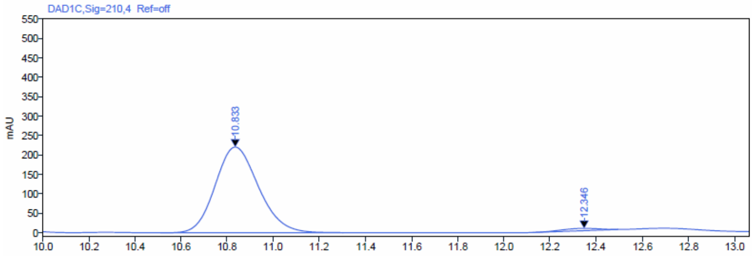

| Signal: | DAD1C,Sig=210,4 Ref=off | | | | | |
|---|---|---|---|---|---|---|
| RT [min] | Type | Width [min] | Area | Height | Area% | Name |
| 10.833 | BBA | 0.79 | 2871.77 | 220.25 | 97.72 | |
| 12.346 | BB | 0.37 | 66.86 | 5.83 | 2.28 | |

Rac-**3d** is a colorless oil.

$^{1}H$ NMR (600 MHz, $CDCl_3$) δ 7.37 – 7.30 (m, 2H), 7.17 – 7.13 (m, 2H), 3.33 (dd, J = 10.6, 5.3 Hz, 1H), 3.03 (tt, J = 11.7, 6.9 Hz, 1H), 2.36 (ddd, J = 13.7, 10.6, 4.9 Hz, 1H), 2.18 (ddd, J = 13.7, 10.8, 5.3 Hz, 1H), 1.37 (d, J = 7.0 Hz, 3H).

$^{13}C$ NMR (151 MHz, $CDCl_3$) δ 140.64, 133.55, 129.56, 128.28, 112.56, 112.14, 38.63, 37.04, 21.81, 21.03.

HRMS (APCI, m/z) calcd for $C_{12}H_{12}ClN_2$ $[M+H]^+$: 219.0689, found: 219.0713.

HRMS spectrum of **3d**:

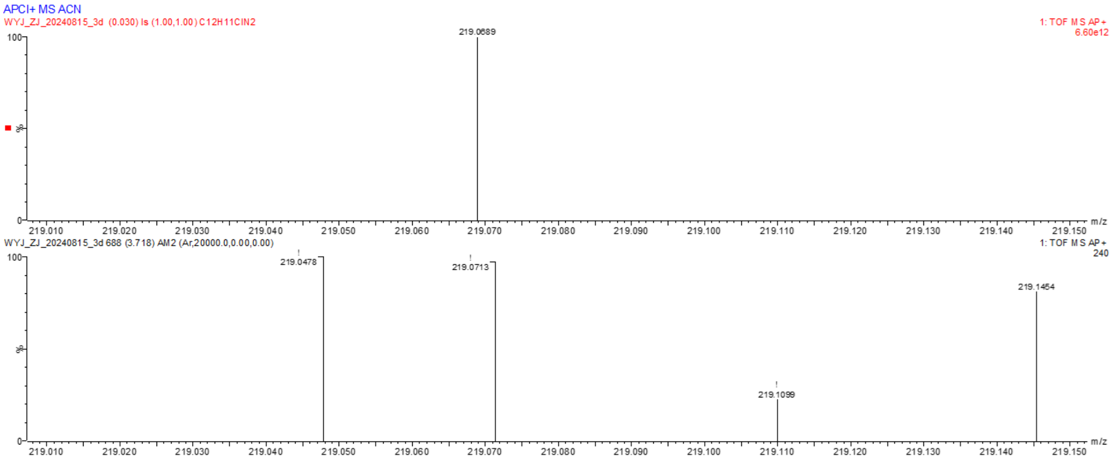

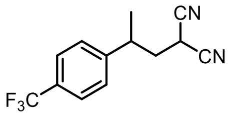

**3e**

2-(2-(4-(Trfluoromethyl) phenyl) propyl) malononitrile **3e**

By using the typical procedure described in Section 5, the reaction of malononitrile **1a** (0.008 mmol) and 1-(prop-1-en-2-yl)-4-(trifluoromethyl)benzene **2e** (0.004 mmol) catalyzed by SYE2 (1 mol%) with FI (5 mol%) afforded **3d** in 27.8% reverse phase HPLC yield (average of duplicate runs), with a 90.3:9.7 er. The corresponding reaction catalyzed by GluER-Y343A (1 mol%) afforded **3d** in 29.7% reverse phase HPLC yield (average of duplicate runs), with a 17.2:82.8 er (the other enantiomer as the major).

Reverse phase HPLC calibration curve for **3e** is displayed below:

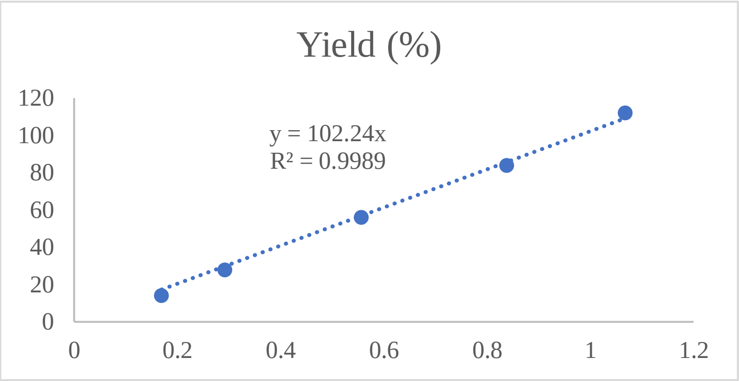

Enantiomeric ratio was determined by chiral HPLC using a Chiralpak IC column (4.6 x 250 mm, 5 μm), (HPLC: IF, 210 nm, n-hexane/isopropanol = 90:10, flow rate 1 mL/min, 25 °C, Retention Time: 7.929 min, 8.796 min).

HPLC trace of rac-**3e**:

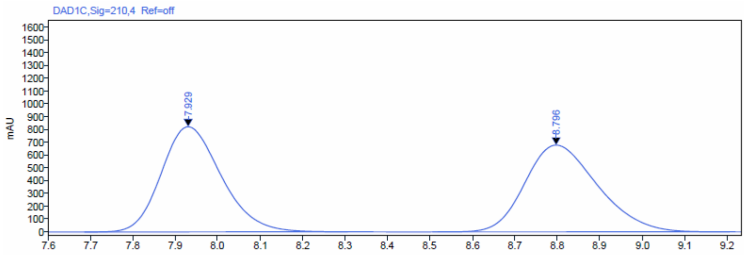

Signal: DAD1C,Sig=210,4 Ref=off

| RT [min] | Type | Width [min] | Area | Height | Area% | Name |
|---|---|---|---|---|---|---|
| 7.929 | BBA | 0.62 | 8071.80 | 821.56 | 50.03 | |
| 8.796 | BBA | 0.70 | 8061.48 | 677.69 | 49.97 | |

HPLC trace of enantioenriched-**3e** obtained from the reaction catalyzed by SYE2:

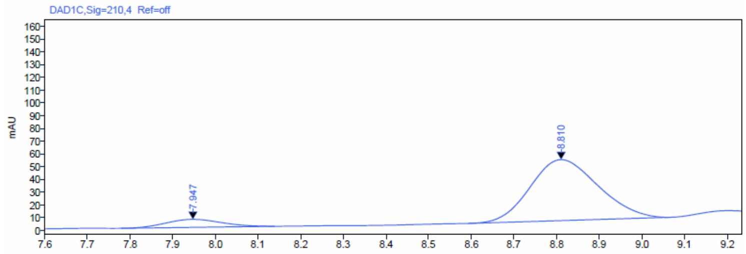

Signal: DAD1C,Sig=210,4 Ref=off

| RT [min] | Type | Width [min] | Area | Height | Area% | Name |
|---|---|---|---|---|---|---|
| 7.947 | BB | 0.36 | 56.70 | 6.27 | 9.71 | |
| 8.810 | BB | 0.47 | 527.05 | 47.74 | 90.29 | |

HPLC trace of enantioenriched-**3e** obtained from the reaction catalyzed by GluER-Y343A:

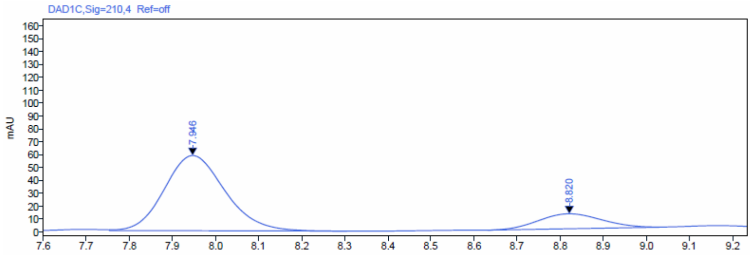

Signal: DAD1C,Sig=210,4 Ref=off

| RT [min] | Type | Width [min] | Area | Height | Area% | Name |
|---|---|---|---|---|---|---|
| 7.946 | VBA | 0.55 | 568.48 | 58.29 | 82.77 | |
| 8.820 | BV | 0.40 | 118.33 | 11.60 | 17.23 | |

Rac-**3e** is a pale yellow oil.

$^{1}$H NMR (600 MHz, $CDCl_3$) δ 7.64 (d, J = 8.1 Hz, 2H), 7.36 (d, J = 8.0 Hz, 2H), 3.33 (dd, J = 10.5, 5.5 Hz, 1H), 3.13 (dqd, J = 13.9, 6.9, 5.1 Hz, 1H), 2.40 (ddd, J = 13.8, 10.5, 5.2 Hz, 1H), 2.25 (ddd, J = 13.8, 10.6, 5.5 Hz, 1H), 1.41 (d, J = 7.0 Hz, 3H).

$^{13}$C NMR (151 MHz, $CDCl_3$) δ 146.33, 130.49, 130.27, 130.06, 129.84, 127.36, 126.62, 126.41, 126.39, 126.36, 126.34, 124.82, 123.01, 121.21, 112.39, 112.02, 38.42, 37.45, 21.64, 21.05.

$^{19}$F NMR (565 MHz, $CDCl_3$) δ -62.61.

HRMS (ESI, m/z) calcd for $C_{13}H_{12}F_3N_2$ $[M+H]^+$: 253.0947, no signal was found under both ESI and APCI detector because **3e** was not ionizable in ESI and APCI.

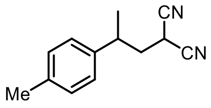

3f

2-(2-(p-Tolyl) propyl) malononitrile **3f**

By using the typical procedure described in Section 5, the reaction of malononitrile **1a** (0.008 mmol) and 1-methyl-4-(prop-1-en-2-yl)benzene **2f** (0.004 mmol) catalyzed by SYE2 (1 mol%) with FI (5 mol%) afforded **3f** in 33.4% reverse phase HPLC yield (average of duplicate runs), with a 90.3:9.7 er. The corresponding reaction catalyzed by GluER-Y343A (1 mol%) afforded **3f** in 49.3% reverse phase HPLC yield (average of duplicate runs), with a 2:98 er (the other enantiomer as the major).

Reverse phase HPLC calibration curve for **3f** is displayed below:

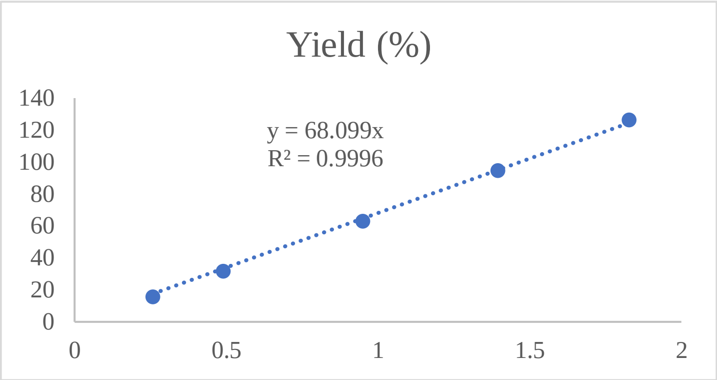

Enantiomeric ratio was determined by chiral HPLC using a Chiralpak IC column (4.6 x 250 mm, 5 μm), (HPLC: IF, 210 nm, n-hexane/isopropanol = 90:10, flow rate 1 mL/min, 25 °C, Retention Time: 7.999 min, 8.889 min).

HPLC trace of rac-**3f**:

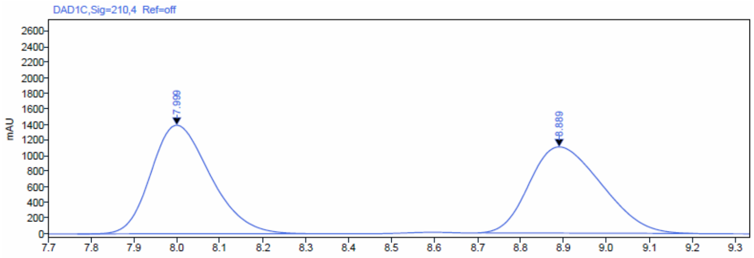

Signal: DAD1C,Sig=210,4 Ref=off

| RT [min] | Type | Width [min] | Area | Height | Area% | Name |
|---|---|---|---|---|---|---|
| 7.999 | BBA | 0.53 | 13421.75 | 1395.60 | 50.92 | |
| 8.889 | BBA | 0.50 | 12936.99 | 1108.36 | 49.08 | |

HPLC trace of enantioenriched-**3f** obtained from the reaction catalyzed by SYE2:

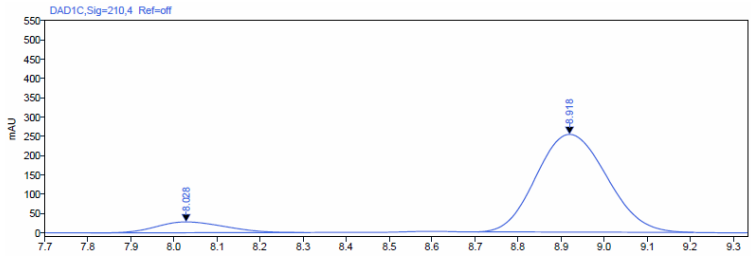

Signal: DAD1C,Sig=210,4 Ref=off

| RT [min] | Type | Width [min] | Area | Height | Area% | Name |
|---|---|---|---|---|---|---|
| 8.028 | BBA | 0.50 | 309.58 | 28.10 | 9.71 | |
| 8.918 | BBA | 0.50 | 2878.60 | 253.06 | 90.29 | |

HPLC trace of enantioenriched-**3f** obtained from the reaction catalyzed by GluER-Y343A:

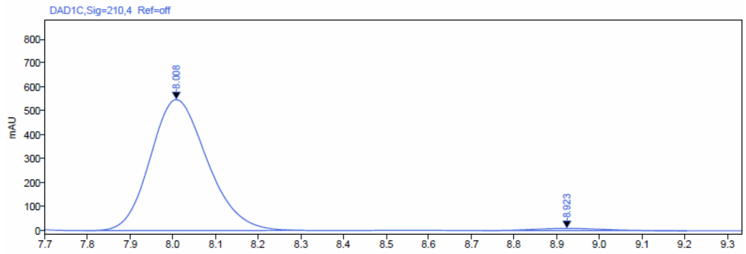

Signal: DAD1C,Sig=210,4 Ref=off

| RT [min] | Type | Width [min] | Area | Height | Area% | Name |
|---|---|---|---|---|---|---|
| 8.008 | BBA | 0.51 | 5079.70 | 547.96 | 97.97 | |
| 8.923 | BBA | 0.48 | 105.42 | 9.45 | 2.03 | |

Rac-**3f** is a colorless oil.

$^1$H NMR (600 MHz, $CDCl_3$) δ 7.17 (d, J = 7.8 Hz, 2H), 7.11 – 7.08 (m, 2H), 3.30 (dd, J = 11.1, 4.9 Hz, 1H), 2.99 (ddq, J = 13.8, 6.9, 3.5 Hz, 1H), 2.39 – 2.35 (m, 1H), 2.34 (s, 3H), 2.16 (ddd, J = 13.6, 11.2, 4.9 Hz, 1H), 1.36 (d, J = 7.0 Hz, 3H).

$^{13}$C NMR (151 MHz, $CDCl_3$) δ 138.99, 137.51, 130.05, 126.73, 112.85, 112.36, 39.00, 37.17, 21.97, 21.05, 21.04.

HRMS (APCI, m/z) calcd for $C_{13}H_{15}N_2$ $[M+H]^+$: 199.1238, found: 199.1235.

HRMS spectrum of **3f**:

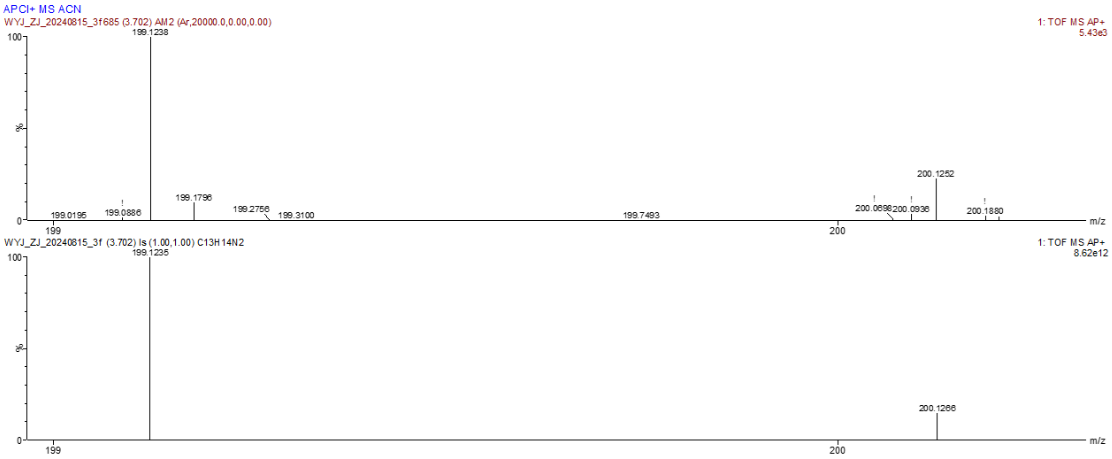

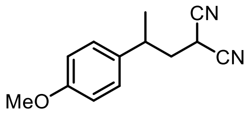

**3g**

2-(2-(4-Methoxyphenyl) propyl) malononitrile **3g**

By using the typical procedure described in Section 5, the reaction of malononitrile **1a** (0.008 mmol) and 1-methoxy-4-(prop-1-en-2-yl)benzene **2g** (0.004 mmol) catalyzed by SYE2 (1 mol%) with FI (5 mol%) afforded **3g** in 78.1% reverse phase HPLC yield (average of duplicate runs), with a 90.5:9.5 er. The corresponding reaction catalyzed by GluER-Y343A (1 mol%) afforded **3g** in 86.1% reverse phase HPLC yield (average of duplicate runs), with a 2.1:97.9 er (the other enantiomer as the major).

Reverse phase HPLC calibration curve for **3g** is displayed below:

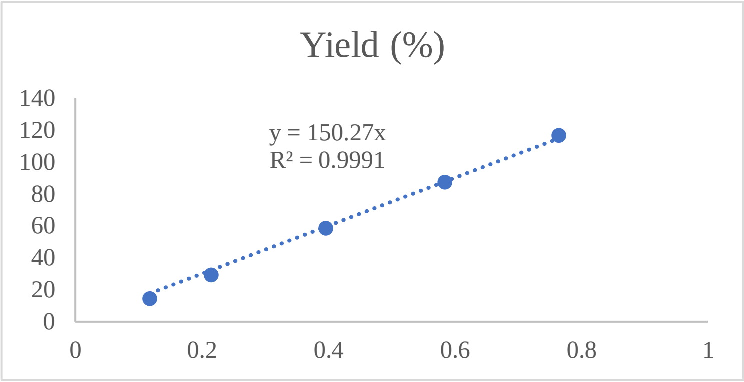

Enantiomeric ratio was determined by chiral HPLC using a Chiralpak IC column (4.6 x 250 mm, 5 μm), (HPLC: IF, 210 nm, n-hexane/isopropanol = 90:10, flow rate 1 mL/min, 25 °C, Retention Time: 13.685 min, 15.732 min).

HPLC trace of rac-**3g**:

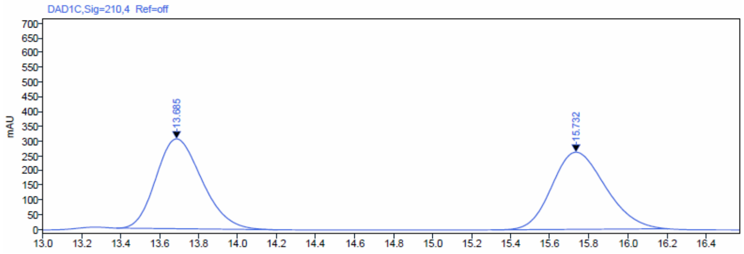

Signal: DAD1C,Sig=210,4 Ref=off

| RT [min] | Type | Width [min] | Area | Height | Area% | Name |
|---|---|---|---|---|---|---|
| 13.685 | BBA | 0.83 | 4994.73 | 304.69 | 49.92 | |
| 15.732 | BBA | 0.90 | 5010.48 | 261.67 | 50.08 | |

HPLC trace of enantioenriched-**3g** obtained from the reaction catalyzed by SYE2

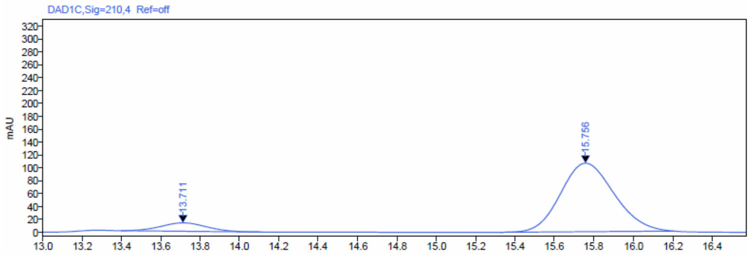

Signal: DAD1C,Sig=210,4 Ref=off

| RT [min] | Type | Width [min] | Area | Height | Area% | Name |
|---|---|---|---|---|---|---|
| 13.711 | BBA | 0.72 | 211.83 | 13.10 | 9.52 | |
| 15.756 | BBA | 0.88 | 2012.19 | 106.27 | 90.48 | |

HPLC trace of enantioenriched-**3g** obtained from the reaction catalyzed by GluER-Y343A:

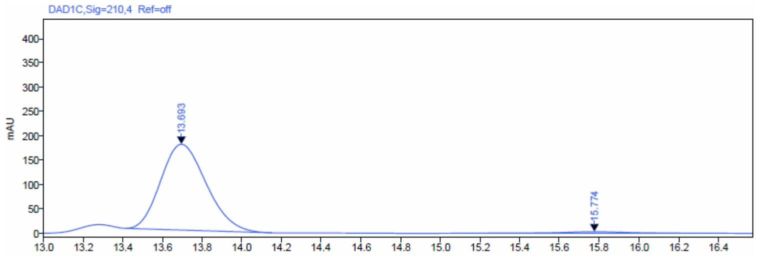

Signal: DAD1C,Sig=210,4 Ref=off

| RT [min] | Type | Width [min] | Area | Height | Area% | Name |
|---|---|---|---|---|---|---|
| 13.693 | BBA | 0.74 | 2817.10 | 176.35 | 97.87 | |
| 15.774 | BBA | 0.97 | 61.38 | 3.09 | 2.13 | |

Rac-**3g** is a colorless oil.

$^{1}$H NMR (600 MHz, $CDCl_3$) δ 7.14 – 7.11 (m, 2H), 6.91 – 6.88 (m, 2H), 3.81 (s, 3H), 3.30 (dd, J = 11.0, 4.9 Hz, 1H), 3.01 – 2.96 (m, 1H), 2.36 (ddd, J = 13.6, 11.0, 4.6 Hz, 1H), 2.14 (ddd, J = 13.6, 11.2, 4.9 Hz, 1H), 1.36 (d, J = 7.0 Hz, 3H).

$^{13}$C NMR (151 MHz, $CDCl_3$) δ 159.08, 133.91, 127.86, 114.74, 112.84, 112.35, 55.35, 39.12, 36.78, 22.05, 21.02.

HRMS (ESI, m/z) calcd for $C_{13}H_{15}N_2O$ $[M+H]^+$: 215.1184, found: 215.1192.

HRMS spectrum of **3g**:

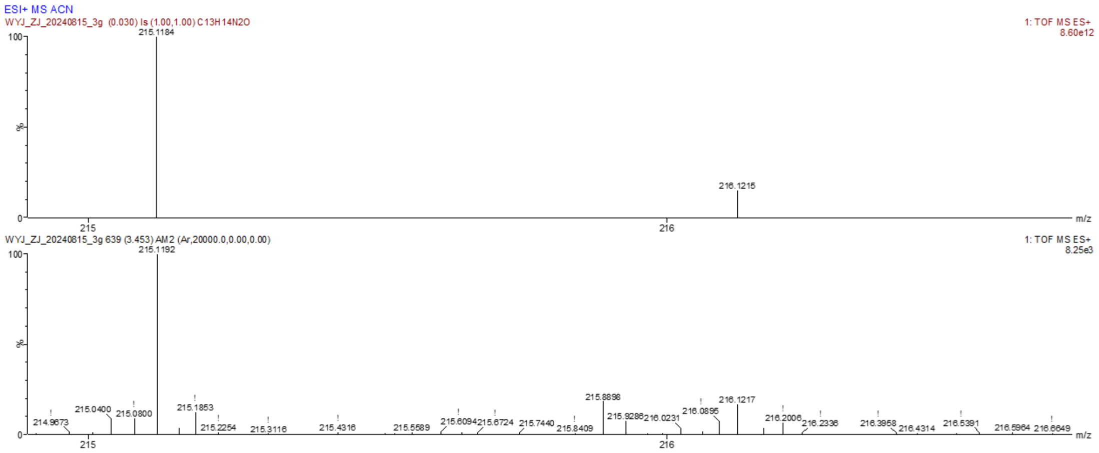

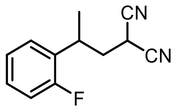

**3h**

2-(2-(2-Fluorophenyl) propyl) malononitrile **3h**

By using the typical procedure described in Section 5, the reaction of malononitrile **1a** (0.008 mmol) and 1-fluoro-2-(prop-1-en-2-yl)benzene **2h** (0.004 mmol) catalyzed by SYE2 (1 mol%) with FI (5 mol%) afforded **3h** in 17.4% reverse phase HPLC yield (average of duplicate runs), with a 87.7:12.3 er. The corresponding reaction catalyzed by GluER-Y343A (1 mol%) afforded **3h** in 38% reverse phase HPLC yield (average of duplicate runs), with a > 1:99 er (the other enantiomer as the major).

Reverse phase HPLC calibration curve for **3h** is displayed below:

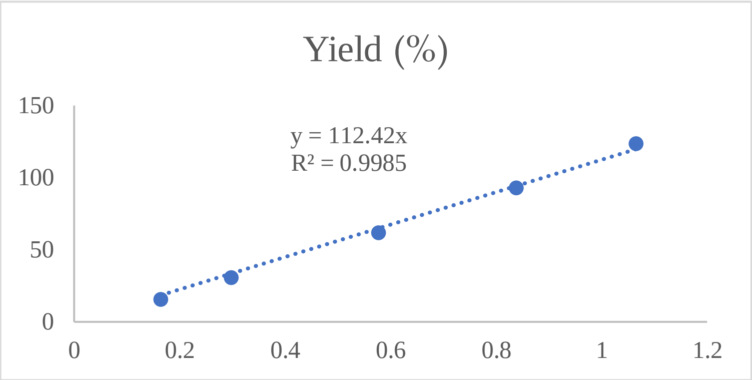

Enantiomeric ratio was determined by chiral HPLC using a Chiralpak IC column (4.6 x 250 mm, 5 μm), (HPLC: IF, 210 nm, n-hexane/isopropanol = 90:10, flow rate 1 mL/min, 25 °C, Retention Time: 8.507 min, 9.755 min).

HPLC trace of rac-**3h**:

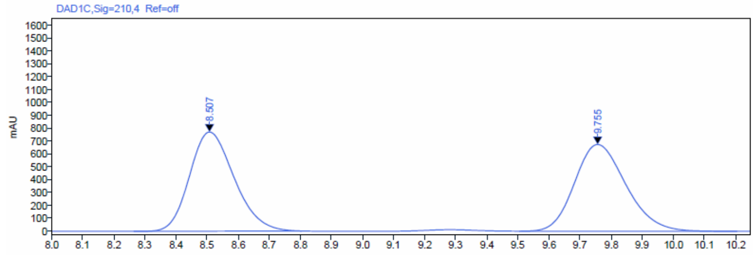

Signal: DAD1C,Sig=210,4 Ref=off

| RT [min] | Type | Width [min] | Area | Height | Area% | Name |
|---|---|---|---|---|---|---|
| 8.507 | BBA | 0.54 | 7674.58 | 770.75 | 49.83 | |
| 9.755 | BBA | 0.70 | 7726.95 | 674.05 | 50.17 | |

HPLC trace of enantioenriched-**3h** obtained from the reaction catalyzed by SYE2:

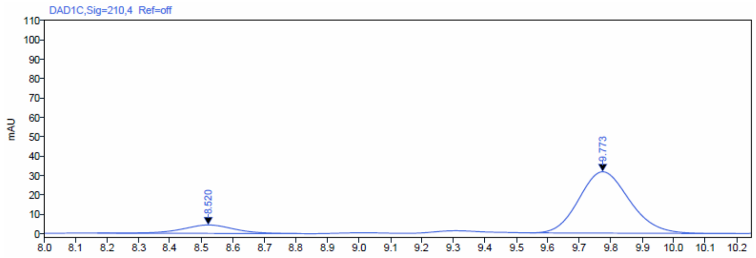

Signal: DAD1C,Sig=210,4 Ref=off

| RT [min] | Type | Width [min] | Area | Height | Area% | Name |
|---|---|---|---|---|---|---|
| 8.520 | BBA | 0.63 | 49.34 | 4.36 | 12.31 | |
| 9.773 | BBA | 0.65 | 351.51 | 31.61 | 87.69 | |

HPLC trace of enantioenriched-**3h** obtained from the reaction catalyzed by GluER-Y343A:

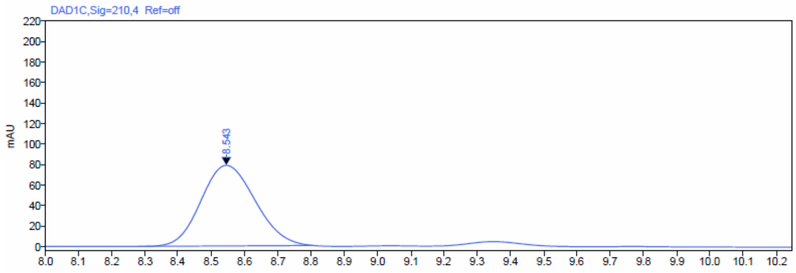

Signal: DAD1C,Sig=210,4 Ref=off

| RT [min] | Type | Width [min] | Area | Height | Area% | Name |
|---|---|---|---|---|---|---|
| 8.543 | BBA | 0.53 | 889.09 | 78.49 | 100.00 | |

Rac-**3h** is a colorless oil.

[1]H NMR (600 MHz, $CDCl_3$) δ 7.28 (dddd, J = 8.2, 7.2, 5.3, 1.8 Hz, 1H), 7.22 (td, J = 7.5, 1.8 Hz, 1H), 7.15 (td, J = 7.5, 1.3 Hz, 1H), 7.08 (ddd, J = 10.9, 8.2, 1.2 Hz, 1H), 3.43 (dd, J = 9.9, 5.8 Hz, 1H), 3.35 – 3.28 (m, 1H), 2.43 – 2.34 (m, 2H), 1.42 (d, J = 7.1 Hz, 3H).

[13]C NMR (151 MHz, $CDCl_3$) δ 161.77, 160.14, 129.37, 128.83, 125.01, 116.44, 116.29, 112.59, 112.23, 37.26, 32.48, 21.15, 20.34.

[19]F NMR (565 MHz, $CDCl_3$) δ -117.07.

HRMS (APCI, m/z) calcd for $C_{12}H_{12}FN_2$ $[M+H]^+$: 203.0984, found: 203.1015.

HRMS spectrum of **3h**:

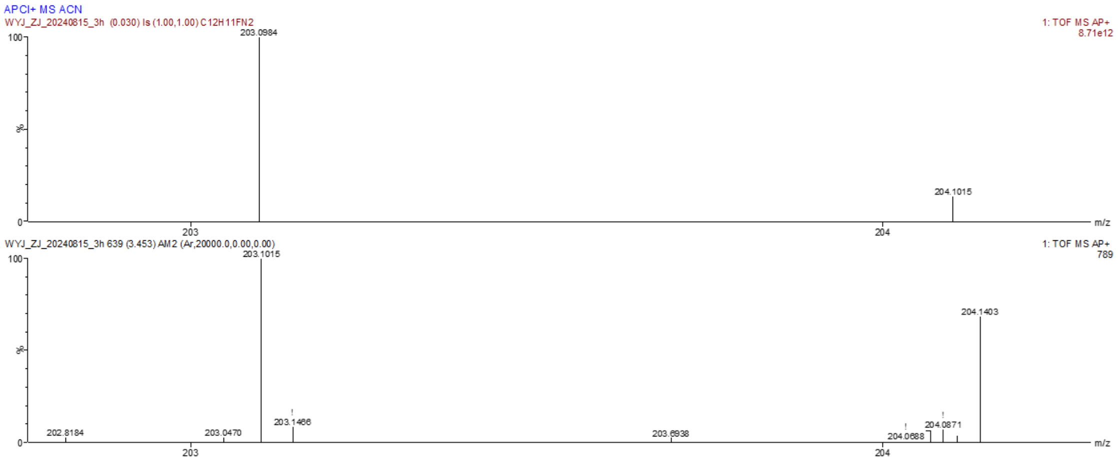

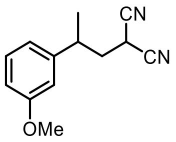

**3i**

2-(2-(3-Methoxyphenyl) propyl) malononitrile **3i**

By using the typical procedure described in Section 5, the reaction of malononitrile **1a** (0.008 mmol) and 1-methoxy-3-(prop-1-en-2-yl)benzene **2i** (0.004 mmol) catalyzed by SYE2 (1 mol%) with FI (5 mol%) afforded **3i** in 70.3% reverse phase HPLC yield (average of duplicate runs), with a 90.2:9.8 er. The corresponding reaction catalyzed by GluER-Y343A (1 mol%) afforded **3i** in 74.7% reverse phase HPLC yield (average of duplicate runs), with a 12.2:87.8 er (the other enantiomer as the major).

Reverse phase HPLC calibration curve for **3i** is displayed below:

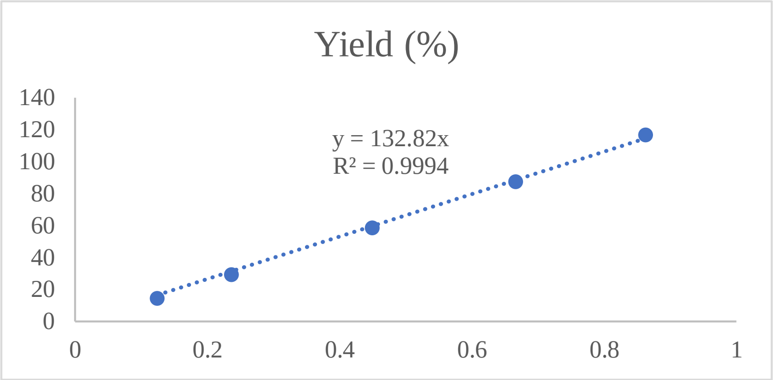

Enantiomeric ratio was determined by chiral HPLC using a Chiralpak IC column (4.6 x 250 mm, 5 μm), (HPLC: IF, 210 nm, n-hexane/isopropanol = 90:10, flow rate 1 mL/min, 25 °C, Retention Time: 9.823 min, 11.099 min).

HPLC trace of rac-**3i**:

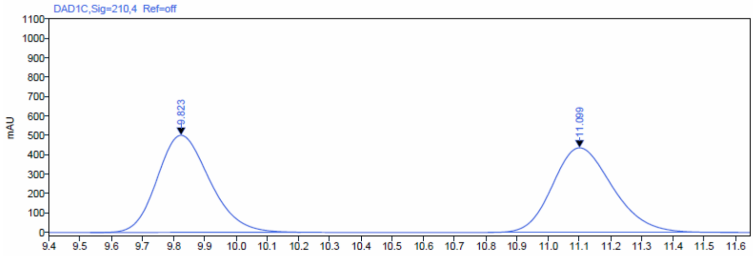

| Signal: | DAD1C,Sig=210,4 Ref=off | | | | | |
|---|---|---|---|---|---|---|
| RT [min] | Type | Width [min] | Area | Height | Area% | Name |
| 9.823 | BBA | 0.67 | 5841.62 | 500.02 | 50.03 | |
| 11.099 | BBA | 0.80 | 5835.07 | 436.08 | 49.97 | |

HPLC trace of enantioenriched-**3i** obtained from the reaction catalyzed by SYE2:

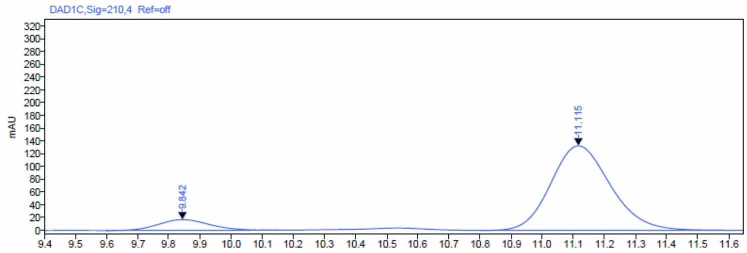

| Signal: | DAD1C,Sig=210,4 Ref=off | | | | | |
|---|---|---|---|---|---|---|
| RT [min] | Type | Width [min] | Area | Height | Area% | Name |
| 9.842 | BB | 0.55 | 190.72 | 16.73 | 9.78 | |
| 11.115 | BBA | 0.80 | 1759.43 | 132.20 | 90.22 | |

HPLC trace of enantioenriched-**3i** obtained from the reaction catalyzed by GluER-Y343A:

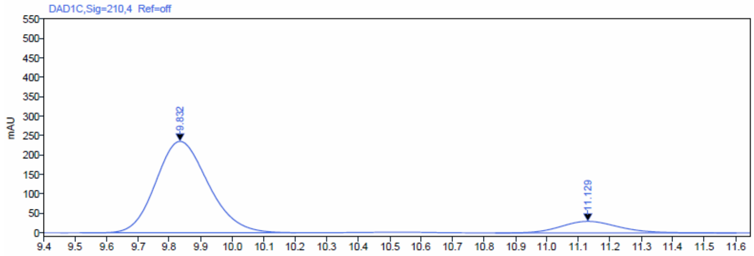

| Signal: | DAD1C,Sig=210,4 Ref=off | | | | | |
|---|---|---|---|---|---|---|
| RT [min] | Type | Width [min] | Area | Height | Area% | Name |
| 9.832 | BBA | 0.68 | 2721.52 | 235.01 | 87.78 | |
| 11.129 | BBA | 0.77 | 378.88 | 29.70 | 12.22 | |

Rac-**3i** is a colorless oil.

$^{1}H$ NMR (600 MHz, $CDCl_3$) δ 7.29 (dd, J = 8.2, 7.6 Hz, 1H), 6.83 (ddd, J = 8.3, 2.6, 0.9 Hz, 1H), 6.79 (dt, J = 7.6, 1.3 Hz, 1H), 6.74 (dd, J = 2.5, 1.7 Hz, 1H), 3.81 (s, 3H), 3.33 (dd, J = 11.0, 4.9 Hz, 1H), 3.03 – 2.97 (m, 1H), 2.36 (ddd, J = 13.6, 11.0, 4.7 Hz, 1H), 2.18 (ddd, J = 13.6, 11.2, 4.9 Hz, 1H), 1.37 (d, J = 7.0 Hz, 3H).

$^{13}C$ NMR (151 MHz, $CDCl_3$) δ 160.31, 143.76, 130.49, 119.02, 112.92, 112.81, 112.68, 112.35, 55.28, 38.86, 37.61, 21.79, 21.05.

HRMS (ESI, m/z) calcd for $C_{13}H_{15}N_2O$ $[M+H]^+$: 215.1184, found: 215.1193.

HRMS spectrum of **3i**:

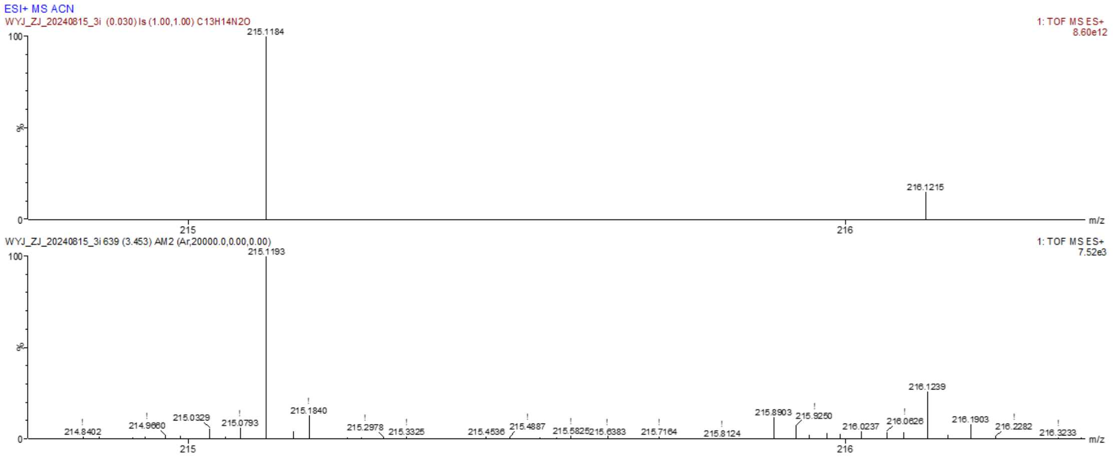

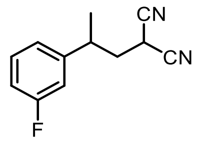

**3j**

2-(2-(3-Fluorophenyl) propyl) malononitrile **3j**

By using the typical procedure described in Section 5, the reaction of malononitrile **1a** (0.008 mmol) and 1-fluoro-3-(prop-1-en-2-yl)benzene **2j** (0.004 mmol) catalyzed by SYE2 (1 mol%) with FI (5 mol%) afforded **3j** in 35.6% reverse phase HPLC yield (average of duplicate runs), with a 94.8:5.2 er. The corresponding reaction catalyzed by GluER-Y343A (1 mol%) afforded **3j** in 44.8% reverse phase HPLC yield (average of duplicate runs), with a > 1:99 er (the other enantiomer as the major).

Reverse phase HPLC calibration curve for **3j** is displayed below:

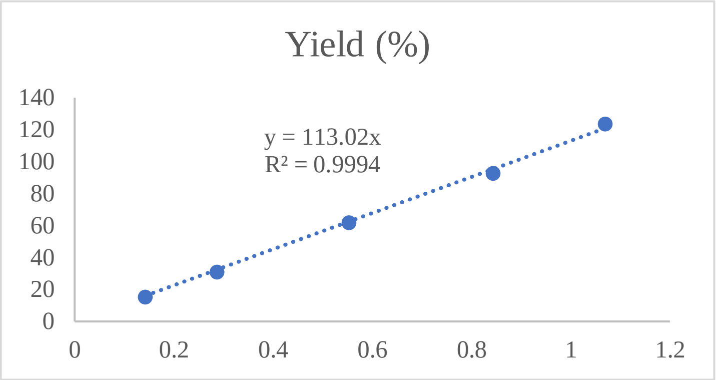

Enantiomeric ratio was determined by chiral HPLC using a Chiralpak IC column (4.6 x 250 mm, 5 μm), (HPLC: IF, 210 nm, n-hexane/isopropanol = 90:10, flow rate 1 mL/min, 25 °C, Retention Time: 9.161 min, 11.154 min).

HPLC trace of rac-**3j**:

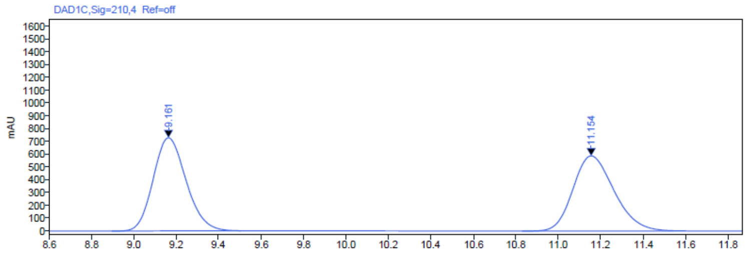

Signal: DAD1C,Sig=210,4 Ref=off

| RT [min] | Type | Width [min] | Area | Height | Area% | Name |
|---|---|---|---|---|---|---|
| 9.161 | BBA | 0.61 | 7843.96 | 727.41 | 49.87 | |
| 11.154 | BBA | 0.77 | 7884.57 | 586.33 | 50.13 | |

HPLC trace of enantioenriched-**3j** obtained from the reaction catalyzed by SYE2:

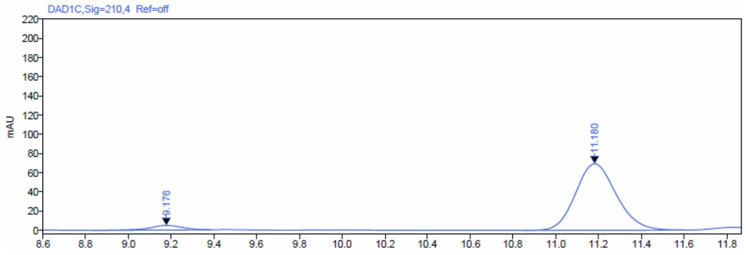

Signal: DAD1C,Sig=210,4 Ref=off

| RT [min] | Type | Width [min] | Area | Height | Area% | Name |
|---|---|---|---|---|---|---|
| 9.176 | BV | 0.56 | 50.30 | 4.73 | 5.21 | |
| 11.180 | BBA | 0.75 | 915.21 | 69.15 | 94.79 | |

HPLC trace of enantioenriched-**3j** obtained from the reaction catalyzed by GluER-Y343A:

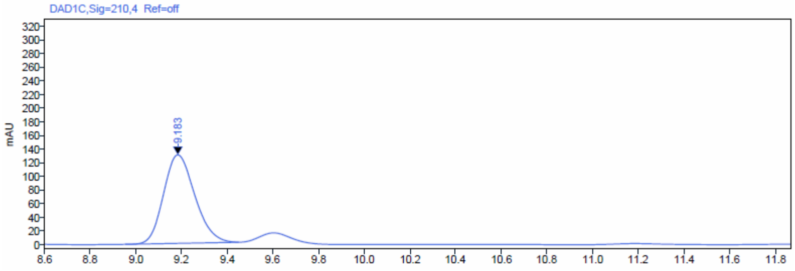

Signal: DAD1C,Sig=210,4 Ref=off

| RT [min] | Type | Width [min] | Area | Height | Area% | Name |
|---|---|---|---|---|---|---|
| 9.183 | BB | 0.50 | 1229.94 | 129.59 | 100.00 | |

Rac-**3j** is a colorless oil.

[1]H NMR (600 MHz, $CDCl_3$) δ 7.34 (td, J = 8.0, 6.0 Hz, 1H), 7.00 (dddd, J = 12.9, 8.4, 2.7, 1.1 Hz, 2H), 6.92 (ddd, J = 9.8, 2.6, 1.8 Hz, 1H), 3.35 (dd, J = 10.6, 5.3 Hz, 1H), 3.08 – 3.01 (m, 1H), 2.37 (ddd, J = 13.7, 10.6, 4.9 Hz, 1H), 2.20 (ddd, J = 13.7, 10.8, 5.3 Hz, 1H), 1.38 (d, J = 7.0 Hz, 3H).

[13]C NMR (151 MHz, $CDCl_3$) δ 164.16, 162.52, 144.85, 131.03, 122.75, 114.73, 113.79, 112.57, 112.17, 38.59, 37.37, 21.69, 21.04.

[19]F NMR (565 MHz, $CDCl_3$) δ -111.37.

HRMS (APCI, m/z) calcd for $C_{12}H_{12}FN_2$ $[M+H]^+$: 203.0984, found: 203.1010.

HRMS spectrum of **3j**:

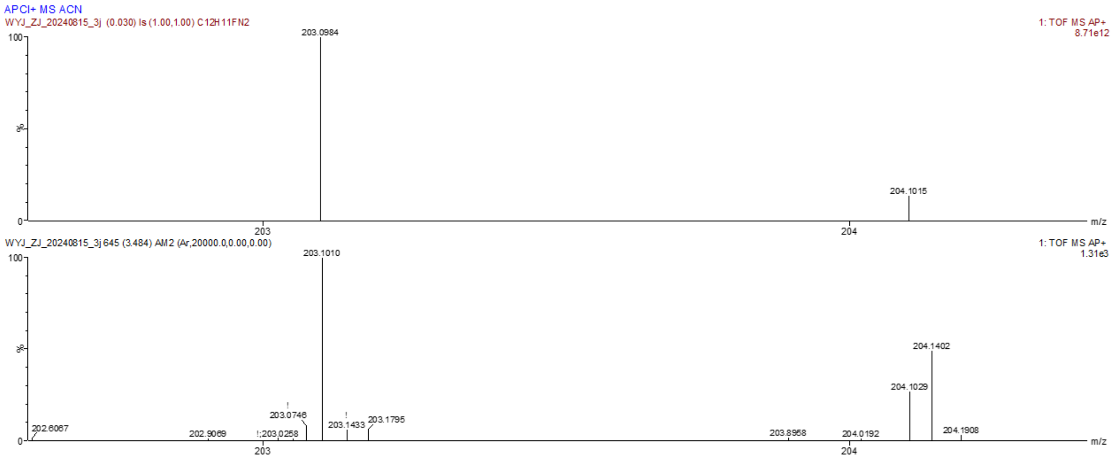

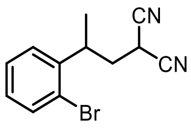

**3k**

2-(2-(2-Bromophenyl) propyl) malononitrile **3k**

By using the typical procedure described in Section 5, the reaction of malononitrile **1a** (0.008 mmol) and 1-bromo-2-(prop-1-en-2-yl)benzene **2k** (0.004 mmol) catalyzed by SYE2 (1 mol%) with FI (5 mol%) afforded **3k** in 4.7% reverse phase HPLC yield (average of duplicate runs), with a 85.7:14.3 er. The corresponding reaction catalyzed by GluER-Y343A (1 mol%) afforded **3j** in 4.1% reverse phase HPLC yield (average of duplicate runs), with a > 1:99 er (the other enantiomer as the major).

Reverse phase HPLC calibration curve for **3k** is displayed below:

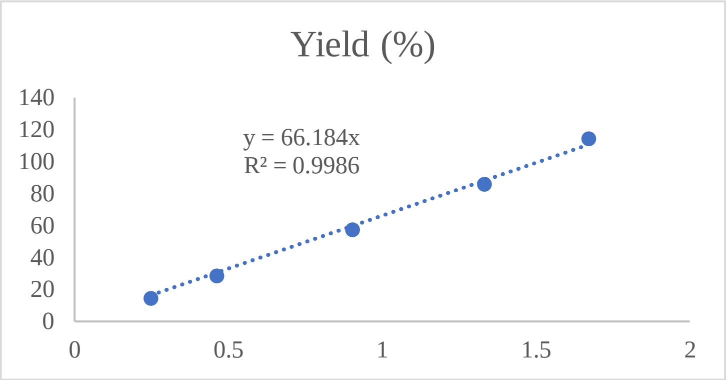

Enantiomeric ratio was determined by chiral HPLC using a Chiralpak IC column (4.6 x 250 mm, 5 μm), (HPLC: IF, 210 nm, n-hexane/isopropanol = 90:10, flow rate 1 mL/min, 25 °C, Retention Time: 9.716 min, 12.860 min).

HPLC trace of rac-**3k**:

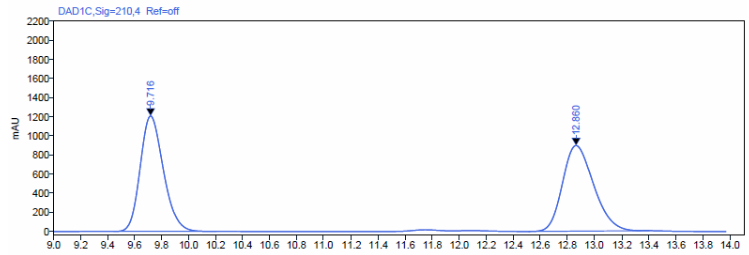

Signal: DAD1C,Sig=210,4 Ref=off

| RT [min] | Type | Width [min] | Area | Height | Area% | Name |
|---|---|---|---|---|---|---|
| 9.716 | BBA | 0.76 | 14375.61 | 1206.98 | 50.13 | |
| 12.860 | BBA | 0.89 | 14300.02 | 895.07 | 49.87 | |

HPLC trace of enantioenriched-**3k** obtained from the reaction catalyzed by SYE2:

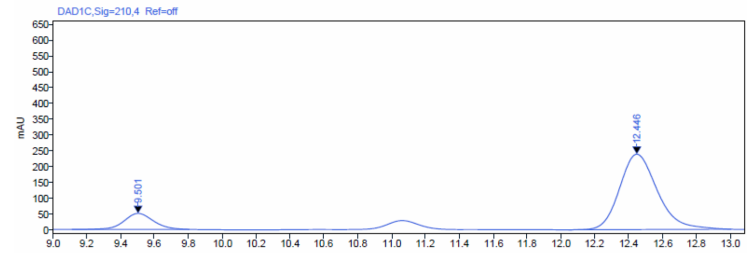

Signal: DAD1C,Sig=210,4 Ref=off

| RT [min] | Type | Width [min] | Area | Height | Area% | Name |
|---|---|---|---|---|---|---|
| 9.501 | BBA | 0.69 | 616.10 | 50.71 | 14.29 | |
| 12.446 | BBA | 0.96 | 3694.90 | 238.33 | 85.71 | |

HPLC trace of enantioenriched-**3k** obtained from the reaction catalyzed by GluER-Y343A:

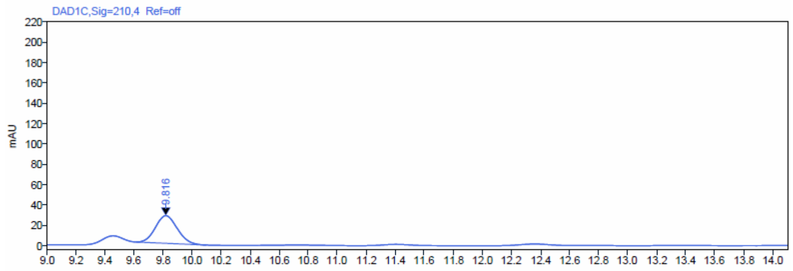

Signal: DAD1C,Sig=210,4 Ref=off

| RT [min] | Type | Width [min] | Area | Height | Area% | Name |
|---|---|---|---|---|---|---|
| 9.816 | BBA | 0.48 | 283.78 | 27.13 | 100.00 | |

Rac-**3k** is a colorless oil.

[1]H NMR (600 MHz, $CDCl_3$) δ 7.60 (dd, J = 8.0, 1.3 Hz, 1H), 7.35 (td, J = 7.5, 1.3 Hz, 1H), 7.21 (dd, J = 7.8, 1.7 Hz, 1H), 7.15 (ddd, J = 8.0, 7.3, 1.7 Hz, 1H), 3.65 (dp, J = 9.2, 6.8 Hz, 1H), 3.53 (dd, J = 9.1, 6.2 Hz, 1H), 2.41 (ddd, J = 13.8, 9.2, 6.3 Hz, 1H), 2.32 (ddd, J = 13.9, 9.1, 6.0 Hz, 1H), 1.37 (d, J = 7.0 Hz, 3H).

[13]C NMR (151 MHz, $CDCl_3$) δ 141.49, 133.75, 129.03, 128.45, 127.05, 124.66, 112.55, 38.01, 36.04, 20.86, 20.75.

HRMS (APCI, m/z) calcd for $C_{12}H_{12}BrN_2$ $[M+H]^+$: 263.0184, found: 263.0167.

HRMS spectrum of **3k**:

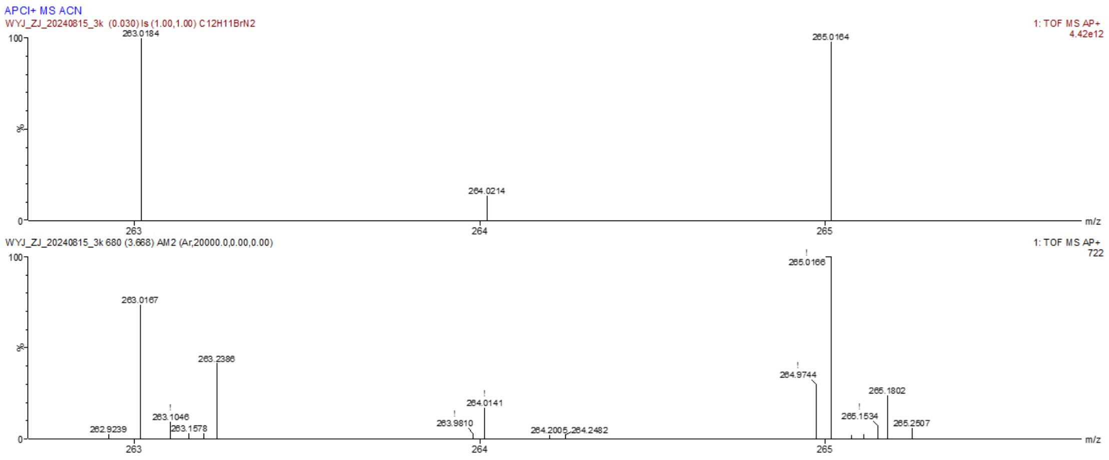

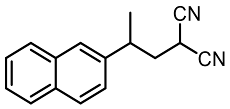

**3l**

2-(2-(Naphthalen-2-yl) propyl) malononitrile **3l**

By using the typical procedure described in Section 5, the reaction of malononitrile **1a** (0.008 mmol) and 2-(prop-1-en-2-yl)naphthalene **2l** (0.004 mmol) catalyzed by SYE2 (1 mol%) with FI (5 mol%) afforded **3l** in 36.5% reverse phase HPLC yield (average of duplicate runs), with a 91:9 er. The corresponding reaction catalyzed by GluER-Y343A (1 mol%) afforded **3l** in 37.6% reverse phase HPLC yield (average of duplicate runs), with a 14.5:85.5 er (the other enantiomer as the major).

Reverse phase HPLC calibration curve for **3l** is displayed below:

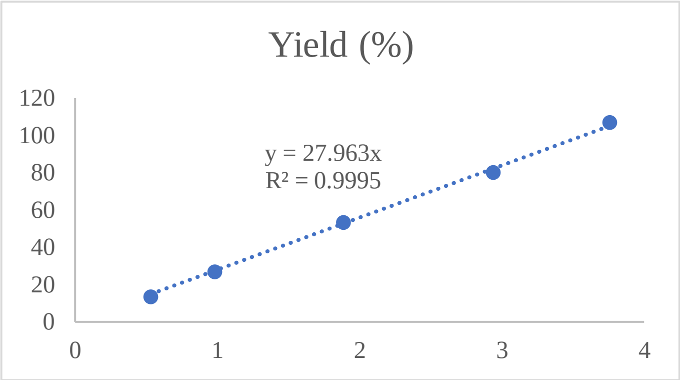

Enantiomeric ratio was determined by chiral HPLC using a Chiralpak IC column (4.6 x 250 mm, 5 μm), (HPLC: IF, 210 nm, n-hexane/isopropanol = 90:10, flow rate 1 mL/min, 25 °C, Retention Time: 10.472 min, 11.954 min).

HPLC trace of rac-**3l**:

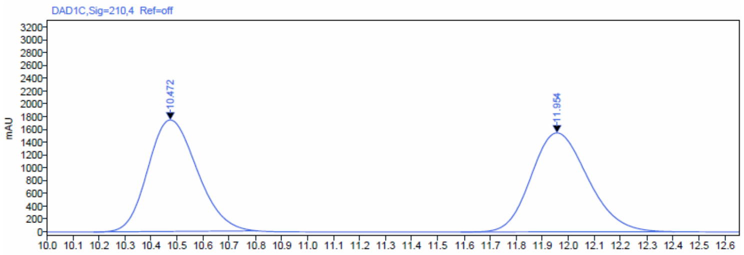

**Signal:** DAD1C,Sig=210,4 Ref=off

| RT [min] | Type | Width [min] | Area | Height | Area% | Name |
|---|---|---|---|---|---|---|
| 10.472 | BBA | 0.63 | 22823.29 | 1741.49 | 49.43 | |
| 11.954 | BBA | 0.82 | 23346.72 | 1545.28 | 50.57 | |

HPLC trace of enantioenriched-**3l** obtained from the reaction catalyzed by SYE2:

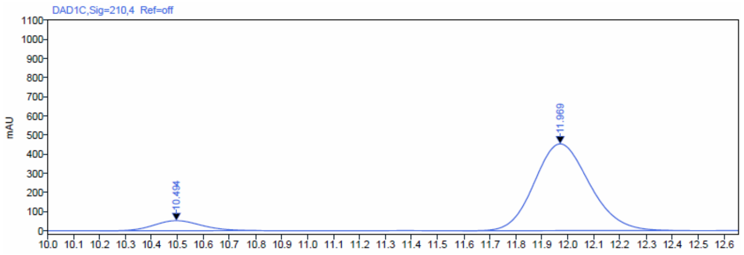

**Signal:** DAD1C,Sig=210,4 Ref=off

| RT [min] | Type | Width [min] | Area | Height | Area% | Name |
|---|---|---|---|---|---|---|
| 10.494 | BBA | 0.60 | 661.13 | 52.35 | 9.02 | |
| 11.969 | BBA | 0.81 | 6668.57 | 452.75 | 90.98 | |

HPLC trace of enantioenriched-**3l** obtained from the reaction catalyzed by GluER-Y343A:

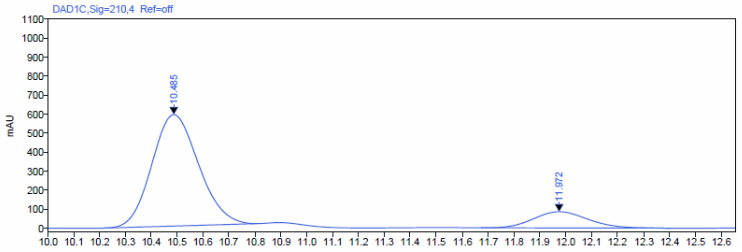

**Signal:** DAD1C,Sig=210,4 Ref=off

| RT [min] | Type | Width [min] | Area | Height | Area% | Name |
|---|---|---|---|---|---|---|
| 10.485 | BBA | 0.59 | 7334.00 | 585.59 | 85.53 | |
| 11.972 | BBA | 0.73 | 1240.56 | 85.28 | 14.47 | |

Rac-**3l** is a colorless oil.

$^1$H NMR (600 MHz, $CDCl_3$) δ 7.85 (d, J = 8.5 Hz, 1H), 7.83 – 7.78 (m, 2H), 7.66 – 7.62 (m, 1H), 7.51 – 7.45 (m, 2H), 7.29 (dd, J = 8.5, 1.9 Hz, 1H), 3.26 (dd, J = 11.0, 5.0 Hz, 1H), 3.21 – 3.14 (m, 1H), 2.40 (ddd, J = 13.7, 11.0, 4.7 Hz, 1H), 2.27 (ddd, J = 13.7, 11.1, 5.1 Hz, 1H), 1.44 (d, J = 7.0 Hz, 3H).

$^{13}$C NMR (151 MHz, $CDCl_3$) δ 139.40, 133.59, 132.89, 129.49, 127.80, 127.76, 126.73, 126.27, 126.22, 124.11, 112.82, 112.40, 38.69, 37.74, 21.91, 21.10.

HRMS (APCI, m/z) calcd for $C_{16}H_{15}N_2$ $[M+H]^+$: 235.1235, found: 235.1243.

HRMS spectrum of **3l**:

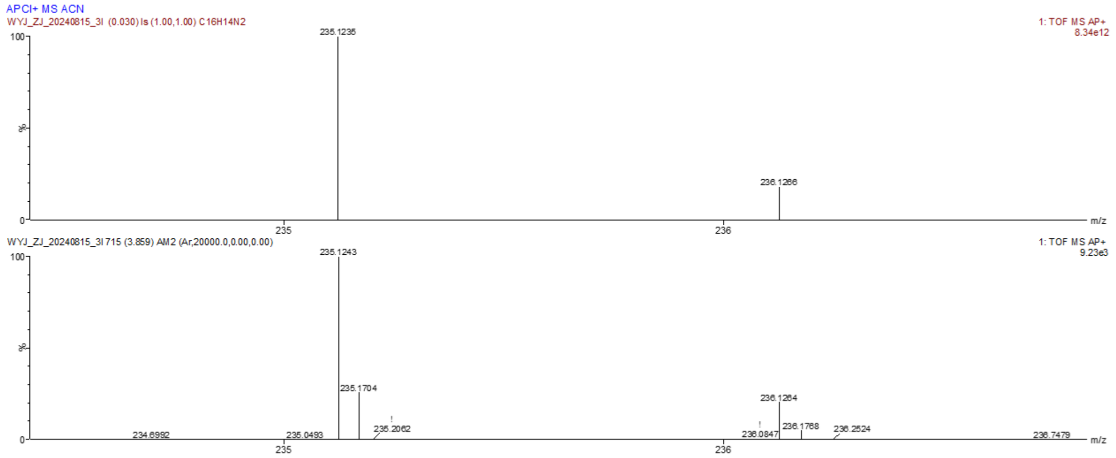

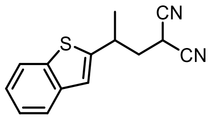

**3m**

2-(2-(benzo[b]thiophen-2-yl)propyl)malononitrile **3m**

By using the typical procedure described in Section 5, the reaction of malononitrile **1a** (1 mmol) and 2-(prop-1-en-2-yl)benzo[b]thiophene **2m** (0.5 mmol) catalyzed by SYE2 (1 mol%) with FI (5 mol%) afforded **3m** in 26.2% isolated yield (average of duplicate runs), with a 80:20 er. The corresponding reaction catalyzed by GluER-Y343A (1 mol%) afforded **3m** in 27.5% isolated yield (average of duplicate runs), with a 5.3:94.7 er (the other enantiomer as the major).

Enantiomeric ratio was determined by chiral HPLC using a Chiralpak IC column (4.6 x 250 mm, 5 μm), (HPLC: IF, 210 nm, n-hexane/isopropanol = 90:10, flow rate 1 mL/min, 25 °C, Retention Time: 11.547 min, 12.837 min).

HPLC trace of rac-**3m**, which was obtained from scale up reaction with GluER-WT.

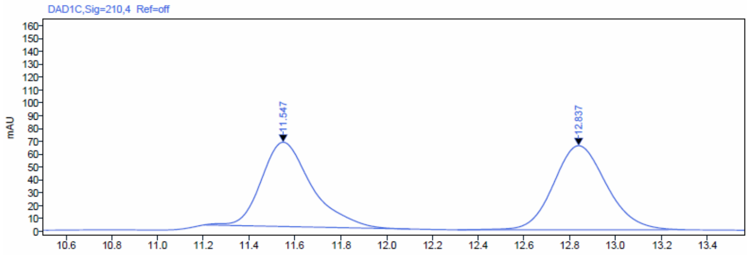

Signal: DAD1C,Sig=210,4 Ref=off

| RT [min] | Type | Width [min] | Area | Height | Area% | Name |
|---|---|---|---|---|---|---|
| 11.547 | BBA | 0.90 | 1002.09 | 65.48 | 48.84 | |
| 12.837 | BBA | 0.99 | 1049.56 | 65.28 | 51.16 | |

HPLC trace of enantioenriched-**3m** obtained from the reaction catalyzed by SYE2:

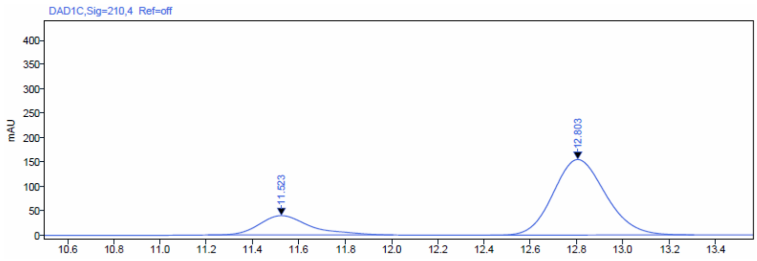

Signal: DAD1C,Sig=210,4 Ref=off

| RT [min] | Type | Width [min] | Area | Height | Area% | Name |
|---|---|---|---|---|---|---|
| 11.523 | BBA | 0.80 | 608.09 | 39.76 | 19.99 | |
| 12.803 | BBA | 0.88 | 2434.58 | 154.89 | 80.01 | |

HPLC trace of enantioenriched-**3m** obtained from the reaction catalyzed by GluER-Y343A:

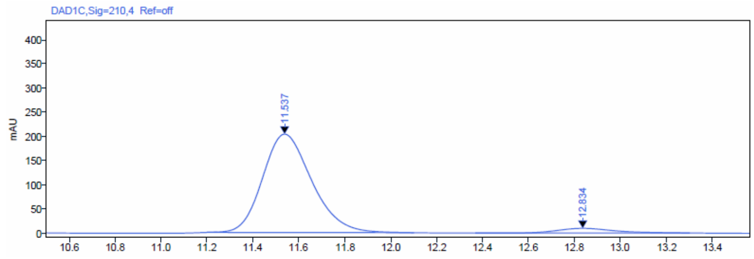

Signal: DAD1C,Sig=210,4 Ref=off

| RT [min] | Type | Width [min] | Area | Height | Area% | Name |
|---|---|---|---|---|---|---|
| 11.537 | BBA | 0.90 | 3024.99 | 203.37 | 94.71 | |
| 12.834 | BBA | 0.93 | 168.89 | 9.77 | 5.29 | |

Rac-**3m** is a colorless oil.

$^{1}$H NMR (600 MHz, $CDCl_3$) δ 7.80 (dt, *J* = 7.9, 1.0 Hz, 1H), 7.72 (dt, *J* = 7.9, 1.0 Hz, 1H), 7.34 (dddd, *J* = 24.7, 8.3, 7.2, 1.2 Hz, 2H), 7.17 (s, 1H), 3.53 – 3.42 (m, 2H), 2.47 – 2.42 (m, 1H), 2.25 (ddd, *J* = 13.7, 11.1, 4.9 Hz, 1H), 1.51 (d, *J* = 6.9 Hz, 3H).

$^{13}$C NMR (151 MHz, $CDCl_3$) δ 146.24, 139.37, 138.96, 124.81, 124.65, 123.49, 122.54, 122.05, 112.50, 112.15, 39.21, 34.19, 22.73, 21.04.

HRMS (ESI, m/z) calcd for $C_{14}H_{13}N_2S$ $[M+H]^+$: 241.0799, found: 241.0797.

HRMS spectrum of **3m**:

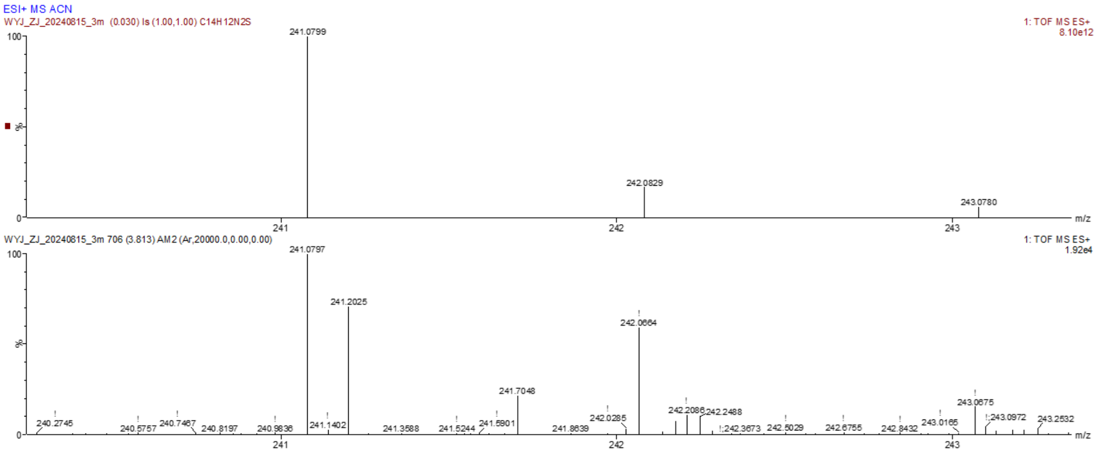

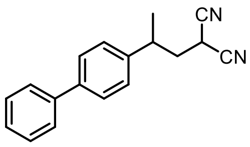

**3n**

2-(2-([1,1'-Biphenyl]-4-yl) propyl) malononitrile **3n**

By using the typical procedure described in Section 5, the reaction of malononitrile **1a** (0.008 mmol) and 4-(prop-1-en-2-yl)-1,1'-biphenyl **2n** (0.004 mmol) catalyzed by SYE2 (1 mol%) with FI (5 mol%) afforded **3n** in 9.4% reverse phase HPLC yield (average of duplicate runs), with a 54.3:45.7 er. The corresponding reaction catalyzed by GluER-Y343A (1 mol%) afforded **3n** in 21.7% reverse phase HPLC yield (average of duplicate runs), with a 96.8:3.2 er.

Reverse phase HPLC calibration curve for **3n** is displayed below:

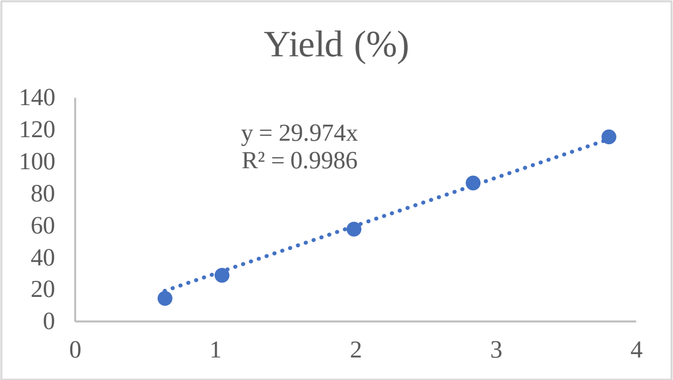

Enantiomeric ratio was determined by chiral HPLC using a Chiralpak IC column (4.6 x 250 mm, 5 µm), (HPLC: IF, 210 nm, n-hexane/isopropanol = 90:10, flow rate 1 mL/min, 25 °C, Retention Time: 13.136 min, 13.799 min).

HPLC trace of rac-**3n**:

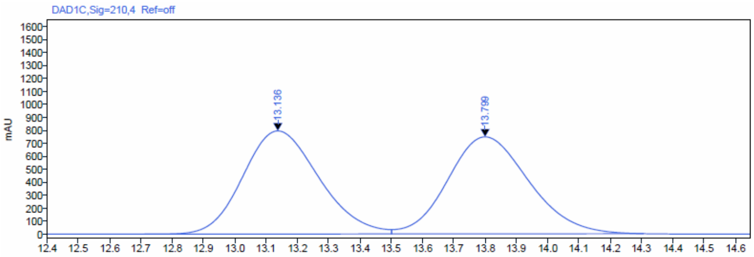

Signal: DAD1C,Sig=210,4 Ref=off

| RT [min] | Type | Width [min] | Area | Height | Area% | Name |
|---|---|---|---|---|---|---|
| 13.136 | BV | 0.79 | 13433.47 | 793.52 | 49.95 | |
| 13.799 | VBA | 0.80 | 13457.99 | 746.19 | 50.05 | |

HPLC trace of enantioenriched-**3n** obtained from the reaction catalyzed by SYE2:

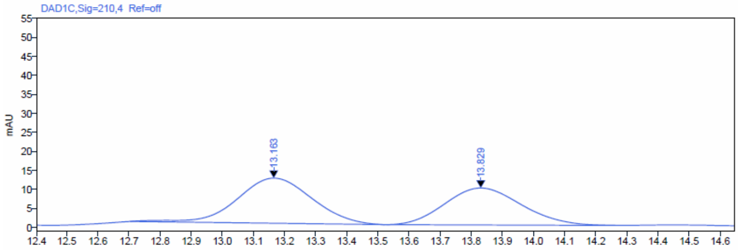

Signal: DAD1C,Sig=210,4 Ref=off

| RT [min] | Type | Width [min] | Area | Height | Area% | Name |
|---|---|---|---|---|---|---|
| 13.163 | BB | 0.82 | 197.80 | 11.85 | 54.33 | |
| 13.829 | BB | 0.74 | 166.25 | 9.71 | 45.67 | |

HPLC trace of enantioenriched-**3n** obtained from the reaction catalyzed by GluER-Y343A:

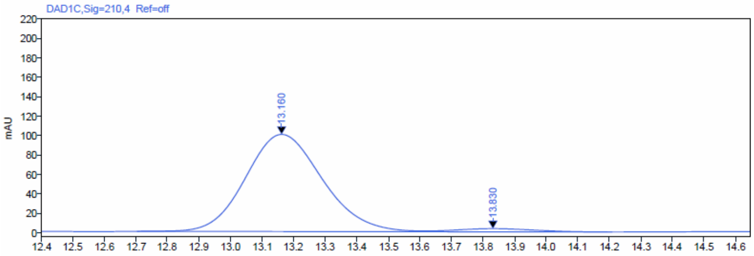

Signal: DAD1C,Sig=210,4 Ref=off

| RT [min] | Type | Width [min] | Area | Height | Area% | Name |
|---|---|---|---|---|---|---|
| 13.160 | BV | 0.92 | 1690.08 | 99.83 | 96.79 | |
| 13.830 | VB | 0.59 | 56.05 | 3.31 | 3.21 | |

Rac-**3n** is a white solid.

$^{1}H$ NMR (600 MHz, $CDCl_3$) δ 7.66 – 7.61 (m, 4H), 7.50 (t, *J* = 7.7 Hz, 2H), 7.44 – 7.38 (m, 1H), 7.34 – 7.31 (m, 2H), 3.41 (dd, *J* = 11.0, 5.1 Hz, 1H), 3.15 – 3.09 (m, 1H), 2.44 (ddd, *J* = 13.6, 11.0, 4.8 Hz, 1H), 2.28 (ddd, *J* = 13.6, 11.0, 5.1 Hz, 1H), 1.46 (d, *J* = 7.0 Hz, 3H).

$^{13}C$ NMR (151 MHz, $CDCl_3$) δ 141.11, 140.79, 140.42, 128.93, 128.10, 127.57, 127.35, 127.08, 112.83, 112.37, 38.86, 37.26, 21.92, 21.13.

HRMS (APCI, m/z) calcd for $C_{18}H_{17}N_2$ $[M+H]^+$: 261.1392, found: 261.1376.

HRMS spectrum of **3n**:

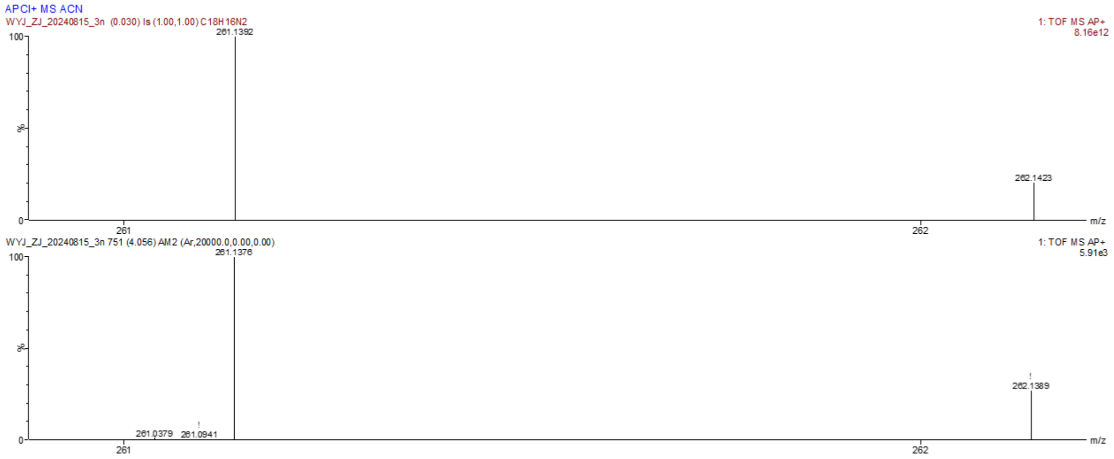

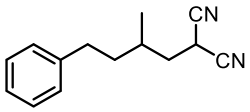

**3o**

2-(2-methyl-4-phenylbutyl)malononitrile **3o**

By using the typical procedure described in Section 5, the reaction of malononitrile **1a** (0.008 mmol) and (3-methylbut-3-en-1-yl)benzene **2o** (0.004 mmol) catalyzed by SYE2 (1 mol%) with FI (5 mol%) afforded **3o** in 9% GC yield (average of duplicate runs), with a 65.8:34.2 er. The corresponding reaction catalyzed by GluER-Y343A (1 mol%) afforded **3o** in 18% GC yield (average of duplicate runs), with a 54.9:45.1 er.

GC calibration curve for **3o** is displayed below:

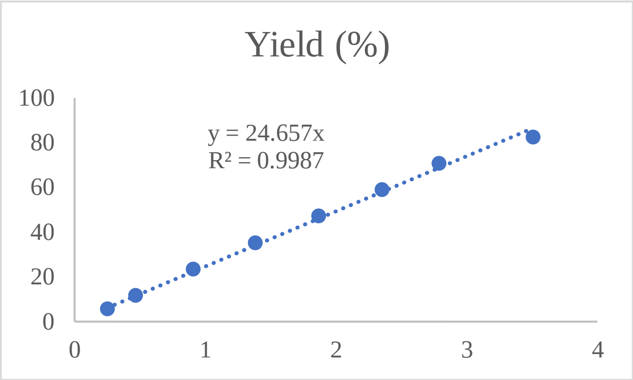

Enantiomeric ratio was determined by chiral HPLC using a Chiralpak IC column (4.6 x 250 mm, 5 μm), (HPLC: IF, 210 nm, n-hexane/isopropanol = 90:10, flow rate 1 mL/min, 25 ºC, Retention Time: 10.200 min, 11.137 min).

HPLC trace of rac-**3o**:

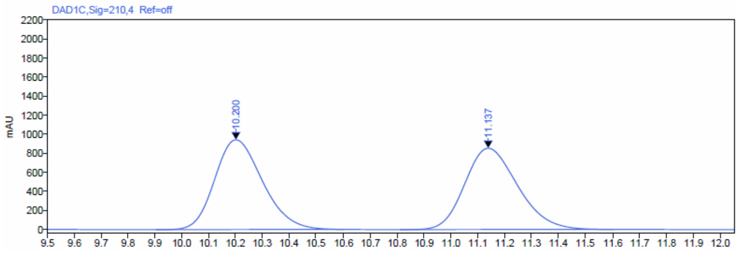

| Signal: | DAD1C,Sig=210,4 Ref=off | | | | | |
|---|---|---|---|---|---|---|
| RT [min] | Type | Width [min] | Area | Height | Area% | Name |
| 10.200 | BBA | 0.70 | 11631.00 | 941.18 | 49.94 | |
| 11.137 | BBA | 0.79 | 11658.78 | 852.49 | 50.06 | |

HPLC trace of enantioenriched-**3o** obtained from the reaction catalyzed by SYE2:

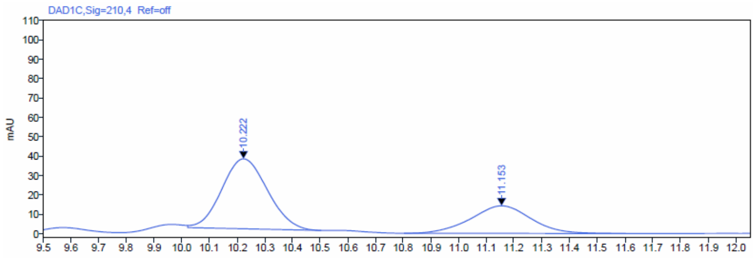

| Signal: | DAD1C,Sig=210,4 Ref=off | | | | | |
|---|---|---|---|---|---|---|
| RT [min] | Type | Width [min] | Area | Height | Area% | Name |
| 10.222 | VB | 0.48 | 413.22 | 36.04 | 65.86 | |
| 11.153 | BBA | 0.80 | 214.19 | 14.19 | 34.14 | |

HPLC trace of enantioenriched-**3o** obtained from the reaction catalyzed by GluER-W66Q-Y343A:

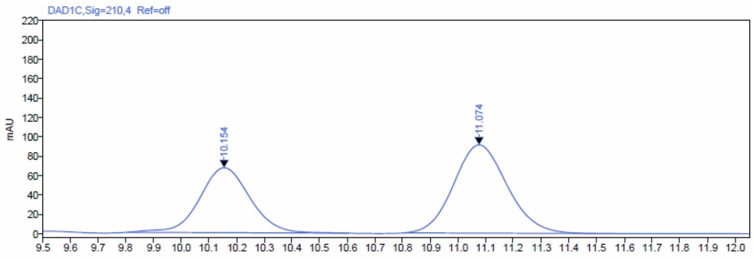

| Signal: | DAD1C,Sig=210,4 Ref=off | | | | | |
|---|---|---|---|---|---|---|
| RT [min] | Type | Width [min] | Area | Height | Area% | Name |
| 10.154 | BBA | 0.80 | 848.41 | 66.94 | 40.93 | |
| 11.074 | BBA | 0.76 | 1224.61 | 90.89 | 59.07 | |

Rac-**3o** is a colorless oil.

$^1$H NMR (600 MHz, $CDCl_3$) δ 7.29 (t, *J* = 7.6 Hz, 2H), 7.23 – 7.14 (m, 3H), 3.69 (dd, *J* = 8.8, 6.2 Hz, 1H), 2.70 (ddd, *J* = 13.7, 10.3, 5.7 Hz, 1H), 2.60 (ddd, *J* = 13.7, 10.3, 6.0 Hz, 1H), 2.13 – 2.06 (m, 1H), 1.88 – 1.80 (m, 2H), 1.72 – 1.65 (m, 1H), 1.57 – 1.52 (m, 1H), 1.05 (d, *J* = 6.2 Hz, 3H).

$^{13}$C NMR (151 MHz, $CDCl_3$) δ 141.48, 128.61, 128.30, 126.18, 112.86, 112.66, 37.88, 37.56, 32.95, 30.59, 20.72, 18.58.

HRMS (APCI, m/z) calcd for $C_{14}H_{17}N_2$ $[M+H]^+$: 213.1392, found: 213.1393.

HRMS spectrum of **3o**:

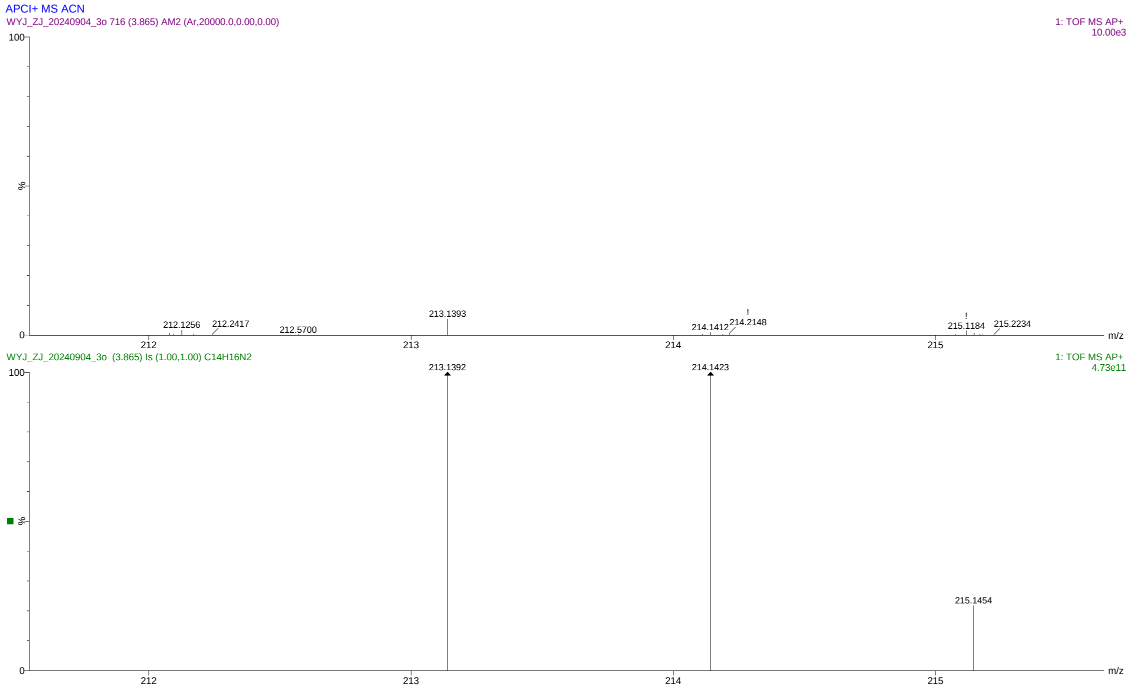

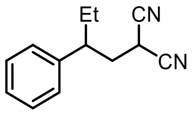

**3p**

2-(2-Phenylbutyl) malononitrile **3p**

By using the typical procedure described in Section 5, the reaction of malononitrile **1a** (0.008 mmol) and but-1-en-2-ylbenzene **2p** (0.004 mmol) catalyzed by SYE2 (1 mol%) with **FI** (5 mol%) afforded **3p** in 39.5% reverse phase HPLC yield (average of duplicate runs), with a 97.8:2.2 er. The corresponding reaction catalyzed by GluER-Y343A (1 mol%) afforded **3p** in 63.8% reverse phase HPLC yield (average of duplicate runs), with a 3.8:96.2 er (the other enantiomer as the major).

Reverse phase HPLC calibration curve for **3p** is displayed below:

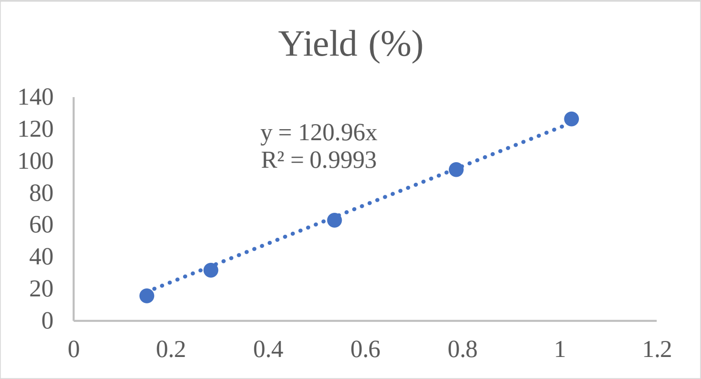

Enantiomeric ratio was determined by chiral HPLC using a Chiralpak IC column (4.6 x 250 mm, 5 μm), (HPLC: IF, 210 nm, n-hexane/isopropanol = 90:10, flow rate 1 mL/min, 25 °C, Retention Time: 7.775 min, 8.346 min).

HPLC trace of rac-**3p**:

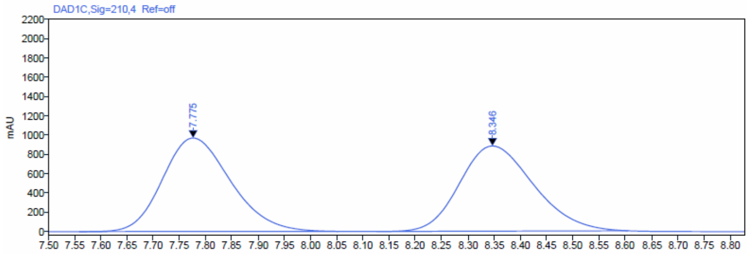

| Signal: | DAD1C,Sig=210,4 Ref=off | | | | | |
|---|---|---|---|---|---|---|
| **RT [min]** | **Type** | **Width [min]** | **Area** | **Height** | **Area%** | **Name** |
| 7.775 | BBA | 0.49 | 8750.29 | 969.28 | 50.21 | |
| 8.346 | BBA | 0.48 | 8675.60 | 883.52 | 49.79 | |

HPLC trace of enantioenriched-**3p** obtained from the reaction catalyzed by SYE2:

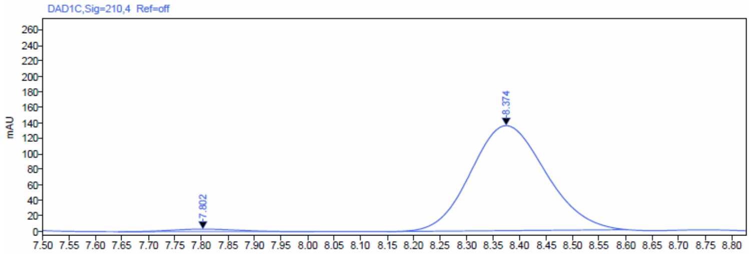

| Signal: | DAD1C,Sig=210,4 Ref=off | | | | | |
|---|---|---|---|---|---|---|
| **RT [min]** | **Type** | **Width [min]** | **Area** | **Height** | **Area%** | **Name** |
| 7.802 | BB | 0.37 | 29.51 | 3.30 | 2.16 | |
| 8.374 | BBA | 0.45 | 1335.04 | 135.69 | 97.84 | |

HPLC trace of enantioenriched-**3p** obtained from the reaction catalyzed by GluER-Y343A:

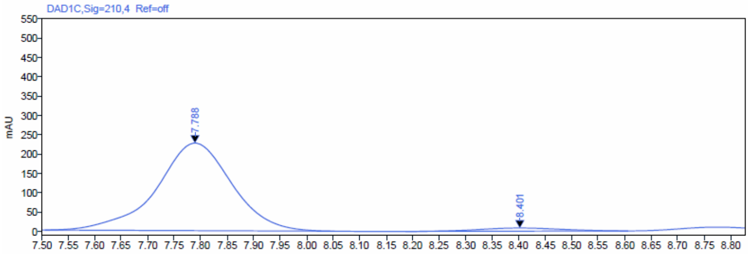

| Signal: | DAD1C,Sig=210,4 Ref=off | | | | | |
|---|---|---|---|---|---|---|
| **RT [min]** | **Type** | **Width [min]** | **Area** | **Height** | **Area%** | **Name** |
| 7.788 | BBA | 0.55 | 2229.11 | 226.35 | 96.25 | |
| 8.401 | BBA | 0.42 | 86.79 | 8.22 | 3.75 | |

Rac-**3p** is a colorless oil.

$^{1}H$ NMR (600 MHz, $CDCl_3$) δ 7.36 (t, J = 7.7 Hz, 2H), 7.29 (td, J = 7.2, 1.4 Hz, 1H), 7.19 – 7.15 (m, 2H), 3.26 (dd, J = 11.5, 4.6 Hz, 1H), 2.73 (m, J = 11.4, 9.4, 4.9 Hz, 1H), 2.43 (ddd, J = 13.2, 11.5, 4.1 Hz, 1H), 2.20 – 2.15 (m, 1H), 1.77 – 1.67 (m, 2H), 0.83 (t, J = 7.3 Hz, 3H).

$^{13}C$ NMR (151 MHz, $CDCl_3$) δ 140.45, 129.37, 127.82, 127.53, 112.91, 112.32, 45.09, 37.46, 29.24, 20.99, 11.94.

HRMS (APCI, m/z) calcd for $C_{13}H_{15}N_2$ $[M+H]^+$: 199.1235, found: 199.1227.

HRMS spectrum of **3p**:

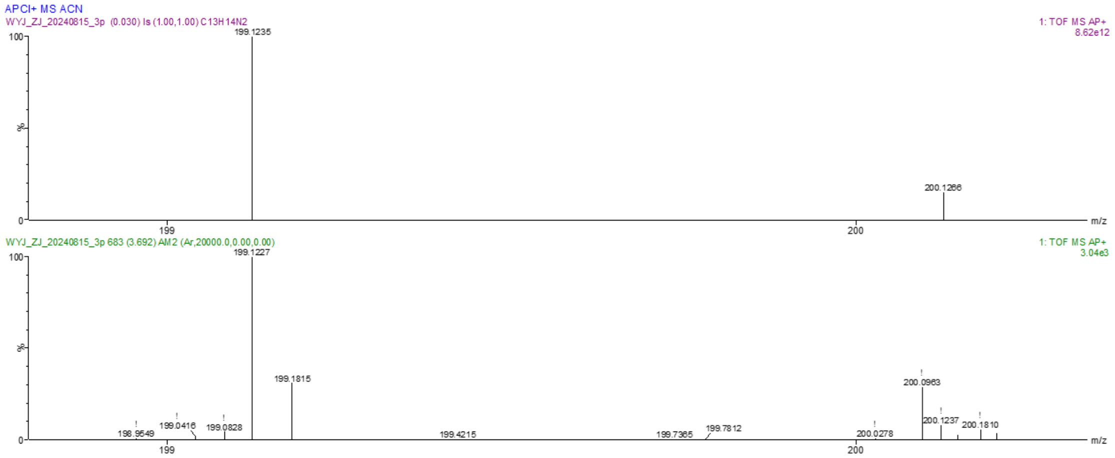

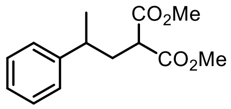

**3q**

dimethyl 2-(2-phenylpropyl)malonate **3q**

By using the typical procedure described in Section 5, the reaction of dimethyl malonate **1b** (0.008 mmol) and prop-1-en-2-ylbenzene **2a** (0.004 mmol) catalyzed by SYE2 (1 mol%) with FI (5 mol%) afforded **3q** in 46.9% GC yield (average of duplicate runs), with a 96:4 er. The corresponding reaction catalyzed by GluER-Y343A (1 mol%) afforded **3q** in 75.9% GC yield (average of duplicate runs), with a > 1:99 er (the other enantiomer as the major).

GC calibration curve for **3q** is displayed below:

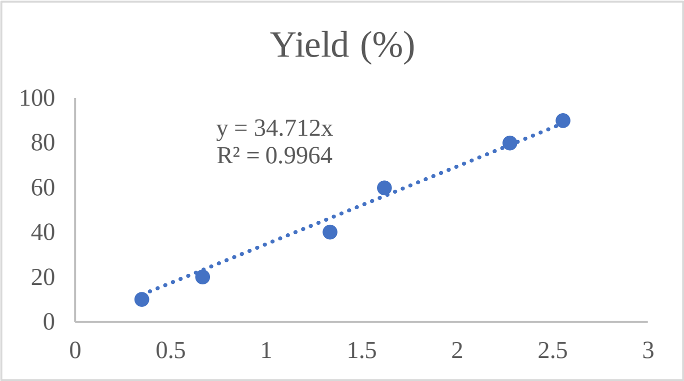

Enantiomeric ratio was determined by chiral HPLC using a Chiralpak IC column (4.6 x 250 mm, 5 μm), (HPLC: IF, 210 nm, n-hexane/isopropanol = 90:10, flow rate 1 mL/min, 25 °C, Retention Time: 6.637 min, 7.013 min).

HPLC trace of rac-**3q**:

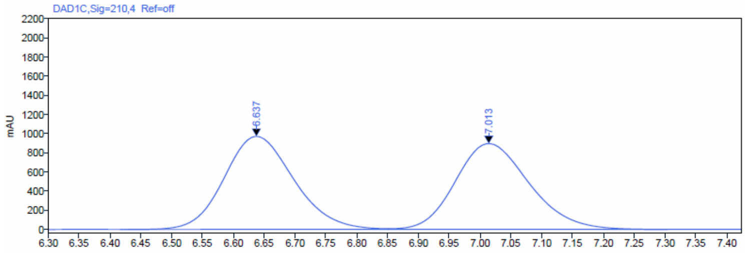

Signal: DAD1C,Sig=210,4 Ref=off

| RT [min] | Type | Width [min] | Area | Height | Area% | Name |
|---|---|---|---|---|---|---|
| 6.637 | BV | 0.44 | 7648.71 | 972.09 | 50.57 | |
| 7.013 | VBA | 0.45 | 7477.69 | 896.24 | 49.43 | |

HPLC trace of enantioenriched-**3q** obtained from the reaction catalyzed by SYE2:

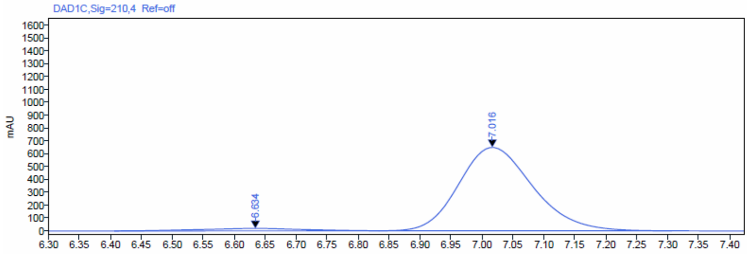

Signal: DAD1C,Sig=210,4 Ref=off

| RT [min] | Type | Width [min] | Area | Height | Area% | Name |
|---|---|---|---|---|---|---|
| 6.634 | BV | 0.43 | 227.19 | 19.44 | 4.03 | |
| 7.016 | VBA | 0.47 | 5409.60 | 649.00 | 95.97 | |

HPLC trace of enantioenriched-**3q** obtained from the reaction catalyzed by GluER-Y343A:

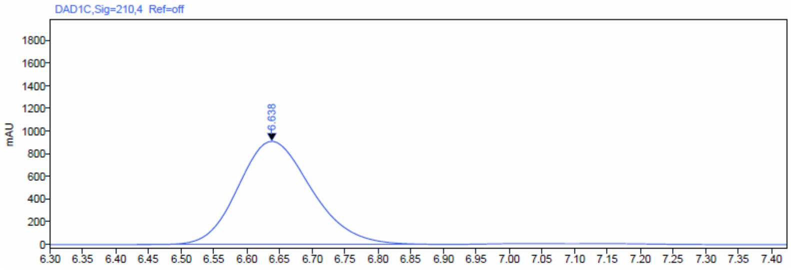

Signal: DAD1C,Sig=210,4 Ref=off

| RT [min] | Type | Width [min] | Area | Height | Area% | Name |
|---|---|---|---|---|---|---|
| 6.638 | BV | 0.47 | 7086.70 | 909.78 | 100.00 | |

Rac-**3q** is a colorless oil, and was synthesized through a sequence of a Knoevenagel condensation followed by a Pd/C catalyzed hydrogenation.

$^{1}$H NMR (600 MHz, $CDCl_3$) δ 7.29 (t, *J* = 7.6 Hz, 2H), 7.22 – 7.13 (m, 3H), 3.73 (s, 3H), 3.62 (s, 3H), 3.21 (dd, *J* = 9.2, 5.9 Hz, 1H), 2.74 – 2.68 (m, 1H), 2.23 (ddd, *J* = 14.5, 9.2, 5.6 Hz, 1H), 2.16 (ddd, *J* = 14.0, 9.7, 5.9 Hz, 1H), 1.28 (d, *J* = 7.0 Hz, 3H).

$^{13}$C NMR (151 MHz, $CDCl_3$) δ 169.91, 169.83, 145.24, 128.60, 127.11, 126.53, 52.47, 52.45, 50.02, 37.91, 37.03, 22.49.

HRMS (ESI, m/z) calcd for $C_{14}H_{19}O_4$ $[M+H]^+$: 251.1283, found: 251.1281.

HRMS spectrum of **3q**:

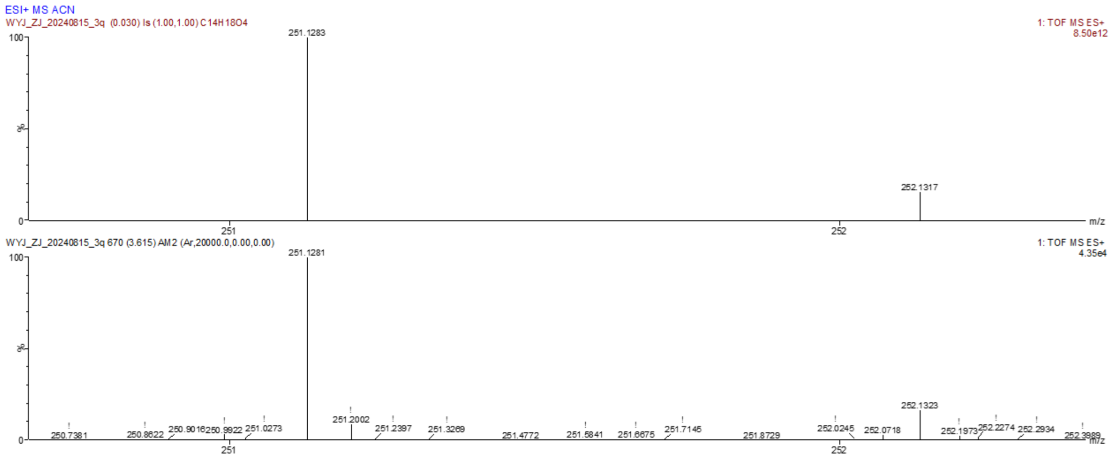

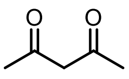

**1c**, n.d.

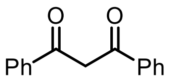

**1d**, n.r.

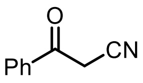

**1e**, n.d.

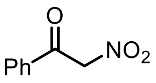

**1f**, n.d.

$CH_3NO_2$

**1g**, n.r.

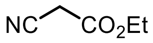

**1h**, n.r.

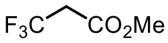

**1i**, n.r.

**Supplementary Fig. 34 |Unsuccessful substrates.** n.d. Not detected; n.r. No reaction.

### 12. NMR Spectra of Substrates and Products

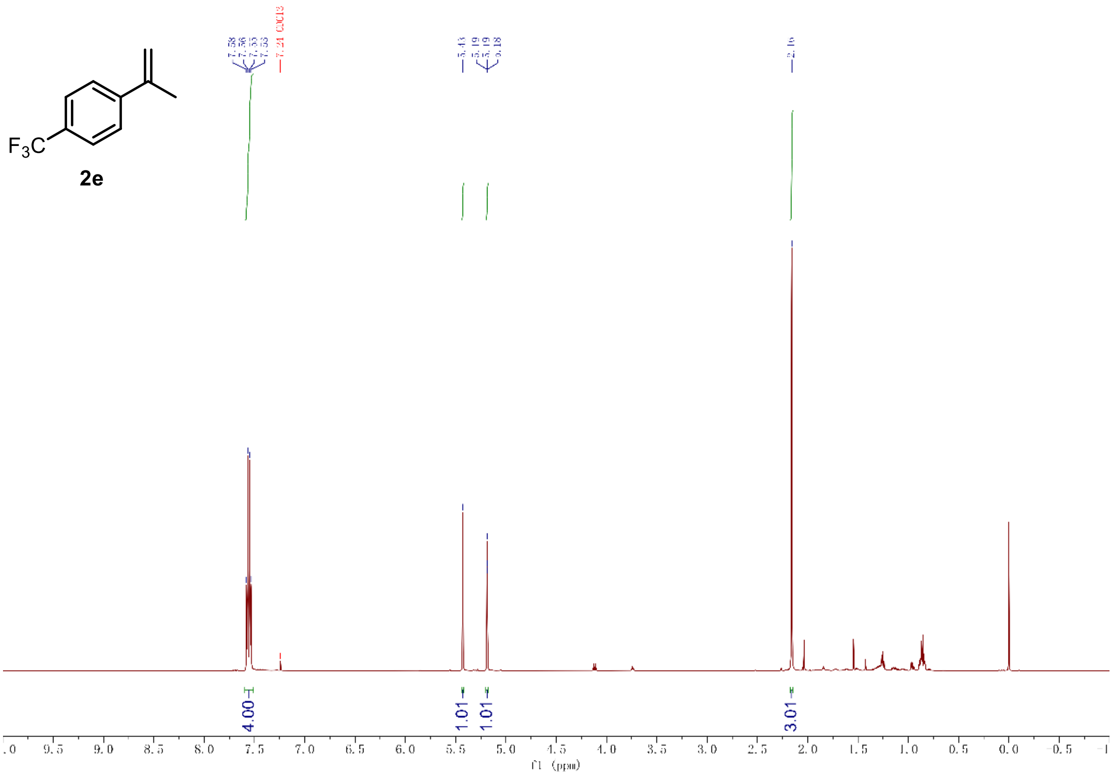

[1]H-NMR of compound **2e** (600 MHz, $CDCl_3$)

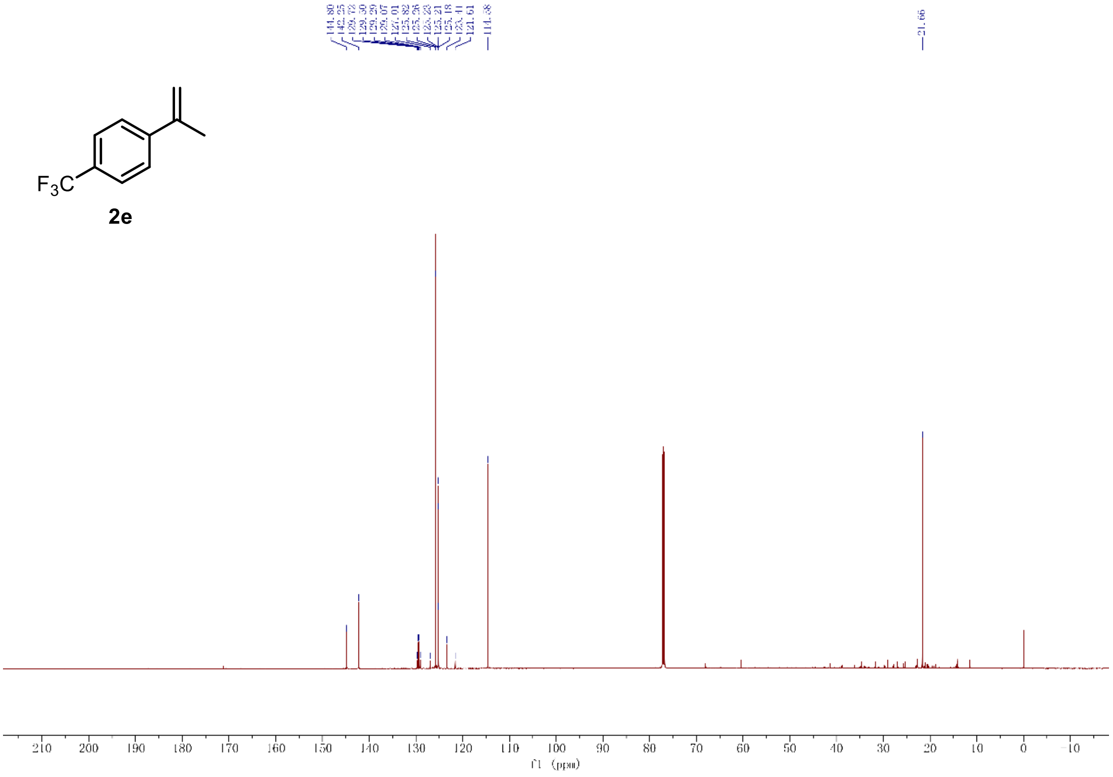

[13]C-NMR of compound **2e** (151 MHz, $CDCl_3$)

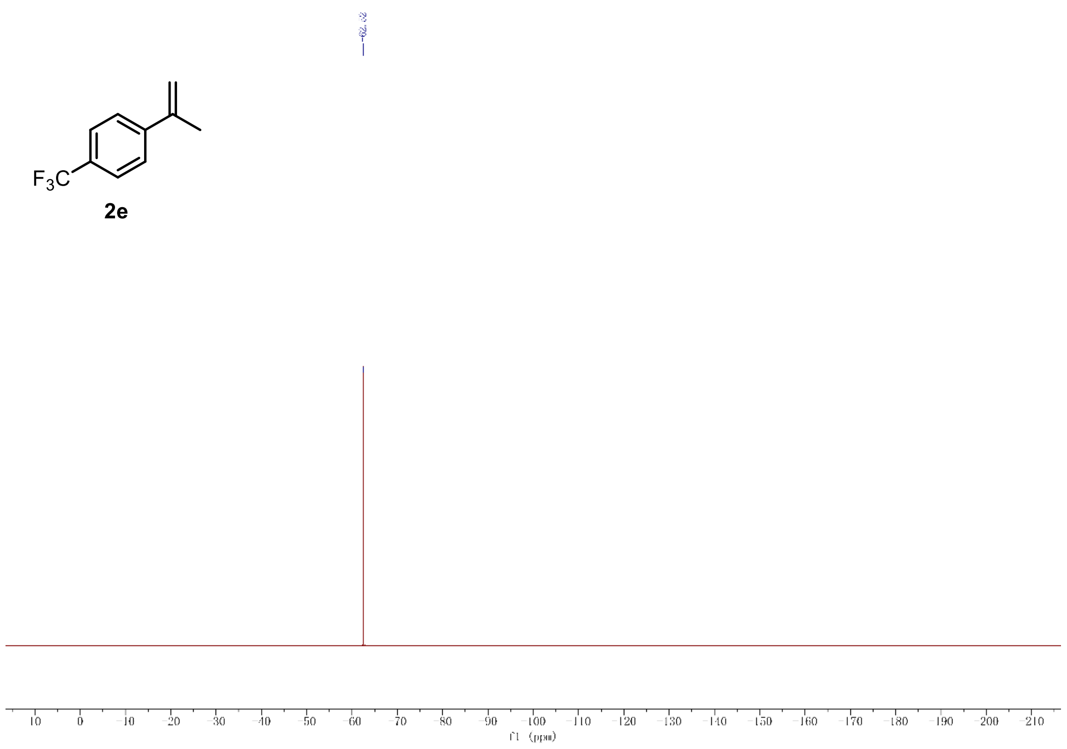

$^{19}F$-NMR of compound **2e** (565 MHz, $CDCl_3$)

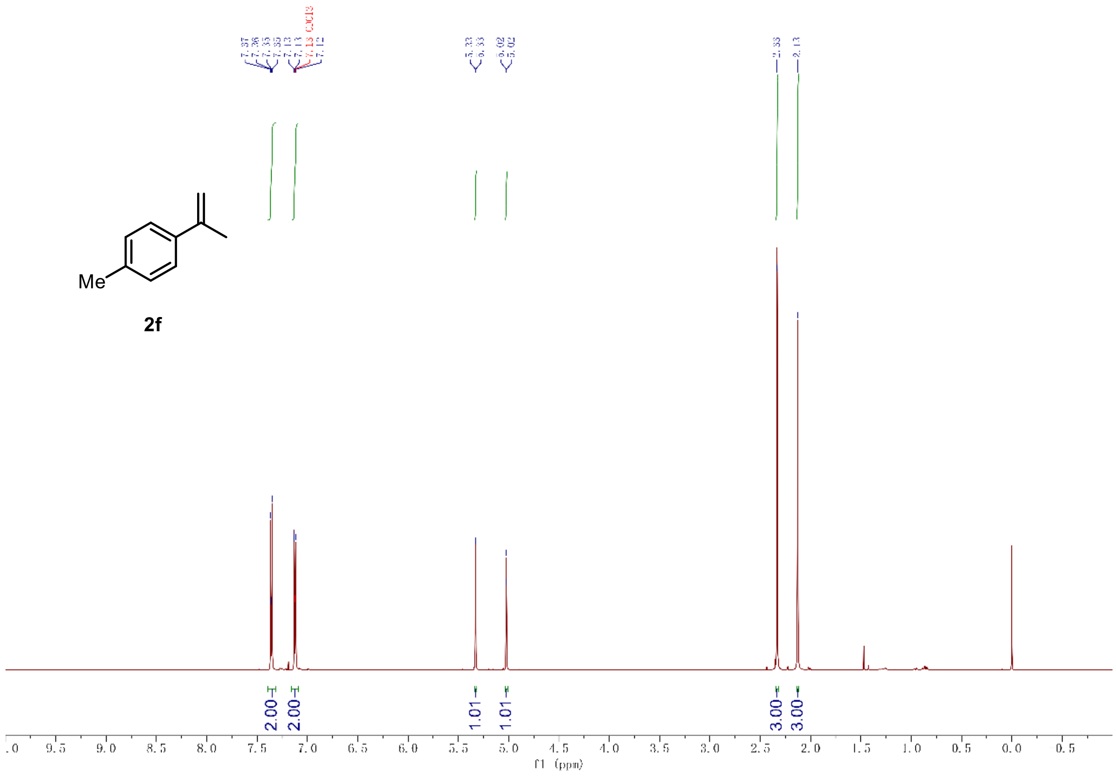

[1]H-NMR of compound **2f** (600 MHz, $CDCl_3$)

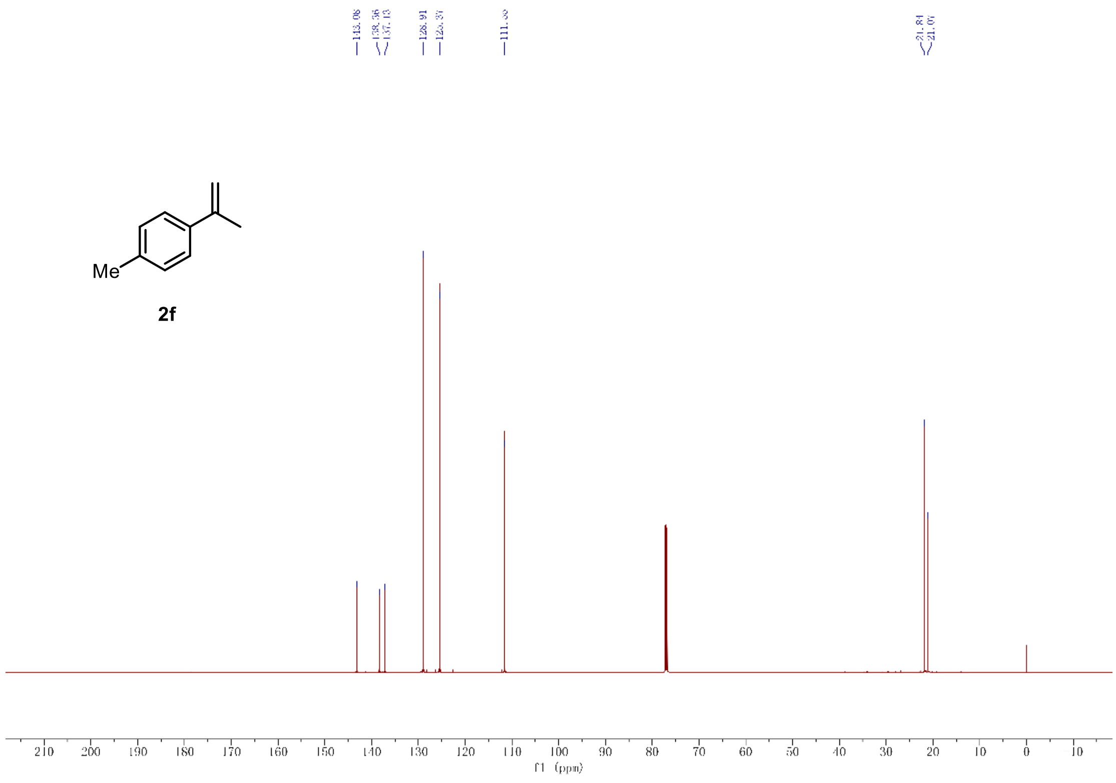

[13]C-NMR of compound **2f** (151 MHz, $CDCl_3$)

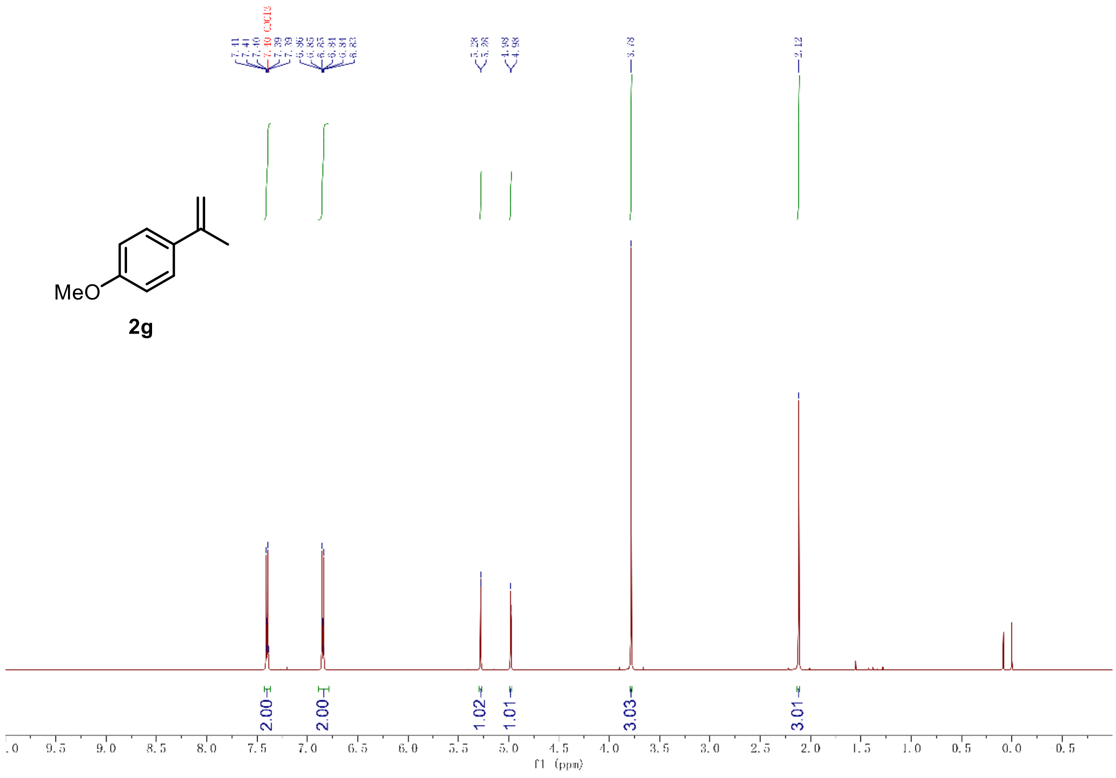

[1]H-NMR of compound **2g** (600 MHz, $CDCl_3$)

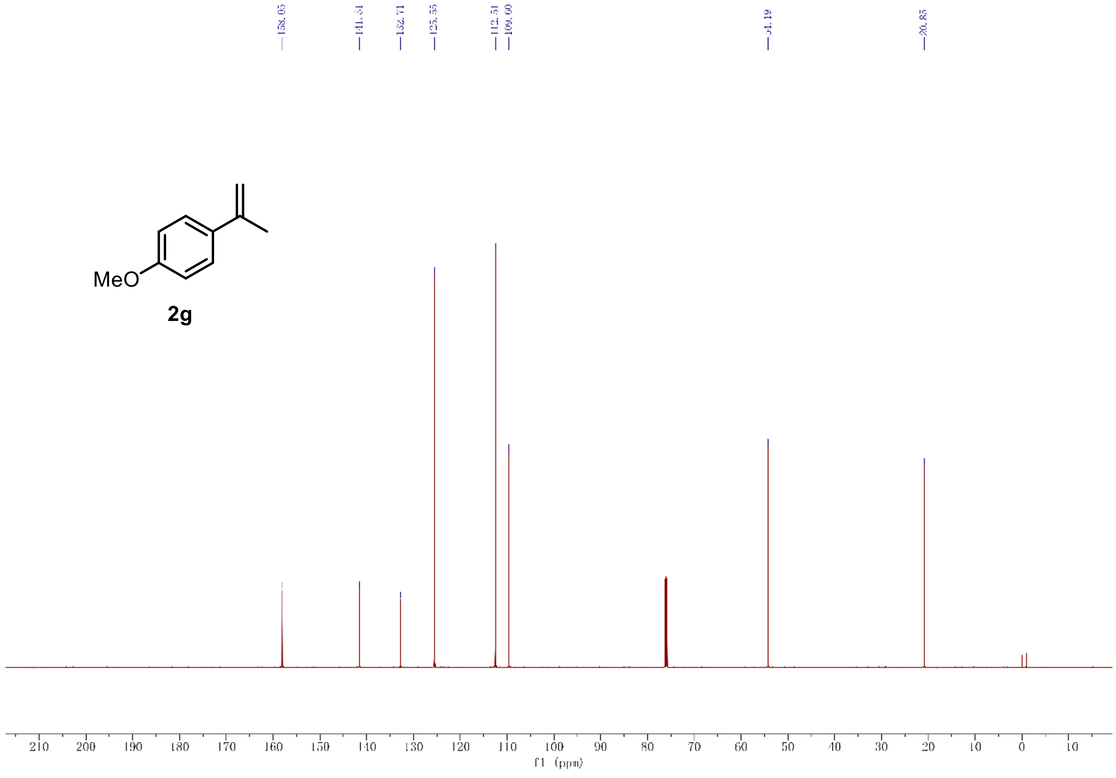

[13]C-NMR of compound **2g** (151 MHz, $CDCl_3$)

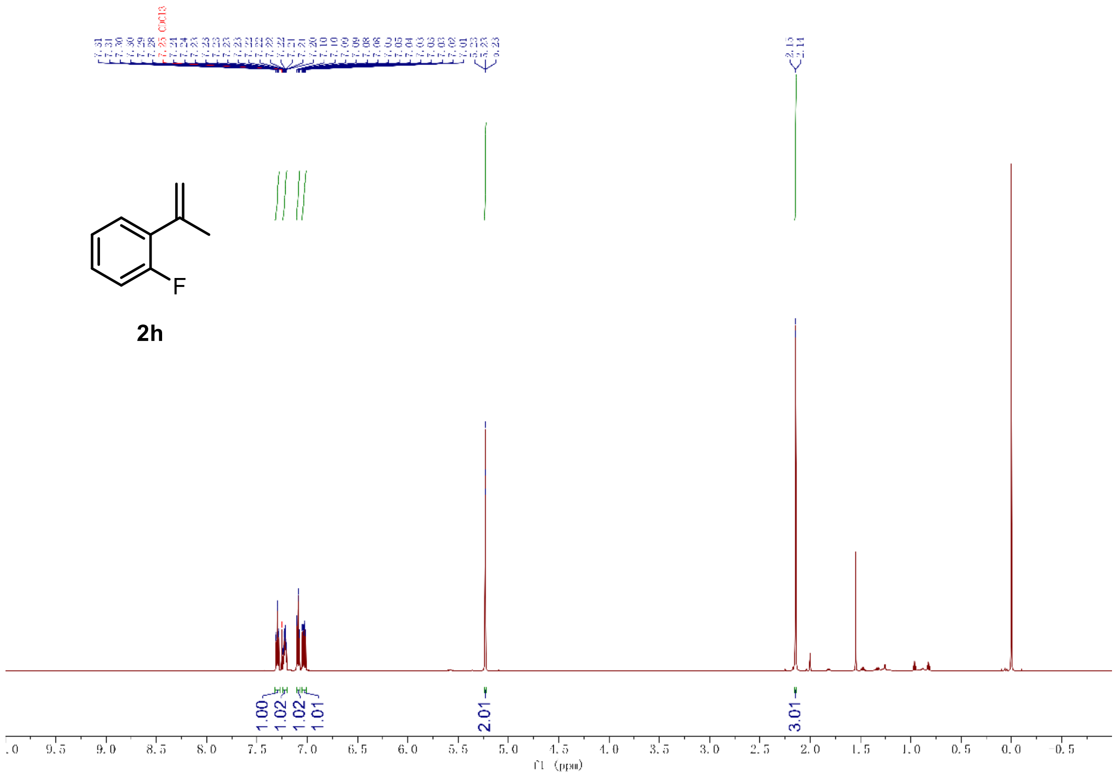

$^1$H-NMR of compound **2h** (600 MHz, $CDCl_3$)

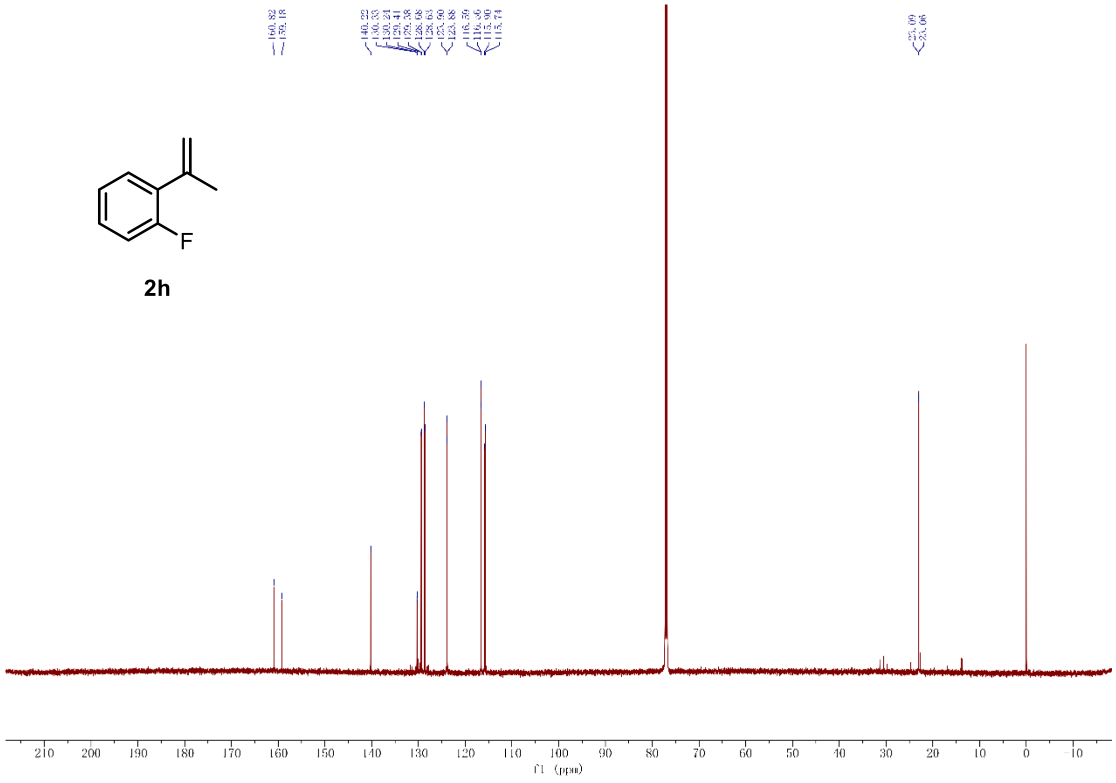

$^{13}$C-NMR of compound **2h** (151 MHz, $CDCl_3$)

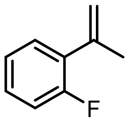

**2h**

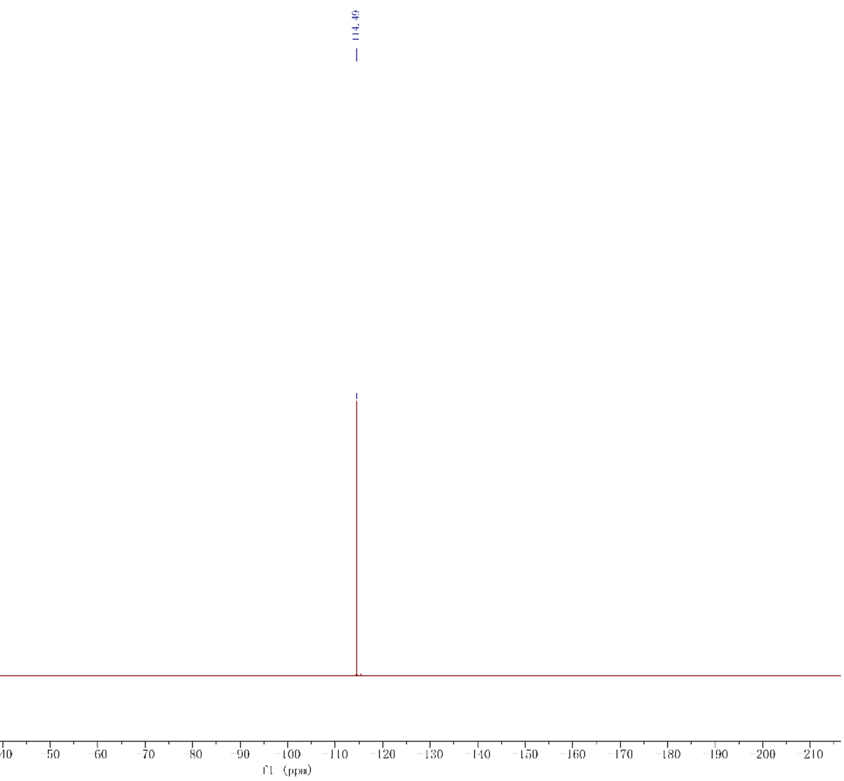

[19]F-NMR of compound **2h** (565 MHz, $CDCl_3$)

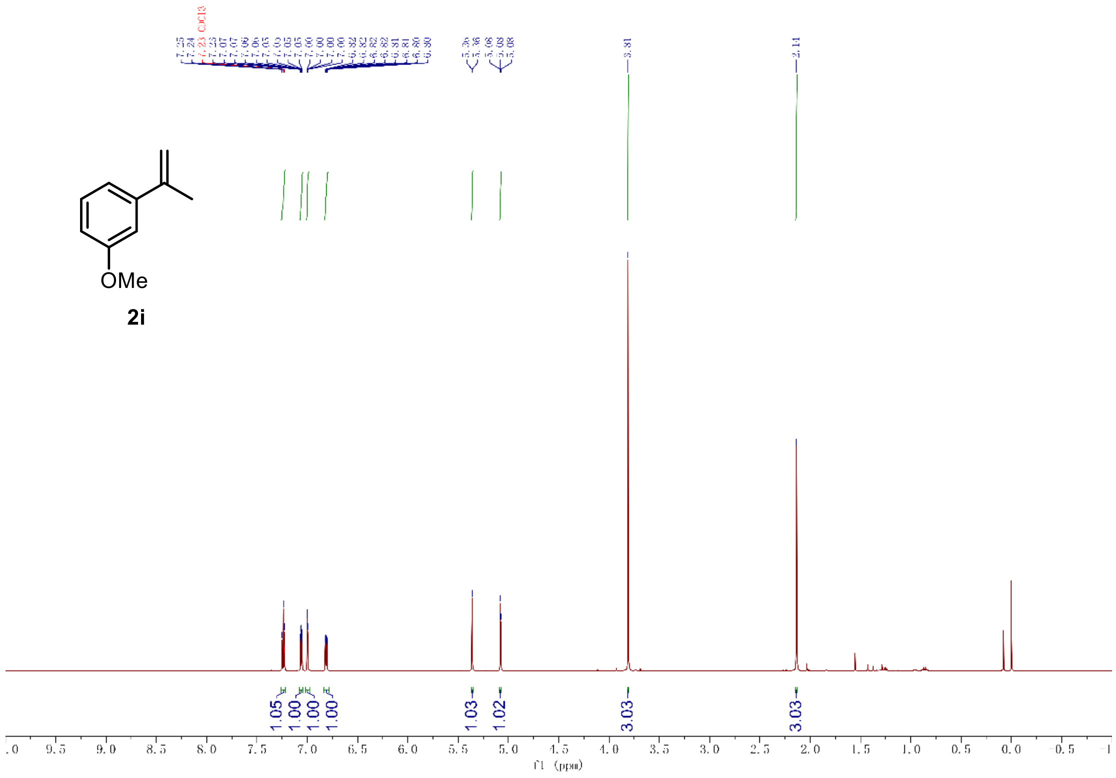

[1]H-NMR of compound **2i** (600 MHz, $CDCl_3$)

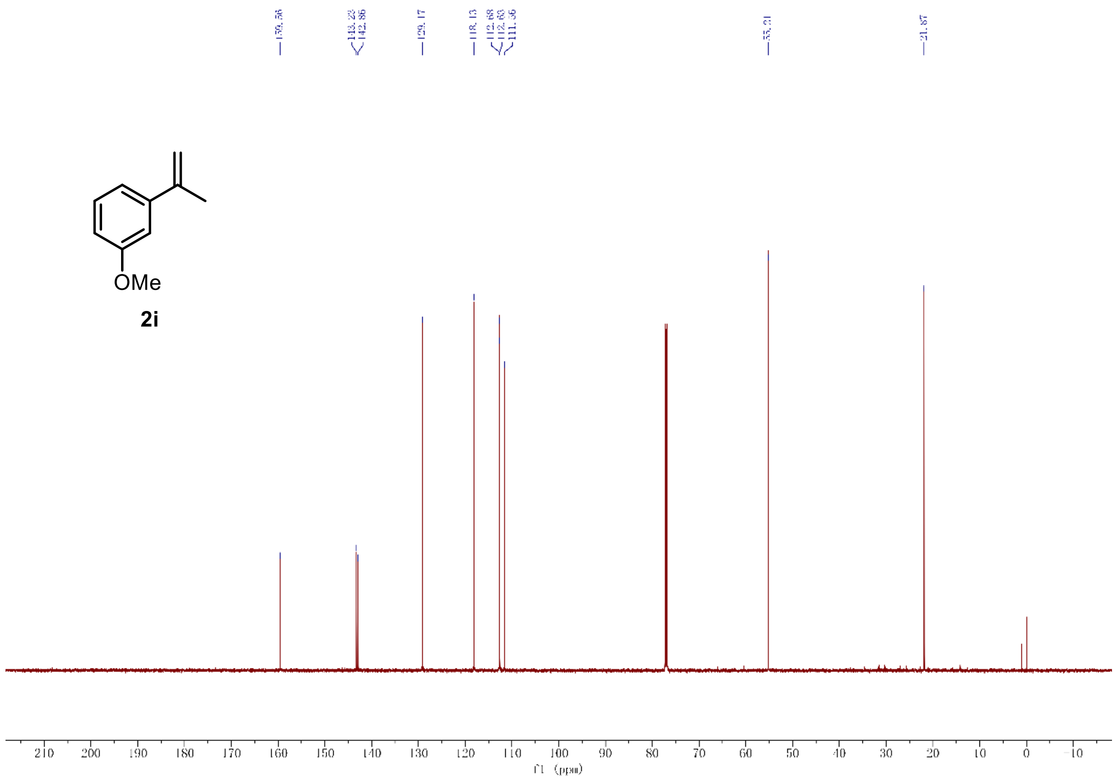

[13]C-NMR of compound **2i** (151 MHz, $CDCl_3$)

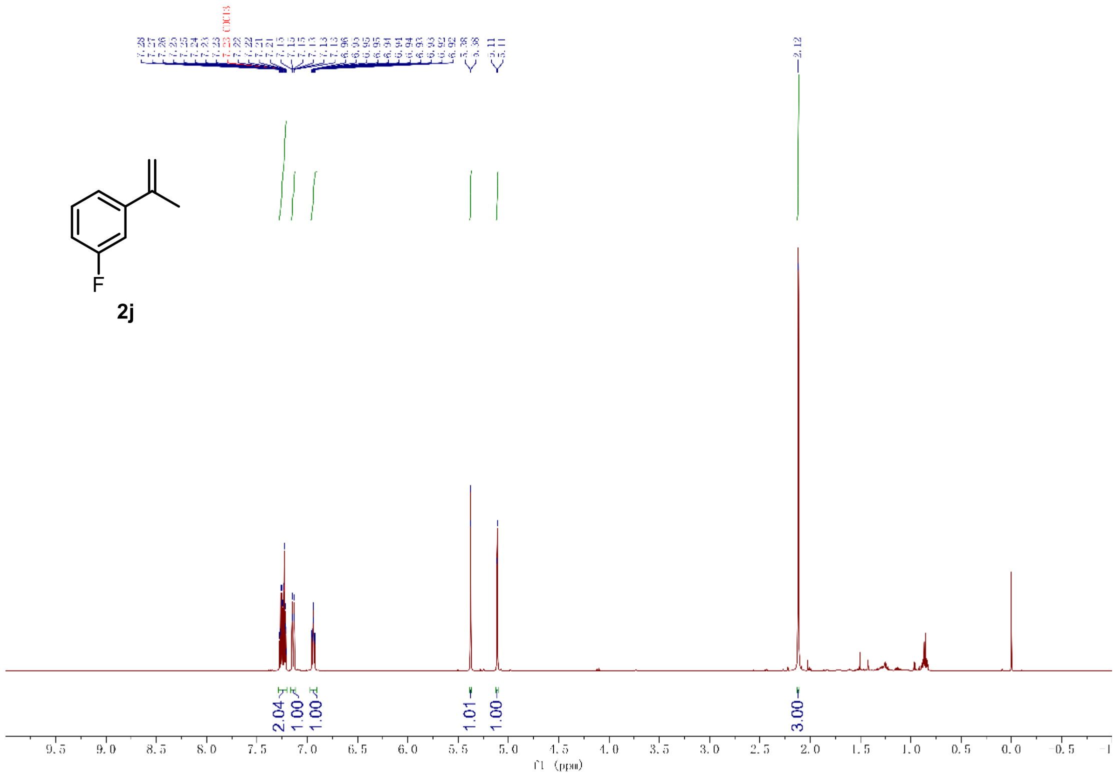

[1]H-NMR of compound **2j** (600 MHz, $CDCl_3$)

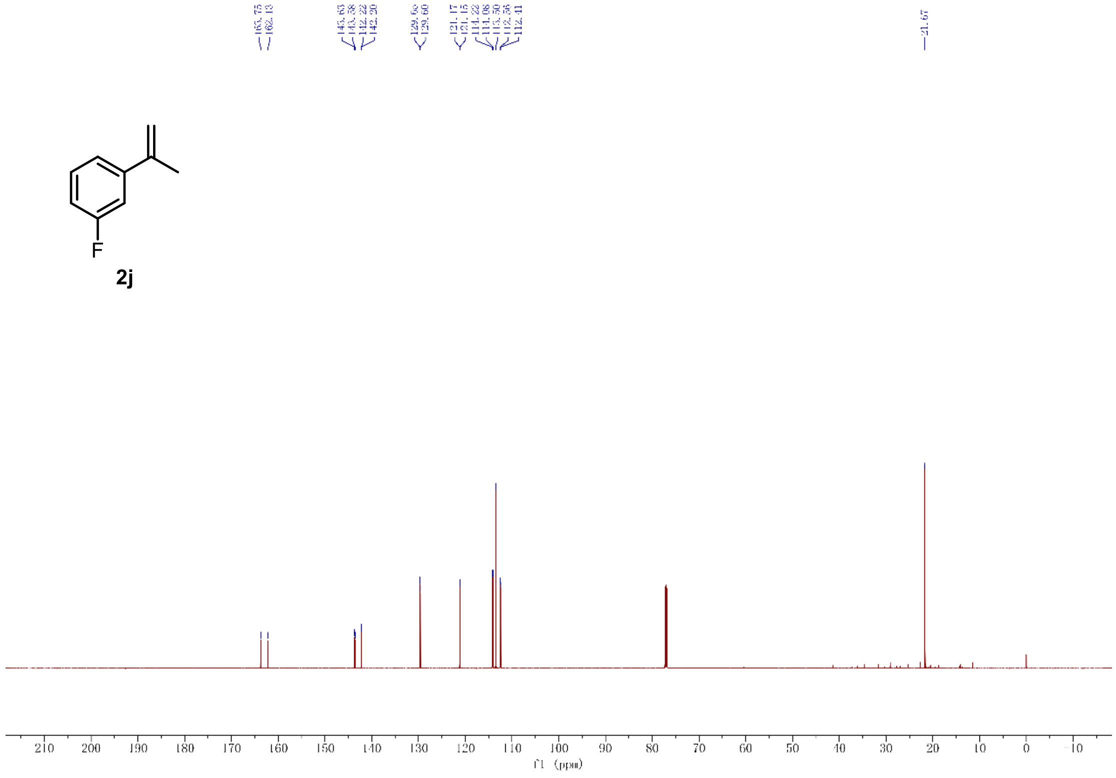

[13]C-NMR of compound **2j** (151 MHz, $CDCl_3$)

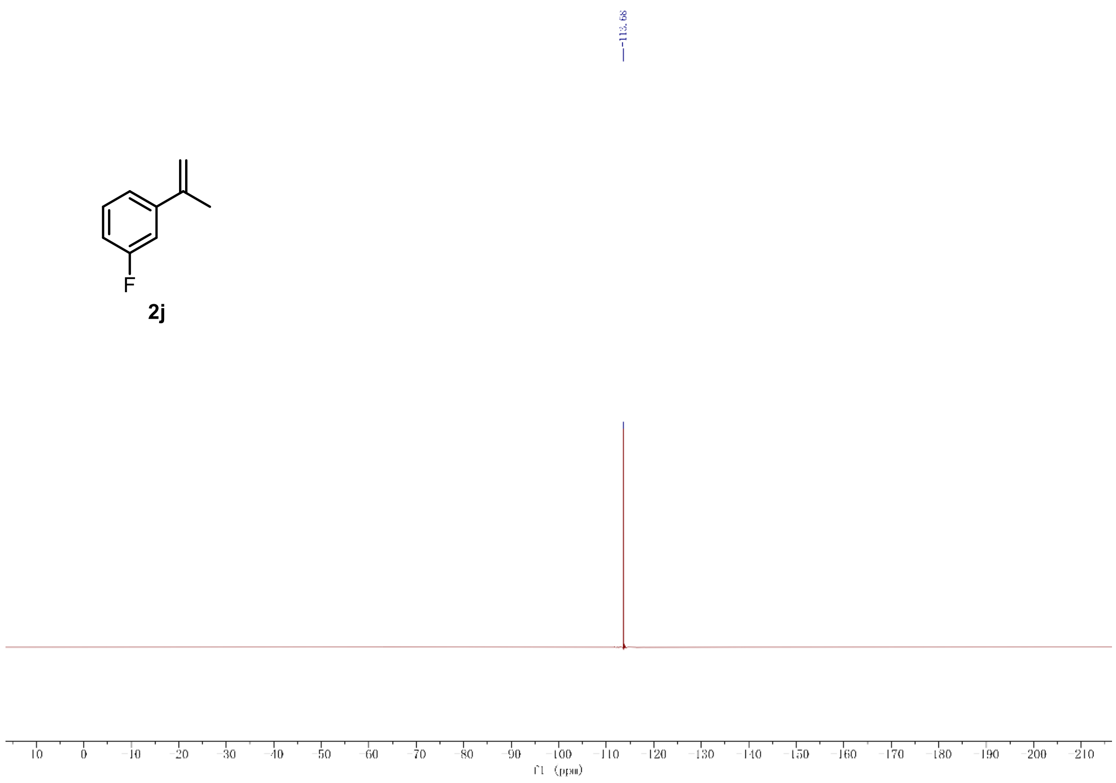

$^{19}$F-NMR of compound **2j** (565 MHz, $CDCl_3$)

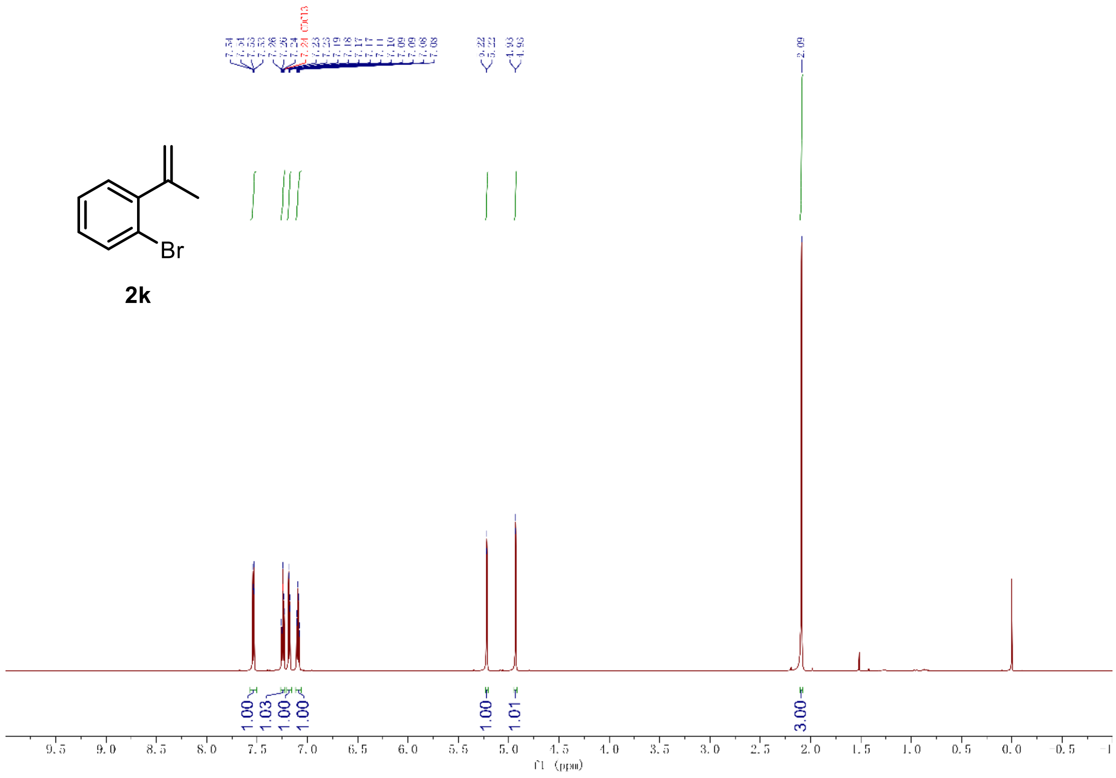

$^{1}$H-NMR of compound **2k** (600 MHz, $CDCl_3$)

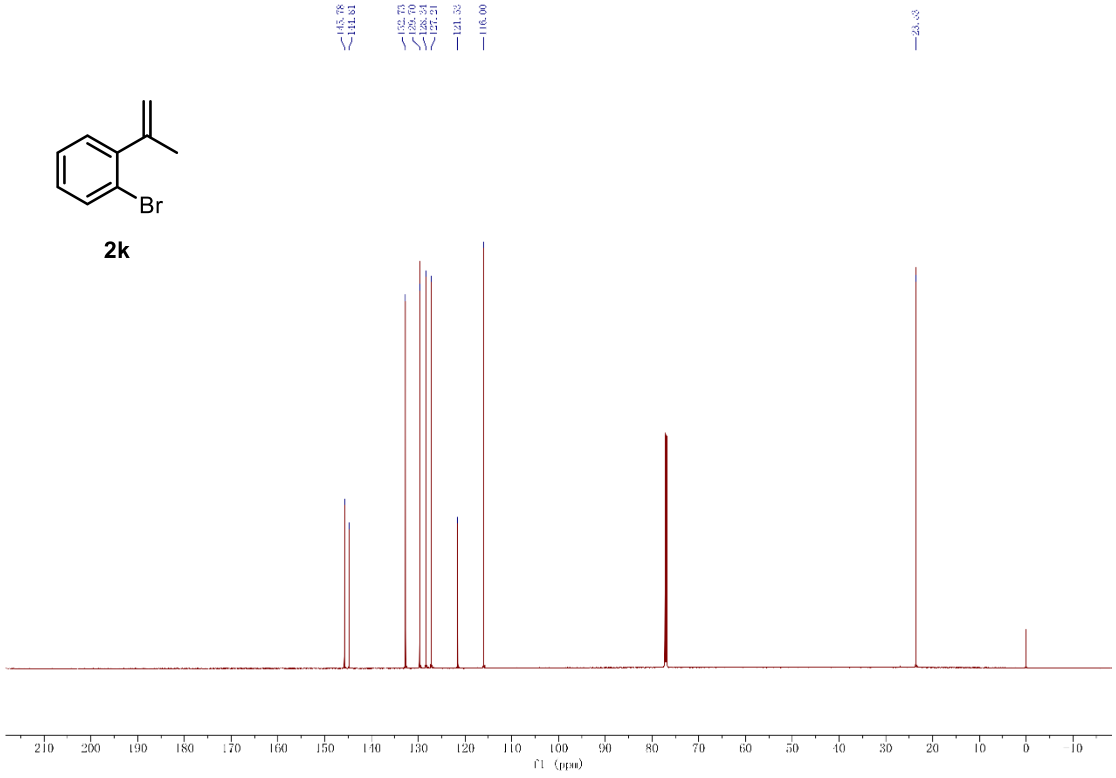

$^{13}$C-NMR of compound **2k** (151 MHz, $CDCl_3$)

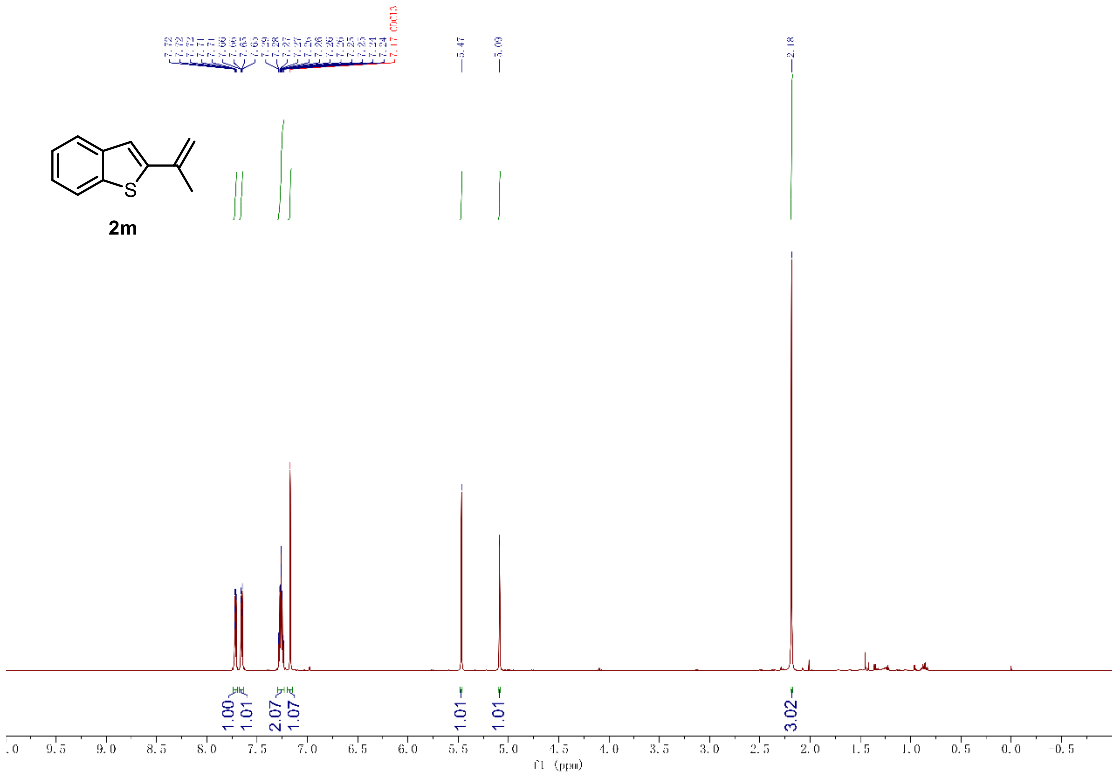

[1]H-NMR of compound **2m** (600 MHz, $CDCl_3$)

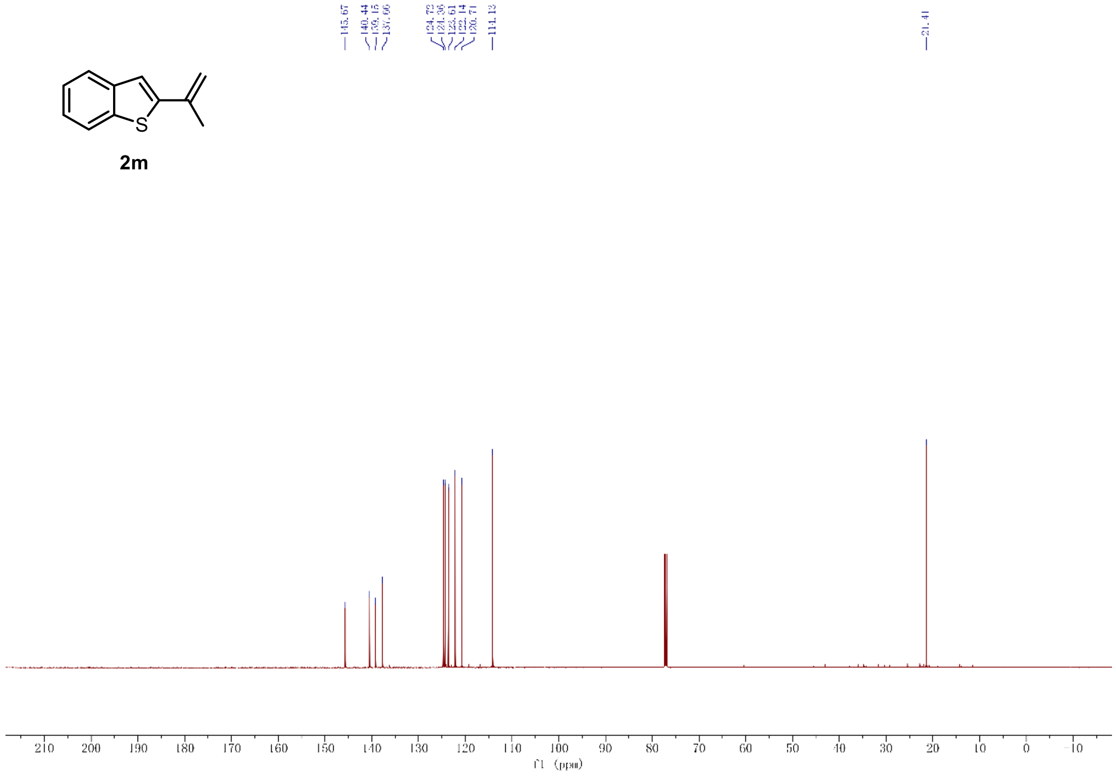

[13]C-NMR of compound **2m** (151 MHz, $CDCl_3$)

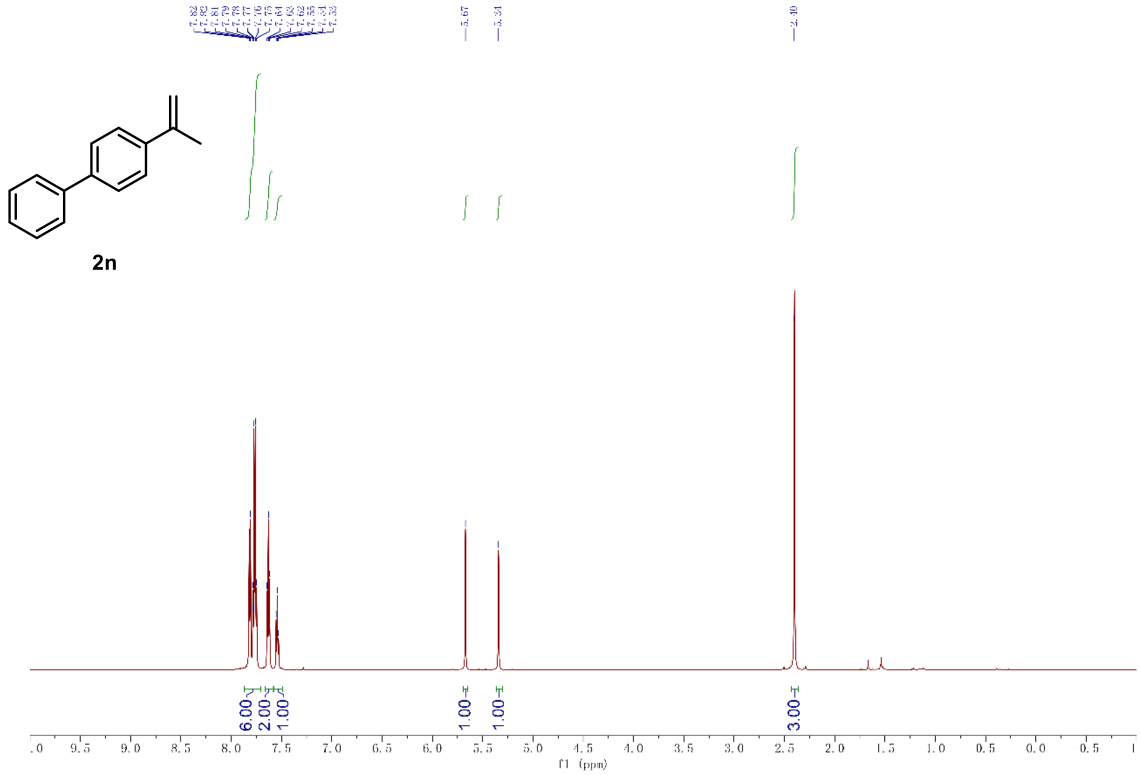

$^{1}$H-NMR of compound **2n** (600 MHz, $CDCl_3$)

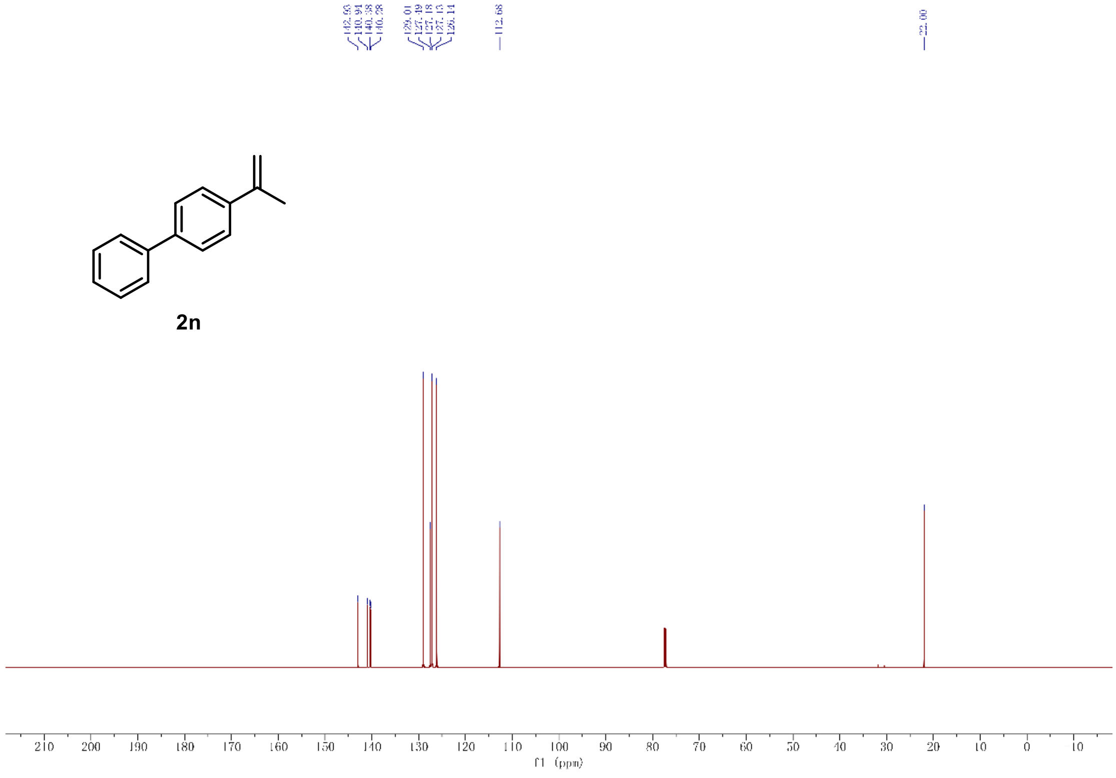

$^{13}$C-NMR of compound **2n** (151 MHz, $CDCl_3$)

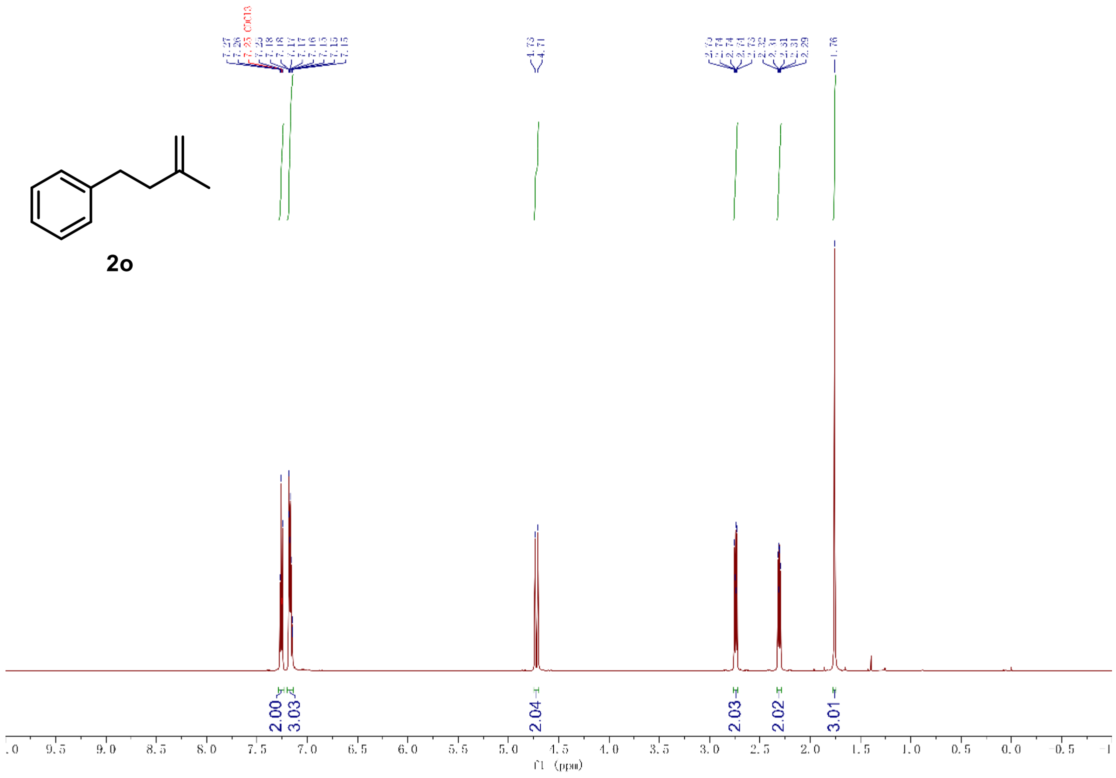

$^{1}$H-NMR of compound **2o** (600 MHz, $CDCl_3$)

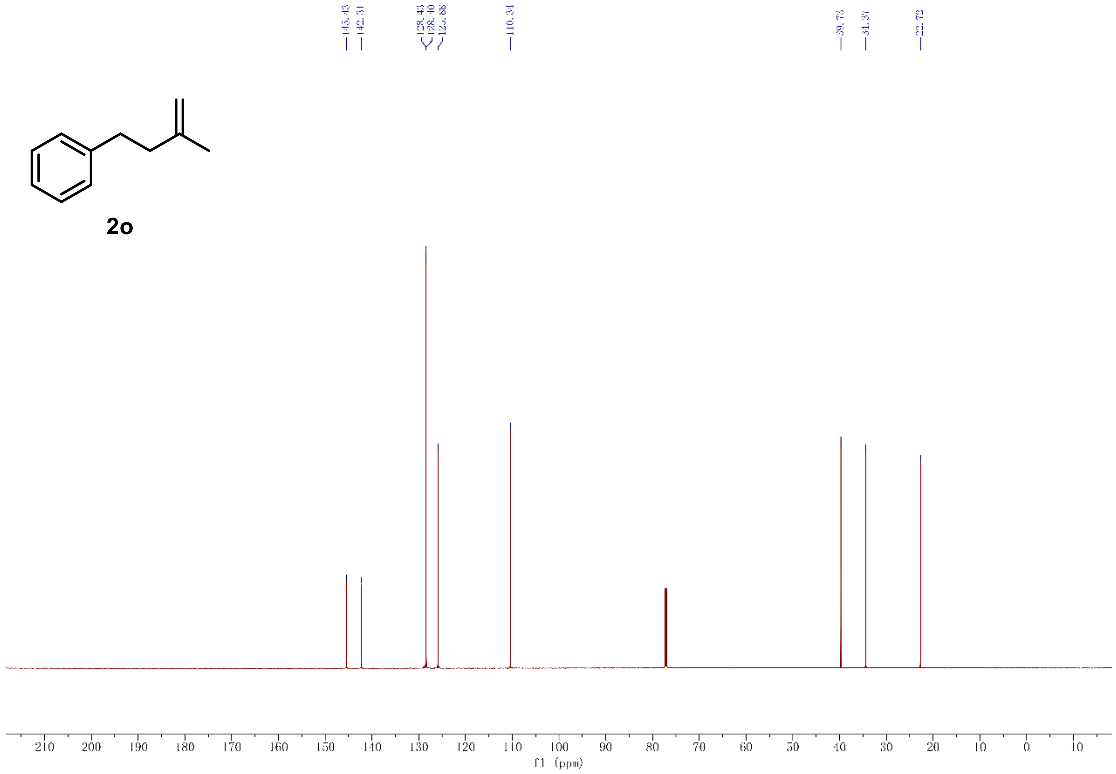

$^{13}$C-NMR of compound **2o** (151 MHz, $CDCl_3$)

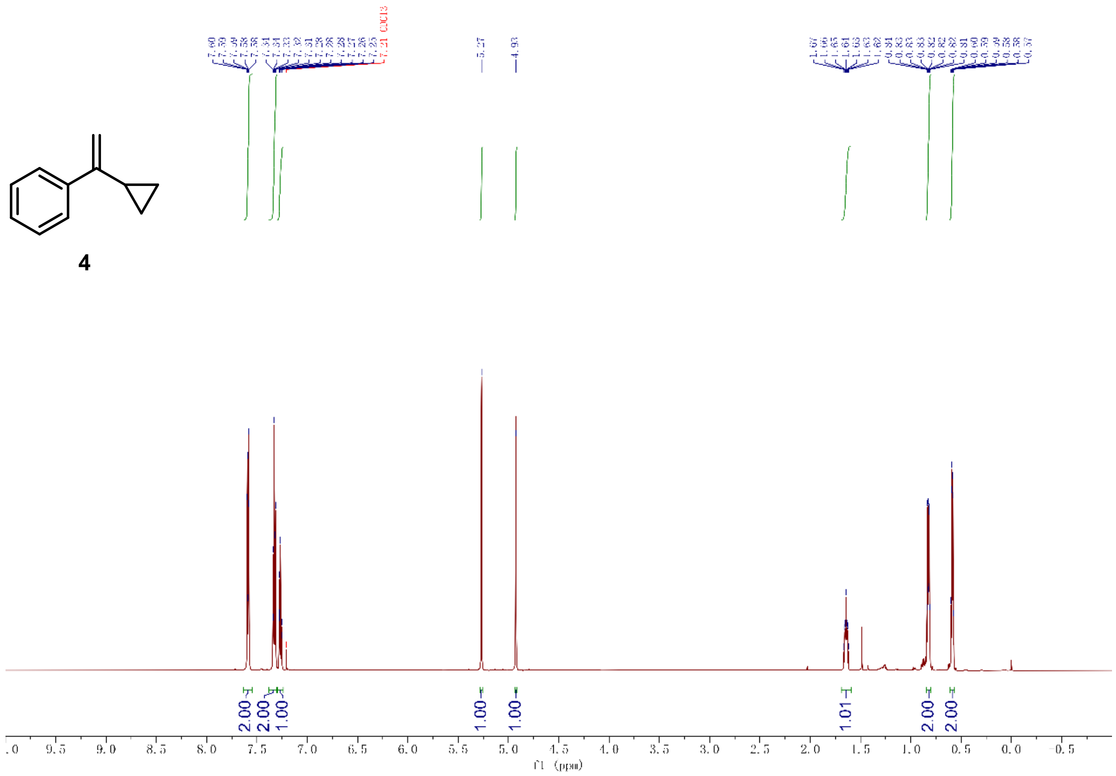

[1]H-NMR of compound **4** (600 MHz, $CDCl_3$)

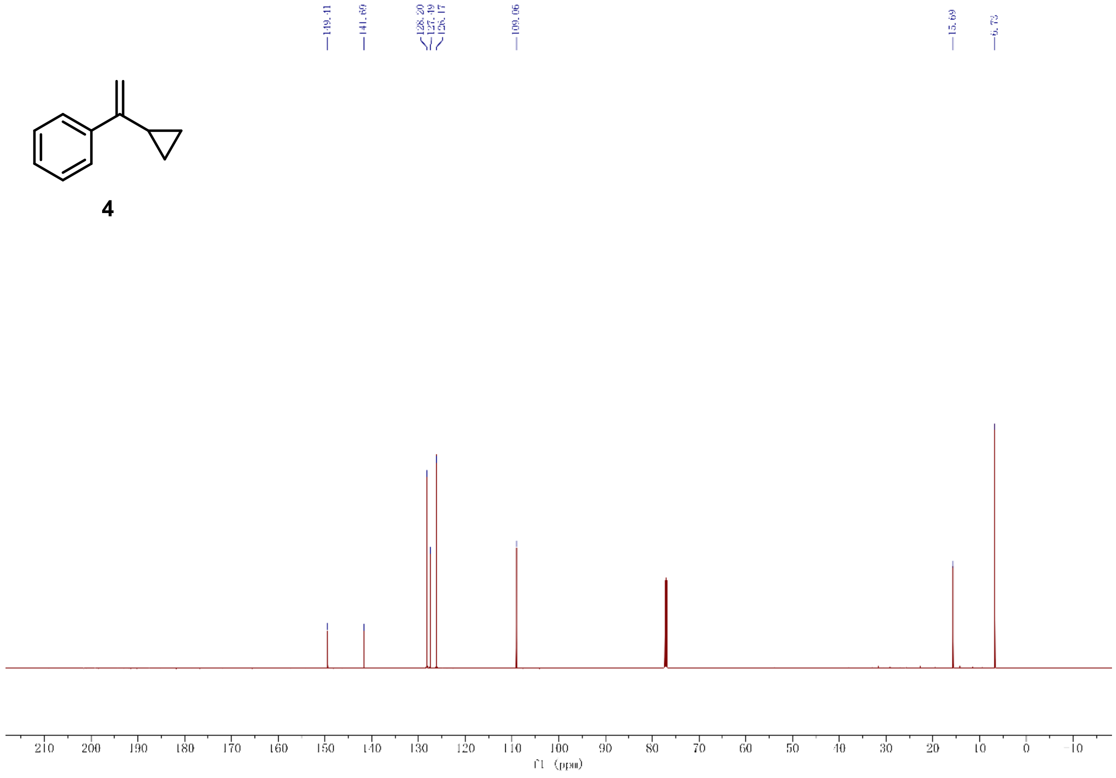

[13]C-NMR of compound **4** (151 MHz, $CDCl_3$)

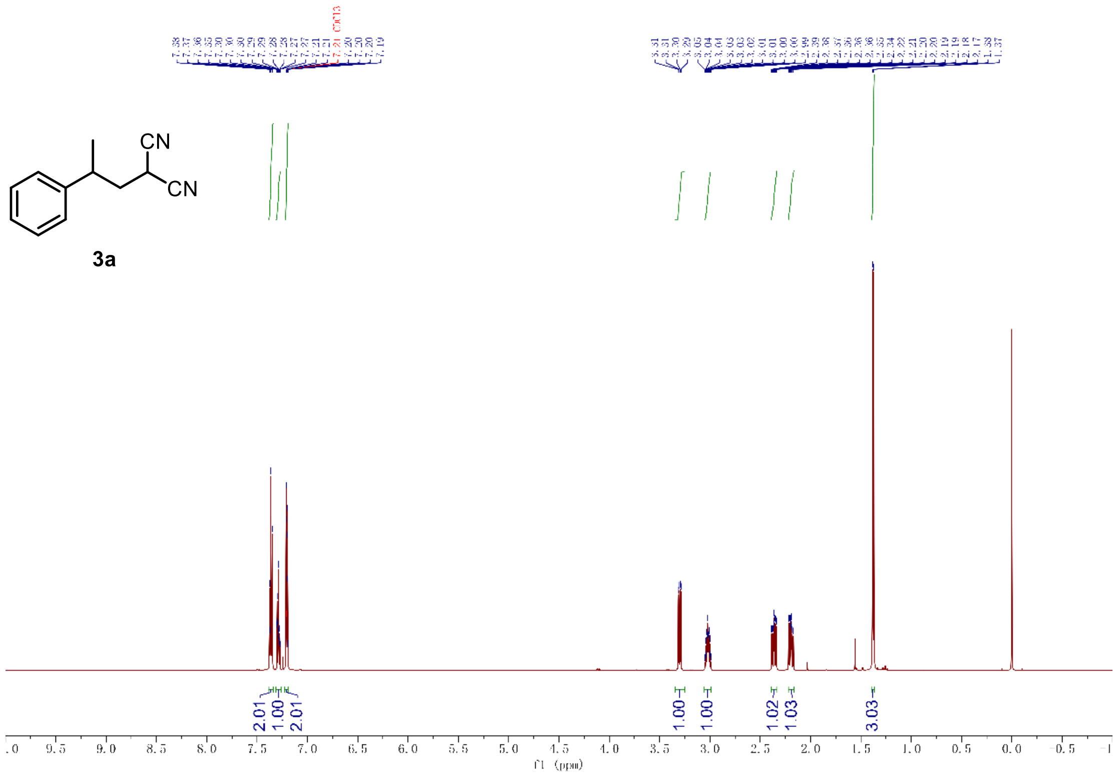

[1]H-NMR of compound **3a** (600 MHz, $CDCl_3$)

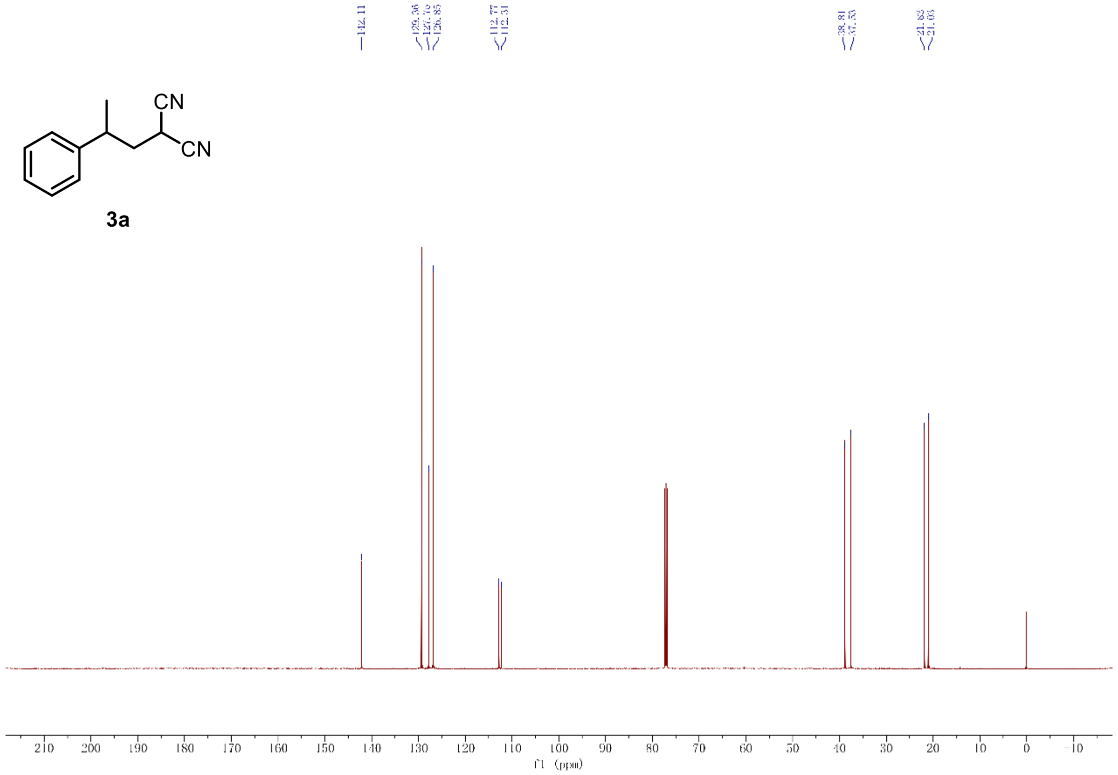

[13]C-NMR of compound **3a** (151 MHz, $CDCl_3$)

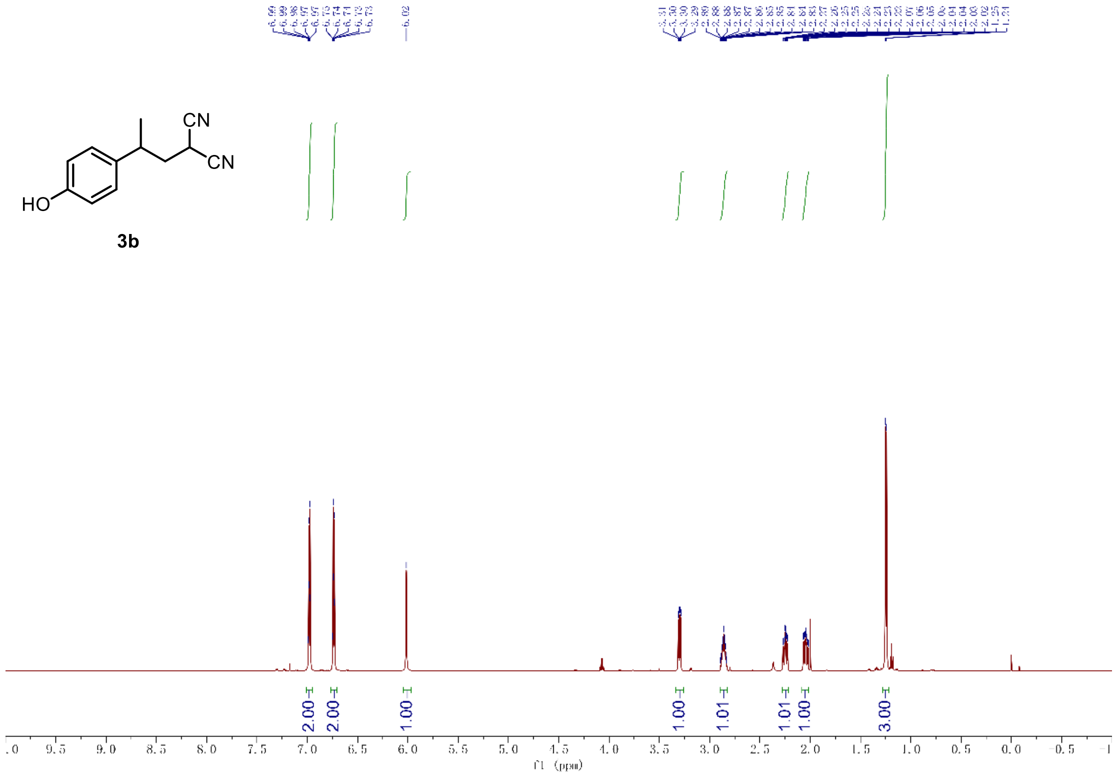

$^{1}$H-NMR of compound **3b** (600 MHz, $CDCl_3$)

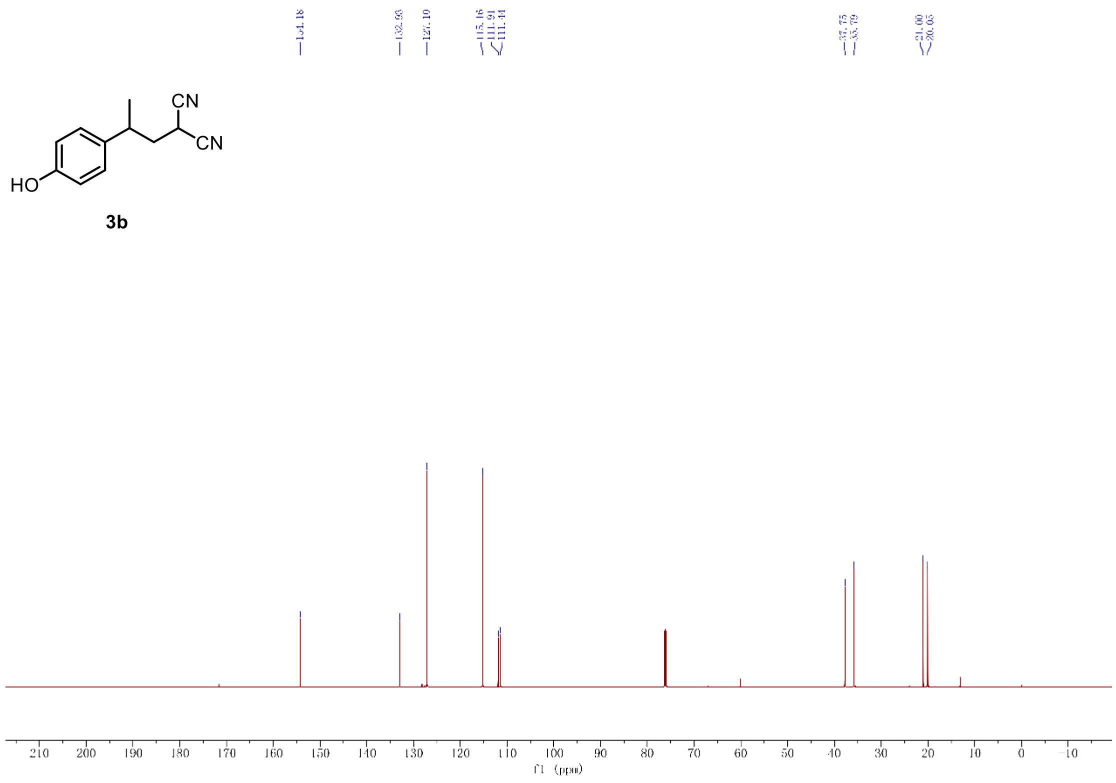

$^{13}$C-NMR of compound **3b** (151 MHz, $CDCl_3$)

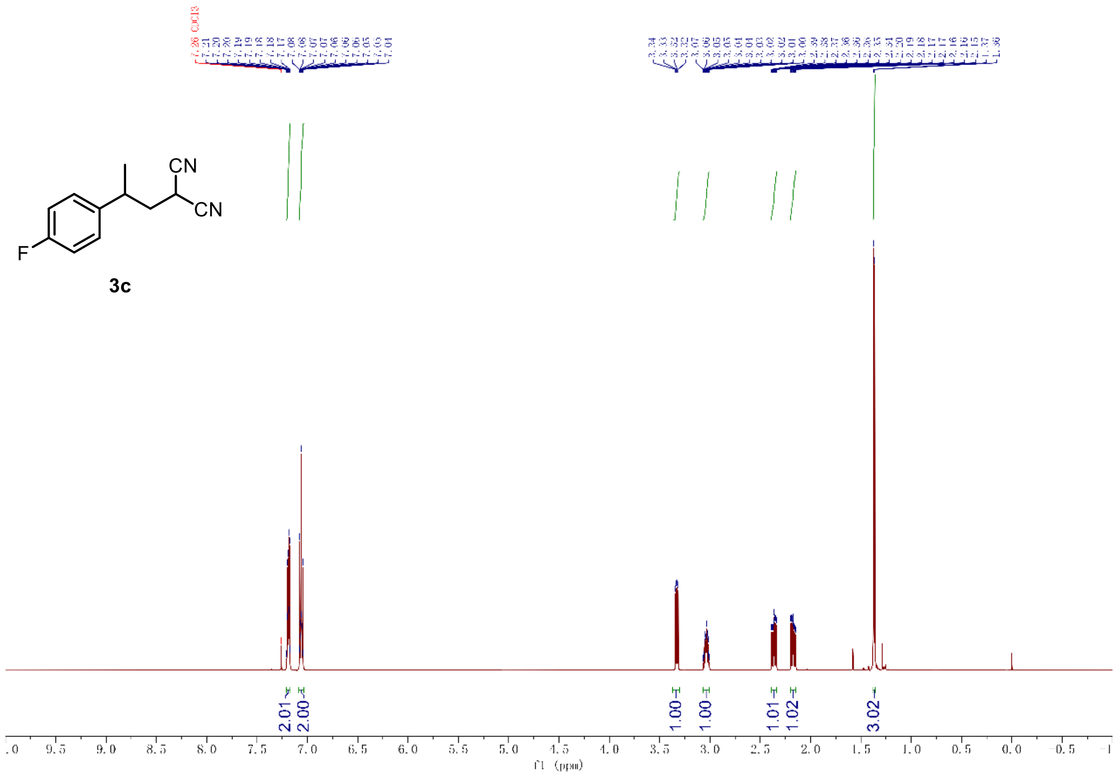

$^{1}$H-NMR of compound **3c** (600 MHz, $CDCl_3$)

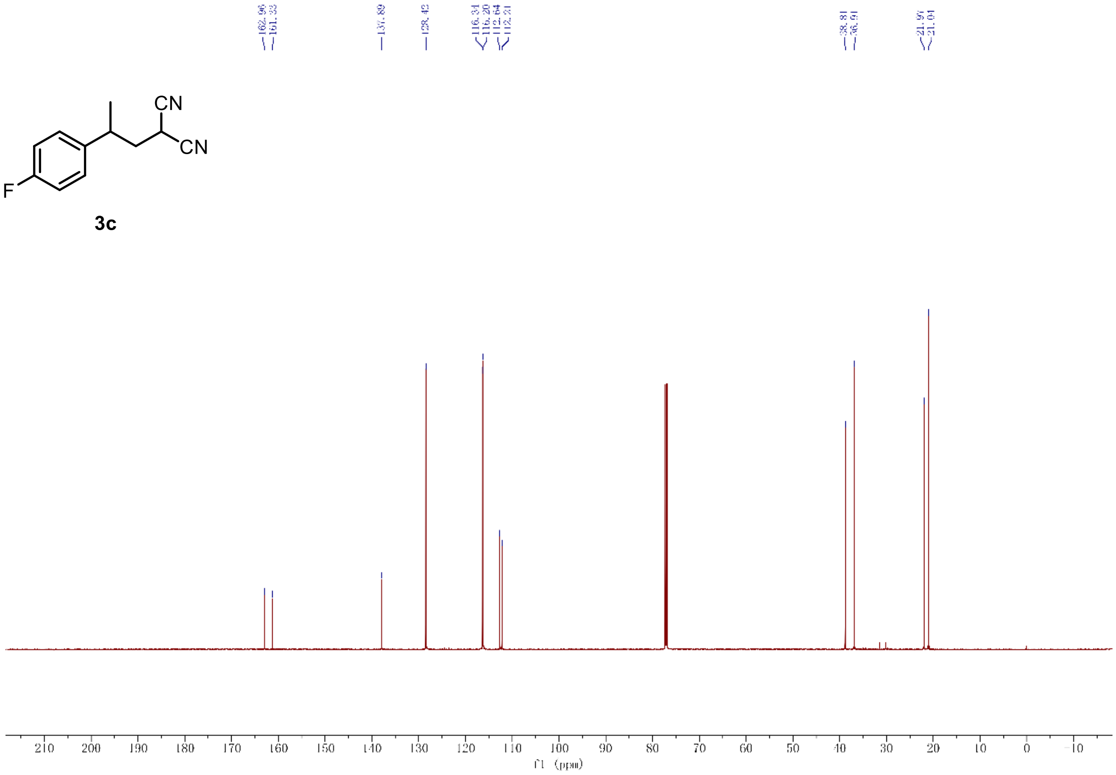

$^{13}$C-NMR of compound **3c** (151 MHz, $CDCl_3$)

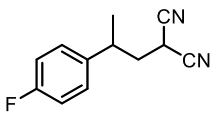

**3c**

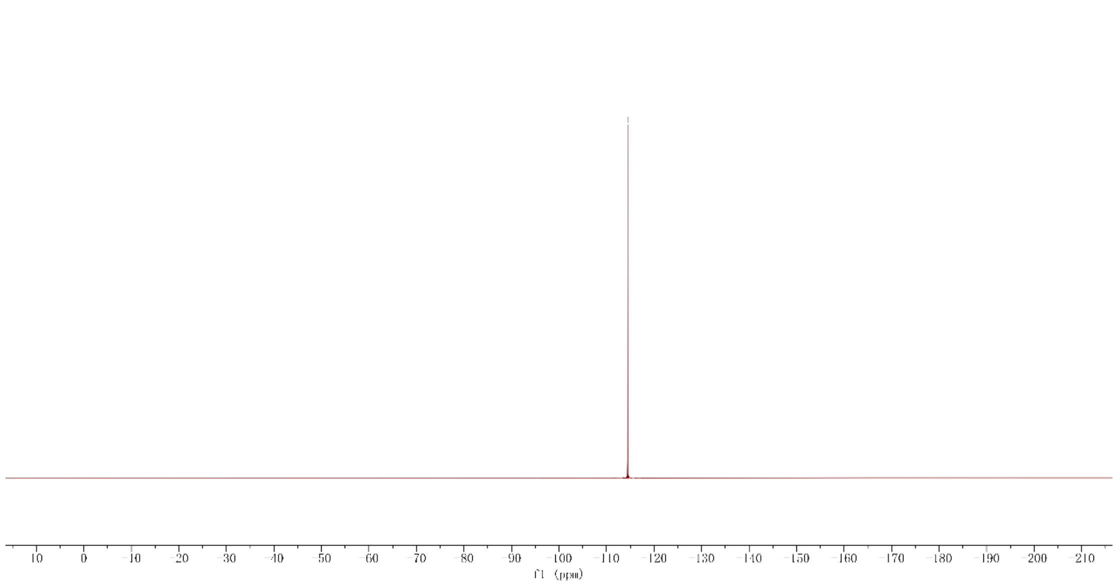

[19]F-NMR of compound **3c** (565 MHz, $CDCl_3$)

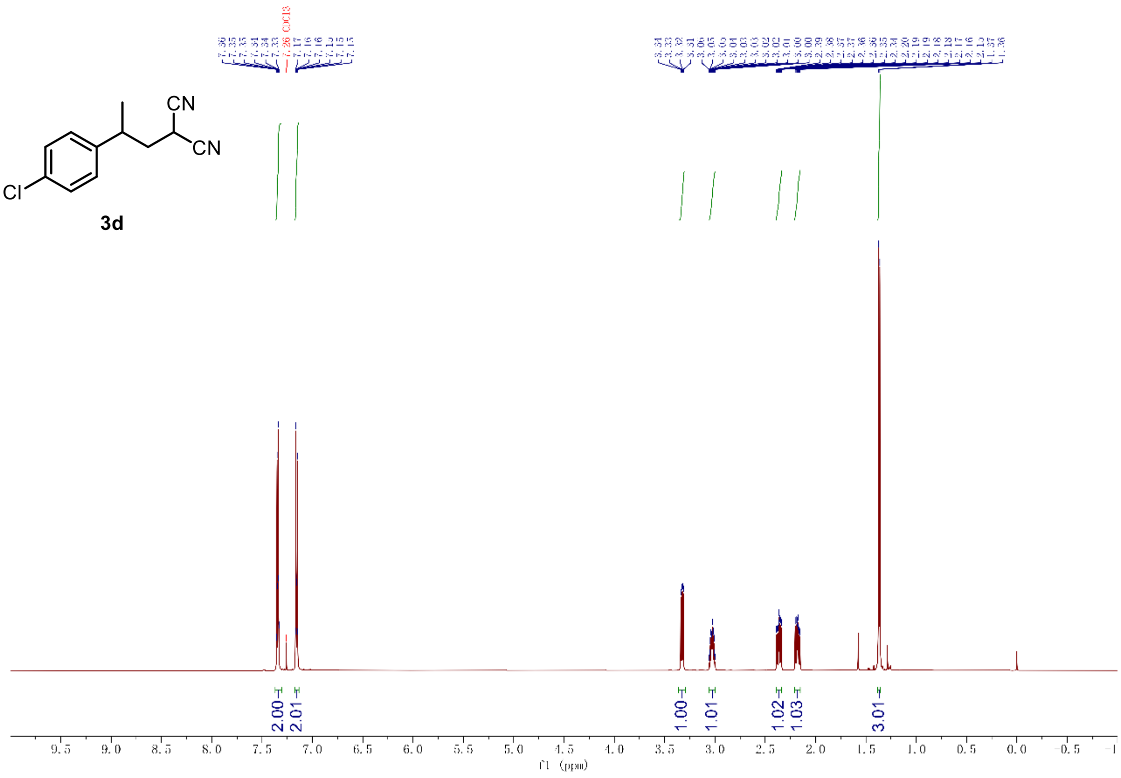

$^{1}$H-NMR of compound **3d** (600 MHz, $CDCl_3$)

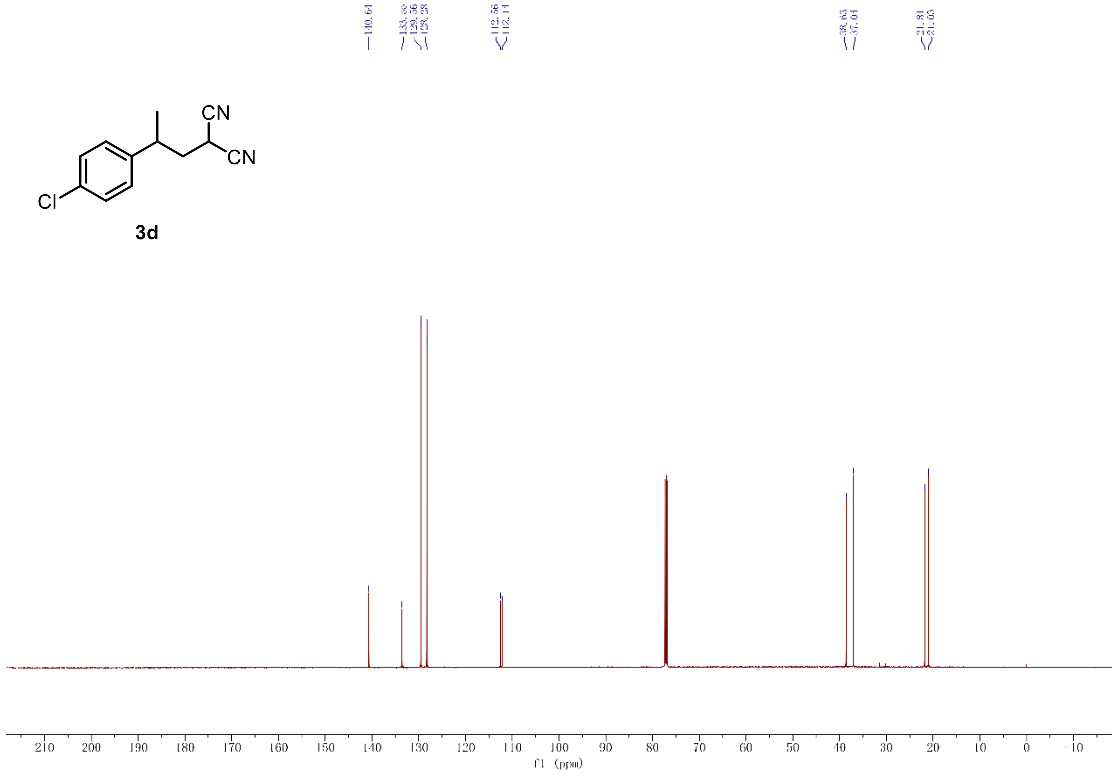

$^{13}$C-NMR of compound **3d** (151 MHz, $CDCl_3$)

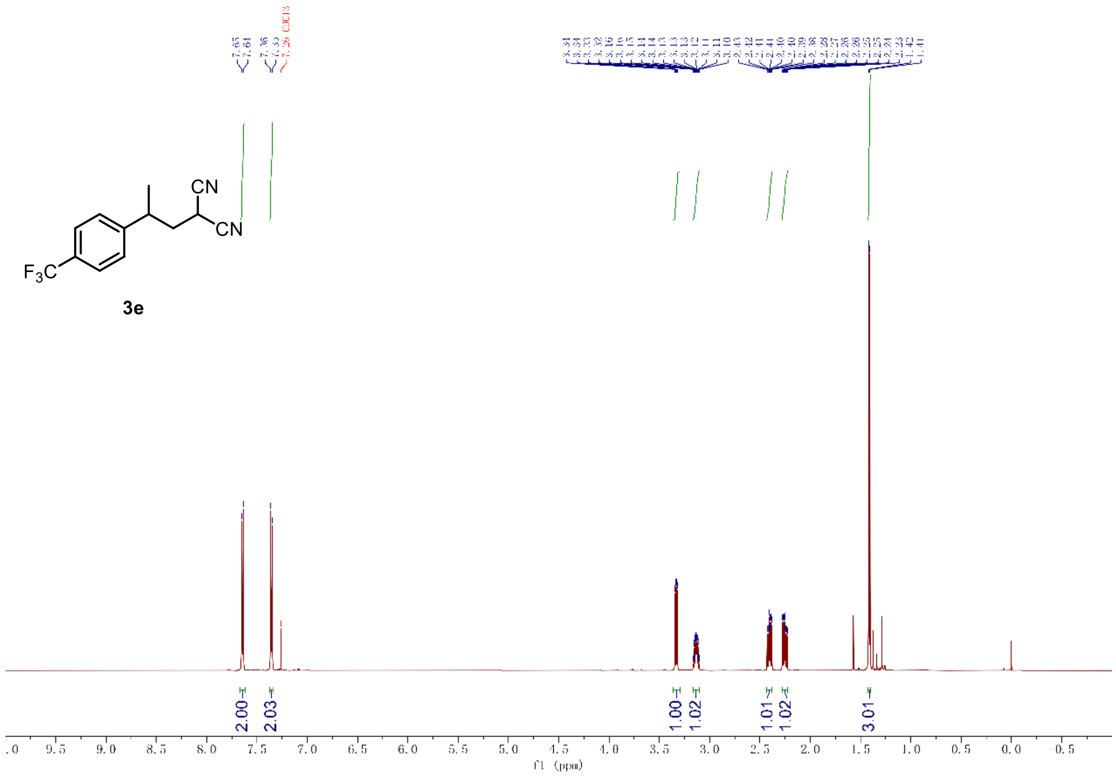

[1]H-NMR of compound **3e** (600 MHz, $CDCl_3$)

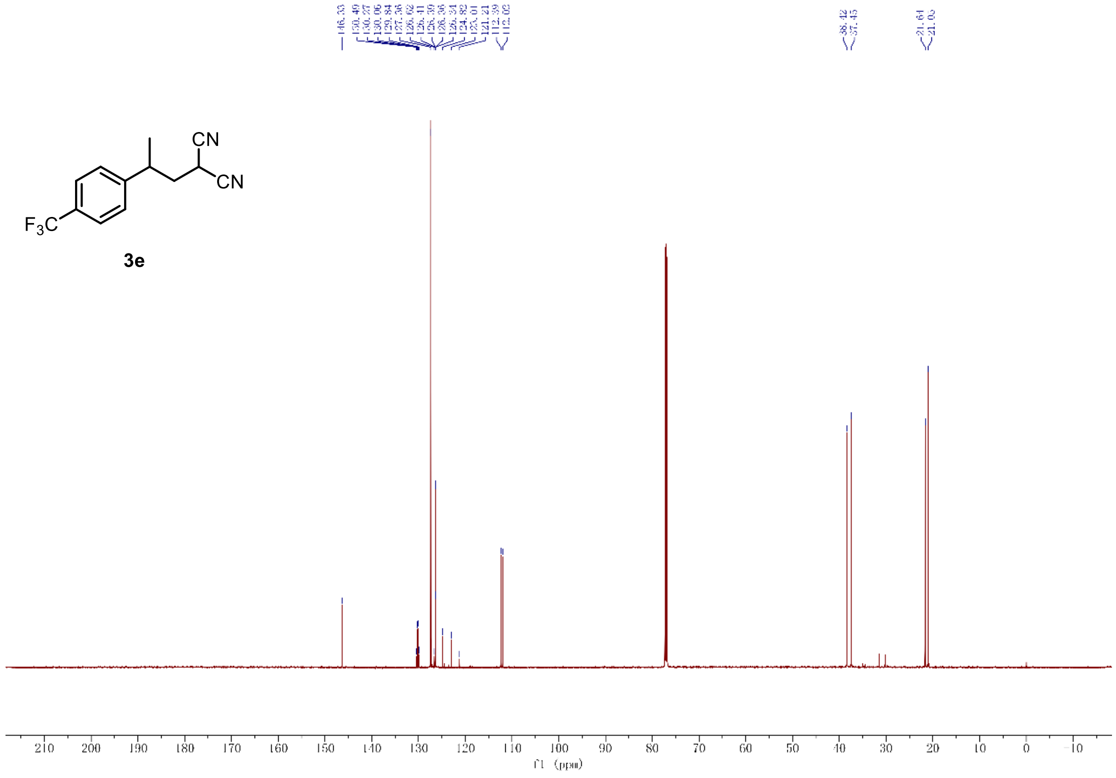

[13]C-NMR of compound **3e** (151 MHz, $CDCl_3$)

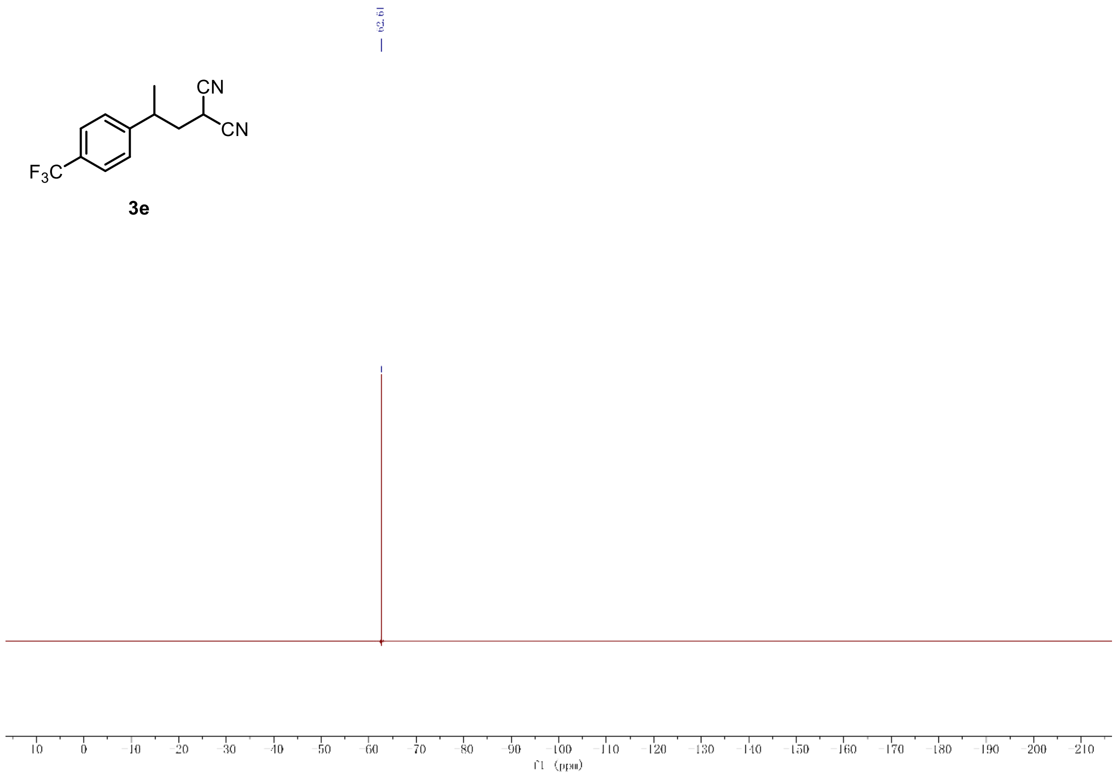

$^{19}$F-NMR of compound **3e** (565 MHz, $CDCl_3$)

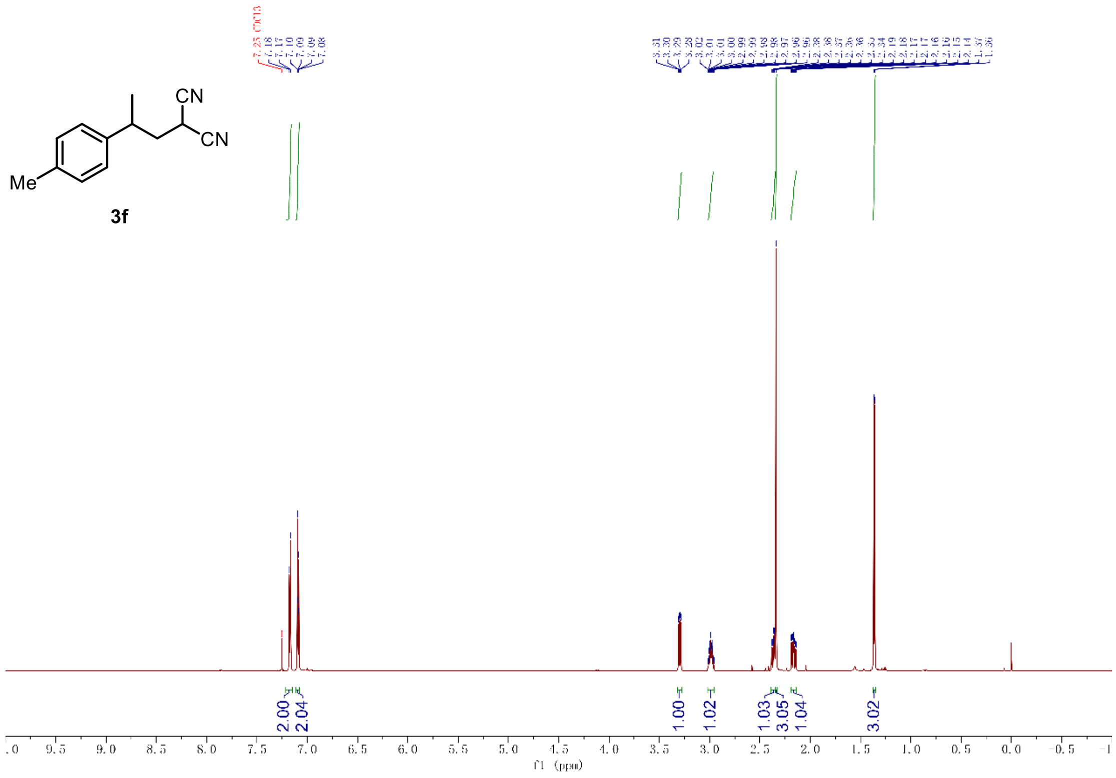

[1]H-NMR of compound **3f** (600 MHz, $CDCl_3$)

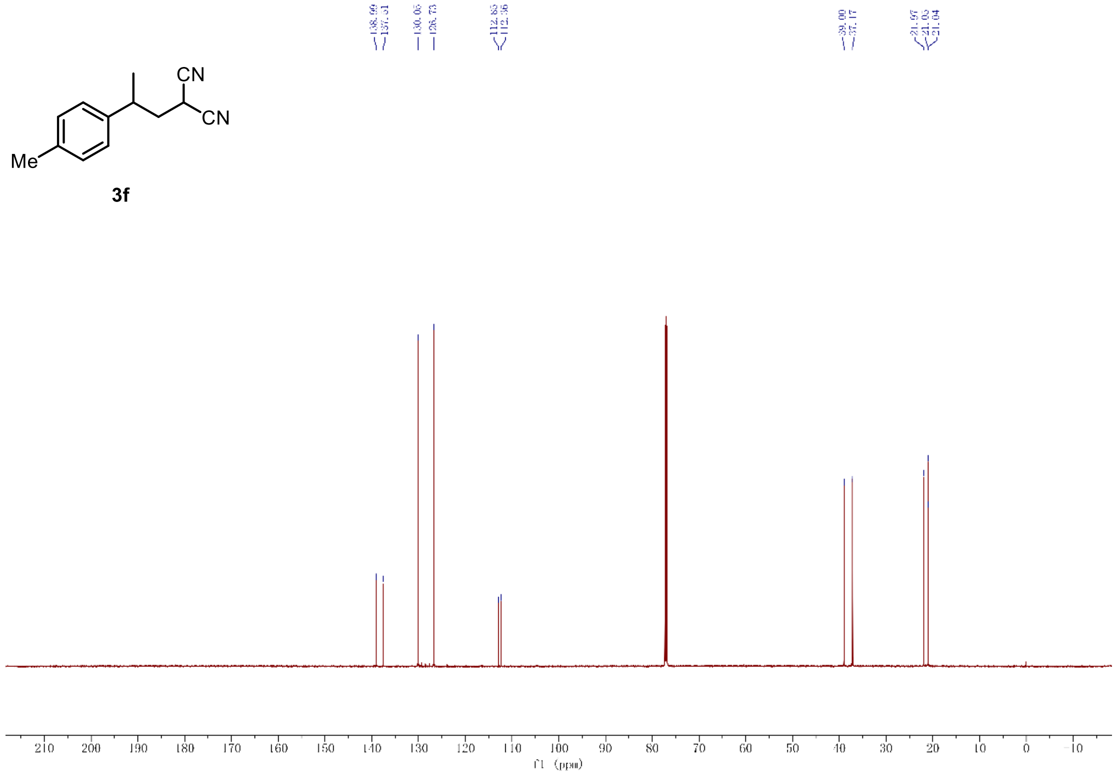

[13]C-NMR of compound **3f** (151 MHz, $CDCl_3$)

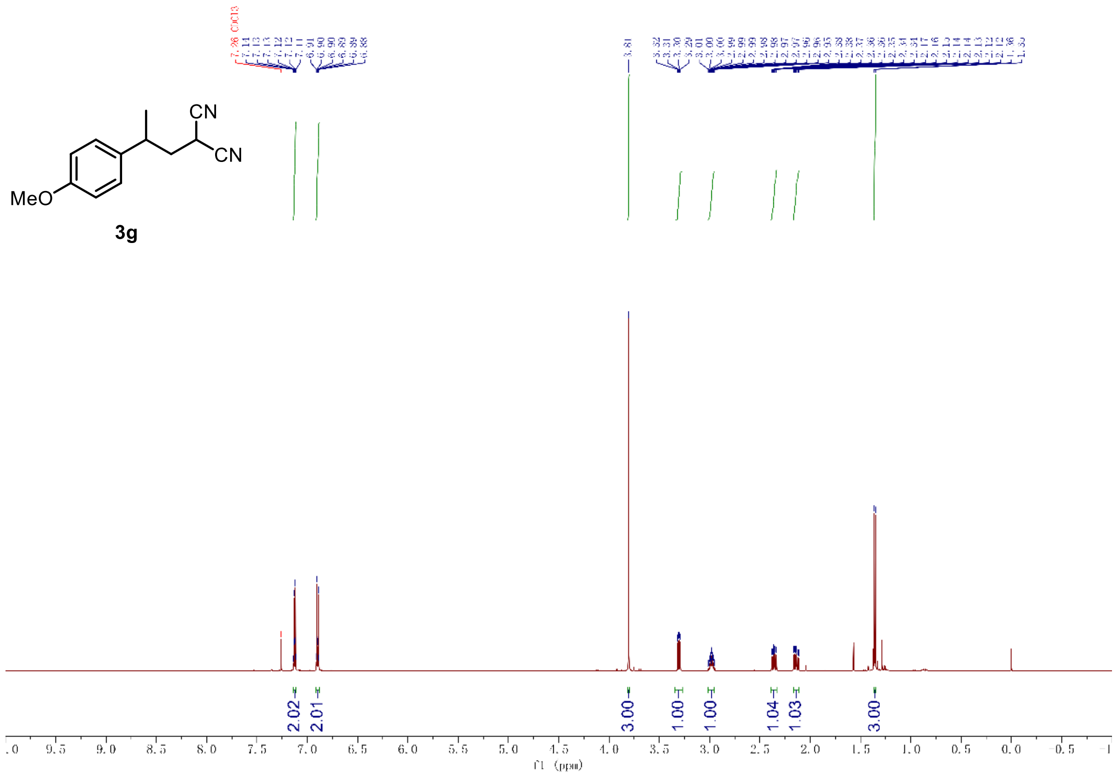

$^{1}$H-NMR of compound **3g** (600 MHz, $CDCl_3$)

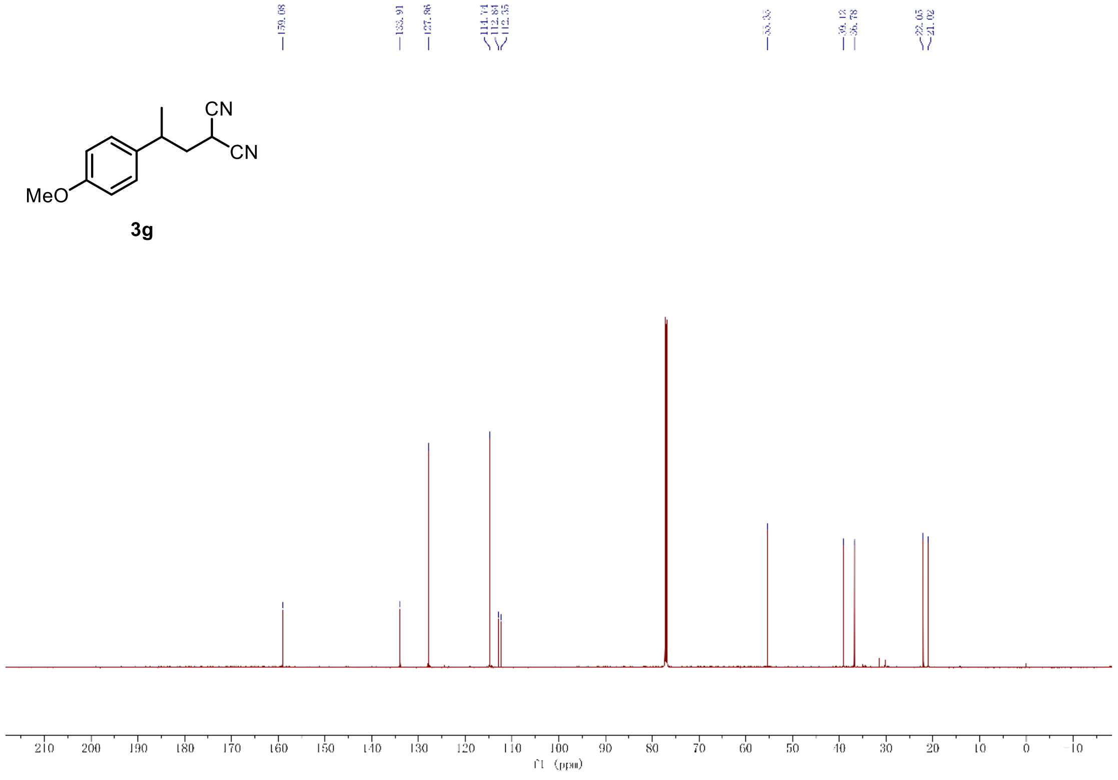

$^{13}$C-NMR of compound **3g** (151 MHz, $CDCl_3$)

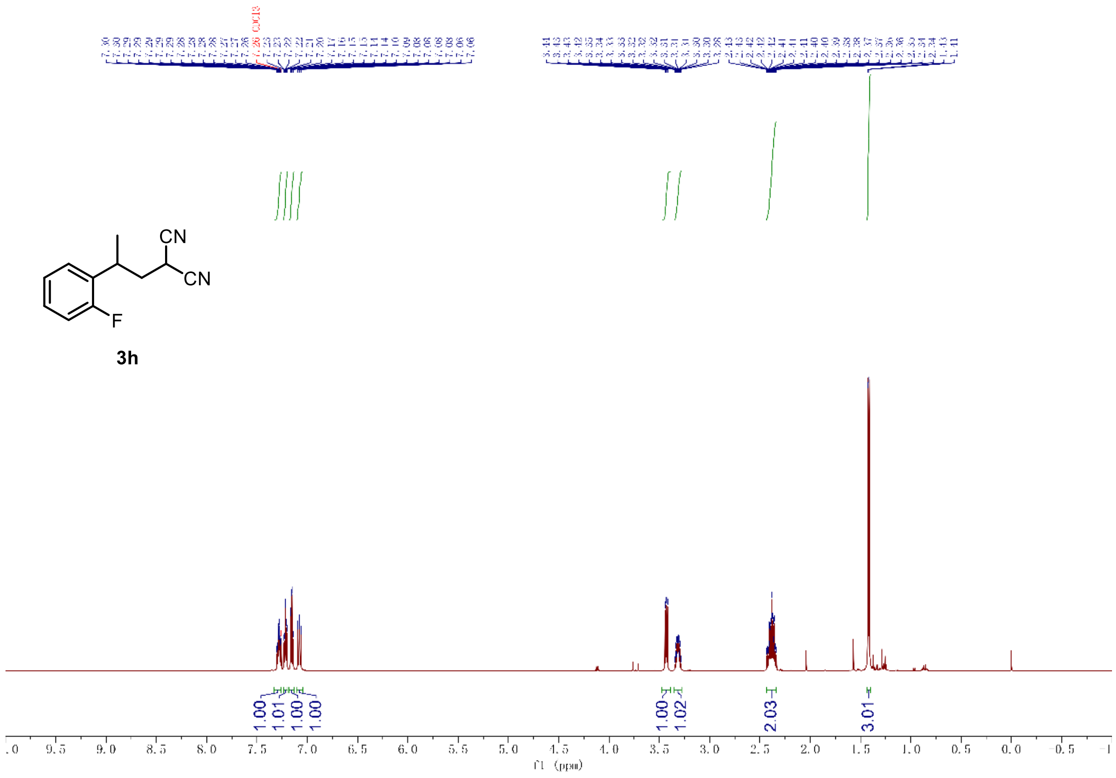

[1]H-NMR of compound **3h** (600 MHz, $CDCl_3$)

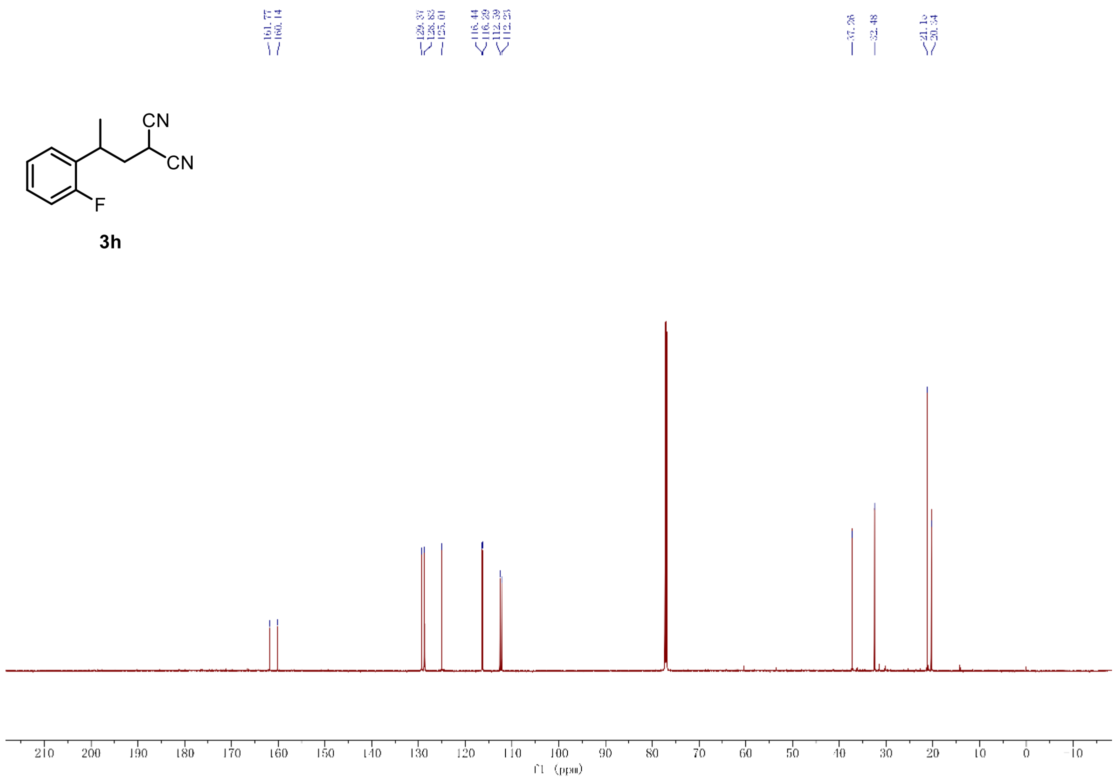

[13]C-NMR of compound **3h** (151 MHz, $CDCl_3$)

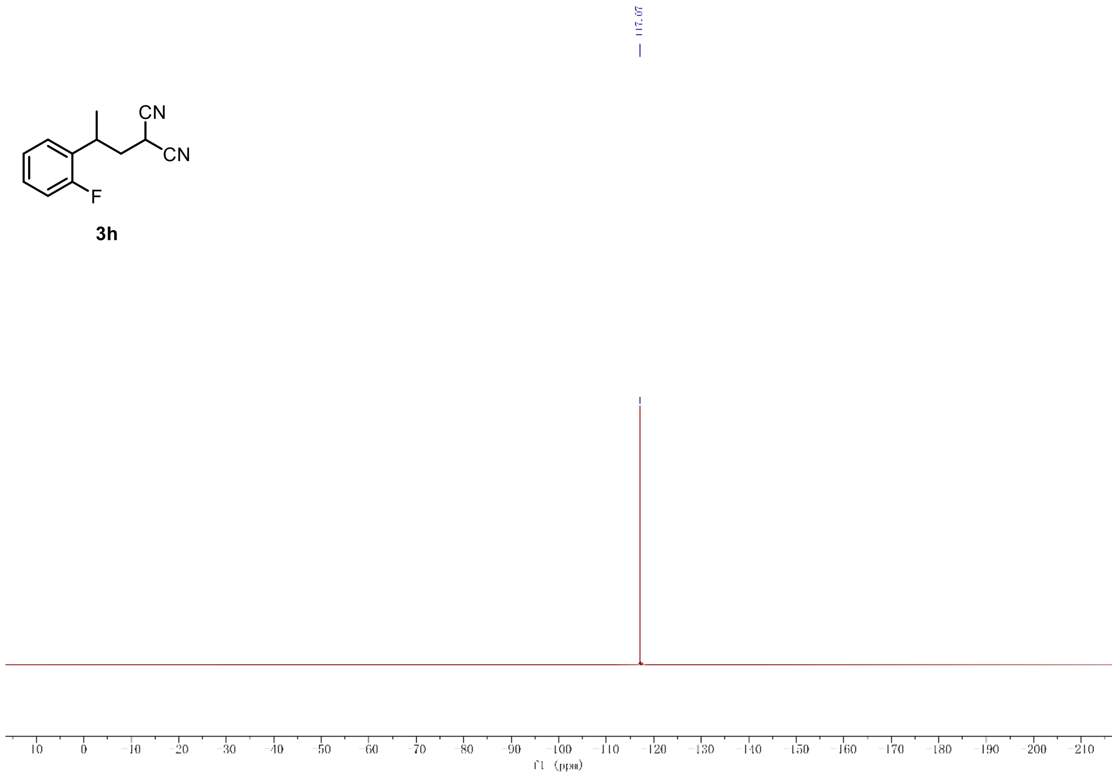

3h

$^{19}$F-NMR of compound **3h** (565 MHz, $CDCl_3$)

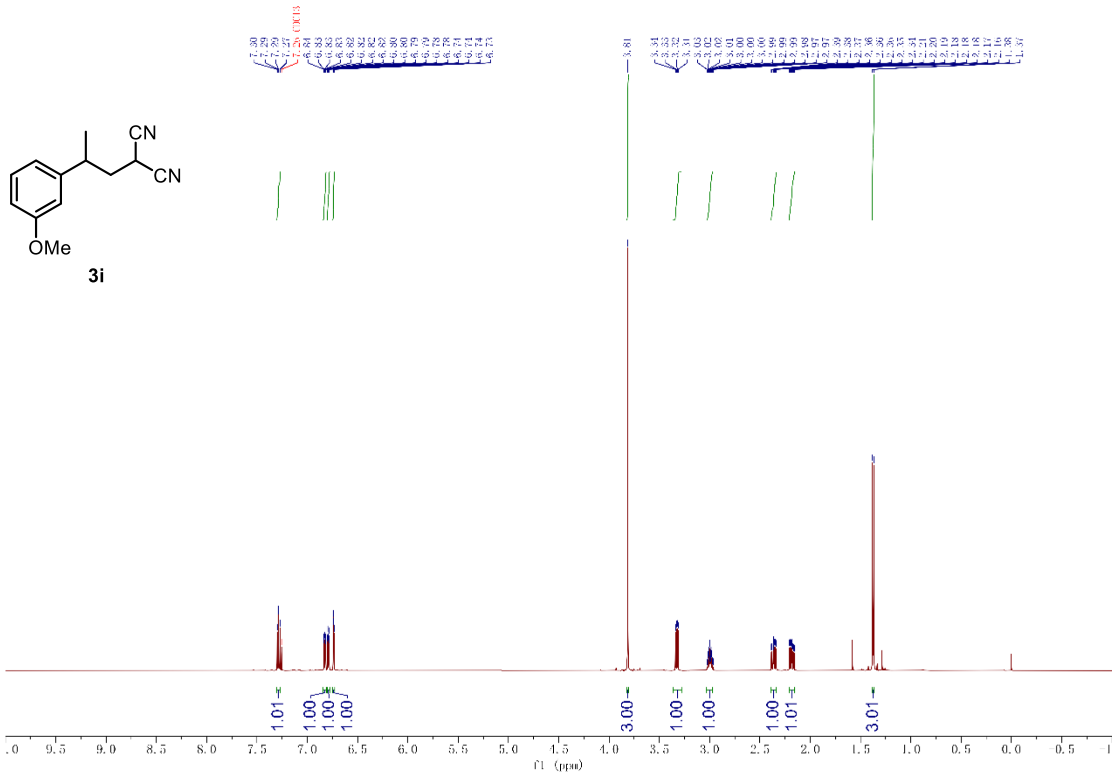

$^{1}$H-NMR of compound **3i** (600 MHz, $CDCl_3$)

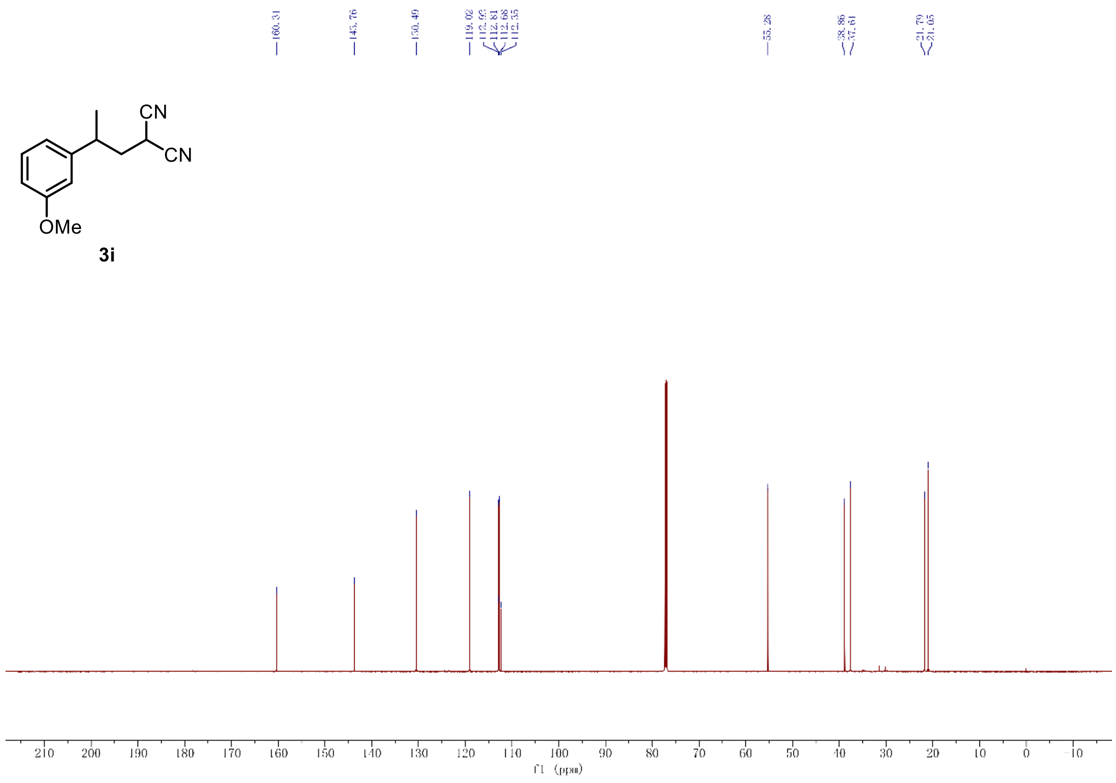

$^{13}$C-NMR of compound **3i** (151 MHz, $CDCl_3$)

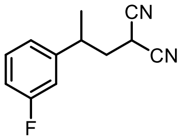

**3j**

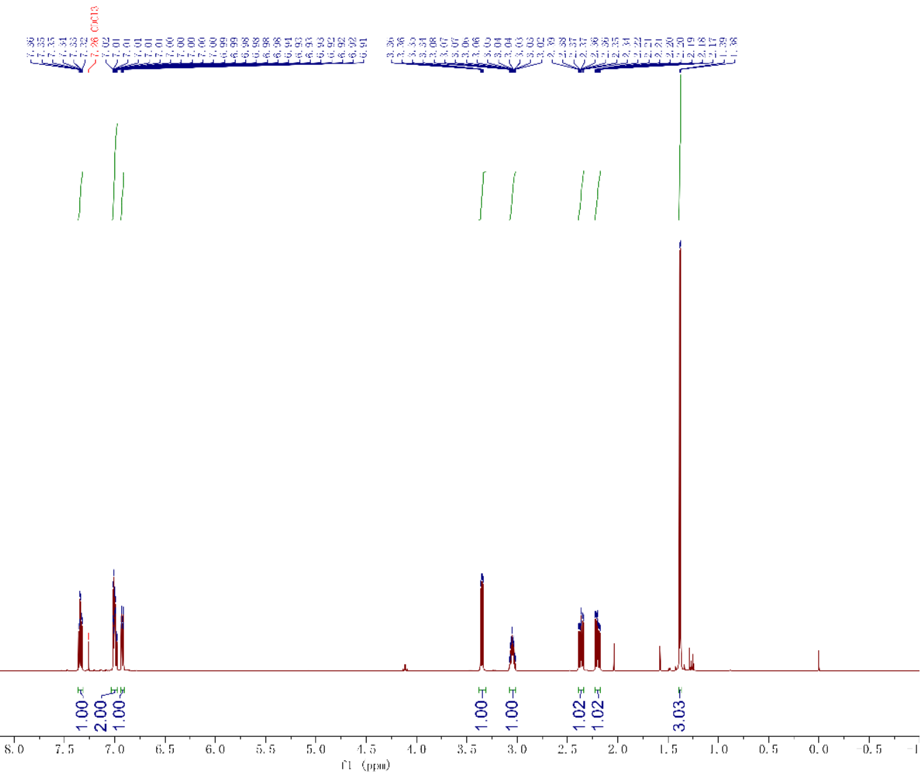

[1]H-NMR of compound **3j** (600 MHz, $CDCl_3$)

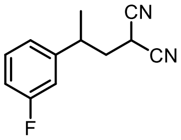

**3j**

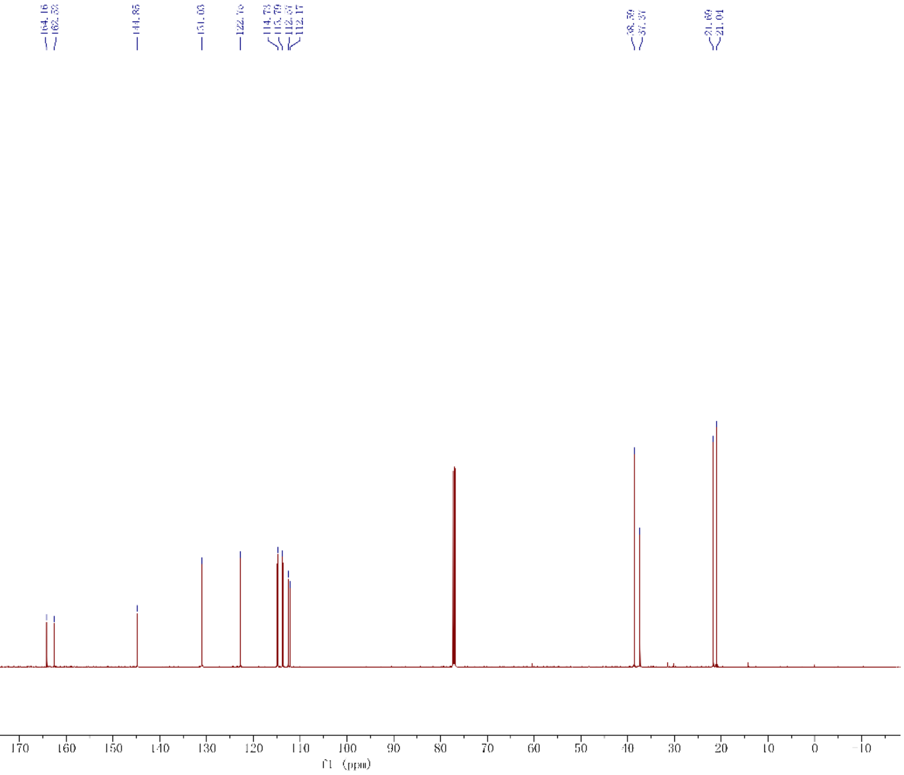

[13]C-NMR of compound **3j** (151 MHz, $CDCl_3$)

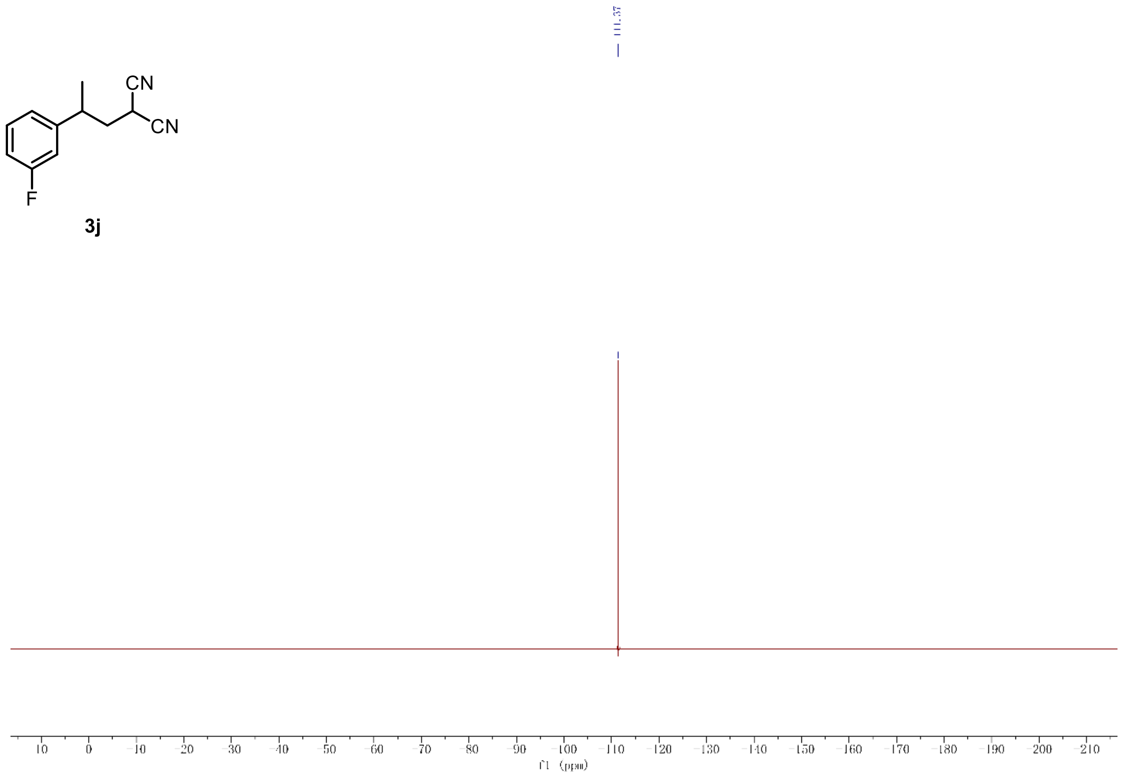

[19]F-NMR of compound **3j** (565 MHz, $CDCl_3$)

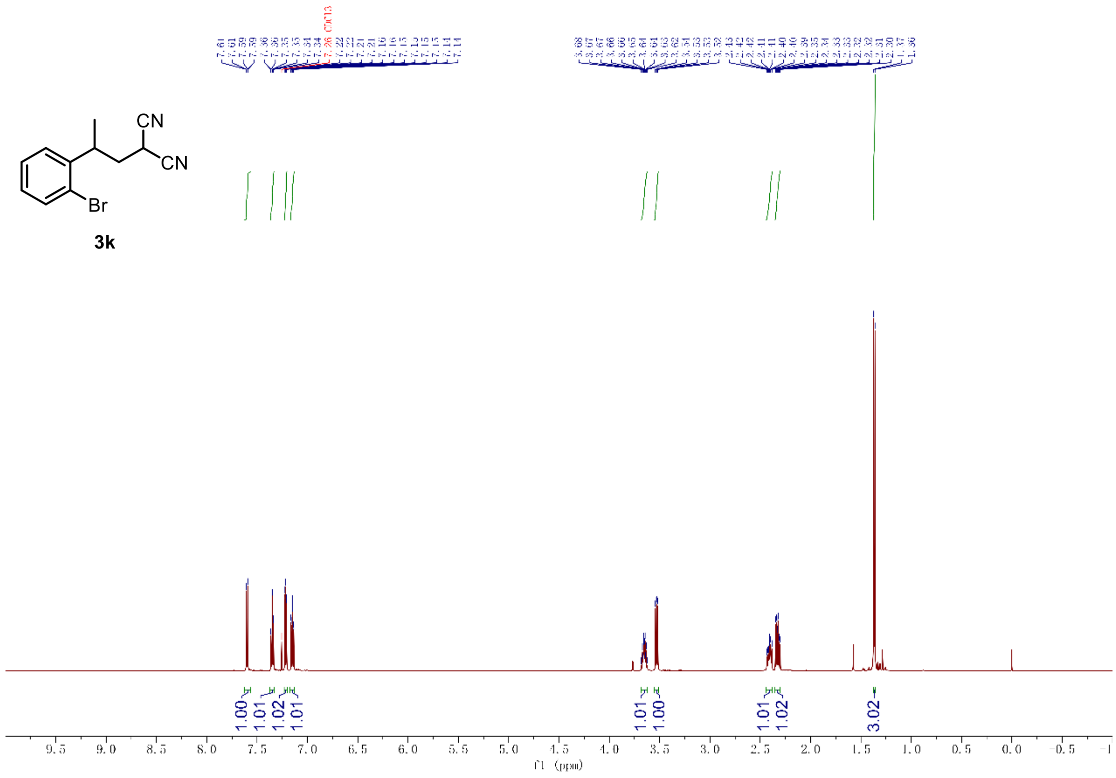

$^{1}$H-NMR of compound **3k** (600 MHz, $CDCl_3$)

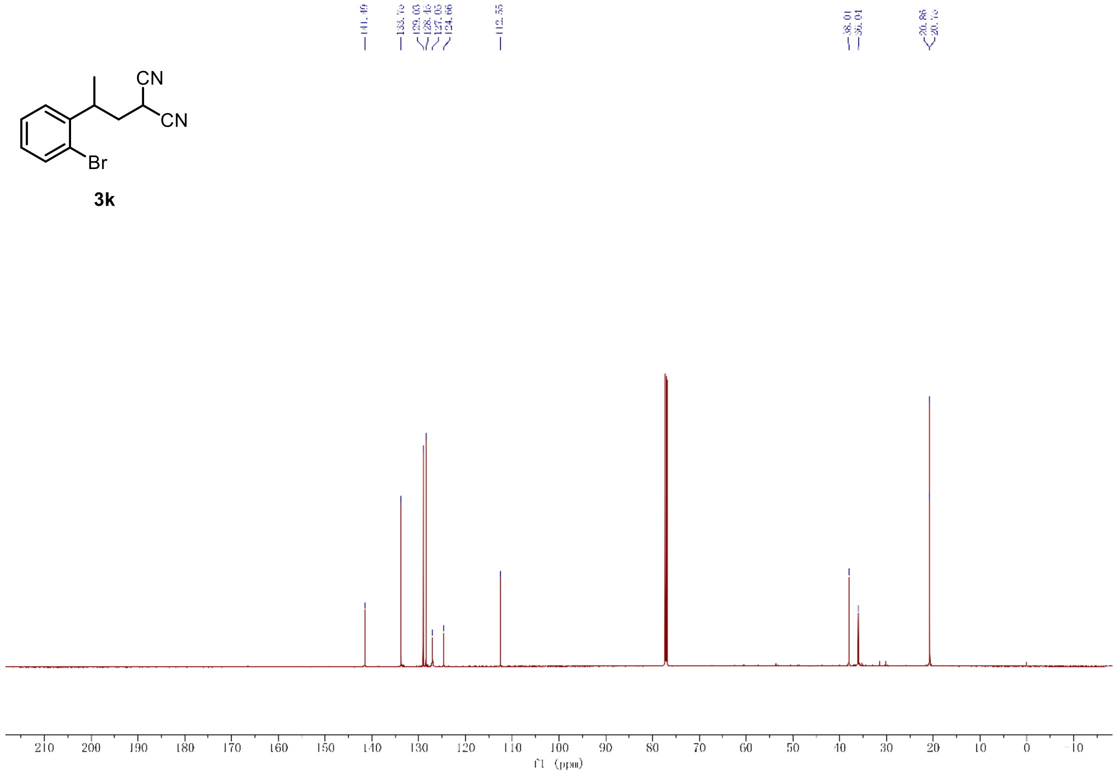

$^{13}$C-NMR of compound **3k** (151 MHz, $CDCl_3$)

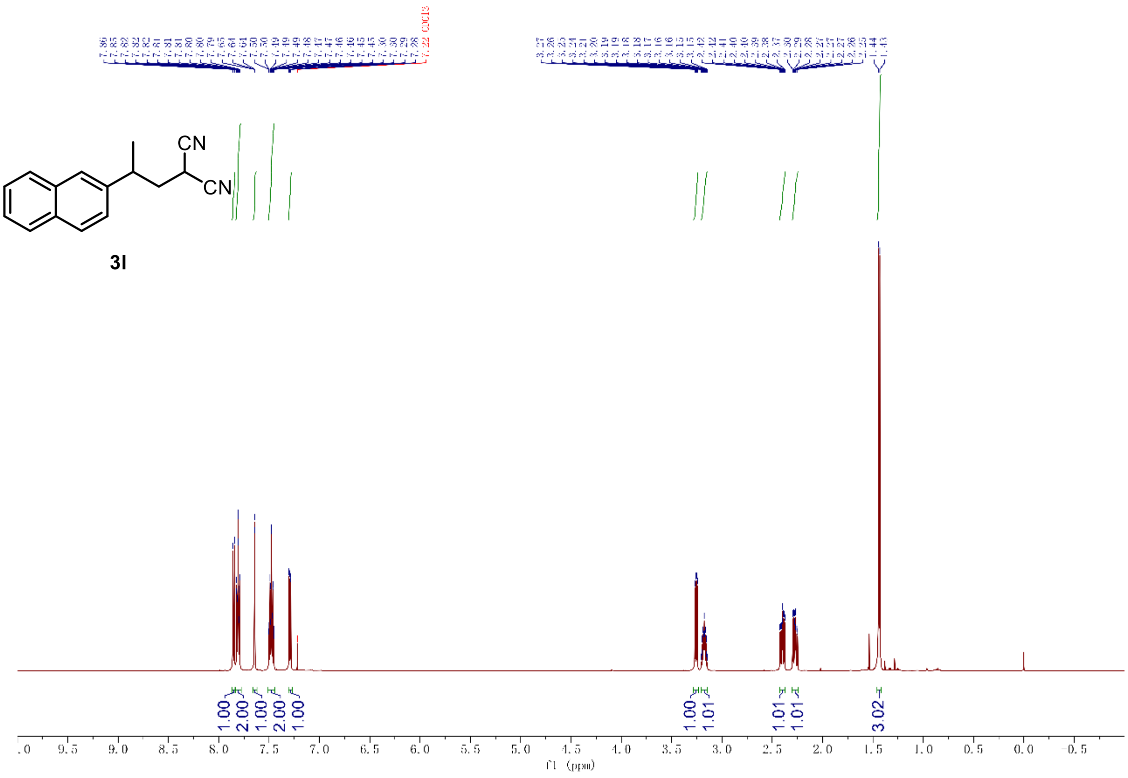

$^{1}$H-NMR of compound **3l** (600 MHz, $CDCl_3$)

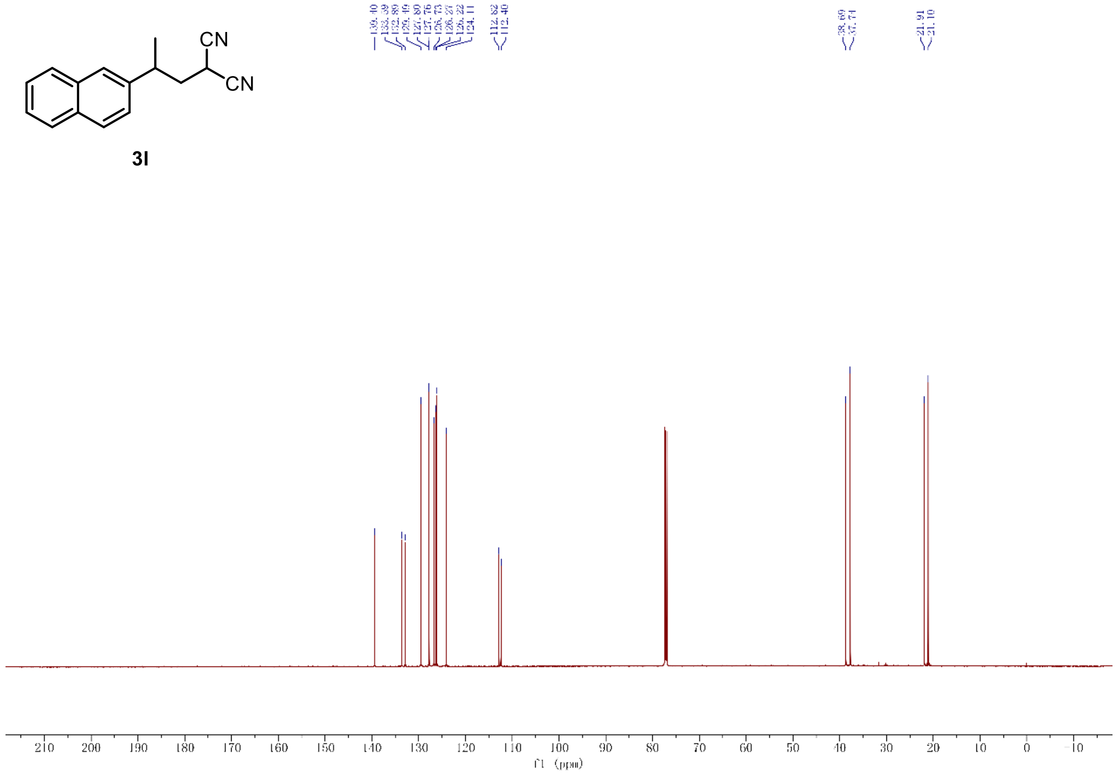

$^{13}$C-NMR of compound **3l** (151 MHz, $CDCl_3$)

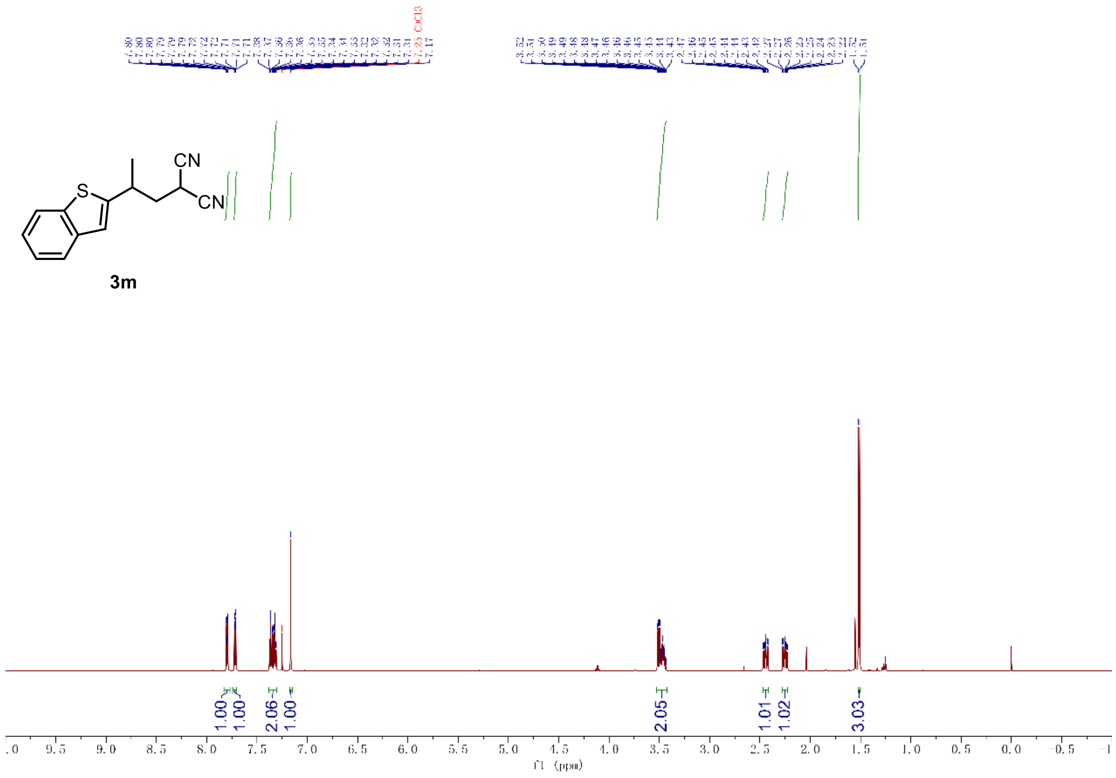

$^{1}$H-NMR of compound **3m** (600 MHz, $CDCl_3$)

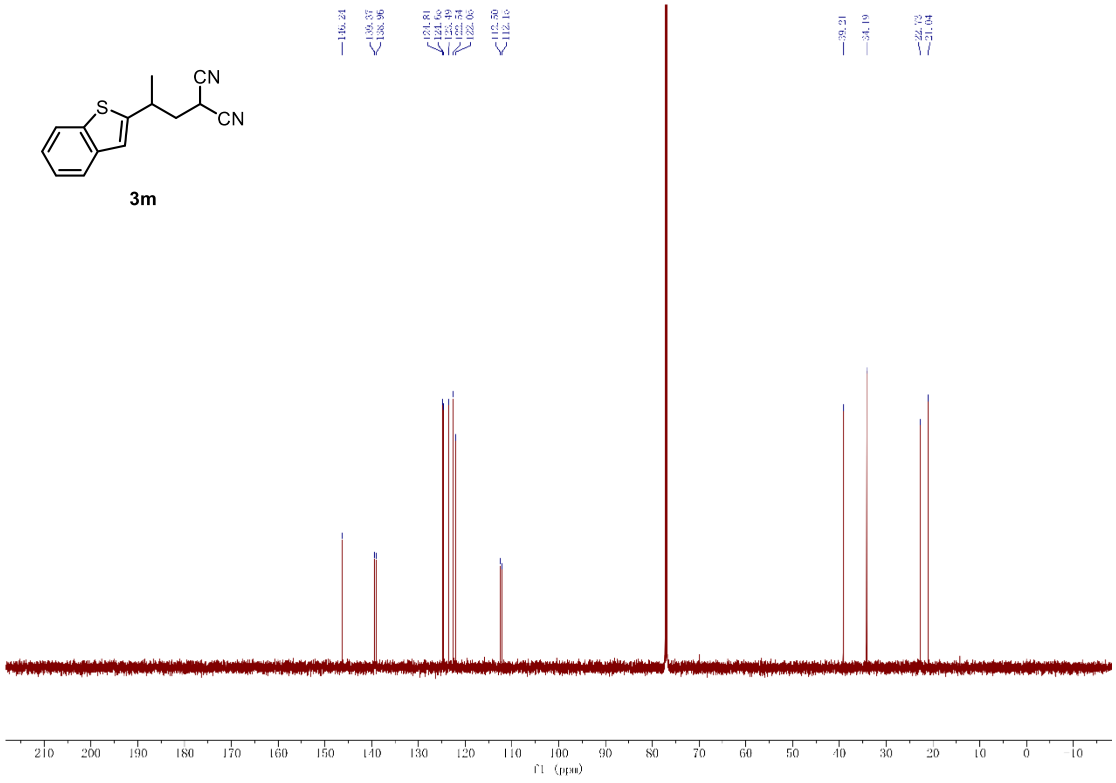

$^{13}$C-NMR of compound **3m** (151 MHz, $CDCl_3$)

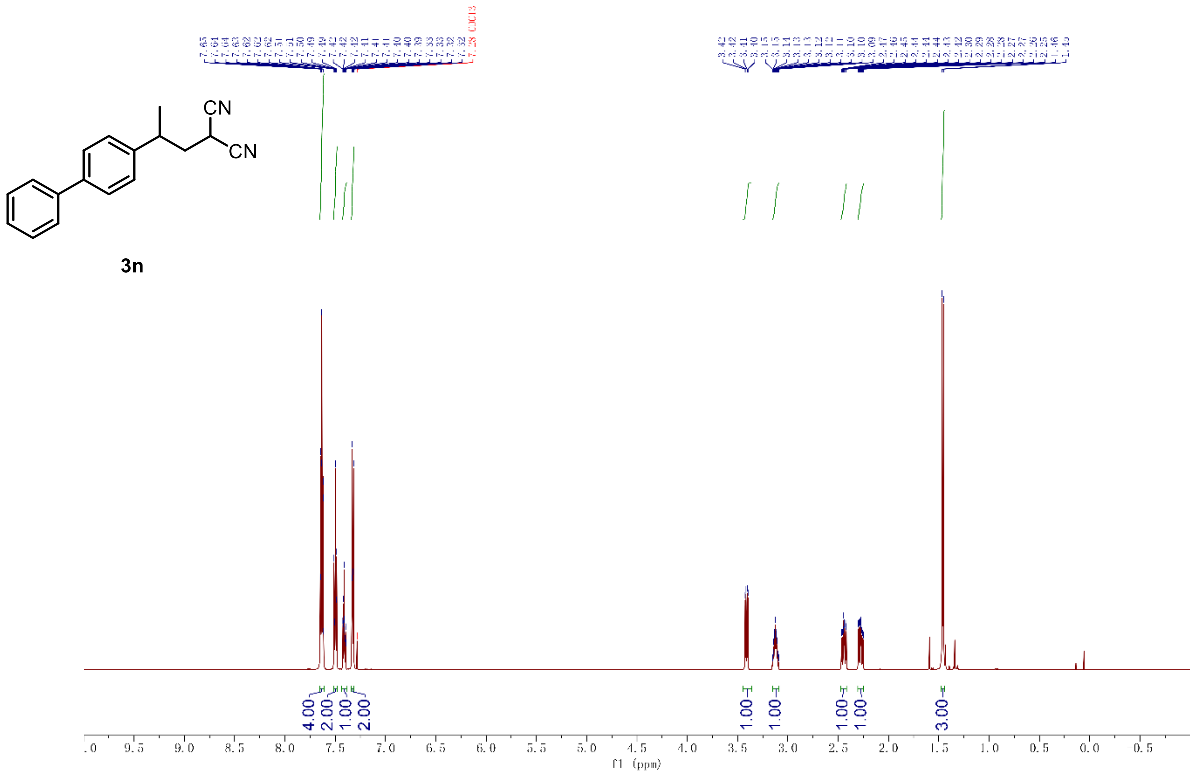

$^{1}$H-NMR of compound **3n** (600 MHz, $CDCl_3$)

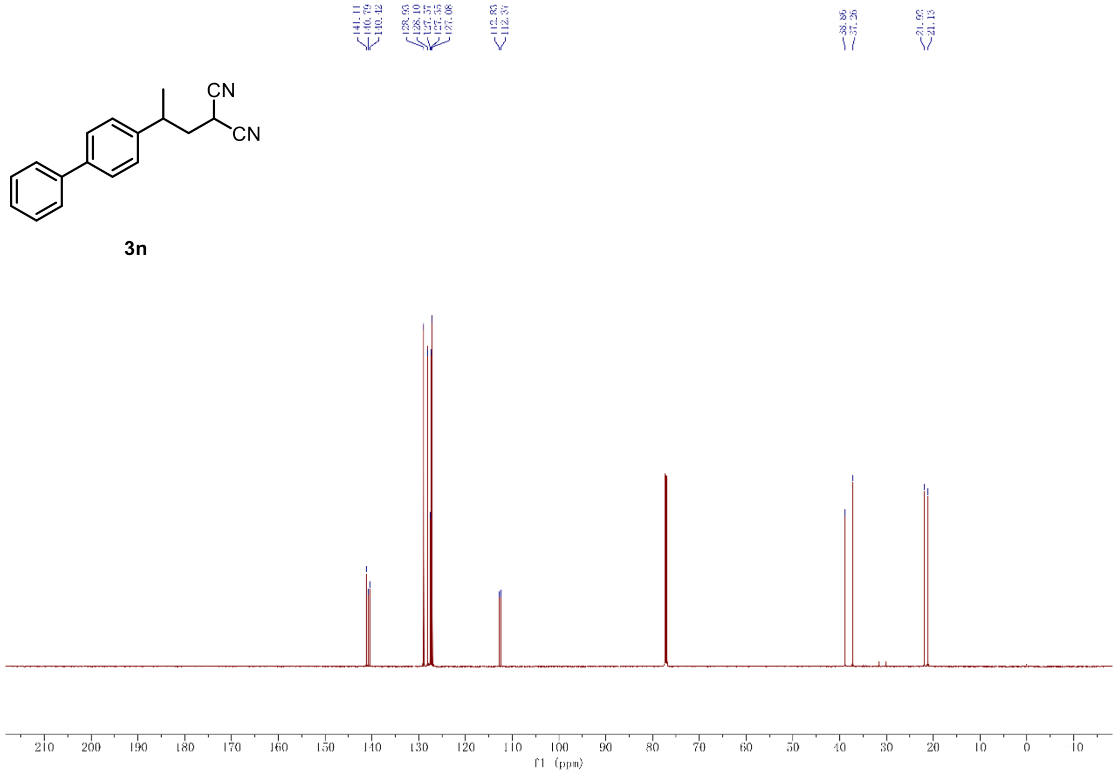

$^{13}$C-NMR of compound **3n** (151 MHz, $CDCl_3$)

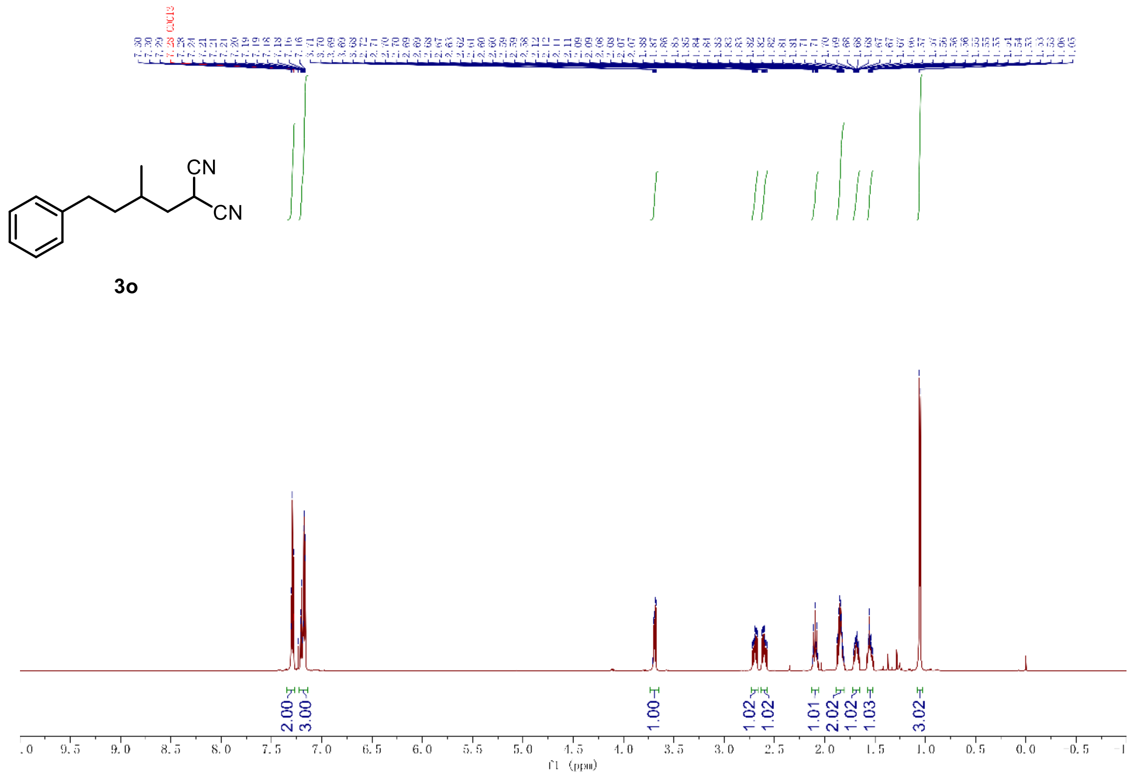

[1]H-NMR of compound **3o** (600 MHz, $CDCl_3$)

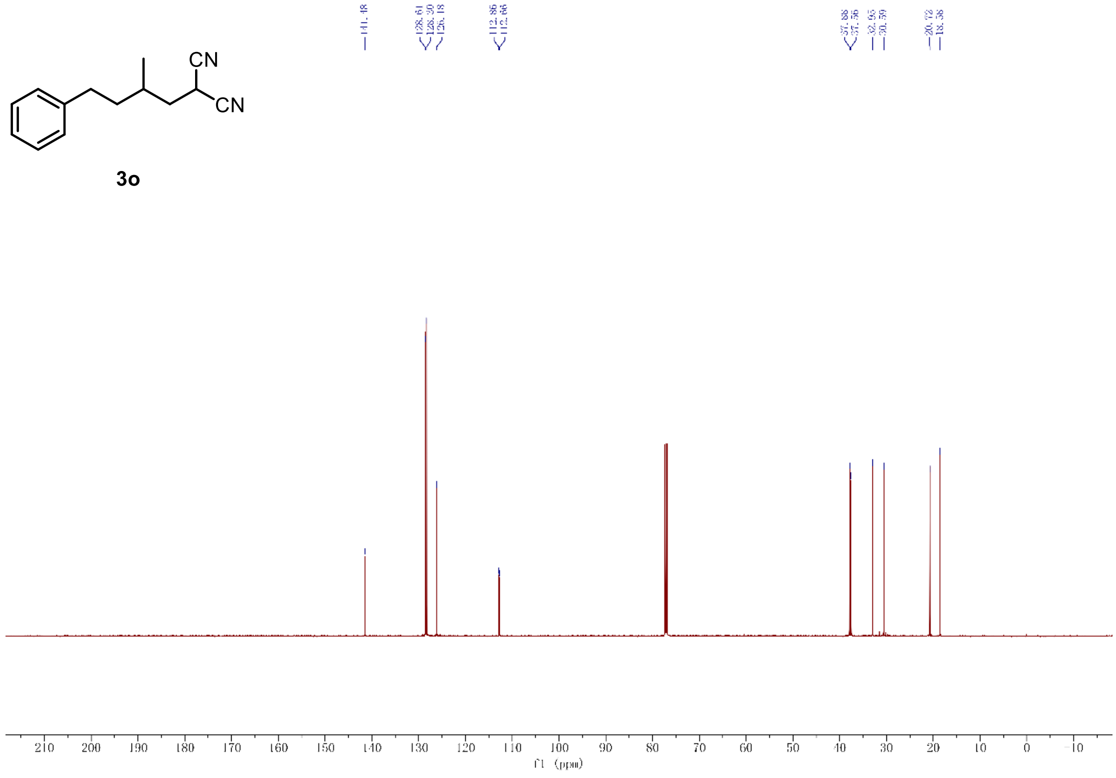

[13]C-NMR of compound **3o** (151 MHz, $CDCl_3$)

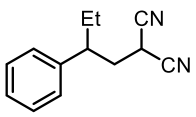

3p

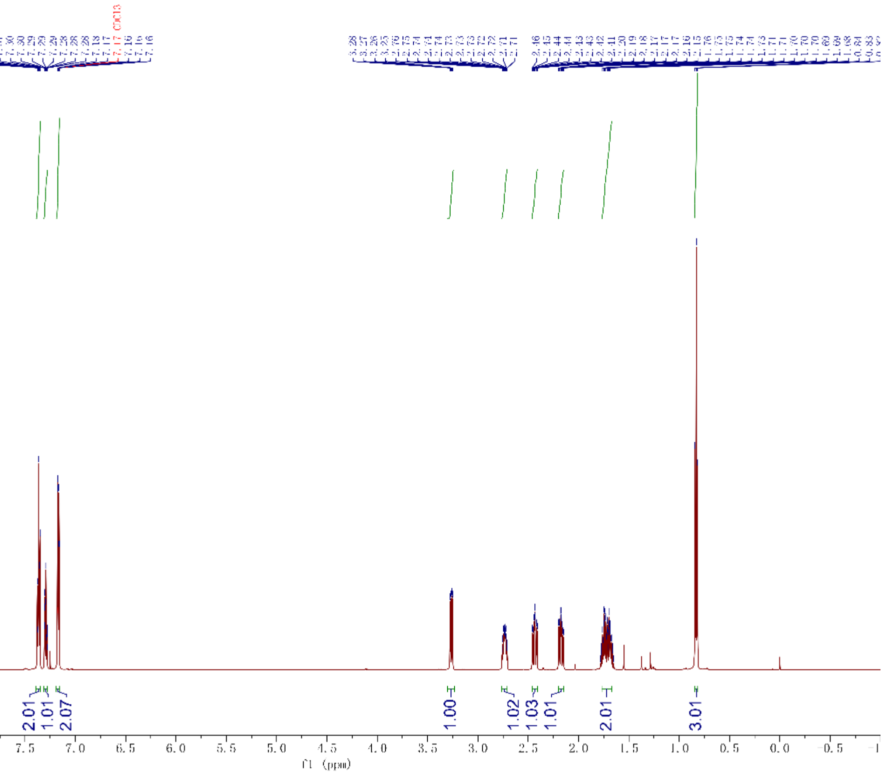

[1]H-NMR of compound **3p** (600 MHz, $CDCl_3$)

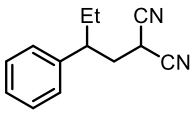

3p

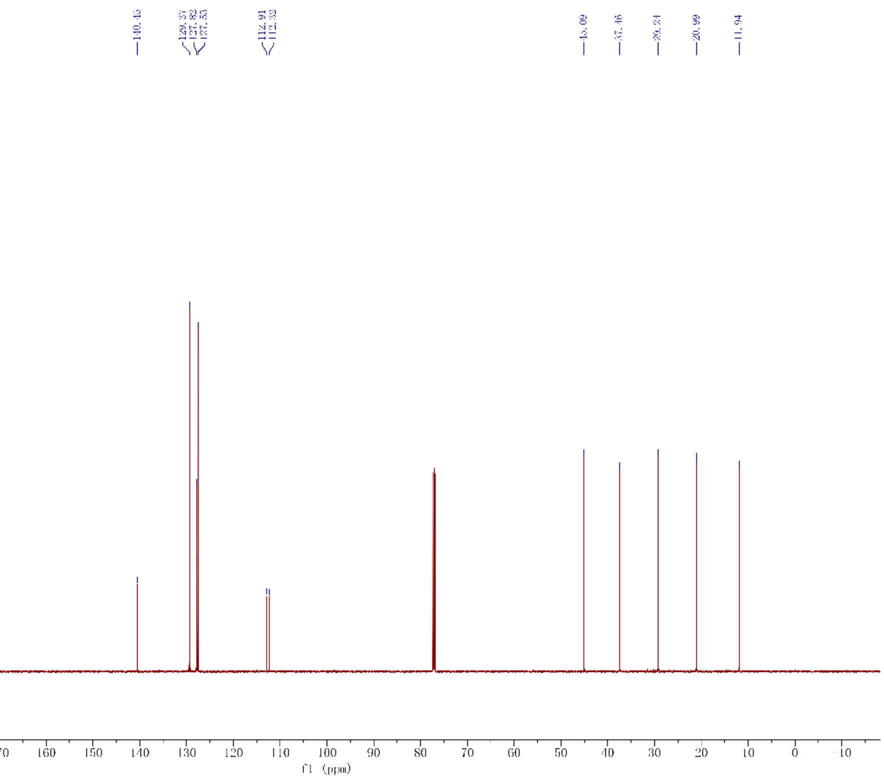

[13]C-NMR of compound **3p** (151 MHz, $CDCl_3$)

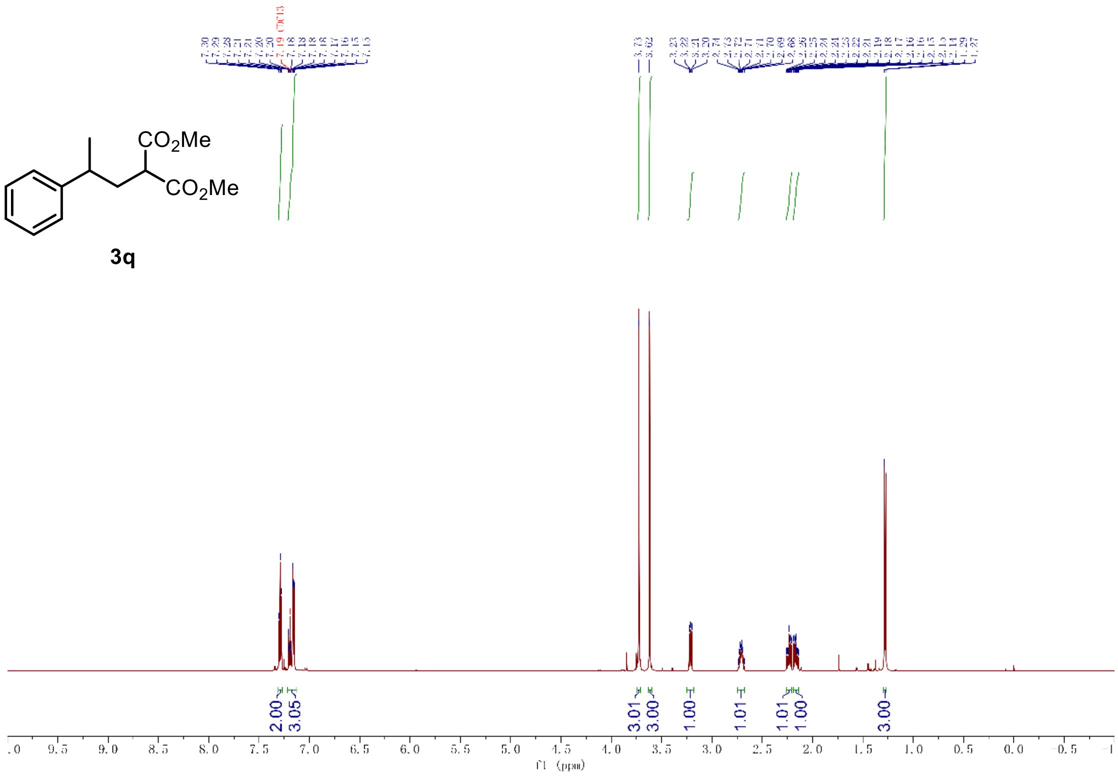

[1]H-NMR of compound **3q** (600 MHz, $CDCl_3$)

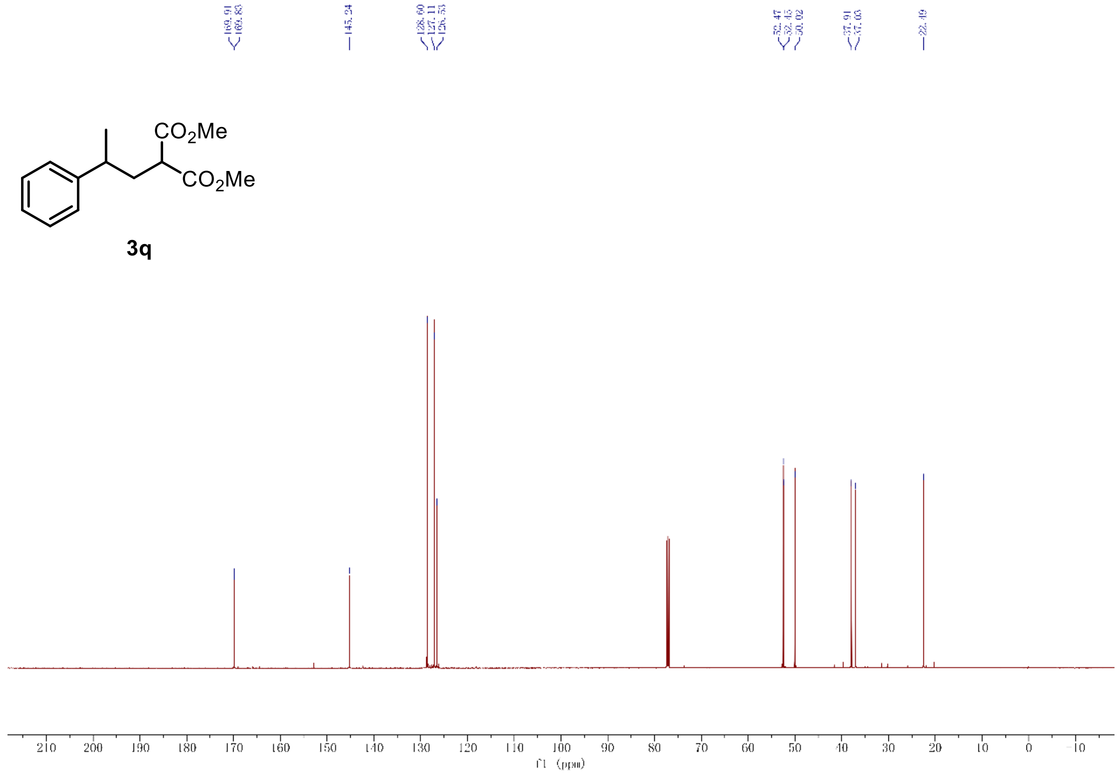

[13]C-NMR of compound **3q** (151 MHz, $CDCl_3$)

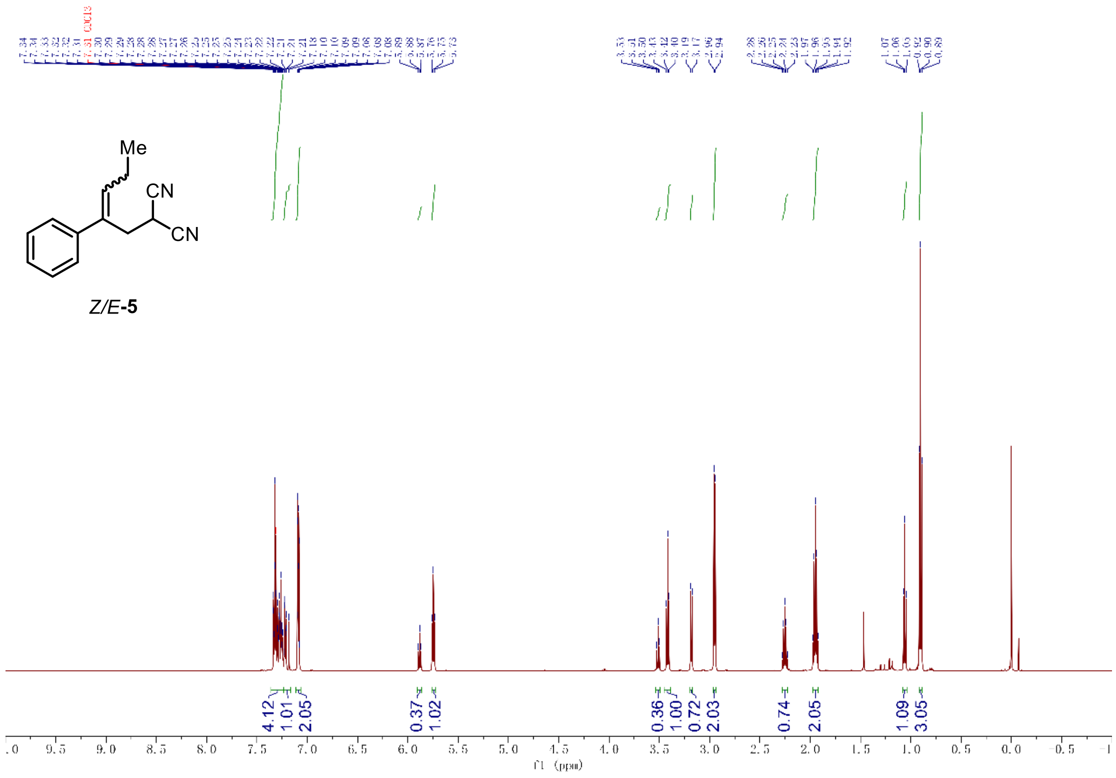

[1]H-NMR of compound **5** (600 MHz, $CDCl_3$)

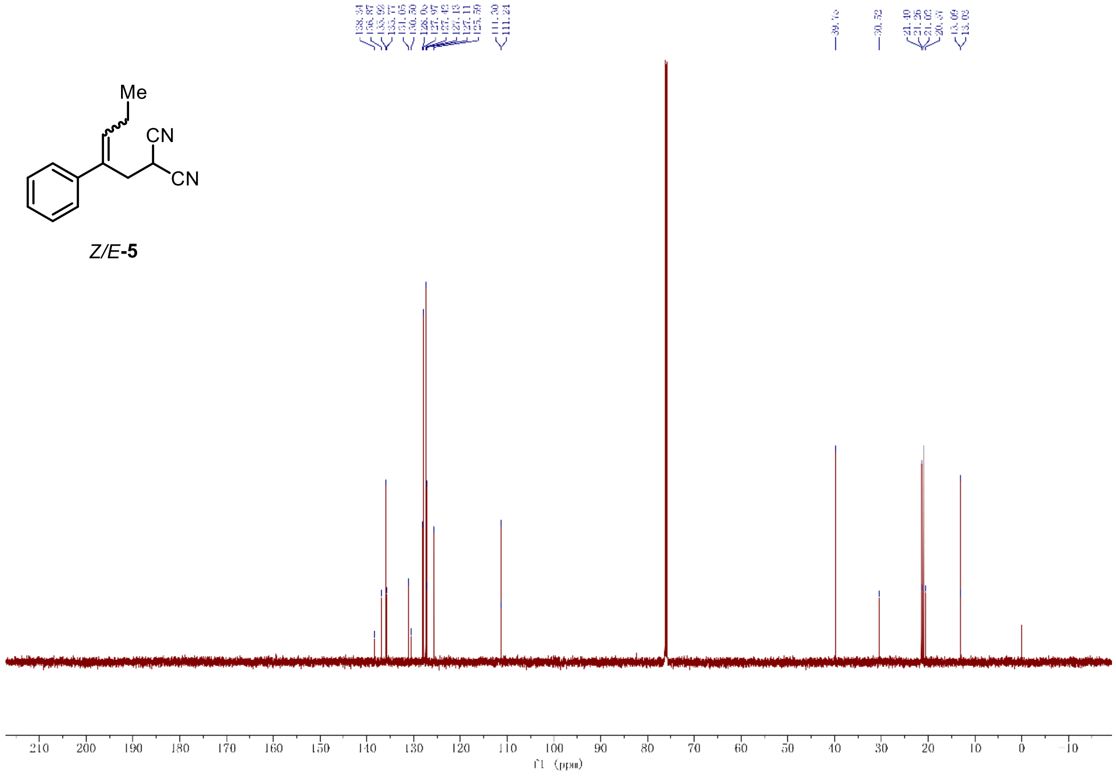

[13]C-NMR of compound **5** (151 MHz, $CDCl_3$)
